# The nail mesenchyme creates a regeneration-specific ligand environment that orchestrates mammalian regeneration versus fibrotic wound healing

**DOI:** 10.64898/2026.04.18.719303

**Authors:** Sruthi Purushothaman, Sandeep Saxena, Christine Eisner, Konstantina Karamboulas, David R. Kaplan, Fabio M.V. Rossi, Freda D. Miller

## Abstract

The adult digit tip is one of a few privileged mammalian tissues that regenerate. Here, we asked why this is so. Using single cell spatial transcriptomics and genetic mouse models we show regeneration requires the nail organ and associated nail mesenchyme, and that in their absence fibrotic wound-healing occurs. We show the nail organ/mesenchyme orchestrates a highly-organized regenerative response that includes reprogramming of tissue-resident mesenchymal cells to a blastema state. It does this by creating a regeneration-specific ligand environment that includes multiple BMPs. Loss of the resultant downstream BMP signaling by genetic deletion of *Smad4* in all mesenchymal cells or specifically in the nail mesenchyme inhibits multiple aspects of the regenerative response, including expression of regeneration-specific ligands. Thus, the nail mesenchyme creates a BMP-rich local ligand environment that reprograms mesenchymal cells and orchestrate tissues regeneration versus fibrotic wound-healing, findings with implications for tissue repair and anti-fibrosis strategies.

## Introduction

A long-standing biological question is why mammals have largely lost the ability to regenerate. Potential answers come from studying the few privileged locations such as the digit tip where mammalian multi-tissue regeneration does occur^1–6^. In adult mice and humans, if the digit tip is amputated sparing the nailbed, then regenerative growth occurs and faithfully restores the lost tissue. This regeneration requires formation of a blastema largely comprised of local *Pdgfra*-positive tissue-resident mesenchymal cells that have lost their differentiated phenotype and acquired a transcriptionally-distinct blastema state^7–9^. The reprogrammed blastema cells then rebuild lost tissues such as the bone and dermis. Notably, axolotl limb regeneration also requires a blastema that forms by dedifferentiation of mesenchymal cells to an embryonic state^10,11^. However, adult mouse blastema cells express genes associated with both development and adult tissue repair^8,9^, indicating this is not classic dedifferentiation but is more akin to reprogramming.

What is special about the injured digit tip that promotes regeneration rather than wound-healing? An important clue is that regeneration does not occur when the nail organ is removed^12,13^. The nail organ is a specialized skin appendage that, like hair follicles, is comprised of epithelial cells and an inductive mesenchyme^1,14–16^. This nail mesenchyme instructs epithelial cells to build the new nail tissue throughout life and thus can be viewed as a developmental signaling center that persists. Notably, several studies show that when this signaling center is perturbed by even partial loss of the nail mesenchyme, digit tip regeneration is abnormal^12,17,18^. While it is unclear how the nail organ/mesenchyme promotes regeneration, transplant studies indicate it may so by modulating the environment. For example, when dermal fibroblasts were transplanted into the regenerating digit tip they contributed to blastema formation and bone regeneration, but when transplanted into the amputated digit tip lacking the nail organ they instead participated in wound-healing^8^. If we could decipher this pro-regenerative environment then perhaps this would inform strategies for promoting tissue repair and/or inhibiting fibrosis, the latter a particularly widespread clinical problem^19–23^.

Here, we asked if and how the nail organ/mesenchyme creates a pro-regenerative environment. Using single cell spatial transcriptomics and genetic mouse models, we demonstrate that the nail organ and more specifically the nail mesenchyme orchestrate a multicellular regenerative response that includes reprogramming mesenchymal cells to a blastema state. The nail organ/mesenchyme does this, at least in part, by creating a regeneration-specific ligand environment that includes multiple BMP family members. When signaling downstream of this BMP-rich environment is blocked by acute loss of the essential downstream BMP signaling protein SMAD4 in all mesenchymal cells or specifically in the nail mesenchyme, this inhibits multiple aspects of the regenerative response. Thus, the nail mesenchyme creates a regeneration-specific BMP-rich local environment that reprograms mesenchymal cells to a blastema state and in so doing orchestrates tissue regeneration versus fibrotic wound-healing.

## Results

### In the absence of the nail, digit tip mesenchymal cells undergo fibrotic wound healing rather than regeneration

We and others previously showed that when the adult mouse digit tip is amputated proximal to the nail, regeneration does not occur (Fig. S1A)^1,8,12,24^. To better understand this finding, we compared the digit tip response 14 days following nonregenerative amputations in the absence of the nail organ versus regenerative amputations that spare the nail bed (Fig. S1A). Initially, we combined lineage tracing with lightsheet microscopy to analyze unsectioned digit tips where either *Pdgfra*-positive mesenchymal cells or *Cdh5*-positive endothelial vasculature were tagged with a TdTomato (TdT) reporter. For mesenchymal cells, we used a well-characterized mouse line that carries CreERT2 knocked-in to the *Pdgfra* locus^25^, and a floxed *Tdt* transgene knocked-in to the *Rosa26* locus (*Pdgfra-Tdt* mice). We tamoxifen-treated these mice for 5 days to acutely label *Pdgfra*-positive digit tip mesenchymal cells, as previously reported^8,18^. For endothelial cells, we used a mouse line that carries the floxed *Tdt* allele and a constitutive *Cdh5Cre* transgene (*Cdh5Cre-Tdt* mice)^26^. In both cases, we performed amputations and at 14 DPA used tissue clearing and lightsheet microscopy to visualize the entire digit tip, analyzing 6-8 digits from 4 different mice.

These analyses (Fig. 1A, B, Video S1-4) confirmed that the regenerative and non-regenerative digit tip responses were very different. In the regenerative condition, there was robust organized tissue growth distal to the amputation site, as previously reported^8,18,24^. The regenerative tissue was largely comprised of *Pdgfra*-TdT-positive mesenchymal cells surrounded by an epithelium, with *Cdh5*-positive blood vessels in and around the new tissue (Fig. 1A, B, Video S1 and S3). In the non-regenerative condition, there was instead a thin vascularized cap of tissue between the bone and epidermis that was largely comprised of *Pdgfra*-TdT-positive mesenchymal cells (Fig. 1A, B, Video S2 and S4). However, there was much less distal growth, the new tissue was disorganized in terms of overt morphology and the vasculature was relatively dense within the tissue cap over the bone.

**Figure 1.**
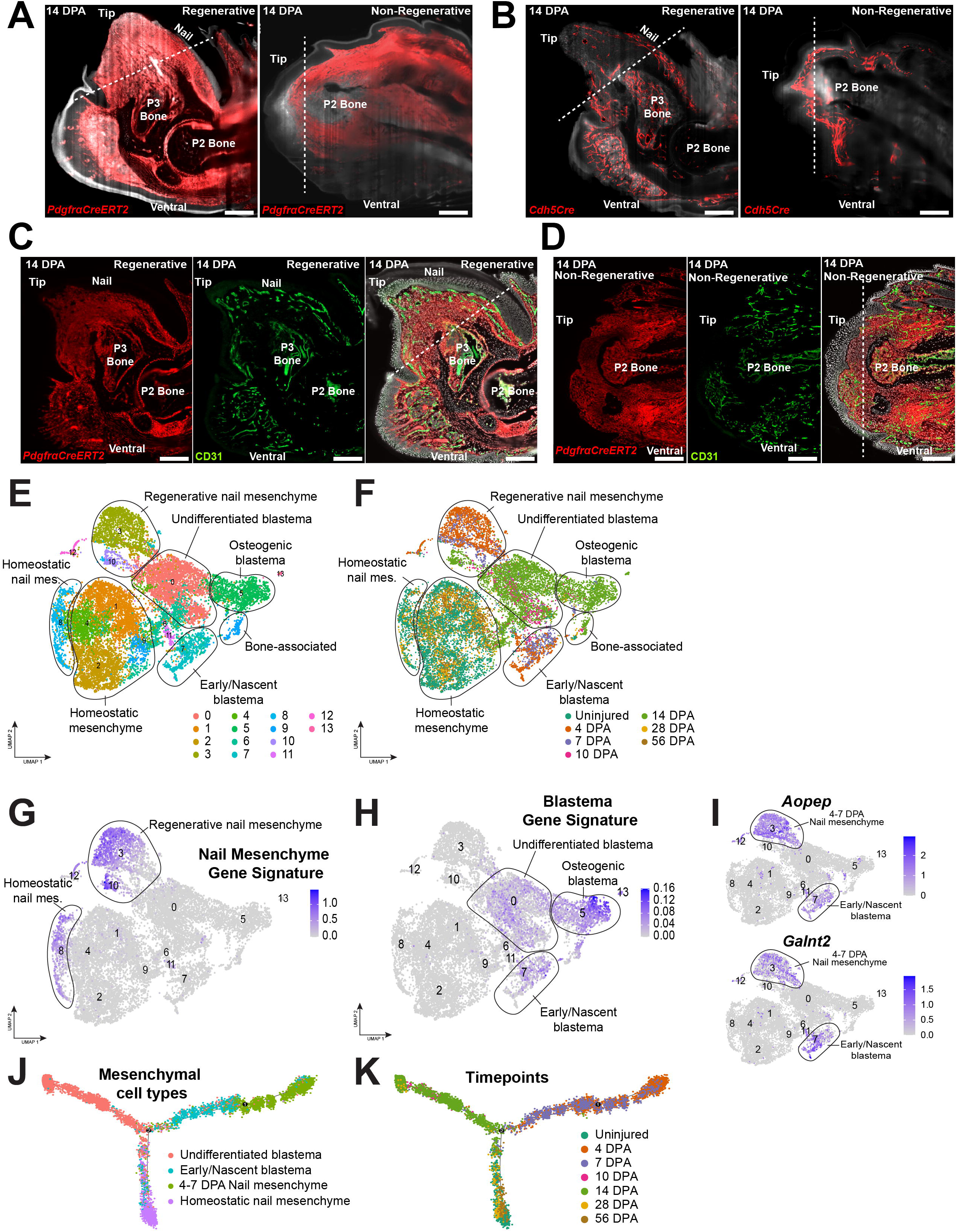
(A-D) *Morphological characterization of amputated digit tip responses in the presence and absence of the nail organ*. (Also see Fig. S1, Table S1, and Video S1-4). **(A)** Representative midline optical sections from lightsheet-generated three-dimensional videos (Video S1 and S2) of cleared whole digit tips of *PdgfraCreERT2*-*TdT* mice at 14 days following regenerative (left panel) or non-regenerative (right panel) amputations, showing tissue morphology and TdT-positive mesenchymal cells (red). The hatched white lines demarcate the amputation plane. Scale bars = 200 µm. **(B)** Representative midline optical sections from lightsheet-generated three-dimensional videos (Video S3 and S4) of cleared whole digit tips of *Cdh5Cre-TdT* mice at 14 days following regenerative (left panel) or non-regenerative (right panel) amputations, showing tissue morphology and TdT-positive endothelial cells (red). The hatched white lines demarcate the amputation plane. Scale bars = 200 µm. **(C, D)** Representative micrographs of midline digit tip sections from 14 DPA *PdgfraCreERT2-TdT* mice following regenerative (C) or non-regenerative (D) amputations, showing mesenchymal cells labelled by the TdT reporter (red), together with immunostaining for CD31 to highlight endothelial cells (green). The merged images on the right also show DAPI counterstaining (white). The white hatched lines indicate the amputation planes. Scale bars = 500 µm. (E-K) *Characterization of 4 and 7 DPA regenerative mesenchymal cell transcriptomes, as defined using scRNA-seq.* (E) UMAP of the merged transcriptomes of *Pdgfra*-positive mesenchymal cells from the new 4 and 7 DPA datasets together with previously-published (Storer et al, 2020, GEO GSE135985; Mahmud et al., 2022, GEO GSE217600) 10, 14, 28 and 56 DPA and uninjured digit tip datasets. Different mesenchymal cell types are annotated and transcriptionally-distinct clusters are color-coded. **(F)** UMAP as in E color-coded to illustrate the datasets of origin of the different transcriptomes. **(G, H)** UMAPs as in E overlaid for expression of the nail mesenchyme gene signature (G) or the blastema gene signature (H), both as previously-defined (Storer et al 2020; Mahmud et al., 2022). Relative enrichment for these signatures is indicated by color-coding, as shown in the adjacent keys. **(I)** UMAPs as in (E) overlaid for expression of genes that are enriched in both the early 4-7 DPA nail mesenchyme (cluster 3) and in the early/nascent blastema (cluster 7) (see Table S1), color-coded as per the adjacent keys. **(J, K)** Monocle trajectory inference including transcriptomes from the 4/7 DPA early nail mesenchyme, the 4/7 DPA nascent blastema, the 10/14 DPA undifferentiated blastema, and the homeostatic nail mesenchyme. (J) shows the different mesenchymal cell types and (K) the datasets of origin, both color-coded as per the adjacent legends.

We extended these findings by analyzing sections of 14 DPA *Pdgfra-Tdt* regenerative and non-regenerative digit tips that were immunostained for CD31 to mark endothelial cells (Fig. 1C, D). This analysis confirmed that *Pdgfra*-TdT-positive mesenchymal cells comprised much of the new tissue between the amputation site and distal epidermis of the regenerating digit tip. The CD31-positive vasculature was well-organized and was less dense within the central blastemal region of the regenerating tissue, consistent with previous reports^8,27^. By contrast, in the non-regenerative condition there was a relatively thin region of *Pdgfra*-TdT-positive mesenchymal tissue between the amputated bone and the epidermis. This new tissue was well-vascularized by CD31-positive blood vessels. Thus, when the nail organ is absent, there is formation of an epithelialized cap of mesenchymal tissue over the amputated bone rather than regeneration.

### scRNA-seq analyses show that in the presence of the nail organ nascent mesenchymal blastema cells are present as early as 4-7 DPA

To understand how the nail organ initiates regeneration versus wound healing, we used single cell transcriptomic approaches. Since the digit tip blastema is first observed at 7 DPA^18^ we characterized regenerating digit tips at 4 and 7 DPA. We first did this by acquiring scRNA-seq data at both timepoints (2 independent runs each) using the10X Genomics Chromium platform. We analyzed the individual datasets using a Seurat-based pipeline (see Experimental Methods). We filtered the datasets for doublets and low-quality cells, confirmed that each individual dataset was of high quality, and then merged the two 4 DPA datasets together and the two 7 DPA datasets together (Fig. S1B, C). We used genes with high variance to compute principal components as inputs for projecting cells in two-dimensions using UMAPs. We performed clustering at a range of resolutions using a shared nearest neighbors-cliq (SNN-cliq)-inspired approach built into the Seurat R package.

Analysis of the merged 4 and 7 DPA regenerative datasets for well-characterized marker genes identified all predicted cell types, including *Pdgfra*-positive mesenchymal cells, immune cells, endothelial cells, vasculature-associated pericytes and vascular smooth muscle (VSM) cells, Schwann cells and a small number of epithelial cells (Fig. S1B, C). We focused further analysis on *Pdgfra*-positive mesenchymal cells, since at the peak of regeneration (10 and 14 DPA), these comprise the majority of regenerative blastema cells (see Fig. 1A, C)^7,8^. We performed a comparative analysis, merging the 4 and 7 DPA mesenchymal cell transcriptomes with our previously-published uninjured and 10 to 56 DPA regenerative mesenchymal cell transcriptomes^8,18^ (GEO GSE135985 and GSE217600). This analysis (Fig. 1E, F) identified two major 10-14 DPA regenerative mesenchymal cell populations. One was the regenerative nail mesenchyme (cluster 10) expressing a previously-defined nail mesenchyme gene signature^18^ (see Experimental Methods) that included genes like *Gpm6a, Cyp2f2, Nrg2, Rspo4* and *Snhg11* (Fig. 1G; Fig. S1D). The second population included 10-14 DPA regenerative blastema cells expressing a previously-defined blastema gene signature^8^ (Fig. 1H; see Experimental Methods). These blastema cells segregated into two clusters (5 and 0). Cluster 5 included blastema cells differentiating down the osteogenic lineage as indicated by high relative expression of genes like *Panx3* and *Ibsp* (Fig. S1D, E). The other cluster (0) included the undifferentiated 10-14 DPA blastema cells.

Notably, the 4 and 7 DPA regenerative cells (clusters 3 and 7) were transcriptionally-distinct from the 10-14 DPA regenerative and uninjured mesenchymal cells, but co-clustered with each other (Fig. 1E, F). The 4-7 DPA cells in cluster 3 expressed the nail mesenchyme gene signature while those in cluster 7 expressed the blastema gene signature (Fig. 1G, H; Fig. S1D), identifying them as the early regenerative nail mesenchyme and the early/nascent blastema, respectively. We asked about the transcriptional state of these early regenerative mesenchymal cells by performing differential gene expression analysis (Table S1). This analysis, coupled with UMAP gene expression overlays showed that relative to all other mesenchymal cells, the nascent blastema was specifically enriched for a gene subset that included *Sfrp4, Ebf3* and *Ccn5* (Fig. S1D, F; Table S1). Moreover, this analysis identified a second group of genes that were differentially-enriched in both the 4-7 DPA nascent blastema and in the early nail mesenchyme including *Aopep, Galnt2, Cacna1c* and *Tent5a* (Fig. 1I; Fig. S1G).

One potential explanation for the commonality in differential gene expression between the 4-7 DPA nail mesenchyme and nascent blastema cells is that they are lineally-related. To explore this idea, we performed Monocle trajectory inference, including the 4/7 DPA nail mesenchyme, the nascent blastema, the 10/14 DPA undifferentiated blastema, and the homeostatic nail mesenchyme. This analysis (Fig. 1J, K) defined a trajectory with the 4/7 DPA nail mesenchyme at one end, followed by the 4 to 7 DPA nascent blastema and then the 10/14 DPA blastema. It also defined a branchpoint where the early nail mesenchyme and nascent blastema gave rise to the homeostatic nail mesenchyme rather than the undifferentiated blastema. This analysis supports a model where following regenerative amputations some 4/7 DPA nail mesenchyme cells differentiate into blastema cells and others ultimately regenerate the homeostatic nail mesenchyme, consistent with our previous lineage tracing analyses^18^.

### Single cell spatial analysis defines the earliest stages of digit tip regeneration

To define the spatial location of these different regenerative populations, we performed single cell multiplexed *in situ* gene expression analysis using the 10X Xenium platform^29–32^. We analyzed sagittal fresh-frozen sections from 5 to 14 DPA with a custom 480 gene probeset (Table S2, panel 1) optimized based on our scRNA-seq datasets to distinguish different digit tip cell types, including transcriptionally-distinct mesenchymal cell populations. We defined a region of interest (ROI) that encompassed the regenerating digit tip distal to the *Spp1/osteopontin*-positive amputated bone (see Fig. 2C). In total, we analyzed multiple sections from at least three different mice at each timepoint (see Experimental Methods). After initial processing of individual sections and removal of a small number of objects with low transcript counts (likely cellular fragments), we merged all cells from a given timepoint. We ultimately obtained 16418, 18964, 25420 and 52642 well-integrated transcriptomes at 5, 7, 10 and 14 DPA, respectively. We analyzed the 10 and 14 DPA datasets on their own, but since there were fewer cellular transcriptomes at 5 and 7 DPA, we merged these two datasets (Fig. 2A; Fig. S2A).

**Figure 2.**
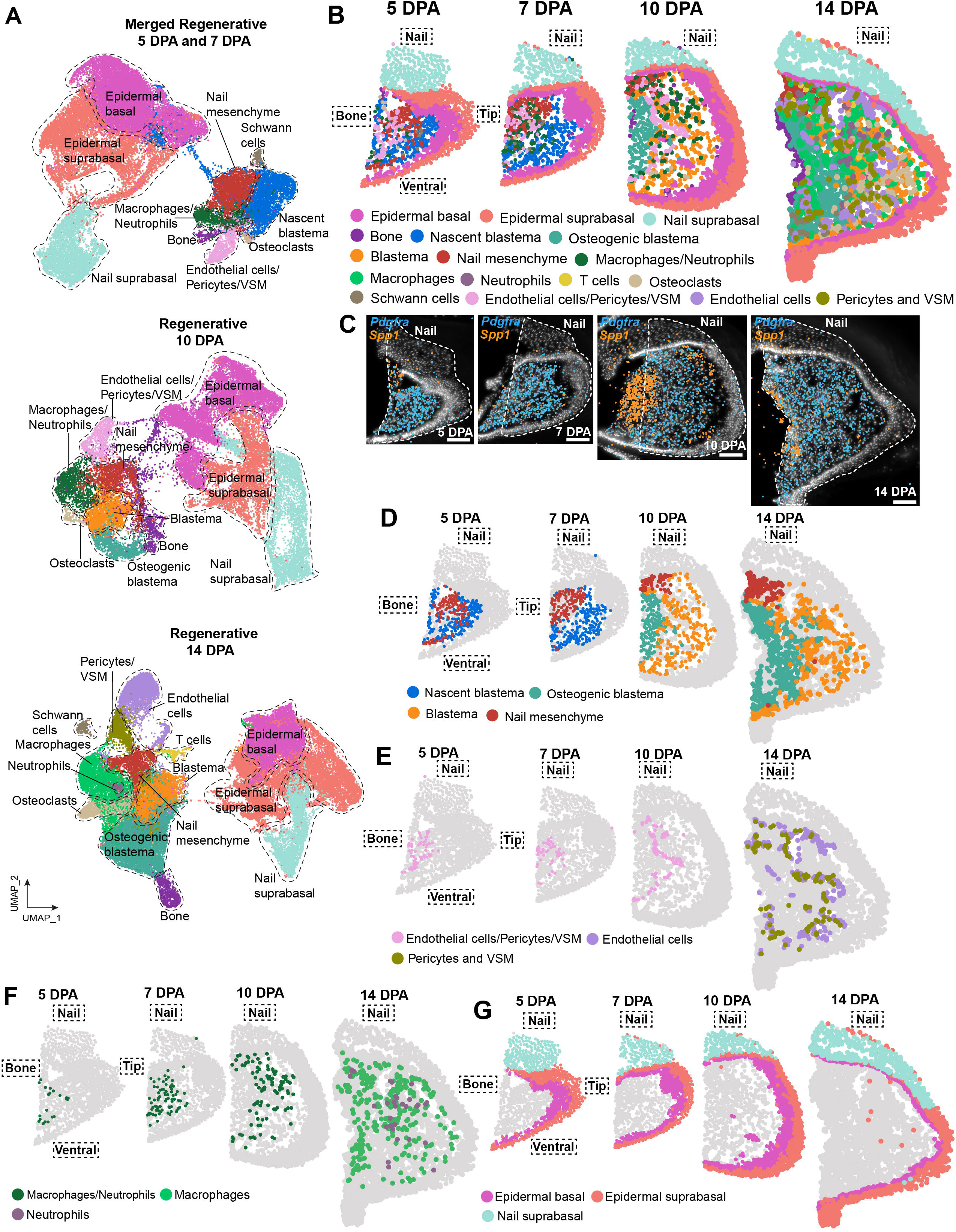
*Single cell spatial transcriptomic characterization of the regenerating digit tip from 5 to 14 DPA.* (Also see Fig. S2 and Table S2). **(A)** UMAPs of regenerating digit tip transcriptomes generated using Xenium-based single cell spatial transcriptomics with a custom 480 gene probeset (Table 2) optimized based on our scRNA-seq datasets to distinguish different digit tip cell types, including transcriptionally-distinct mesenchymal cell populations. The UMAPs show the merged 5 and 7 DPA datasets (top panel), the 10 DPA dataset (middle panel) and the 14 DPA dataset (bottom panel). In all cases, the UMAPs are annotated and color-coded to show cell types as identified by marker gene analysis (see Fig. S2). **(B)** Spatial plots of the datasets in (A), showing all detected cell types on representative regenerating digit tip midline sections. Individual cell types are color-coded as per the legend. **(C)** High-resolution Xenium Explorer images of the same representative sections as in (B), showing expression of *Pdgfra* mRNA to highlight mesenchymal cells (blue dots) and *Spp1* mRNA to highlight the proximal bone (orange). Hatched white lines outline the regenerated tissue. Scale bars = 100 µm. **(D-G)** Spatial plots of the same representative sections as in (B), showing (D) transcriptionally-distinct mesenchymal cell populations, (E) vasculature-associated endothelial and mural cells (pericytes and VSM cells), (F) macrophages and neutrophils, or (G) epithelial cell populations. For both vasculature and immune cells, they did not cleanly segregate transcriptionally at earlier timepoints due to lower cell numbers, and thus are shown together. All Seurat-based spatial plots (B, D, E-G) are sized relative to the unprocessed Xenium Explorer images of the same sections (as in C).

These analyses provided a global spatial overview of the regenerating digit tip. UMAPs, spatial plots and well-validated marker genes (Fig. 2A, B; Fig. S2A-E) identified the predicted cell types at all timepoints, including various populations of epidermal and immune cells, osteoclasts, vasculature endothelial and mural cells, Schwann cells and *Pdgfra*-positive mesenchymal cells. At all timepoints epithelial cells, including epidermal and nail epithelial cells, were the most abundant, followed by mesenchymal cells, including nail mesenchyme and blastema cells. Spatial plots of these datasets (Fig. 2B-D) demonstrated robust regeneration and mesenchymal tissue growth from 5 to 14 DPA and identified the location of all cell types. For vasculature-associated cells, at early timepoints these were located closer to the bone stump while at 14 DPA they were also present distally (Fig. 2E). In all cases, the vascular density was lower in the central blastema, consistent with the morphological analyses (Fig. 1B, C; Video S3) and reports that the blastema is largely avascular^27,33^. Immune cells, predominantly macrophages and neutrophils, were scattered throughout the regenerating tissue at all timepoints (Fig. 2F). At early timepoints monocytes/macrophages and neutrophils did not cluster distinctly, but many immune cells expressed the neutrophil-specific mRNA *Csf3r* (Fig. S2A-E). At 14 DPA, when there were many more total cells, neutrophils and macrophages clustered separately, although there were proportionately fewer neutrophils at this later timepoint (Fig. 2A, F). As for epithelial cells, the Xenium probeset readily distinguished *Cspg4*-positive, *Itga6*-positive basal epidermal cells versus *Erbb3*-positive, *Krt10*-positive suprabasal epidermal cells. By 5 DPA these were already distributed around the entire distal digit tip (Fig. 2G). The Xenium probeset also distinguished suprabasal cells of the nail versus the epidermis (Fig. 2A, G); the epidermal suprabasal cells were enriched for *Krt10* and *Krt17* while the nail suprabasal cells were instead enriched for *Cybrd1, Fhod3, Fgf18, Car2* and *Slit2* (Fig. S2A-E).

To better-understand blastema initiation and formation, we performed a more detailed analysis of the mesenchymal cells, subsetting and merging all *Pdgfra*-positive cells from all timepoints (Fig. 3A, B). This analysis showed that mesenchymal cells evolved transcriptionally over regenerative time; the 5 and 7 DPA mesenchymal cells clustered separately from the 14 DPA cells, as in the scRNA-seq analyses, and the 10 DPA cells were split between the early and late timepoint clusters. Spatial plots showed that at both 5 and 7 DPA the early nail mesenchyme cells were located dorsally beneath the nail epithelium, with the nascent blastema cells localized throughout the remainder of the regenerating digit tip between the bone and epithelium (Fig. 3C). Notably, at 5 DPA some nail mesenchyme cells were also intermingled with nascent blastema cells (Fig. 3C).

**Figure 3.**
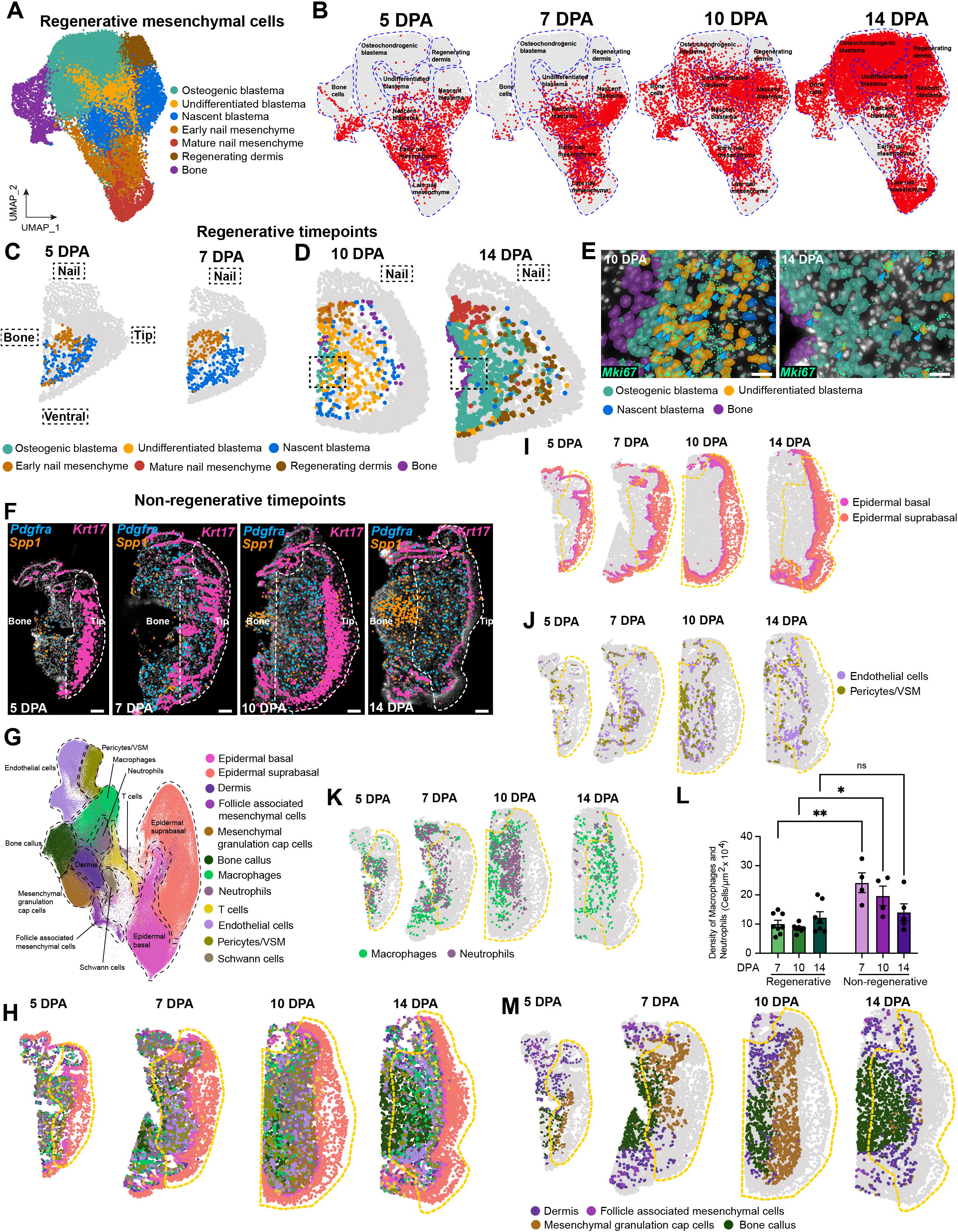
(A-E) *Characterization of regenerative mesenchymal cells over time using single cell spatial transcriptomics*. (Also see Fig. S3). **(A)** UMAP of merged 5 to 14 DPA *Pdgfra*-positive mesenchymal cell transcriptomes, extracted from the datasets in Fig. 2A. The UMAP shows color-coded, transcriptionally-distinct mesenchymal cell types, annotated as informed by regenerating digit tip scRNA-seq-based analyses^8,18^ (Fig. 1E, F). **(B)** The same UMAP as in (A), color-coded to identify the datasets of origin for the different transcriptomes. **(C, D)** Spatial plots of the same 5 to 14 DPA regenerative midline sections as in Fig. 2B, showing the spatial location of transcriptionally-distinct regenerating *Pdgfra*-positive mesenchymal cell populations identified in the UMAP in (A). (C) shows the 5 and 7 DPA timepoints, and (D) the 10 and 14 DPA timepoints, in both cases color-coded as per the adjacent legend. **(E)** High-resolution Xenium Explorer images of the 10 and 14 DPA regenerating digit tip, showing the boxed regions indicated in (D). The images show the different blastema populations, color-coded as per the adjacent legend, as well as expression of *Mki67* mRNA (green dots), which is largely limited to the orange undifferentiated blastema cells (arrowheads). Scale bars = 20 µm. (F-M) *Characterization of the non-regenerative wound-healing response using single cell spatial transcriptomics*. **(F)** High resolution Xenium Explorer images of representative midline digit tip sections at 5 to 14 days following non-regenerative amputations showing expression of *Pdgfra* mRNA to highlight mesenchymal cells (blue dots), *Spp1* mRNA to highlight the proximal bone (orange dots) and *Krt17* mRNA (pink dots) to show the epidermis. Hatched white lines outline the regions distal to the amputations. Scale bars = 100 µm. **(G)** UMAP showing merged Xenium-based single cell transcriptomes from the digit tip 5 to 14 days following non-regenerative amputations annotated and color-coded to denote cell types identified by marker gene analysis. **(H)** Spatial plots of the dataset in (G), showing all detected cell types on the same representative non-regenerative digit tip midline sections as shown in (F). Individual cell types are color-coded as per the adjacent legend and the yellow hatched lines show the tissue distal to the amputation. **(I-K)** Spatial plots of the same representative sections as in (F), showing (I) the different epithelial cell populations, (J) vasculature-associated endothelial and mural cells and (K) macrophages and neutrophils. In (J) the mural cells (pericytes and VSM cells) did not cleanly segregate transcriptionally, and thus are shown together. **(L)** Graph showing the density of macrophages/neutrophils in regenerating versus non-regenerating digit tip midline sections from 7 to 14 DPA. Quantification was performed using high resolution Xenium Explorer images, and each dot represents an individual section. n = 3-4 mice per group. *p<0.05, **p<0.01, ns=non-significant. **(M)** Spatial plots of the same representative sections as in (F), showing transcriptionally-distinct mesenchymal cell populations. The hatched lines outline tissue distal to the amputation plane. All Seurat-based spatial plots (C, H-K, M) are sized relative to the unprocessed Xenium Explorer images of the same sections (as in F).

At 10 and 14 DPA (Fig. 3D) the nail mesenchyme cells were still appropriately localized beneath the nail epithelium, although the 14 DPA cells were transcriptionally-distinct from 5/7 DPA nail mesenchyme cells (Fig. 3A), as seen with the scRNA-seq. At this time of peak regeneration there were also two transcriptionally and spatially-distinct blastema populations (Fig. 3A, B, D), both of which expressed blastema signature genes such as *Ltbp2* and *Arsi* (Fig. S3A). One included more proliferative, *Ki67*-high undifferentiated blastema cells (Fig. 3D, E). These comprised the majority of blastema cells at 10 DPA when they were localized throughout the non-epithelial regenerating tissue. The second 10/14 DPA blastema population included cells differentiating down the osteogenic lineage. These osteogenic blastema cells were enriched for genes like *Alpl* (Fig. S3B) and had a lower proliferative index, as indicated by *Ki67* expression (Fig. 3E). At 14 DPA they comprised the majority of blastema cells, and were located relatively closer to the regenerating bone (Fig. 3D). Notably, at 10 DPA, these osteogenic blastema cells were only located immediately adjacent to the bone (Fig. 3D, E; Fig. S3B), implying a spatial gradient of osteogenic differentiation originating at the bone surface. Finally, there was a third 14 DPA mesenchymal population that we have called regenerating dermis cells (Fig. 3A, B). These cells were located between the osteogenic blastema and the distal/ventral epidermis (Fig. 3D) and expressed both blastema genes and genes characteristic of the dermis such as *Cpxm2, Col26a1, Cxcl14, Ari, Aff3, Npr3* and *Mmp9* (Fig. S3A, C). Based on their location and transcriptional phenotype we posit these are blastema cells differentiating down the dermal lineage. Thus the blastema is first initiated at 5 to 7 DPA, it expands so that the undifferentiated proliferative blastema is maximal around 10 DPA, and then spatially-determined differentiation ensues to regenerate the bone and dermis.

### In the absence of the nail organ there are no blastema-like cells and mesenchymal cells instead become bone callus and mesenchymal granulation cap cells

We next performed a similar single cell spatial transcriptome analysis of the non-regenerative situation, using the same custom probeset to analyze the digit tip at 5 to 14 days following amputations that remove the nail organ. We defined ROIs encompassing the digit tip distal to the amputation plane, using Xenium Explorer and expression of *Spp1/osteopontin*, *Pdgfra*, and *Krt17* to define the bone, mesenchyme and epidermis, respectively (Fig. 3F). Any tissue containing intact hair follicles was considered to be proximal to the amputation. We then used these ROIs to analyze multiple sections from at least 3 different mice at each of 5, 7, 10 and 14 DPA.

UMAP and marker gene analysis of the individual datasets defined similar cell types at all timepoints (Fig. S3D-G), and we therefore merged all datasets together. Analysis of this merged dataset (Fig. 3G, Fig. S3H) identified epithelial cells, immune cells and osteoclasts, vasculature and lymphatic endothelial and mural cells, Schwann cells and *Pdgfra*-positive mesenchymal cells at all times. Spatial plots identified tissue growth between 5 and 14 DPA, but showed this was not distally-oriented and/or spatially well-organized relative to the regenerative condition (Fig. 3F, H). At all timepoints, there was an epidermal covering with underlying wound tissue adjacent to the amputated bone (Fig. 3H, I). However, at 5 and 7 DPA the new epidermis was not as thick as in the regenerative situation (compare Fig. 3I to 2G). There were also no apparent dorsal-ventral differences in tissue growth at any timepoint, and there were no hair follicles in the tissue distal to the amputation plane.

The wound tissue between the epidermis and bone was largely comprised of vasculature, immune and mesenchymal cells (Fig. 3H). Consistent with the morphological studies (Fig. 1B, D) the wound tissue was well-vascularized at all timepoints, as visualized by the distribution of endothelial and mural cells (Fig. 3J). As for immune cells, at 5 to 14 DPA there were many macrophages and neutrophils (Fig. 3K) and quantification of Xenium Explorer images demonstrated these were approximately two-fold more dense than in the regenerative condition (Fig. 3L; also compare Fig. 3K and 2F).

There were also many *Pdgfra*-positive mesenchymal cells within the non-regenerative wound tissue, and at 10 and 14 DPA these were the most abundant cell type after the epidermal cells (Fig. 3M). Detailed analysis defined 4 major mesenchymal cell populations (Fig. 3G). One included hair follicle-associated mesenchymal cells that were only present in tissue proximal to the amputation plane (Fig. 3M; Fig. S3I). A second population included bone callus cells that were enriched for osteogenic genes like *Comp, Alpl* and *Acan* (Fig. 3M; Fig. S3J). These were adjacent to the amputated bone and were already present in small numbers at 5 DPA. By 14 DPA the bone callus cells had expanded so they comprised much of the mesenchymal tissue distal to the bone. A third population included mesenchymal cells that were beneath the epidermis and above the amputated bone/bone callus (Fig. 3M). These were already present at 5 DPA and formed a well-defined sub-epidermal layer at 7 and 10 DPA. Analysis of gene expression showed that these cells had characteristics of wound fibroblasts within granulation tissue, including expression of matrix-associated mRNAs like *Fmod*, *Fbn2*, *Col24a1* and *Mmp11*, as well as profibrotic mRNAs such as *Prss35* (Fig. S3J). We have therefore called these mesenchymal granulation cap cells. These cells were still present at 14 DPA, but were partially replaced by cells that were transcriptionally-similar to dermal cells proximal to the amputation plane (Fig. 3M). Thus, in the absence of the nail organ the multicellular response to amputation is similar to wound-healing and scarring and not to regeneration.

### The regenerative and nonregenerative mesenchymal cell responses differ as early as 5 DPA

These findings indicate that mesenchymal cell responses in the presence or absence of the nail bed are completely different, with blastema formation and differentiation occurring in the regenerative condition, and scar and bone callus formation in the non-regenerative. To better-understand these differences, we merged *Pdgfra*-positive mesenchymal cells from the 14 DPA datasets. UMAPs showed that the 14 DPA regenerative and non-regenerative mesenchymal cells were transcriptionally distinct, since with the exception of bone stump cells they completely segregated from each other (Fig. 4A, B). Of the regenerative cells, 80% were blastema cells (undifferentiated and osteogenic), and a further 7% were nail mesenchyme cells. Relative to the non-regenerative cells, the regenerative mesenchymal cells were highly enriched for transcription factor and growth factor genes such as *Alx4, Bmp5, Fgf10* and *Gldn* (Fig. 4C; Fig. S4A). By contrast, 21% of the nonregenerative mesenchymal cells were mesenchymal granulation cap cells and 36% bone callus cells in varying states of differentiation (Fig. 4A, B). The remainder were hair follicle-associated and dermal mesenchymal cells. Relative to the regenerative cells, the nonregenerative mesenchymal cells were enriched for genes implicated in fibrosis and inflammation, including *Cilp, Thbs4, Angptl1* and *Sulf2* (Fig. 4D; Fig. S4A). Notably, differentiating osteogenic cells in the two conditions clustered separately (Fig. 4A, B), suggesting that bone repair and bone regeneration follow different transcriptional trajectories.

**Figure 4.**
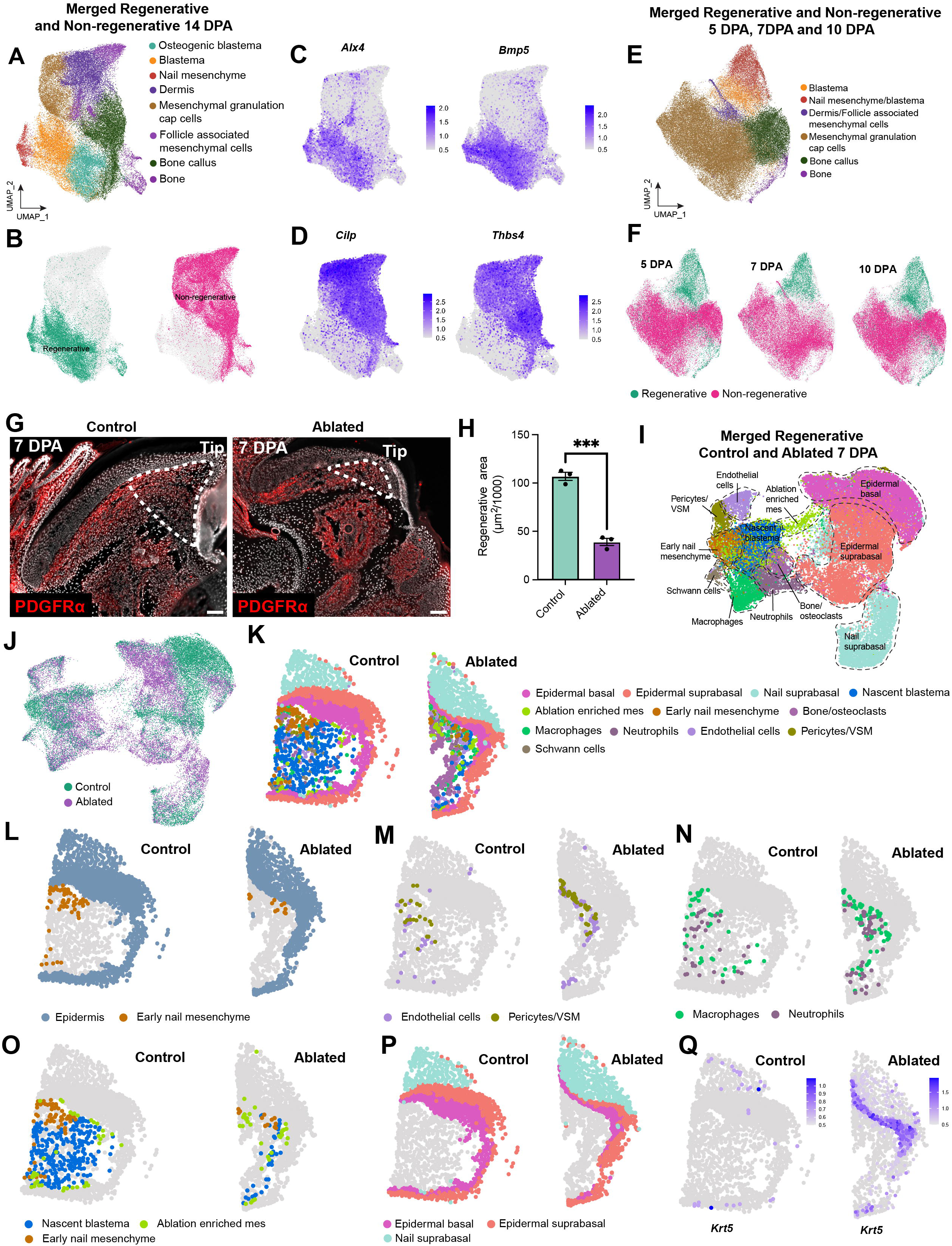
(A-F) *Comparison of Pdgfra-positive mesenchymal cells from 5 to 14 days following regenerative versus non-regenerative amputations.* (Also see Fig. S4). **(A)** UMAP of merged 14 DPA regenerative and non-regenerative *Pdgfra*-positive mesenchymal cell transcriptomes, extracted from the datasets in Fig. 2A and 3G. The UMAP shows transcriptionally-distinct mesenchymal cell types, annotated and color-coded. **(B)** The same UMAP as in (A), color-coded to show transcriptomes from the 14 DPA regenerative (green) versus non-regenerative (pink) mesenchymal cell datasets to highlight their transcriptional segregation. **(C, D)** Expression overlays on the UMAP in (A), showing relative levels of expression of two regeneration-enriched mRNAs, *Alx4* and *Bmp5* (C) and two non-regenerative mRNAs, *Cilp* and *Thbs4* (D), color-coded as per the adjacent keys. **(E)** UMAP of merged 5, 7 and 10 DPA regenerative and non-regenerative *Pdgfra*-positive mesenchymal cell transcriptomes, extracted from the datasets in Fig. 2A and 3G. The UMAP shows transcriptionally-distinct mesenchymal cell types, annotated and color-coded. **(F)** The same UMAP as in (E), color-coded to show transcriptomes from the 5, 7 and 10 DPA regenerative (green) versus non-regenerative (pink) mesenchymal cell datasets to highlight their transcriptional segregation. (G-Q) Characterization of digit tip regeneration at 7 DPA with and without partial nail mesenchyme ablation. (G, H) Lmx1b-DTA (Ablated) or *Lmx1b-WT* (Control) mice were treated with tamoxifen, their digit tips were amputated 8 days later, and regeneration was characterized after a further 7 days (7 DPA). (G) shows representative midline sections immunostained for PDGFRα (red), and counterstained with DAPI (white) to highlight cellular nuclei. The hatched lines demarcate the regenerated tissue area exclusive of the epithelium. Scale bars = 100 µm. (H) shows quantification of the regenerated tissue area in the two different conditions. Each dot represents an individual mouse, with three midline sections per mouse quantified, and the average area plotted. n = 3 per condition, ***p<0.001. **(I)** UMAP showing single cell Xenium-based transcriptomes from 7 DPA *Lmx1b-DTA* digit tips merged with 7 DPA control digit tip transcriptomes (the 7 DPA transcriptomes from the dataset in Fig. S2A), annotated and color-coded for cell types as identified by marker gene analysis. **(J)** UMAP as in (I), color-coded to show 7 DPA *Lmx1b-DTA* (Ablated) versus control transcriptomes. **(K)** Spatial plots of the dataset in (I), showing all detected cell types on representative 7 DPA *Lmx1b-DTA* (Ablated) and control midline sections. Cell types are color-coded as per the adjacent legend. **(L-P)** Spatial plots of the same representative 7 DPA *Lmx1b-DTA* (Ablated) and control sections as in (K), showing (L) nail mesenchyme cells and all epithelial cells, (M) endothelial cells and mural cells (pericytes and VSM shown together), (N) macrophages and neutrophils, (O) transcriptionally-distinct mesenchymal populations, including the nail mesenchyme, nascent blastema, and a population of mesenchymal cells that were enriched in the *Lmx1b-DTA* digit tips (Ablation enriched mes), and (P) the different epithelial cell populations. **(Q)** Spatial plots of the same representative 7 DPA *Lmx1b-DTA* (Ablated) and control sections as in (K), overlaid for expression of *Krt5* mRNA, color-coded as per the adjacent keys.

We asked when these differences arose by merging mesenchymal cells from the 5, 7 and 10 DPA regenerative and nonregenerative Xenium datasets. This analysis (Fig. 4E, F) showed that mesenchymal cells from the two conditions were already transcriptionally distinct by 5 DPA with the exception of the bone stump cells. As at 14 DPA, the 5 to 10 DPA regenerative mesenchymal cells were enriched for *Alx4, Bmp5, Fgf10* and *Gldn,* and the nonregenerative for *Cilp, Thbs4,* and *Angptl1* (Fig. S4B). Thus, the transcriptional differences between regenerative and non-regenerative mesenchymal cells are evident by 5 DPA and persist until at least 14 DPA.

### The nail mesenchyme determines the early cellular regenerative response

These findings indicate that the nail organ is required for a regenerative response by 5 DPA. Since the nail organ is largely comprised of two cell types, epithelial and nail mesenchyme cells, we asked about the relative importance of these two cell types. To do so, we took advantage of a mouse line carrying CreERT2 knocked-in to the *Lmx1b* locus^85^ and a floxed *Diptheria toxin A* (DTA) allele knocked-in to the *Rosa26* locus (*Lmx1b-DTA* mice). We previously showed *Lmx1b* is specifically expressed in the adult nail mesenchyme and that tamoxifen treatment of *Lmx1b-DTA* mice causes selective partial nail mesenchyme ablation and a deficit in regenerative tissue size at 14 DPA^18^. We therefore treated *Lmx1b-DTA* mice with tamoxifen, performed regenerative amputations 8 days later and after a further 7 days (7 DPA) analyzed the regenerating digit tips. As controls, we used mice carrying the same *Lmx1b-CreERT2* allele but without the *DTA* transgene (*Lmx1b-WT* mice). PDGFRα immunostaining and quantification of midline sections from these 7 DPA digit tips demonstrated there was two to three-fold less regenerated tissue when the nail mesenchyme was partially ablated in the *Lmx1b-DTA* digit tips (Fig. 4G, H).

We asked about the cellular basis of this deficit using single cell spatial transcriptomics, analyzing sections from 7 different 7 DPA *Lmx1b-DTA* mice that had been treated as for the immunostaining experiments. UMAPs and marker gene analysis of this dataset identified epithelial cells, *Pdgfra*-positive mesenchymal cells, vasculature endothelial and mural cells, Schwann cells, immune cells and osteoclasts (Fig. S4C). We then merged the 7 DPA ablated and control regenerating datasets to allow a direct comparison of the two conditions (Fig. 4I, J). UMAP and spatial plot analyses of the merged dataset showed that following partial nail mesenchyme ablation there was much less tissue (Fig. 4K) and, as predicted, fewer nail mesenchyme cells (Fig. 4L). Several cell types were nonetheless spatially and transcriptionally similar in the two conditions; vasculature-associated cells, Schwann cells, and immune cells from the ablated and control conditions co-clustered transcriptionally (Fig. 4I, J) and their localization was approximately similar (Fig. 4M, N).

This was not however the case for mesenchymal or epidermal cells. In the ablated condition, mesenchymal cells were greatly reduced in number and were spatially disorganized (Fig. 4O). Moreover, in addition to the reduction in nail mesenchyme cells, there were few nascent blastema cells and many of the mesenchymal cells that were present clustered separately from the nail mesenchyme and blastema cells, indicating they were transcriptionally-distinct (Fig. 4I, 4O; ablation-enriched cells). These transcriptionally-distinct cells comprised approximately 50% of the mesenchymal cells in the ablated digit tips, and only approximately 16% in control regenerating digit tips (Fig. 4O). Thus, following partial nail mesenchyme ablation, blastema formation was deficient and the total numbers and proportions of different mesenchymal cell populations were perturbed.

The epidermal cells were also altered, with many of the ablated regenerating basal epidermal cells clustering distinctly (Fig. 4I, J). Spatial plots demonstrated that re-epithelialization was also perturbed, since in many sections the basal cell layer did not entirely cover the ablated digit tips (Fig. 4P). In addition, many of the basal epidermal cells in the ablated situation expressed high levels of *Krt5* (Fig. 4Q). Given the key role KRT5 plays in early re-epithelialization^34,35^, this finding suggests the ablated digits may be at an earlier stage of epithelial wound-healing. Thus, partial nail mesenchyme ablation caused deficits in mesenchymal blastema formation, re-epithelialization, and regenerative tissue growth at 7 DPA.

To exclude the possibility that these phenotypes were due to the presence of the *Lmx1bCreERT2* allele and/or tamoxifen treatment, we performed single cell spatial transcriptomics on sections from tamoxifen-treated *Lmx1b-WT* digit tips at 7 DPA. We used a different probeset for this analysis (Table S2) that could nonetheless distinguish the different digit tip cell types. This analysis identified the same transcriptionally-distinct cell types in the *Lmx1b-WT* 7 DPA digit tips as in the other control dataset (Fig. S4D) and confirmed their spatial locations (Fig. S4E-I). Thus, the perturbations observed in the ablated *Lmx1b-DTA* digit tips are not due to the presence of the *Lmx1bCreERT2* allele and/or tamoxifen treatment.

### The nail mesenchyme and nascent blastema create an early regeneration-specific ligand environment that includes expression of multiple BMPs

These data indicate the early nail mesenchyme is essential for blastema formation so we asked whether the nail mesenchyme secretes potential pro-regenerative ligands. To do so, we identified ligands expressed by the 4 and 7 DPA early regenerative digit tip mesenchymal cells in the scRNA-seq datasets (Fig. 1E, F) using our previously-published ligand-receptor database^31,36^. We defined a ligand mRNA as expressed if it was detectable in at least 5% of the relevant cells. We identified 80 and 65 early regenerative nail mesenchyme and nascent blastema ligands, respectively, with 52 expressed by both cell types (Table S3), including, for example, *Bmp2* and *Pdgfc*.

To ask if these ligands might be important for regenerative mesenchymal responses, we defined receptors expressed in the 4-7 DPA nail mesenchyme and blastema cells using the same database and including receptors if they expressed in 20% or more cells. We identified 49 and 39 receptors in the nail mesenchyme and nascent blastema cells, respectively with 31 co-expressed by both populations (Table S4), including BMP/TGFβ family receptors and tyrosine kinase receptors for the FGFs, PDGFs and IGFs.

We used this information to predict ligands that were biologically active within the early regenerative mesenchyme. This modeling (Fig. 5A, Table S5) predicted 46 ligand-receptor interactions between the early nail mesenchyme and the nascent blastema. 28 ligands were expressed by both cell types, 11 by only the nail mesenchyme (*Bmp7, Gdf6, Igf2, Inhbb, Pgf, Rgma, Sema3a, Sema3b, Sema6a, Vegfd* and *Wnt11*) and 6 by only the nascent blastema (*Ereg, Hbegf, Il1b, Il6, Lif* and *Wnt4*). A similar profile was obtained when modeling early potential autocrine nail mesenchyme interactions (nail mesenchyme ligands on to nail mesenchyme receptors) (Fig. S5A, Table S5). Probes for more than half of these ligands (24/45) were present in the Xenium panel (see Table S2), so we used the 5-10 DPA spatial transcriptomic and scRNA-seq datasets to identify ligands that were (i) highly enriched in the regenerative nail mesenchyme or blastema cells relative to non-regenerative mesenchymal cells and (ii) expressed in early regenerating mesenchymal cells at levels greater than or similar to other digit tip cell types.

**Figure 5.**
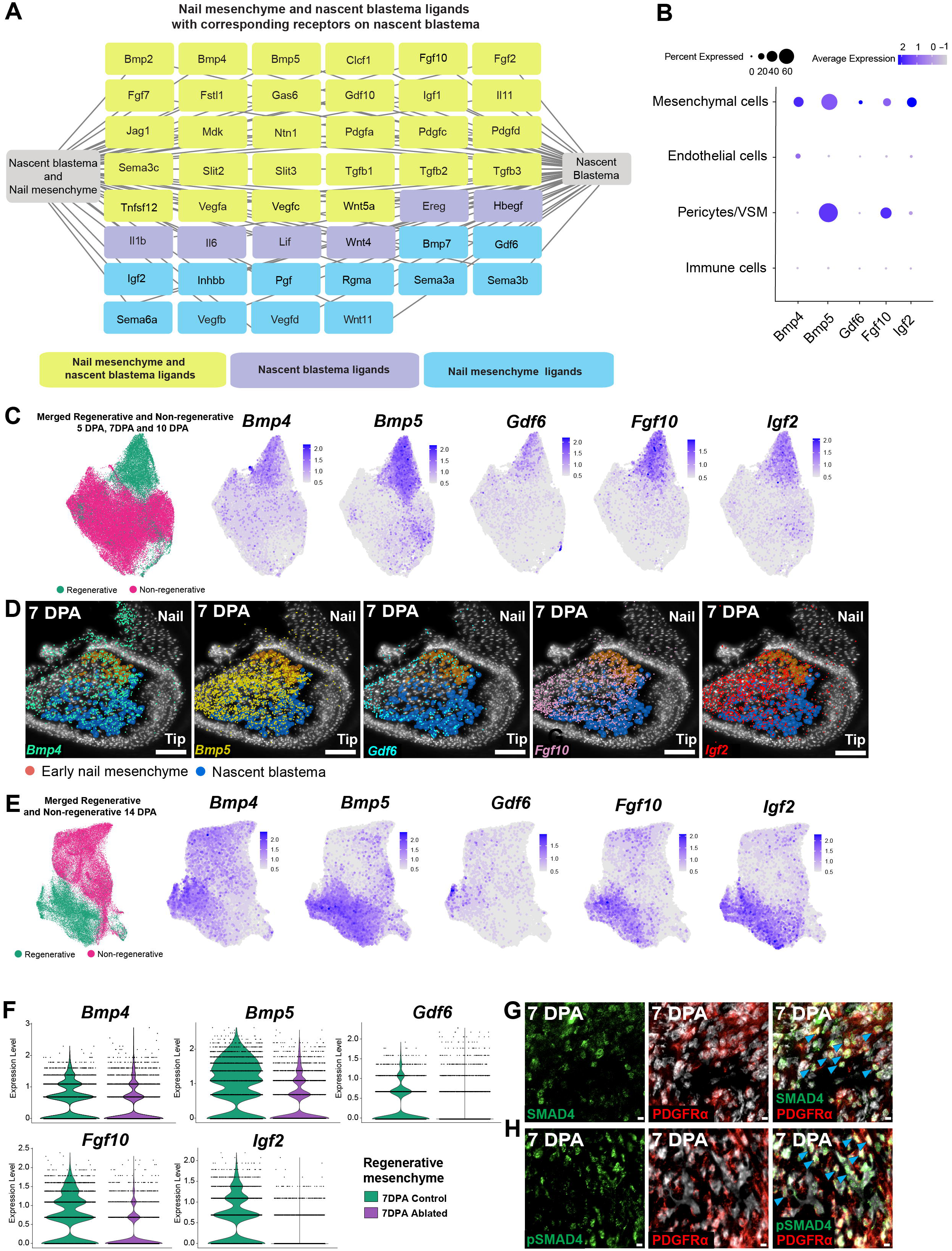
*Identification and characterization of mesenchymal ligands predicted to act on the initial stages of blastema formation and nail mesenchyme regeneration*. (Also see Fig. S5 and Tables S3-5). **(A)** 5-7 DPA early nail mesenchyme and nascent blastema scRNA-seq-derived cellular transcriptomes (as in Fig. 1E) were analyzed for their expression of ligand and ligand receptor mRNAs. Ligand mRNAs were included if they were expressed in ≥ 5% of the nascent blastema or early nail mesenchyme cells while receptor mRNAs were included if they were expressed in ≥ 20% of the nascent blastema cells. These data were used to generate predictive models of the bioactive ligand environment within the early regenerating digit tip. Each box includes a ligand known to bind to a corresponding receptor expressed by the nascent blastema cells. The ligands are color-coded to show those that are expressed in both cell populations (yellow/green), those that are only expressed in the nascent blastema (purple) and those that are expressed only by the nail mesenchyme (blue). **(B)** Dot plot showing expression levels of *Bmp4, Bmp5, Gdf6, Fgf10* and *Igf2* mRNAs in various 7 DPA regenerating digit tip cell types, as determined from the scRNA-seq data in Fig. S1C. The size of the dot indicates the percentage of cells detectably expressing the mRNA, and the color indicates relative expression level, coded as per the adjacent key. **(C)** UMAPs of the merged 5, 7 and 10 DPA regenerative and non-regenerative *Pdgfra*-positive mesenchymal cell transcriptomes (as in Fig. 4E), overlaid to show expression of the 5 selected ligand mRNAs, *Bmp4, Bmp5, Gdf6, Fgf10* and *Igf2.* The panel on the left is color-coded to show the regenerative (green) versus non-regenerative (pink) digit tip transcriptomes. **(D)** High-resolution Xenium Explorer images of a 7 DPA regenerating digit tip section, showing the early nail mesenchyme and nascent blastema, color-coded as per the adjacent legend, as well as *Bmp4* mRNA (green dots), *Bmp5* mRNA (yellow dots), *Gdf6* mRNA (turquoise dots), *Fgf10* mRNA (pink dots) and *Igf2* mRNA (red dots). Scale bars = 100 µm. **(E)** UMAPs of the 14 DPA regenerative and non-regenerative *Pdgfra*-positive mesenchymal cell transcriptomes (as in Fig. 4A), overlaid to show expression of the 5 selected ligand mRNAs, *Bmp4, Bmp5, Gdf6, Fgf10* and *Igf2.* The panel on the left is color-coded to show the regenerative (green) versus non-regenerative (pink) 14 DPA digit tip transcriptomes. **(F)** Violin plots showing relative expression of the five selected ligand mRNAs, *Bmp4, Bmp5, Gdf6, Fgf10* and *Igf2* in 7 DPA *Lmx1b-DTA* (Ablated) versus control *Pdgfra*-positive mesenchymal cell transcriptomes subsetted from the dataset shown in Fig. 4I. Each dot corresponds to expression in an individual cell. **(G, H)** Representative high-magnification images of regenerating 7 DPA digit tip sections immunostained for PDGFRα to identify mesenchymal cells (red) and for either total SMAD4 protein (G, red) or for the phosphorylated, activated form of SMAD4 (pSMAD4, H, red). Sections were also counterstained for DAPI (white) to show all cellular nuclei. Blue arrowheads in the right merged panels denote double-labelled cells. Scale bars = 20 µm.

This analysis identified 5 ligands that fulfilled our criteria, *Fgf10, Igf2*, and three members of the BMP/TGFβ family, *Bmp4, Bmp5* and *Gdf6*, as follows. First, analysis of the 7 DPA scRNA-seq regenerating digit tip dataset showed that all five ligands were expressed in mesenchymal cells at levels similar to or greater than other cell types, with the caveat that epithelial cells could not be assessed due to low cell numbers (Fig. 5B). Second, gene overlays on the 5-10 DPA mesenchyme-only spatial transcriptomic UMAP showed these ligands were also highly enriched in the regenerative versus nonregenerative mesenchyme (Fig. 5C). Third, high-resolution Xenium Explorer spatial analysis confirmed all five ligands were expressed in the nail mesenchyme and blastema cells but not epidermal cells at 7 DPA (Fig. 5D). Fourth, ligand mRNA overlays on the 14 DPA mesenchyme-only spatial transcriptomic UMAP showed the five ligands were still enriched in the regenerative versus nonregenerative mesenchyme (Fig. 5E). Moreover, Xenium Explorer analysis confirmed these expression patterns and showed *Gdf6* was relatively specific to the nail mesenchyme at 14 DPA (Fig. S5B). Finally, we asked if ligand expression was dependent upon the nail mesenchyme by analyzing the 7 DPA *Lmx1b-DTA* single cell spatial transcriptomic dataset. This analysis demonstrated that expression of all five ligands was decreased in the regenerative mesenchyme following partial nail mesenchyme ablation (Fig. 5F). Thus, establishment of a regeneration-specific ligand environment requires the presence of the nail mesenchyme.

### Mesenchymal cell BMP family signaling via SMAD4 is essential for blastema initiation and digit tip regeneration

Of the 5 ligands fulfilling our criteria, three (BMP4, BMP5 and GDF6) bind to and activate BMP receptors. Notably, exogenous application of BMPs 2 and 7 following non-regenerative digit tip amputations in neonates is sufficient to enhance skeletogenesis^37^. We therefore asked if this ligand family was essential for initiating and/or maintaining a regenerative response. Since there are many BMP and BMP receptor family members, we chose to do this by acutely knocking-out the transcription factor SMAD4, which is a shared essential downstream signaling protein for all BMP/TGFβ family ligands^38–40^. Prior to doing this, we confirmed that *Smad4* mRNA was expressed in 7 and 14 DPA regenerating mesenchymal cells using the scRNA-seq and spatial transcriptomic datasets (Fig. S5C-E), and showed that nuclear SMAD4 protein was detectable in most PDGFRα-positive regenerating mesenchymal cells by immunostaining 7 DPA digit tip sections (Fig. 5G). We also immunostained 7 DPA regenerating digit tip sections for the phosphorylated form of SMAD4; BMP receptor activation causes SMAD4 phosphorylation and nuclear translocation, and phospho-SMAD4 thus indicates pathway activation. The majority of PDGFRα-positive mesenchymal cells were positive for phospho-SMAD4 (Fig. 5H), consistent with a BMP/TGFβ-rich environment in the regenerating digit tip.

We therefore asked if mesenchymal cell SMAD4 activation was essential for regeneration using a mouse line carrying *Pdgfra-CreERT2* and two copies of a floxed *Smad4* allele (called *Pdgfra*-*Smad4KO* mice). When these crossed mice are treated with tamoxifen this causes inducible knockout of *Smad4* in *Pdgfra*-positive mesenchymal cells. As controls, we performed similar experiments with mice that carried *Pdgfra*-*CreERT2*, but were wildtype for *Smad4* (called *Pdgfra-WT* mice). In both cases we treated mice with tamoxifen, performed regenerative amputations after 7 days, and characterized regeneration at 14 DPA.

We first confirmed that tamoxifen treatment caused inducible knockout of *Smad4* in *Pdgfra*-*Smad4KO* mice by immunostaining 14 DPA sections for PDGFRα and SMAD4 (Fig. 6A). Relative to *Pdgfra*-*WT* digits, most mesenchymal cells in the *Pdgfra-Smad4KO* digits were negative for SMAD4 immunoreactivity, confirming the efficacy of the knockout. We then quantified the extent of tissue regeneration at 14 DPA using three measurements. First, we quantified regenerative tissue area in digit tip midline sections immunostained for PDGFRα (Fig. 6B, C). Exclusive of the epidermis, regenerating tissue area was decreased approximately 4-fold in the 14 DPA *Pdgfra*-*Smad4KO* digits. Second, we measured size of the regenerated nails; there was a significant approximately two-fold decrease in *Pdgfra-Smad4KO* nail length at 14 DPA (Fig. S6A). Finally, we performed three-dimensional imaging of cleared, unsectioned 14 DPA digit tips. To do this, we utilized *Pdgfra-Smad4KO* and *Pdgfra-WT* mice that also carried a floxed *Tdt* allele. We tamoxifen-treated and amputated the mice as for the other experiments to induce *Smad4* knockout and *Tdt* expression in *Pdgfra*-positive mesenchymal cells, and then cleared and performed lightsheet microscopy on the 14 DPA digit tips (Video S5 and S6). Quantification demonstrated that the volume of regenerated tissue was decreased more than 5-fold in the *Pdgfra-Smad4KO* digit tips (Fig. 6D, E).

**Figure 6.**
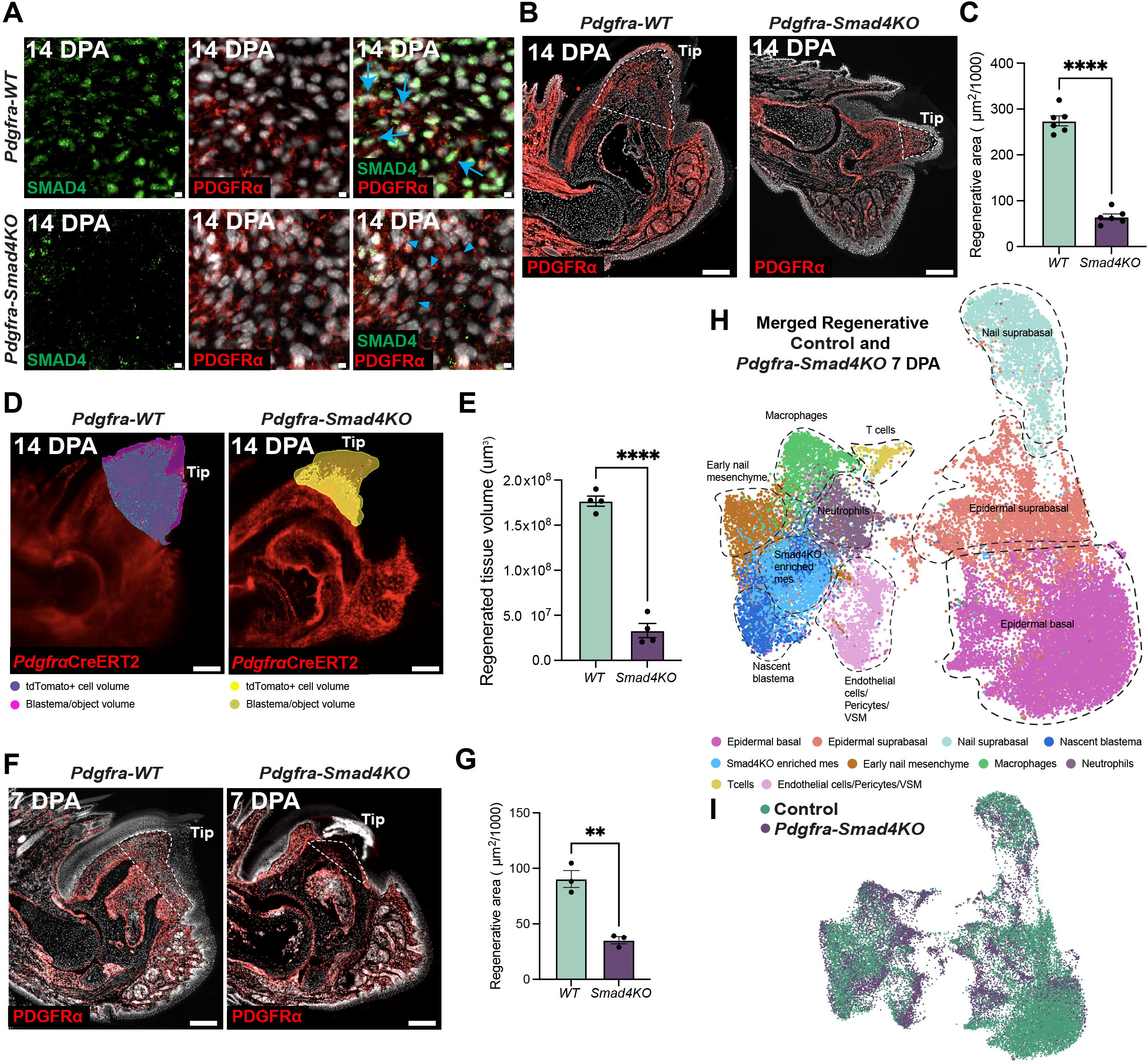
*Mesenchymal-specific deletion of Smad4 leads to deficits in digit tip regeneration at both 7 and 14 DPA*. (Also see Fig. S6 and Video S5 and S6). *Pdgfra*-*Smad4KO* or *Pdgfra-WT* mice were treated with tamoxifen, regenerative amputations were performed after 7 days, and digit tips were characterized after a further 14 days (14 DPA, A-E) or 7 days (7 DPA, F-I). **(A)** Representative high-magnification images of 14 DPA regenerating digit tip sections from *Pdgfra*-*Smad4KO* or *Pdgfra-WT* mice that were immunostained for PDGFRα to identify mesenchymal cells (red) and for total SMAD4 protein (green). Sections were also counterstained for DAPI (white) to show all cellular nuclei. Blue arrows denote double-labelled cells and arrowheads cell positive for PDGFRα but negative for SMAD4. Scale bars = 20 µm. **(B)** Representative images of 14 DPA midline sections from *Pdgfra*-*Smad4KO* or *Pdgfra-WT* mice that were immunostained for PDGFRα (red), and counterstained with DAPI (white) to highlight cellular nuclei. The hatched lines demarcate the regenerated tissue exclusive of the epithelium. Scale bars = 200 µm. **(C)** Quantification of digit tip sections as in (B) for the total regenerated area exclusive of the epithelium. n = 6 mice, ****p< 0.0001. **(D)** Representative midline optical sections from lightsheet-generated three-dimensional videos (Video S5 and S6) of cleared whole digit tips of *Pdgfra*-*Smad4KO* or *Pdgfra-WT* mice. The mice also carried a floxed *Tdt* reporter allele to allow visualization of TdT-positive mesenchymal cells (red). The masks show the regenerated tissue volume and the TdT-positive mesenchymal tissue area, color-coded as per the adjacent legends. Scale bars = 200 µm. **(E)** Quantification of lightsheet videos (such as those shown in Videos S5 and S6) for the volume of regenerated tissue in 14 DPA digit tips from *Pdgfra*-*Smad4KO* versus *Pdgfra-WT* mice. n = 4 mice, with one digit imaged per mouse. ****p< 0.0001. **(F)** Representative images of 7 DPA midline sections from *Pdgfra*-*Smad4KO* or *Pdgfra-WT* mice that were immunostained for PDGFRα (red) and counterstained with DAPI (white) to highlight cellular nuclei. The hatched lines demarcate the regenerated tissue exclusive of the epithelium. Scale bars = 200 µm. **(G)** Quantification of digit tip sections as shown in (F) for the total regenerated area exclusive of the epithelium. Each dot represents an individual mouse, with three midline sections per mouse quantified, and the average area plotted. n = 3 each, **p=0.0026. **(H)** UMAP showing merged 7 DPA *Pdgfra-Smad4KO* and control (from the dataset in Fig. S2A) digit tip transcriptomes as acquired using Xenium-based single cell spatial transcriptomics, annotated and color-coded for cell types. **(I)** UMAPs as in (H), color-coded to highlight 7 DPA *Pdgfra-Smad4KO* versus control transcriptomes.

We asked when this regenerative deficit first appeared by performing similar experiments at 7 DPA. Immunostaining of midline digit tip sections for PDGFRα at this earlier timepoint identified a 2 to 3-fold decrease in *Pdgfra*-Smad4KO regenerative tissue area relative to *Pdgfra-WT* controls (Fig. 6F, G). We then asked about the cellular basis of this deficit by performing Xenium-based single cell spatial transcriptomics on the 7 DPA *Pdgfra*-*Smad4KO* digit tips, analyzing sections from 4 independent animals. Analysis of the resultant dataset identified epithelial cells, *Pdgfra*-positive mesenchymal cells, vasculature endothelial and mural cells, Schwann cells, immune cells and osteoclasts (Fig. S6B), as was seen in the other regenerative datasets. We therefore merged the 7 DPA *Pdgfra*-*Smad4KO* and control datasets to directly compare the two conditions, using marker genes to identify the different cell types (Fig. 6H, I).

UMAP and spatial plot analysis of this merged dataset identified several major changes in the *Pdgfra-Smad4KO* condition relative to controls. In particular, as seen in the morphological analyses (Fig. 6F, G), there was much less tissue between the epidermis and the amputated bone and overall tissue regeneration was reduced (Fig. 7A). Nonetheless, some cell types were relatively unaffected transcriptionally since immune cells and vasculature-associated endothelial and mural cells from the two groups co-clustered transcriptionally (Fig. 6H, I) and their localization was roughly similar (Fig. 7A-C). However, in the nonregenerative condition, the density of macrophages and neutrophils was significantly increased by approximately two-fold (Fig. 7D).

**Figure 7.**
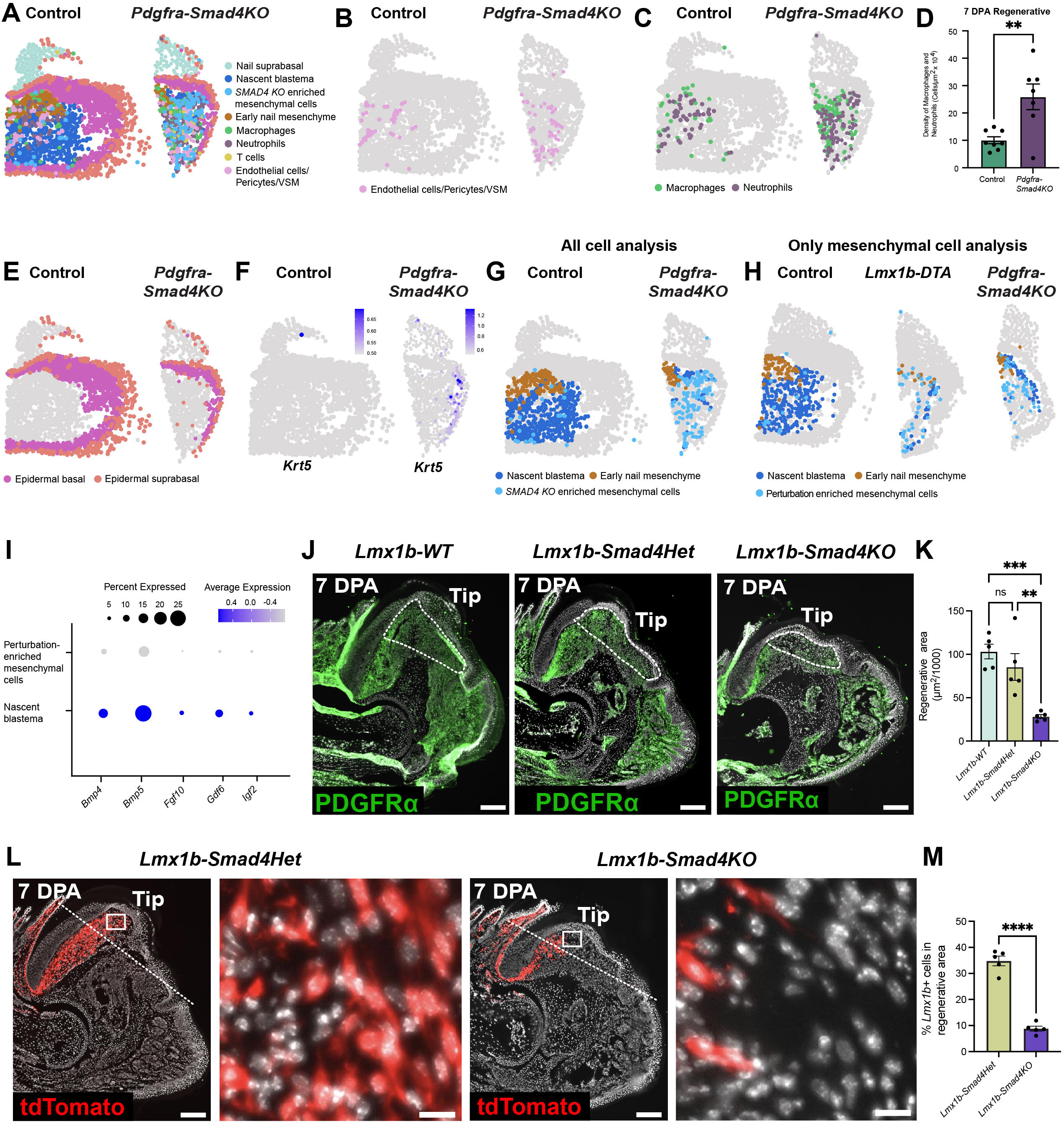
*(A-G) Loss of Smad4 in all mesenchymal cells perturbs multiple aspects of digit tip regeneration by 7 DPA.* (Also see Fig. S6 and Table S6). **(A)** Spatial plots of the merged 7 DPA *Pdgfra-Smad4KO* and control digit tip transcriptomes from the dataset in Fig. 6H, showing all detected cell types on representative 7 DPA *Pdgfra-Smad4KO* and control midline sections. Cell types are color-coded as per the adjacent legend. **(B-D)** Spatial plots of the same representative 7 DPA *Pdgfra-Smad4KO* and control sections as in (A), showing (B) vasculature-associated endothelial and mural cells and (C) macrophages and neutrophils. **(D)** Graph showing the density of macrophages/neutrophils in regenerating 7 DPA *Pdgfra-Smad4KO* and control digit tip midline sections as in (C). Quantification was performed using high resolution Xenium Explorer images, and each dot represents an individual section. n = 3-4 mice per group. **p<0.01. **(E)** Spatial plots of the same representative 7 DPA *Pdgfra-Smad4KO* and control sections as in (A), showing the different epithelial cell populations. **(F)** Spatial plots of the same representative 7 DPA *Pdgfra-Smad4KO* and control sections as in (A), overlaid for expression of *Krt5* mRNA, color-coded as per the adjacent keys. **(G)** Spatial plot of the same 7 DPA *Pdgfra-Smad4KO* and control sections as in (A), showing transcriptionally-distinct mesenchymal cell populations, including the nail mesenchyme, nascent blastema, and a population of mesenchymal cells that was proportionately enriched in the *Pdgfra-Smad4KO* digit tips. **(H)** Spatial plots of the merged 7 DPA regenerative *Lmx1b-DTA*, *Pdgfra-Smad4KO*, and control mesenchymal cell transcriptomes from the dataset in Fig. S6C, showing transcriptionally-distinct mesenchymal cell clusters, including the nail mesenchyme, nascent blastema, and a population of mesenchymal cells that was proportionately enriched in both the *Lmx1b-DTA* and *Pdgfra-Smad4KO* digit tips (perturbation-enriched mesenchymal cells). Note these are different control and *Pdgfra-Smad4KO* sections from those shown in (A-G). **(I)** Dot plot showing expression levels of *Bmp4, Bmp5, Gdf6, Igf2* and *Fgf10* mRNAs in the 7 DPA nascent blastema cells versus perturbation-enriched mesenchymal cells, as determined from the merged Xenium single cell transcriptomic data in Fig. S6C. The size of the dot indicates the percentage of cells detectably expressing the mRNA, and the color indicates relative expression level, coded as per the adjacent key. *(J-M) Loss of Smad4 specifically in the nail mesenchyme is sufficient to inhibit digit tip regeneration by 7 DPA*. *Lmx1b*-*Smad4KO, Lmx1b-Smad4Het* or *Lmx1b-WT* mice were treated with tamoxifen, regenerative amputations were performed after 7 days, and digit tips were characterized after a further 7 days (7 DPA). **(J)** Representative images of 7 DPA midline sections from *Lmx1b*-*Smad4KO, Lmx1b-Smad4Het* or *Lmx1b-WT* mice that were immunostained for PDGFRα (green), and counterstained with DAPI (white) to highlight cellular nuclei. The hatched lines demarcate the regenerated tissue exclusive of the epithelium. Scale bars = 200 µm. **(K)** Quantification of digit tip sections as in (J) for the total regenerated area exclusive of the epithelium. Each dot represents an individual mouse, with three midline sections per mouse quantified, and the average area plotted. n = 5 mice per group, **p< 0.01, ***p< 0.001, ns = nonsignificant. **(L)** Representative images of 7 DPA midline sections from *Lmx1b-Smad4Het* or *Lmx1b-Smad4KO* mice that also carried a floxed *Tdt* reporter allele to allow visualization of TdT-positive nail mesenchyme cells and their progeny (red). Sections were counterstained with DAPI to visualize cellular nuclei. The hatched lines denote the amputation planes, and the boxed regions are shown at higher magnification to the right of each lower magnification image. Scale bars = 200 µm. **(M)** Quantification of digit tip sections as in (L) for the percentage of TdT-positive cells within the regenerated tissue exclusive of the epithelium. Each dot represents an individual mouse, with three midline sections per mouse quantified, and the average percent plotted. n = 5 mice per group, ****p<0.0001.

Epidermal and mesenchymal cells were also perturbed in the *Pdgfra-Smad4KO* condition relative to controls, with deficits similar to those following nail mesenchyme ablation. Specifically, the *Pdgfra*-*Smad4KO* epidermal basal cells clustered distinctly from the controls, and expressed higher levels of *Krt5* (Fig. 6H, I; Fig. 7E, F), indicative of perturbed re-epithelialization. As for mesenchymal cells, in the *Pdgfra*-*Smad4KO* digit tips they were reduced and spatially disorganized (Fig. 7G), with fewer nail mesenchyme and blastema cells. Instead, many of the *Pdgfra-Smad4KO* mesenchymal cells were transcriptionally distinct from the control blastema cells, as indicated by the segregation of these two populations in the clustering UMAP (Fig. 6H, I; Fig. 7G). Thus, when SMAD4 signaling is lost in mesenchymal cells, this results in deficient nail mesenchyme regeneration, decreased genesis of blastema cells and perturbed re-epithelialization.

Since the 7 DPA *Pdgfra-Smad4KO* mesenchymal cell deficits were similar to those seen following nail mesenchyme ablation, we merged mesenchymal cell transcriptomes from the 7 DPA *Lmx1b-DTA*, *Pdgfra-Smad4KO* and control datasets to allow a direct comparison. UMAP and spatial plot analyses (Fig. 7H; Fig. S6C) identified, in addition to bone cells, three distinct types of mesenchymal cells; the nail mesenchyme, nascent blastema cells and a third population that was proportionately enriched in the *Lmx1b-DTA* and *Pdgfra-Smad4KO* conditions. Differential gene expression analysis showed that these perturbation-enriched cells expressed significantly less of all five ligands, *Fgf10*, *Igf2, Gdf6, Bmp4* and *Bmp5* relative to the nascent blastema cells (Fig. 7I; Table S6). Thus, in both the *Lmx1b-DTA* and *Pdgfra-Smad4KO* 7 DPA digit tips, nail mesenchyme and nascent blastema cells are depleted, and there is a proportionate increase in transcriptionally-distinct mesenchymal cells that are deficient in expression of regeneration-specific ligands.

### Loss of Smad4 signaling specifically in the nail mesenchyme inhibits nail mesenchyme and digit tip regeneration

These results indicate that BMP-SMAD4 signaling is essential for nail mesenchyme regeneration and suggest that the other phenotypes may be due, at least in part, to the nail mesenchyme deficit. To directly test this idea, we performed a more selective deletion of the *Smad4* gene specifically in the nail mesenchyme, crossing mice carrying the floxed *Smad4* and *Tdt* alleles to the *Lmx1bCreERT2* mice. Tamoxifen-mediated induction of CreERT2 in these *Lmx1b-Smad4KO* mice causes nail mesenchyme-specific knockout of *Smad4*, and allows lineage tracing of nail mesenchyme cells and their progeny. As controls, we analyzed *Lmx1bCreERT2* mice that did not carry the floxed *Smad4* or *Tdt* alleles (*Lmx1b-WT*) or mice carrying the floxed *Tdt* allele and one floxed *Smad4* allele (*Lmx1b-Smad4Het*). We treated all groups with tamoxifen, performed regenerative amputations, and analyzed the digit tips at 7 DPA.

Initially we asked whether nail mesenchyme-specific *Smad4* knockout affected the extent of digit tip regeneration, quantifying regenerated tissue on midline digit tip sections immunostained for PDGFRα. This analysis (Fig. 7J, K) demonstrated that 7 DPA regenerative tissue area was similar in the *Lmx1b-WT* and *Lmx1b-Smad4Hets* digit tips, but was decreased 3 to 4-fold in the *Lmx1b-Smad4KOs*. We then took advantage of the TdT reporter to specifically visualize nail mesenchyme cells and their progeny. In the 7 DPA *Lmx1b-Smad4Het* regenerating tissue almost 35% of cells were TdT-positive, and these were not just limited to the dorsal region immediately under the nail (Fig. 7L, M). By contrast, the percentage of TdT-positive regenerating cells was reduced 3 to 4-fold in the *Lmx1b-Smad4KO* digit tips, and these were more spatially-restricted (Fig. 7L, M). Thus, SMAD4 signaling within the nail mesenchyme is essential for nail mesenchyme regeneration, and when this is disrupted, other aspects of digit tip regeneration are also perturbed.

## Discussion

One key biological question is why a few privileged mammalian tissues have maintained the evolutionary ability to regenerate following injury. Here, we have addressed this question by studying the digit tip, which regenerates in both adult mice and humans. Our findings argue that the digit tip’s regenerative ability is due to the presence of the nail mesenchyme, a signaling center and inductive mesenchymal tissue that persists into adult life in order to instruct homeostatic nail growth. In particular, our data indicate that the nail mesenchyme promotes regeneration by creating a BMP-rich ligand environment that facilitates reprogramming of local tissue-resident mesenchymal cells and in so doing initiates blastema formation. When the nail organ/mesenchyme are absent and/or perturbed, this regeneration-specific environment does not occur, and mesenchymal cells do not reprogram to a blastema state but instead undergo bone callus or fibrotic wound healing. Moreover, perturbation of the nail mesenchyme and/or blastema formation alters re-epithelialization arguing that mesenchymal cells coordinate a highly-organized multicellular regenerative response. Thus, the digit tip is likely privileged with regard to regeneration because it is one of the few sites in the body where a developmental signaling center has persisted and can create the necessary pro-regenerative environment. If we can fully define this regeneration-specific ligand environment, as we have started to do here, then perhaps we can use that information to promote repair and/or prevent inappropriate fibrotic responses in other tissues.

Based upon our findings, we propose that regeneration-specific ligands, including the BMPs, are sufficient to reprogram tissue-resident adult mesenchymal cells to a blastema state. How and why does this reprogramming happen? Potential clues come from considering previously-defined cellular transitions that occur during regeneration. In particular, lineage tracing studies have shown that the regenerative blastema is largely comprised of mesenchymal cells that originate from multiple sources including the periosteum/bone, the mesenchymal components of local injured nerves, and the nail mesenchyme, all of which are transcriptionally very distinct from each other^8,9,18,41,42^. Regardless of their tissue-of-origin, these cells lose expression of genes associated with their differentiated states, acquire a blastema state involving genes important for both development and tissue repair, and then contribute to bone and dermis regeneration^8^. We posit that the reason mesenchymal cells so readily undergo this reprogramming is because they are amongst the most “plastic” of differentiated mammalian cell types. Indeed, it has long been known that when mesenchymal cells such as dermal fibroblasts are cultured under the appropriate conditions, they can generate adipocytes, chondrocytes and osteoblasts^21,43–49^. Moreover, even *in vivo*, mesenchymal cells are known to, for example, undergo inappropriate osteogenesis or adipogenesis in response to environmental perturbations^50–56^. Thus, it seems likely that following regenerative digit tip amputations, at least a subset of local mesenchymal cells are in a flexible/plastic state that is receptive to ligands such as nail mesenchyme-derived BMPs that reprogram them to a blastema state. It may well be that this same cellular flexibility results in fibrosis when mesenchymal cells are exposed to a less positive environment such as non-regenerative injuries and/or persistent tissue inflammation.

Our data identify five phenotypes that occur when *Smad4* is acutely knocked-out in all mesenchymal cells; decreased nail mesenchyme regeneration, aberrant blastema cell reprogramming, decreased expression of regeneration-specific ligands, perturbed re-epithelialization and decreased net regenerative growth. At least some of these perturbations, the deficits in nail mesenchyme regeneration and total regenerative growth, are also seen when *Smad4* is knocked-out specifically in the nail mesenchyme. Thus, SMAD4-mediated signaling in nail mesenchyme cells is essential for their regeneration, and loss of nail mesenchyme regeneration may be sufficient to at least partially cause the other phenotypes, as seen in the ablation studies. These findings are consistent with a key role for BMPs within the nail mesenchyme, but it is important to note that SMAD4 is downstream of both BMP and TGFβ family receptors. Nonetheless, several compelling arguments suggest that the pro-regenerative SMAD4 effects we document here involve BMPs and not TGFβs. First, previous work showed that exogenous local application of BMPs 2 and 7 was sufficient to enhance skeletogenesis following non-regenerative digit tip amputations in neonatal mice^27^. Second, BMPs are pro-regenerative in other vertebrates, including, for example, during dedifferentiation of regenerating newt limb mesenchymal cells^59^. Third, TGFβ family ligands and receptors are widely expressed in both the regenerative and non-regenerative digit tips (data not shown), whereas we show here that several BMPs are selectively expressed during regeneration. Finally, TGFβs have been broadly-implicated in fibrosis^60–63^, whereas the BMPs are generally thought to be anti-fibrotic^27,64,65^. We therefore propose that mesenchymally-derived BMPs are important for both nail mesenchyme regeneration and blastema formation.

While our studies have focused on the BMPs, they do not rule out other potential pro-regenerative ligands. Indeed, two of the regeneration-specific ligands we identified were FGF10 and IGF2. Notably, mesenchymal FGF10 is required for embryonic limb development^66,67^, and IGF2 regulates bone growth throughout life^68,69^. It may be that ligands like FGF10 and IGF2 act in cooperation with the BMPs to promote blastema formation and regeneration as they do during embryonic limb development^66,67,69^. It is also possible that epithelial cells contribute ligands to the regenerating digit tip environment; previous work showed that nail organ epithelial:mesenchymal interactions are important for digit tip regeneration, and have implicated Wnt signaling, Hedgehog signaling and Cxcr4 in these interactions^14,70–72^. Relevant to this, our Xenium analyses indicate that epithelial cells are the predominant source of BMPs 7 and 8a within the regenerating digit tip (data not shown). Thus, it may be that the BMPs important for nail mesenchyme regeneration come from both the nail mesenchyme itself and from the adjacent nail epithelium.

In this study, we predominantly focused on the cellular transitions that underpin blastema formation and regeneration. However, our Xenium-based analyses also speak to patterning of the newly-formed digit tip, another key aspect of regeneration. In particular, we show that from 7 DPA on, there are major differences in the shape of the regenerated tissue in the presence and absence of the nail organ; in the regenerative condition the tissue grows distally while in the non-regenerative, it expands laterally as much or more than it does distally. In addition, there are differentiation gradients within the regenerating digit tip. At 10 DPA the first osteogenic differentiation occurs immediately adjacent to the amputated bone and by 14 DPA the osteogenic domain has expanded distally, while the first dermal differentiation is observed adjacent to the distal epidermis at 14 DPA. Thus, we posit there are local differentiation cues originating from the three major spatially-segregated digit tip components, the bone, the epidermis, and the nail organ. Since the regenerating mouse digit tip is very small, then these local cues may play a major role in shaping the structure of the regenerating tissue. This does not preclude other potential patterning mechanisms such as those seen during digit/limb development^73–76^ or amphibian limb regeneration^77,78^. However, it does suggest that perhaps the digit tip is regeneration-privileged in part due to its small size and well-defined tissue architecture.

Our findings have implications for aberrant mesenchymal tissue repair and pathological fibrosis. Specifically, our data argue that the mesenchymal cell decision to repair/regenerate rather than undergo wound-healing or fibrosis is driven by the local ligand environment. When digit tip bone and dermal cells are exposed to a regeneration-specific ligand environment, they acquire a blastema fate and rebuild the lost tissues, but when this positive environment is absent due to loss of the nail organ or nail mesenchyme, they instead undergo a wound-healing scarring response. Moreover, when dermal fibroblasts are transplanted into the regenerating digit tip, they contribute to the blastema and ultimately regenerating bone, but when transplanted into the non-regenerative digit tip, they instead contribute to the mesenchymal scar cap^8^. In this regard, many previous studies have argued that a negative environment, largely as determined by an inflammatory immune response, is responsible for pathological fibrosis^79–83^. Here, we show that the density of macrophages/neutrophils is higher in the non-regenerative digit tip from 5 to 10 DPA. However, while fewer in number, many immune cells are still present in the regenerating condition, and the monocyte lineage plays a positive role during digit tip regeneration^84^, arguing that if these immune cells do provide negative cues, then the pro-regenerative environment created by the nail organ is sufficient to override them. Together, our findings identify pro-regenerative ligands that can potentially promote mesenchymal repair and inhibit fibrosis, thereby paving the way for new pro-repair and anti-fibrosis strategies.

## Acknowledgements

This work was funded by grants from the Canadian Institutes of Health Research (PJT-180308 to FDM and DRK, and FDN-159908 to FMVR) and SP had a Michael Smith Health Research BC fellowship. We thank Dr. Jasmine Yang for her technical assistance.

## Author contributions

SP, SS and CE conceptualized, performed and analyzed experiments and co-wrote the manuscript. SP performed and analyzed the regenerative, non-regenerative and nail mesenchyme ablation Xenium experiments, including the mergers, and some of the light sheet experiments. SS performed and analyzed most of the immunocytochemical analyses and lightsheet experiments, and performed and analyzed the SMAD4 knockout experiments, including the Xenium analyses. CE collected and analyzed the scRNA-seq datasets, performed the ligand-receptor modeling, and performed some of the immunocytochemical analyses. KK collected the scRNA-seq datasets with CE. FDM, DRK and FMVR all conceptualized experiments and co-wrote the manuscript, and FDM also analyzed data.

## Materials and Methods

### Resource Availability

#### Lead contact

Further information and requests for resources and reagents should be directed to and will be fulfilled by the lead contact, Freda Miller.

#### Data availability

Previously published scRNA-seq datasets (Uninjured 1, Uninjured 2, 7DPA regenerative 1, 10DPA regenerative, 14DPA regenerative 1, 14DPA regenerative 2, *Lmx1bCreERT2;TdT* 14DPA regenerative, *PdgfraCreERT;TdT* 14DPA regenerative, *Dmp1CreERT;TdT* 14DPA regenerative, 28DPA regenerative, and 56DPA regenerative) are available from GEO (GEO:GSE135985 and GEO:GSE217600). New scRNA-seq datasets (Uninjured 3, 4DPA regenerative 1, 4DPA regenerative 2, 7DPA regenerative 2) and Xenium datasets (5 DPA regenerative, 7 DPA regenerative, 10 DPA regenerative, 14 DPA regenerative, 5 DPA non-regenerative, 7 DPA non-regenerative, 10 DPA non-regenerative, 14 DPA non-regenerative, 7 DPA regenerative *Lmx1b-DTA* and 7 DPA regenerative *Pdgfra-Smad4KO*) are also available from GEO (GEO:GSE326776 and GEO:GSE326786). Any additional information required to reanalyze the data reported in this paper is available from the Lead Contact upon request.

### Experimental Model and Subject Details

#### Mice

All mice used in this study were maintained in accordance with Canadian Council on Animal Care guidelines, with approval from the Animal Care Committees of the Hospital for Sick Children and the University of British Columbia. In all studies, mice were maintained under a 12-h light/dark cycle with unrestricted access to rodent chow and water. All animals were healthy with no observable behavioral abnormalities. Adult mice aged 8–12 weeks, of either sex, were used and randomly assigned to experimental groups. Wild type C57BL6 mice were purchased either from Charles River Laboratories (strain codes: 027) or from Jackson Laboratories (JAX stock #000664). The following transgenic mouse lines were obtained from Jackson Laboratories and were genotyped as recommended: *Lmx1bCreERT2* (*Lmx1b^tm2(cre/ERT2)Rjo^/J*; JAX stock #031289)^85^; *PdgfraCreERT2* (B6.129S-*Pdgfra^tm^*^1.1^*^(cre/ERT^*^2^*^)Blh^*/J; Jax stock #032770)^35^; *R26-LSL-TdTomato* (Cg-*Gt(ROSA) 26Sor^tm^*^9^*^(CAG-tdTomato)Hze^/*J; JAX stock #007909)^86^; *R26-LSL-DTA* (129-Gt(ROSA) *26Sor^tm^*^1^*^(DTA)Mrc^/*J; JAX stock #010527)^87^; *Cdh5-Cre* (*B6.FVB-Tg(Cdh5-cre)7Mlia/J*; JAX stock #006137)^26^ and *Smad4^Flox^*(*Smad ^tm^*^2^*^.1Cxd^/J*, JAX Stock #017462)^88^.

#### Animal surgeries and tamoxifen injections

We performed mouse digit tip amputations as previously described^4,8,18^. Briefly, 8–12-week-old mice were anesthetized and administered meloxicam (5 mg/kg) subcutaneously for analgesia. The distal portion of the third phalanx (P3) was amputated to induce a regenerative response, whereas complete removal of the nail bed was performed to generate a non-regenerative response by amputating through the second phalanx (P2) (see Fig. S1A). For transgenic mice carrying an inducible *Creert2* allele, tamoxifen (30 mg/mL; Sigma) was dissolved in sunflower oil and administered via intraperitoneal injection for five consecutive days. Amputation procedures were performed seven days after the final tamoxifen injection. For diphtheria toxin A (DTA) ablation experiments, tamoxifen was administered intraperitoneally for seven consecutive days, and digit amputation was performed the following day. Digits were harvested at the indicated time points according to the specific experimental requirements.

### Method Details

#### Tissue preparation, immunostaining and microscopy

Harvested digits were fixed in 4% paraformaldehyde (PFA) for 24 h at 4°C, followed by decalcification in 0.5 M EDTA (pH 7.0) for 14 days at room temperature. After decalcification, the tissues were cryoprotected in 30% sucrose for 24 h, embedded in optimal cutting temperature (O.C.T.) compound, and snap-frozen. Samples were sectioned sagittally at a thickness of 14 μm using a cryostat. Sections were air-dried in a 37°C incubator for 30 min and subsequently washed in phosphate-buffered saline (PBS) for 10 min. Tissue sections were blocked for 1 h at room temperature in blocking buffer containing 5% bovine serum albumin (BSA; Jackson ImmunoResearch) and 0.3% Triton X-100 (Fisher Scientific) in PBS. Appropriate secondary antibodies (1:500 - 1:1000 dilution in 5% BSA in PBS) were applied and incubated for 16–24 h at 4°C. Sections were washed three times in PBS for 5 min each. Nuclei were counterstained with DAPI (Thermo Fisher Scientific) for 5 min, followed by PBS washing. Slides were then air-dried and mounted using Vectashield mounting medium.

Fluorescent images were acquired using either a Zeiss LSM900 confocal microscope equipped with an Airyscan 2 detector (ZEN 3.4 software) or a Zeiss AxioImager M2 system equipped with either an X-Cite 120 LED light source and a Hamamatsu C11440 camera (ZEN acquisition software) or an Illuminator microLED light source and an Axiocam 712 mono camera (ZEN 3.1 software). Optical sections were acquired at a thickness of 0.3–1 μm and processed as z-stacks. Images were stitched where necessary and further analyzed using ImageJ software.

#### Antibodies

Primary antibodies used were as follows: goat anti-PDGFRα ([1:100 or 1:250], R & D Systems; cat#AF1062); goat anti-CD31 ([1:100], R&D Systems; cat# AF3628); rat anti-mouse CD68 ([1:250], Invitrogen; cat#14-0681-82); rabbit anti-SMAD4 ([1:200], Cell Signaling Technology; cat#46535); rabbit anti-phospho-SMAD4 ([1:200], Affinity Biosciences; cat# AF8316). Fluorescently labeled highly cross-absorbed secondary antibodies were purchased from Jackson ImmunoResearch or Invitrogen and used at a dilution of 1:500 - 1:1000.

#### Tissue clearing and light sheet microscope

The SHIELD method^89^ was used for tissue clearing of digits (according to Life Canvas Technologies protocol). Mice subjected to regenerative or non-regenerative amputations (described above) were anesthetized with 2,2,2-tribromoethanol (Avertin). Animals were transcardially perfused with 20 mL of 2 mM EDTA in PBS, followed by 20 mL of 4% PFA. Digits were dissected and fixed overnight in 4% PFA at 4°C. The following day, digits were washed with PBS and nails were removed. Samples were decalcified in 0.5 M EDTA (pH 7.0) at 4°C for 4 days. Decalcified digits were then incubated in fresh SHIELD OFF solution for 4 days at 4°C in the dark. Subsequently, digits were transferred to SHIELD ON solution and incubated for 24 hours at 37°C. For tissue clearing, samples were incubated in delipidation buffer at 37°C for 7 days. After clearing, digits were washed overnight at 37°C in PBST (1% Triton X-100 in PBS). Samples were then refractive index–matched in 100% Easy Index solution (RI=1.465 or 1.52) for 24 hours.

Imaging was performed using a Zeiss Lightsheet Z.1 microscope with ZEN Black 2014 software. Images were acquired using the standard chamber (5x objective, NA 1.45) with a 0.7x zoom, 5x/0.16 NA detection optics, and 5x/0.1 NA illumination optics. Z-stacks were collected with an optical slice thickness of 5 μm. A 561 nm laser was used to detect red fluorescence (tdTomato), and a 488 nm laser was used to capture background signal (gray channel). Image analysis was performed using ImageJ, and 3D reconstructions were generated using Arivis Vision4D software (Version 3.5, Zeiss).

#### Single cell isolation and 10X Genomics single cell RNA sequencing

For the regenerative 4DPA (4DPAR1 - 13 mice, 78 digits; 4DPAR2 – 12 mice, 72 digits) and 7DPA (7DPAR2 12 mice – 72 digits) single cell datasets, regenerating digit tips distal to the original amputation plane were collected at 4DPA or 7DPA. Any skin or nail was removed, and the tissue was digested using Collagenase D (1.2U/ml)/Dispase II (2.4U/ml) at 37^°^C for 30-40 minutes, with constant agitation. Enzymatic digestion was halted by the addition of 5% serum and the cells were mechanically dissociated using a small-bore-hole glass pipette. Dissociated cells were passed through a 20um cell strainer and pelleted by centrifugation at 600*g* for 5 minutes. The cell pellet was then washed in 1X Hank’s balanced salt solution (1X HBSS) + 5% serum and re-filtered through a 20um cell strainer to remove any additional debris. Cells were pelleted by centrifugation at 600*g* for 5 minutes, and cells resuspended in 1X HBSS + 5% serum and counted using a hemocytometer. For the additional uninjured digit tip scRNA-seq dataset that was uploaded to GEO (GEO:GSE326786 - uninjured3), freshly dissected tissue was digested in Liberase (280mg/ml, Roche) at 37^°^C for 4 h with constant agitation and then processed as for the regenerating digit tips above. At least 150,000 cells were isolated for each experiment. 10X Genomics single cell RNA sequencing, including droplet collection, cDNA amplification and sequencing library preparation, was then carried out using the 10X Genomics Chromium system as per the manufacturer’s guidelines and sequenced on an Illumina NextSeq500 at the Princess Margaret Genomics facility (Toronto, ON) or The Hospital for Sick Children Centre for Applied Genomics (Toronto, ON). FASTQ sequencing reads were processed, aligned to the wild-type mouse genome (mm10) and converted to digital gene expression matrices using the Cell Ranger count function within the Cell Ranger Single-Cell Software Suite with settings as recommended by the manufacturer (https://support.10xgenomics.com/single-cell-gene-expression/software/overview/welcome).

#### scRNA-seq data analysis

Dataset count matrices were processed using a Seurat-based pipeline (Seurat version 5). Briefly, datasets were filtered to remove cells with low unique molecular identifier (UMI) counts, red blood cells, cells with high mitochondrial DNA content, and putative doublets. Cell transcriptomes were normalized using SCTransform V2^90^ to account for read depth and library size. Using SCTransform, mitochondrial genes and cell cycle scores calculated using Seurat’s CellCycleScoring function, were regressed out. Principal component analysis (PCA) was performed using the top 3000 highly variable genes and the top 30 principal components were used to generate both Uniform Manifold Approximation and Projection (UMAP) embeddings, and to iteratively carry out SNN-Cliq-inspired clustering using Seurat’s FindClusters function at resolutions ranging from 0.4 to 2.4. Each dataset was analyzed by choosing the most conservative resolution at which cell populations of known identity separated into distinct clusters. Higher resolutions were only used to examine cell heterogeneity.

Cell types were identified based on the expression of canonical markers. We used the following gene expression definitions: all mesenchymal cells expressed *Pdgfra*, *Ly6a*, and *Prrx2*; nail mesenchymal cells expressed *Lmx1b*, *Sfrp2*, *Rspo4*, and *Nrg2* in addition to the mesenchymal cell markers; pericytes/vascular smooth muscle (VSM) cells expressed *Rgs5*, *Myh11*, and *Pdgfrb*; endothelial cells expressed *Pecam1*, *Tek*, and *Cdh5*; immune cells expressed *Ptprc* or *Acp5*; schwann cells expressed *Sox10*, *Sox2*, *Plp1*; and epithelial cells expressed *Krt5*, *Krt14*, and *Krt10*.

UMAPs and gene expression overlays, generated using DimPlot and FeaturePlot functions, were used to investigate gene expression and interpret cell clusters. Heatmaps were generated using the DoHeatmap function.

For merged or subsetted datasets, barcodes corresponding to the cells of interest were used to extract the relevant gene expression data from the gene expression matrices. For merged datasets, the gene expression data from each relevant dataset was combined and the cells were run through the pipeline as described above.

#### Batch correction

For the 7 DPA regenerative merge, batch correction was performed to reduce batch effects that manifested between cells originating from different experiments within a Seurat object. Batch correction of the SCT normalized merged object was performed using Harmony^91^ via the Seurat IntegrateLayers function using the minimum number of iterations (2) required for the datasets to intermingle as expected. After using IntegrateLayers(method = HarmonyIntegration, max_iter = 2, normalization.method = ‘SCT’), the data was reclustered using the FindNeighbors and FindClusters functions at a range of resolutions, and new UMAP embeddings were generated using the RunUMAP function.

#### Trajectory inference using transcriptomic data

Trajectory inference and pseudotime ordering of scRNA-seq data was performed using Monocle 2 (V2.30.1) in R^96,97^. As Monocle 2 was designed to work with Seurat V3 objects, Seurat V5 assays were first converted to V3 assays to generate a Monocle cell data set (cds), which was then normalized using Monocle’s size factor normalization method via the estimateSizeFactors function. To perform PCA, variable genes were found by converting the Seurat object to a single cell experiment using as.SingleCellExperiment, normalizing counts using logNormalizeCounts, and modelling gene variance using modelGeneVar followed by the getTopHVGs function. Cells were projected into two-dimensional space using Monocle’s reduceDimension(max_components = 2, reduction_method = ‘tSNE’), and clusters assigned using Monocle’s density peak clustering algorithm using the clusterCells function. To determine genes used for pseudotime ordering, the top 1000 differentially expressed genes between clusters at the relevant resolution using the differentialGeneTest function. To regress out cell cycle related genes from trajectories generated using Monocle, genes used by Cyclone and Seurat to assign cell cycle phase were subtracted from the set of 1000 ordering genes. Expression profiles were reduced to two dimensions using a DDRTree algorithm via Monocle’s reduceDimension(max_components = 2, method = ‘DDRTree’) function and cells ordered using the cell cycle regressed differentially expressed genes to generate trajectories using the Monocle function orderCells. The function plot_cell_trajectory was used for visualizing cell trajectories.

#### Gene signature analysis

Scaled expression of multiple genes was calculated using the Seurat AddModuleScore function^92^. Module scores were calculated using the SCT normalized gene expression scores. To calculate the overall expression of the blastema gene signature, the 230 blastema genes identified in Storer et al.^8^ were used. To calculate the overall expression of the nail mesenchyme gene signature, the 15 nail mesenchyme signature genes identified in Mahmud et al.^18^ were used.

#### Ligand-receptor analysis

Ligand-receptor analysis was performed essentially as described in Willis et al. (2025) using a curated ligand-receptor database and the CCINX package^36^. Ligands were included when they were detectably expressed in >5% of the relevant cell types and receptors when detectably expressed in ≥20%. Cytoscape (v3.10.3, SCR_003032) was used to visualize the predictive ligand-receptor communication models, where ligands included in the model are presented in a central panel of nodes and edges connecting them to their source and target cell type^93^.

#### Xenium panel design

Two different custom gene panels that were targeted for mesenchymal cells and other digit tip cell types were used for the Xenium-based single cell spatial transcriptomic analyses, one targeting 480 genes (10x Genomics, 9XXV3X) and the other 473 genes (10x Genomics, UT7ADC). Genes were selected to distinguish digit tip cell types and transcriptional states based on previous literature and on our previously-published regenerative and non-regenerative digit tip scRNA-seq datasets^8^. The lists of genes in the two panels can be found in Table S2. The scRNA-seq datasets were also used during panel design to inform panel design and minimize the risk of optical crowding for each gene.

### 10X Genomics Xenium tissue preparation and data processing

Unless otherwise specified, C57BL/6 mice were analyzed in all experiments. Digit tip tissue was dissected close to the bone stump and collected into chilled PBS, including multiple digit tips per mouse from a minimum of three independent mice per condition. Digit tips were pooled, embedded in OCT compound, flash-frozen, and stored at -70°C. Sectioning was performed according to 10X Genomics guidelines. The cryostat chamber temperature was maintained at -23°C, and the specimen temperature was maintained at -26°C. Cryosections (10 μm thickness) were mounted onto Xenium slides (chemistry v1) and stored at -70°C until further processing.

Slides were processed according to the Xenium workflow for fresh-frozen tissue as previously-described^30–32^. On day 1, frozen slides were incubated at 37°C for 1 min and fixed in 4% PFA for 30 min at room temperature. Slides were washed with PBS for 1 min, followed by permeabilization and wash steps. Briefly, sections were incubated in 1% SDS for 2 min, washed twice with PBS, and then incubated in chilled 70% methanol for 1 h. Slides were washed twice with PBS and incubated in PBST (0.05% Tween-20 in PBS). Hybridization was performed using a custom 480-gene mesenchymal panel (10x Genomics, 9XXV3X) or a custom 473-gene mesenchymal panel (10x Genomics, UT7ADC) prepared in TE buffer for 18–22 hours at 50°C. The following day, slides were washed three times with PBST (1 minute each) and incubated in post-hybridization wash buffer for 30 minutes at 37°C. Slides were again washed three times with PBST (1 minute each) and incubated in Xenium ligation enzyme mix for 2 hours at 37°C. After three additional PBST washes (1 minute each), slides were incubated in Xenium amplification enzyme solution for 2 h at 30°C. Slides were washed twice with TE buffer (1 minute each) and stored overnight at 4°C. On day 3, slides were washed with PBS and incubated in reducing agent for 10 min at room temperature. Sections were dehydrated sequentially in 70% and 100% ethanol, followed by incubation in Xenium autofluorescence quenching solution for 10 min. Slides were washed three times in 100% ethanol and dried at 37°C for 5 min. Sections were rehydrated with PBS and PBST, incubated in nuclear staining buffer for 1 min, and washed four times with PBST.

Samples were subsequently loaded into the Xenium Analyzer instrument (software version 1.8.2.1–3.2.1.3, depending on the sample), where automated cycles of reagent delivery, probe hybridization, imaging, and probe stripping were performed. The resulting Z-stack images were then processed using the Xenium onboard analysis pipeline (software version 1.7.1.0–3.2.0.8, depending on the sample) for initial image pre-processing, following the procedure described by Janesick et al.^94^. This included decoding fluorescent puncta into individual transcripts and assigning transcript quality scores (Q-scores). Only transcripts with Q-scores >20 were included in downstream analyses.

To ensure decoding accuracy and assay specificity, negative control codewords (not assigned to any probe) and negative control probes (not complementary to any biological sequence) were included. On-instrument cell segmentation was performed based on 3D DAPI nuclear morphology, and nuclei and cell boundaries were subsequently projected into a 2D segmentation mask. Cell IDs were assigned to each identified cell, and transcripts were allocated to cells based on their X–Y spatial coordinates. When the default settings were not used, cell boundaries were expanded by 5 μm from the nuclear mask using Xenium Ranger (version 1.7) to account for tightly packed cells in regenerating and non-regenerating digit tips.

### 10X Xenium data analysis

Standard Xenium output files generated using either of the two panels were used for downstream analyses. Spatial datasets were first examined in Xenium Explorer (version 3.0 or 3.2; RRID: SCR_025847). Regions of interest (ROIs) were manually delineated using the freehand selection tool with reference to *Spp1* transcript representing the amputated bone, to capture the outgrowth area. Cell identifiers corresponding to selected ROIs were exported as previously described^30–32,94^.

For the C57/Bl6 regenerating digit tips 14, 14, 16 and 30 sections were used for analysis at 5, 7, 10 and 14 DPA respectively. For the C57/Bl6 non-regenerative digit tips 21, 36, 34 and 49 sections were analyzed at 5, 7, 10 and 14 DPA respectively, all using the 480 gene panel. For the *Lmx1bCreERT2-R26-LSL-DTA* 7 DPA regenerating digit tips, 34 sections were analyzed and for *PdgfraCreERT2Smad4KO* 7 DPA 7 DPA regenerating digit tips 23 sections were analyzed, both using the 480 gene panel.

Individual Xenium fields of view (FOVs) from each tissue section were first processed separately to generate Seurat objects using the Seurat package (v5.0.1; RRID: SCR_016341). Raw Xenium output files for each FOV were loaded using the LoadXenium function. Cell IDs corresponding to regions of interest (ROIs), exported from Xenium Explorer, were used to subset each FOV. Low-quality cells were flagged based on the mean number of detected genes and total transcript counts, and cells falling outside ±2.5 standard deviations from the mean were removed. Each Seurat object was then annotated with a unique dataset identifier, and FOV image keys were adjusted to avoid conflicts during subsequent merging.

For merging, the previously generated Seurat objects were used as references to ensure high-quality cell selection: cell barcodes from the RDS files were used to subset each FOV, retaining only cells present in these reference objects. Seurat objects corresponding to the same or different timepoint or experimental condition were then merged using the merge function, with unique cell identifiers added to preserve FOV provenance. Normalization, variance stabilization, and feature selection were performed using SCTransform across all merged cells. Dimensionality reduction was performed using principal component analysis (PCA), with the optimal number of components assessed via an elbow plot. Low-dimensional embeddings were generated using Uniform Manifold Approximation and Projection (UMAP), and cell–cell relationships were determined using FindNeighbors function. Unsupervised clustering was performed at multiple resolutions ranging from 0.4 to 2.4 using FindClusters function to define transcriptionally distinct populations and cluster structure was visualized in two dimensions using UMAP embeddings. The fully processed and annotated merged Seurat object was used for downstream analyses. Gene expression and spatial distribution of clusters were assessed using ImageDimPlot, FeaturePlot, and ImageFeaturePlot to interpret both transcriptional and spatial patterns.

Cell-type identities were assigned based on the expression of established marker genes. Epidermal basal cells were identified by *Cspg4*, *Itga6* and *Irx4*. Epidermal suprabasal cells were identified by *Sp9*, *Erbb3* and *Krt10*. Nail suprabasal cells were identified by *Cybrd1*, *Fhod3* and *Plk3*. Nascent blastema cells were identified by *Ltbp2*, *Gpr27* and *Col24a1*. Early nail mesenchyme cells were identified by *Sfrp2*, *Rspo4*, *Zic2* and *Steap1.* Osteogenic blastema cells were identified by *Ltbp2*, *Acan* and *H19*. Undifferentiated blastema cells were identified by *Ltbp2*, *Arsi* and *Matn3*. Mature nail mesenchyme cells were identified by *Sfrp2*, *Rspo4*, *Dio3* and *Lmx1b*. Regenerating dermis cells were identified by *Cpxm2*, *Col26a1* and *Cxcl14*. Macrophages cells were identified by *Lyz2*, *Csf1r* and *Fcgr3*. Neutrophils cells were identified by *Csf3r*, *S100a8* and *Il1b*. T cells were identified by *Cd3e*, *Cd3g* and *Cxcr6*. Osteoclasts cells were identified by *Mmp9*, *Acp5* and *Ocstamp*. Schwann cells were identified by *Sox10*, *Plp1* and *Ngfr*. Endothelial cells were identified by *Cdh5*, *Pecam1* and *Tie1*. Pericytes/VSM cells were identified by *Rgs5*, *Myh11* and *Kcnj8*. Dermis cells were identified by *Cilp*, *Pdgfra* and *Masp1*. Follicle associated mesenchymal cells were identified by *Coch*, *Tnmd* and *Cntfr* and by their spatial location. Mesenchymal granulation cap cells were identified by *Crabp1*, *Prss35* and *Wnt5a*. Bone callus cells were identified by *Col11a1*, *Fmod* and *Tmem119*.

Spatial plots generated in Seurat were rescaled to match the dimensions of the original unprocessed scan images using Xenium Explorer.

#### Resolutions used for Xenium data annotation and visualization

Each dataset was first analyzed using a conservative clustering resolution. The resolution was increased only when necessary to reveal cellular heterogeneity or when two cell populations, identified by canonical marker expression, did not separate into distinct clusters. The following resolutions were used to annotate and visualize Xenium data within this paper. For the merged regenerative 5 DPA and 7 DPA (Figure 2A) cell type annotations were based on clustering at a resolution of 0.6. For the regenerative 10 DPA (Figure 2A) cell type annotations were based on clustering at a resolution of 1.2. For the regenerative 14 DPA (Figure 2A) cell type annotations were based on clustering at a resolution of 0.8. For merged regenerative mesenchymal cells at 5, 7, 10 and 14 DPA (Figure 3A) cell type annotations were based on clustering at a resolution of 0.5. For merged non-regenerative cells at 5, 7, 10 and 14 DPA (Figure 3G) cell type annotations were based on clustering at a resolution of 0.8. For merged non-regenerative cells at 5 DPA (Fig. S3D) cell type annotations were based on clustering at a resolution of 0.6. For merged non-regenerative cells at 7 DPA (Fig. S3E) cell type annotations were based on clustering at a resolution of 0.8. For merged non-regenerative cells at 10 DPA (Fig. S3F) cell type annotations were based on clustering at a resolution of 0.6. For merged non-regenerative cells at 14 DPA (Fig. S3G) cell type annotations were based on clustering at a resolution of 0.8. For merged regenerative and non-regenerative mesenchymal cells at 14 DPA (Figure 4A) cell type annotations were based on clustering at a resolution of 0.4. For merged regenerative and non-regenerative mesenchymal cells at 5, 7 and 10 DPA (Figure 4E) cell type annotations were based on clustering at a resolution of 0.5. For merged regenerative cells from Control and Ablated conditions at 7 DPA (Figure 4I) cell type annotations were based on clustering at a resolution of 1.4. For regenerative cells from Ablated condition at 7 DPA (Fig. S4C) cell type annotations were based on clustering at a resolution of 1.2. For regenerative cells from WT/control condition at 7 DPA (Fig. S4C) using probeset 2, cell type annotations were based on clustering at a resolution of 1.2. For merged regenerative cells from control and *Pdgfra-Smad4KO* at 7 DPA (Figure 6H) cell type annotations were based on clustering at a resolution of 1. For regenerative cells from *Pdgfra-Smad4KO* at 7 DPA (Fig. S6B) cell type annotations were based on clustering at a resolution of 0.8.

#### Xenium Explorer visualizations

Xenium Explorer images highlighting different cell types and transcripts were generated using Xenium Explorer (versions 3.0 or 3.2; RRID: SCR_025847). DAPI staining was used as the spatial reference, onto which cell boundaries, nuclear outlines, and detected transcripts were overlaid. For cell type visualization, Seurat-derived cluster assignments at the appropriate resolution were imported into Xenium Explorer, and clusters were color-coded based on their annotated cell type identities. This enabled the spatial distribution of individual mRNAs to be visualized across tissue sections.

### Quantification and statistical analysis

#### Differential gene expression analysis

Differential gene expression analysis of the scRNA-Seq datasets was conducted using SCT-normalized expression values. Data were first prepared using the PrepSCTFindMarkers function and differential expression testing was then performed using the Seurat FindAllMarkers or FindMarkers functions. Statistical comparisons were carried out using a Wilcoxon Rank Sum Test and genes with a Bonferroni-adjusted p-value of less than 0.05 were considered statistically significant. The differentially expressed genes were further filtered to only include genes with an average log2FC of greater than. 0.6 (FC > 1.5) and those expressed in at least 10% of cells in the cluster of interest.

Differential gene expression analysis of Xenium datasets was conducted using SCT-normalized expression values. Data were first prepared with the PrepSCTFindMarkers function and differential expression testing was then performed using the FindAllMarkers function in Seurat. Statistical comparisons were carried out using a Wilcoxon rank-sum test, and genes with a Bonferroni-corrected adjusted p-value below 0.05 were considered statistically significant.

#### Violin and dot plots

Violin plots were generated using the VlnPlot function to illustrate the distribution of gene expression levels across clusters. Dot plots were generated using the DotPlot function to display both the average expression level and the percentage of cells expressing each gene within each cluster.

#### Morphometric analysis of nail length and area

Digital images of collected digit tips were acquired using an Olympus SZX10 stereomicroscope (Tokyo, Japan) equipped with an Olympus DP72 digital camera and a Plan-Apo 1x objective lens, operated using Olympus cellSens Standard acquisition software. Image analysis was performed using Fiji^95^. The nail area and length were manually outlined and measured in a blinded manner. Measurements were obtained for all experimental digits from each mouse and averaged to represent a single biological replicate. A minimum of three independent mice were analyzed per time point. Statistical significance was determined using a two-tailed Student’s *t*-test, with *p* < 0.05 considered statistically significant. Error bars represent the standard error of the mean (SEM).

#### Quantification of tissue section area

To determine the area of newly-grown tissue in the regenerating and non-regenerative digit tips, midline sections from *Lmx1bCreERT2-R26-LSL-DTA*, *PdgfraCreERT2Smad4KO-tdt* and *Lmx1bCreERT2Smad4KO-tdt* mice were immunostained for PDGFRα, and nuclei were counterstained with DAPI. The PDGFRα-positive regenerated region, excluding the epithelium, was manually outlined and measured using ImageJ. For quantification of both the total regenerated region (DAPI-positive area) and the PDGFRα-positive regenerated region, three sections at comparable anatomical levels were analyzed per regenerating digit tip. Measurements from the three sections were averaged and considered as a single biological replicate. A minimum of three independent animals were analyzed per condition. Statistical significance was determined using a two-tailed Student’s t-test, with p < 0.05 considered statistically significant. Error bars represent the standard error of the mean (SEM).

#### Quantification of regenerating tissue volume

Volumetric analysis was conducted using at least three independent mice per condition. To do that, we drew a mask on the regenerated area across all z-stacks using the software Arivis Pro (4.2.2). The segmented regions from each slice were combined to generate a three-dimensional representation of the regenerated area. Subsequently, the analysis panel in Arivis was used to calculate the total volume of the regenerated tissue. Statistical significance was determined using a two-tailed Student’s t-test, with p < 0.05 considered statistically significant. Error bars represent the standard error of the mean (SEM).

#### Statistical analysis

With the exception of the scRNA-seq analyses, statistical significance was determined using a two-tailed Student’s t-test or an ANOVA when more than two groups were compared (as in Fig. 3L and 7J). In all cases, p < 0.05 was considered to be statistically significant. Error bars indicate the standard error of the mean (S.E.M.). Statistics used for the computational analyses are described in the relevant sections.

## SUPPLEMENTAL MATERIALS

### Supplemental Figures

**Figure S1. *Characterization of early 4 and 7 DPA regenerating digit tip mesenchymal cells using scRNA-seq*. (Related to Figure 1). (A)** Schematic of regenerative versus non-regenerative digit tip amputations that maintain and remove the nail base, respectively. Hatched lines indicate the amputation planes. **(B, C)** UMAP of transcriptomes of the regenerating digit tip at 4 (B) and 7 (C) DPA, as defined using scRNA-seq. The 4 and 7 DPA datasets include cells from 2 independent runs each. Transcriptionally-distinct clusters are color-coded and cell types are annotated as determined using cell type-specific marker genes. **(D)** Single-cell heatmap showing expression of 5 nail mesenchyme signature mRNAs (*Gpm6a, Cyp2f2, Nrg2, Rspo4, Snhg11*), *Panx3*, which is associated with osteogenic differentiation, and *Sfrp4*, which is specifically enriched in the nascent blastema. Every row represents expression levels in an individual cell, and levels are color-coded as per the adjacent key. NM = nail mesenchyme. **(E-G)** UMAPs of the merged transcriptomes of *Pdgfra*-positive mesenchymal cells from the new 4 and 7 DPA datasets (from B and C) together with those from the previously-published^8,18^ (GEO GSE135985; GEO GSE217600) 10, 14, 28 and 56 DPA and uninjured digit tip datasets as shown in Fig. 1E, F. UMAPs are overlaid for expression of mRNAs associated with osteogenic differentiation (E) or that were identified as enriched specifically in the early/nascent blastema (F) or in both the early 4-7 DPA nail mesenchyme and in the early/nascent blastema (G) (see Table S1). Expression levels are coded as per the adjacent keys.

**Figure S2. *Characterization of the 5 and 7 DPA Xenium-based single cell spatial transcriptomic datasets*. (Related to Figure 2). (A)** UMAP of the regenerating digit tip Xenium-based single cell transcriptomes from the merged 5 and 7 DPA datasets, annotated as in Fig. 2A. The left panel shows the annotated dataset while the middle and right panels show the datasets of origin for the different transcriptomes. **(B)** Annotated UMAPs of the regenerating digit tip Xenium-based single cell transcriptomes from 10 (left panel) and 14 (right panel) DPA, as in Fig. 2A. **(C-E)** Gene expression overlays for selected marker genes on UMAPs of the merged 5 and 7 DPA dataset (C) or of the 10 and 14 DPA datasets (D and E, respectively). In each case expression levels are coded as per the adjacent keys.

**Figure S3. (A-C) *Characterization of 10 and 14 DPA regenerative Pdgfra-positive mesenchymal cells***. **(Related to Figure 3). (A)** Spatial plots of the 10 and 14 DPA regenerating digit tip sections shown in Fig. 2D, overlaid for expression of two blastema mRNAs, *Ltbp2* and *Arsi*, color-coded as per the adjacent keys. **(B)** Spatial plots of the same representative 10 and 14 DPA sections as in (A), overlaid for expression of *Alpl* mRNA, with levels coded as per the adjacent keys. **(C)** Spatial plots of the same representative 14 DPA regenerating digit tip section as in (A), overlaid for expression of two mRNAs enriched in the regenerating dermis, *Cpxm2* and *Mmp9*, with levels coded as per the adjacent keys. **(D-J) *Single cell spatial transcriptomics to define cells in the non-regenerative digit tip at 5 to 14 DPA***. **(D-G)** UMAPs of Xenium-based single cell spatial transcriptomes from the non-regenerative digit tip at 5 DPA (D), 7 DPA (E), 10 DPA (F) and 14 DPA (G), annotated for cell types as determined using marker genes. **(H)** Transcriptomes from the individual non-regenerative datasets shown in (D-G) were merged, and annotated, as shown in Fig. 3G. Shown here are the transcriptomes colored to identify their dataset of origin within this merger. **(I)** Spatial plots of the same representative 5 and 7 DPA non-regenerative sections as in Fig. 3H, showing the epidermal cells (blue) and the follicle-associated mesenchymal cells (purple) that are present in areas proximal to the amputation. **(J)** High-resolution Xenium Explorer images of the same representative 10 DPA non-regenerative section as in Fig. 3H, showing the connective tissue cap cells (brown, both panels) and the bone callus cells (green, both panels), overlaid for expression of *Comp* mRNA (green dots, right panel) and *Prss35* mRNA (blue dots, right panel). The right panel is a higher magnification image of the boxed region in the left panel. Scale bars = 100 µm (left) and 20 µm (right).

**Fig. S4. (A, B) *Characterization of regenerative versus non-regenerative mesenchymal cells.* (Related to Figure 4). (A)** UMAP of merged 14 DPA regenerative and non-regenerative *Pdgfra*-positive mesenchymal cell transcriptomes, as shown in Fig. 4A. The UMAP on the left shows color-coded transcriptionally-distinct mesenchymal cell types, annotated. The UMAPS in the centre and right panels show gene expression overlays for one regenerative mRNA, *Fgf10* and one non-regenerative mRNA, *Angptl1*, color-coded as per the adjacent keys. **(B)** UMAP of merged 5, 7 and 10 DPA regenerative and non-regenerative *Pdgfra*-positive mesenchymal cell transcriptomes, as shown in Fig. 4E. The UMAP on the left shows color-coded transcriptionally-distinct mesenchymal cell types, annotated. The other UMAPs are overlaid for expression of three regenerative mRNAs, *Bmp5, Fgf10* and *Gldn*, and two non-regenerative mRNAs, *Cilp* and *Thbs4*, color-coded as per the adjacent keys. ***(C-I) Single cell spatial transcriptomic analysis of 7 DPA Lmx1b-DTA and Lmx1b-WT regenerating digit tips.*** *Lmx1b-DTA* (Ablated) or *Lmx1b-WT* mice were treated with tamoxifen, their digit tips were amputated 8 days later, and single cell spatial transcriptomics was performed after a further 7 days (7 DPA). For the *Lmx1b-DTA* sections, Xenium was performed using the same probeset as for all other datasets (Table S2, probeset 1). For the *Lmx1b-WT* sections, Xenium was performed using a different probeset that also distinguishes the different digit tip cell types (Table S2, probeset 2). **(C)** UMAP of 7 DPA *Lmx1b-DTA* regenerating digit tip transcriptomes acquired using Xenium-based single cell spatial transcriptomics with probeset 1, annotated for cell types. **(D)** UMAP of 7 DPA *Lmx1b-WT* regenerating digit tip transcriptomes acquired using Xenium-based single cell spatial transcriptomics with probeset 2 (see Table S2). Cell types are color-coded and annotated. **(E)** Spatial plots of the dataset in (D), showing all detected cell types on a representative 7 DPA *Lmx1b-WT* midline section. Cell types are color-coded as per the adjacent legend. **(F-I)** Spatial plots of the same representative 7 DPA section as in (E), showing (F) the nail mesenchyme and nascent blastema cells, (G) vascular-associated endothelial cells and mural cells, (H) macrophages/neutrophils, and (I) the different populations of epithelial cells.

**Figure S5. *Expression of selected ligands and Smad4 in mesenchymal cells of the regenerating digit tip*. (Related to Figure 5). (A)** 5-7 DPA early nail mesenchyme scRNA-seq-derived transcriptomes (as in Fig. 1E) were analyzed for their expression of ligand and ligand receptor mRNAs. Ligand mRNAs were included if they were expressed in ≥ 5% of the early nail mesenchyme cells while receptor mRNAs were included if they were expressed in ≥ 20% of the early nail mesenchyme cells. The data were used to generate a predictive model of potential autocrine interactions within the early regenerating nail mesenchyme. Each box includes a ligand known to bind to a corresponding receptor expressed by the nascent blastema cells. Also see Tables S3-5. **(B)** High resolution Xenium Explorer images of a representative 14 DPA regenerating digit tip section, showing expression of *Bmp4* mRNA (green dots), *Bmp5* mRNA (yellow dots), *Gdf6* mRNA (turquoise dots), *Fgf10* mRNA (pink dots) and *Igf2* mRNA (red dots). Scale bars = 100 µm. **(C)** *Smad4* gene expression overlay on the 7 DPA scRNA-seq UMAP as in Fig. S1B, with *Pdgfra*-positive mesenchymal cell clusters outlined. **(D)** *Smad4* gene expression overlay on the 5 plus 7 DPA Xenium UMAP from Fig. 2A with *Pdgfra*-positive mesenchymal cell clusters outlined. **(E)** *Smad4* gene expression overlay on the 14 DPA Xenium UMAP from Fig. 2A with *Pdgfra*-positive mesenchymal cell clusters outlined.

**Figure S6. *Loss of Smad4 in mesenchymal cells inhibits their acquisition of a blastema state***. **(Related to Figures 6 and 7). (A)** *Pdgfra*-*Smad4KO* or *Pdgfra-WT* mice were treated with tamoxifen, regenerative amputations were performed after 7 days, and the length of the regenerated nails was quantified after a further 14 days. n = 8-12 mice, ****p<0.0001. **(B)** UMAP of Xenium-based single cell transcriptomes from 7 DPA regenerative digit tips of *Pdgfra-Smad4KO* mice, annotated for cell types. **(C)** UMAP of merged *Pdgfra*-positive mesenchymal cell transcriptomes from the *Lmx1b-DTA*, *Pdgfra-Smad4KO*, and control 7 DPA regenerating digit tips. The nascent blastema, nail mesenchyme and bone were all identified by marker gene expression.

### Supplemental Videos

**Video S1: *Three-dimensional rendering of SHIELD-cleared regenerative 14 DPA digit tip from a PdgfraCreERT2-TdT mouse.* (Related to Figure 1).** A three-dimensional, rendered whole-mount light-sheet fluorescence video of a regenerating 14 DPA distal digit tip from a *PdgfraCreERT2-TdT* reporter mouse. Tamoxifen was administered for five consecutive days, and the digit tip amputated after a further 7 days. Tissue was collected at 14 DPA, nail plates were removed, and samples were cleared using the SHIELD protocol (see Experimental Methods). The cleared, transparent digits exhibited TdT signal (red) within mesenchymal cells, while background morphology was visualized using a gray signal from the 488 nm laser channel. Post-processing of the images was performed using Arivis software (Zeiss). The masked region, shown in faint gray, indicates the regenerated area. The video begins with a lateral view of the regenerated digit tip. At 0.07 seconds, rotation of the digit reveals TdT-positive cells in the proximal dermal layer, followed by visualization of the TdT-positive cells from the opposite lateral view.

**Video S2. *Three-dimensional rendering of SHIELD-cleared non-regenerative 14 DPA digit tip from a PdgfraCreERT2-TdT mouse.* (Related to Figure 1).** A three-dimensional, rendered whole-mount light-sheet fluorescence video of a 14 DPA non-regenerative digit tip from a *PdgfraCreERT2-TdT* reporter mouse. Tamoxifen was administered for five consecutive days, and non-regenerative digit tip amputations performed after a further 7 days. Tissue was collected at 14 DPA and samples were cleared using the SHIELD protocol (see Experimental Methods). The cleared, transparent digits exhibited TdT signal (red) within mesenchymal cells, while background morphology was visualized using a gray signal from the 488 nm laser channel. Post-processing of the images was performed using Arivis software (Zeiss). The masked region, shown in faint gray, indicates the new tissue distal to the amputation. The video begins with a lateral view of the non-regenerated digit tip. At 0.10 seconds, rotation of the digit reveals TdT-positive cells in the proximal dermal layer, followed by visualization of TdT-positive cells from the opposite lateral view.

**Video S3. *Three-dimensional rendering of SHIELD-cleared regenerative 14 DPA digit tip from a Cdh5Cre-TdT mouse*. (Related to Figure 1).** A three-dimensional rendered whole-mount light-sheet fluorescence image of a regenerating 14 DPA digit tip from a *Cdh5Cre-TdT* reporter mouse. Digit tips were collected 14 days after regenerative amputations, nail plates were removed, and samples were cleared using the SHIELD protocol (see Experimental Methods). The cleared, transparent digits exhibited TdT signal (red) within endothelial cells, while background morphology was visualized using a gray signal from the 488 nm laser channel. Post-processing of the images was performed using Arivis software. The masked region, shown in faint gray, indicates the regenerated area. The video begins with a lateral view of the regenerated digit tip. At 0.11 seconds, rotation of the digit reveals TdT-positive blood vessels in the proximal digit tip, followed by visualization of TdT-positive blood vessels in the regenerated tissue from the opposite lateral view.

**Video S4. *Three-dimensional rendering of SHIELD-cleared non-regenerative 14 DPA digit tip from a Cdh5Cre-TdT mouse*. (Related to Figure 1).** A three-dimensional rendered whole-mount light-sheet fluorescence video of a non-regenerative 14 DPA digit tip from a *Cdh5Cre-TdT* reporter mouse. Digit tips were collected 14 days following non-regenerative amputations and were then cleared using the SHIELD protocol (see Experimental Methods). The cleared, transparent digits exhibited TdT signal (red) within endothelial cells, while background morphology was visualized using a gray signal from the 488 nm laser channel. Post-processing of the images was performed using Arivis software (Zeiss). The masked region (faint gray) delineates the new tissue distal to the amputation plane. The video begins with a lateral view of the blood vessels in the non-regenerative digit tip. At 0.11 seconds, rotation of the digit reveals TdT-positive cells in the proximal digit, followed by visualization of TdT-positive blood vessels in the new tissue from the opposite lateral view.

**Video S5. *Three-dimensional rendering of SHIELD-cleared 14 DPA regenerating digit tip from a PdgfraCreERT2-WT-TdT mouse.* (Related to Figure 6).** A three-dimensional rendered whole-mount light-sheet fluorescence image of a regenerating 14 DPA distal digit tip from a *PdgfraCreERT2-TdT* mouse that was wildtype for the floxed *Smad4* allele. Tamoxifen was administered for five consecutive days, and non-regenerative digit tip amputations performed after a further 7 days. Digit tips were collected at 14 DPA, nail plates were removed, and samples were cleared using the SHIELD protocol (see Experimental Methods). The cleared, transparent digits exhibit TdT signal (red) within mesenchymal cells, while overall morphology is visualized using a gray signal from the 488 nm laser channel. Post-processing of the images was performed using Arivis software. The masked region, shown in faint gray, indicates the regenerated area. The video begins with a lateral view of the regenerated digit tip. At 0.10 seconds, rotation of the digit reveals TdT-positive cells in the proximal dermal layer, followed by visualization of TdT-positive cells in the regenerated tissue from the opposite lateral view.

**Video S6. *Three-dimensional rendering of SHIELD-cleared regenerating 14 DPA digit tip from a PdgfraCreERT2-Smad4KO-TdT mouse.* (Related to Figure 6).** A three-dimensional, rendered whole-mount light-sheet fluorescence video of a regenerating 14 DPA digit tip from a *Pdgfra-Smad4KO-TdT* reporter mouse. Tamoxifen was administered for five consecutive days, and non-regenerative digit tip amputations performed after a further 7 days. Digit tips were collected at 14 DPA, nail plates were removed, and samples were cleared using the SHIELD protocol (see Experimental Methods). The cleared, transparent digits exhibit TdT signal (red) within mesenchymal cells, while overall morphology is visualized using a gray signal from the 488 nm laser channel. Post-processing of the images was performed using Arivis software. The masked region, shown in faint gray, indicates the regenerated area. The video begins with a lateral view of the regenerated digit tip. At 0.10 seconds, rotation of the digit reveals TdT-positive cells in the proximal digit tip, followed by visualization of TdT-positive cells in the regenerated tissue from the opposite lateral view.

### Supplemental Tables

**Table S1. *Differentially expressed genes in the early nail mesenchyme or nascent blastema*. (Related to Figure 1).** Shown are mRNAs differentially expressed in a comparison of the early nail mesenchyme (cluster 3) or nascent blastema (cluster 7) compared to all other cells in the merged dataset shown in Figure 1E. Genes were considered significantly differentially enriched if the gene had a Bonferroni corrected p-value (p_val_adj) < 0.05, the average log2FC (avg_log2FC) > 0.6, and the gene was expressed in at least 10% of cells in the queried cluster (pct.1 > 0.1). Also shown are the uncorrected p-value, and percent expression in all other cell types.

**Table S2. *Probes used for Xenium single-cell spatial transcriptomics* (Related to Figures 2-7 and Figures S2-6).** Shown are the genes included in the custom Xenium probesets 1 and 2, comprised of probes for 480 and 473 genes, respectively. Included are Ensembl IDs for all genes used, as well as the number of probes used for each gene. The number of probes assigned to each gene was optimized individually to achieve strong detection and adequate isoform coverage, while minimizing optical crowding. Probe selection was guided by the 10x Xenium panel designer. The genes highlighted in yellow represent bioactive ligands that are targeted by the Xenium panel.

**Table S3. *Ligands expressed in the early nail mesenchyme or nascent blastema.* (Related to Figure 5 and Figure S5).** Shown are the ligands expressed by the early nail mesenchyme (cluster 3) or nascent blastema (cluster 7) from the merged scRNA-seq dataset shown in Figure 1E. Ligands were identified using a previously published database^36^ (see Experimental Methods) and were considered to be expressed if the mRNA was detected in at least 5% of the cells within the cluster. The gene abundance is shown for each ligand.

**Table S4. *Receptors expressed in the early nail mesenchyme or nascent blastema.* (Related to Figure 5 and Figure S5).** Shown are the receptors expressed by the early nail mesenchyme (cluster 3) or nascent blastema (cluster 7) from the merged scRNA-seq dataset shown in Figure 1E. receptors were identified using a previously published database (see Experimental Methods) and were considered to be expressed if the mRNA was detected in at least 20% of the cells within the cluster. The gene abundance is shown for each ligand.

**Table S5. *Ligand-receptor modelling between the early nail mesenchyme and nascent blastema*. (Related to Figure 5 and Figure S5).** Predicted ligand and receptor communication models were generated from the merged scRNA-seq dataset in Figure 1E using the ligands and ligand receptors defined in Tables S3 and S4, as described in the Experimental Methods. Shown are ligand-receptor communication models for autocrine and paracrine interactions between the early nail mesenchyme and the nascent blastema. Gene abundance for the listed ligands and receptors are indicated.

**Table S6. *Xenium probes differentially detected in a comparison between nascent blastema and perturbation enriched mesenchymal cells*. (Related to Figure 7)** Shown are Xenium probes differentially detected in a comparison between the nascent blastema and perturbation enriched mesenchymal cell clusters shown in the UMAP of the merged 7 DPA *Lmx1b-DTA*, *Pdgfra-Smad4KO* and control Xenium single cell datasets as shown in Figure S6C. Probes were considered to be significantly enriched if the gene had a Bonferroni-adjusted p-value of less than 0.05 (p adj < 0.05) and the gene was expressed in at least 5% of nascent blastema cells. Also shown are the average fold-change (avg. fold-change) and the log-normalized fold-change (avg. log2FC).

