## Supplementary figures and images for "The nail mesenchyme creates a regeneration-specific ligand environment that orchestrates mammalian regeneration versus fibrotic wound healing"

### Supplemental_Figure_1

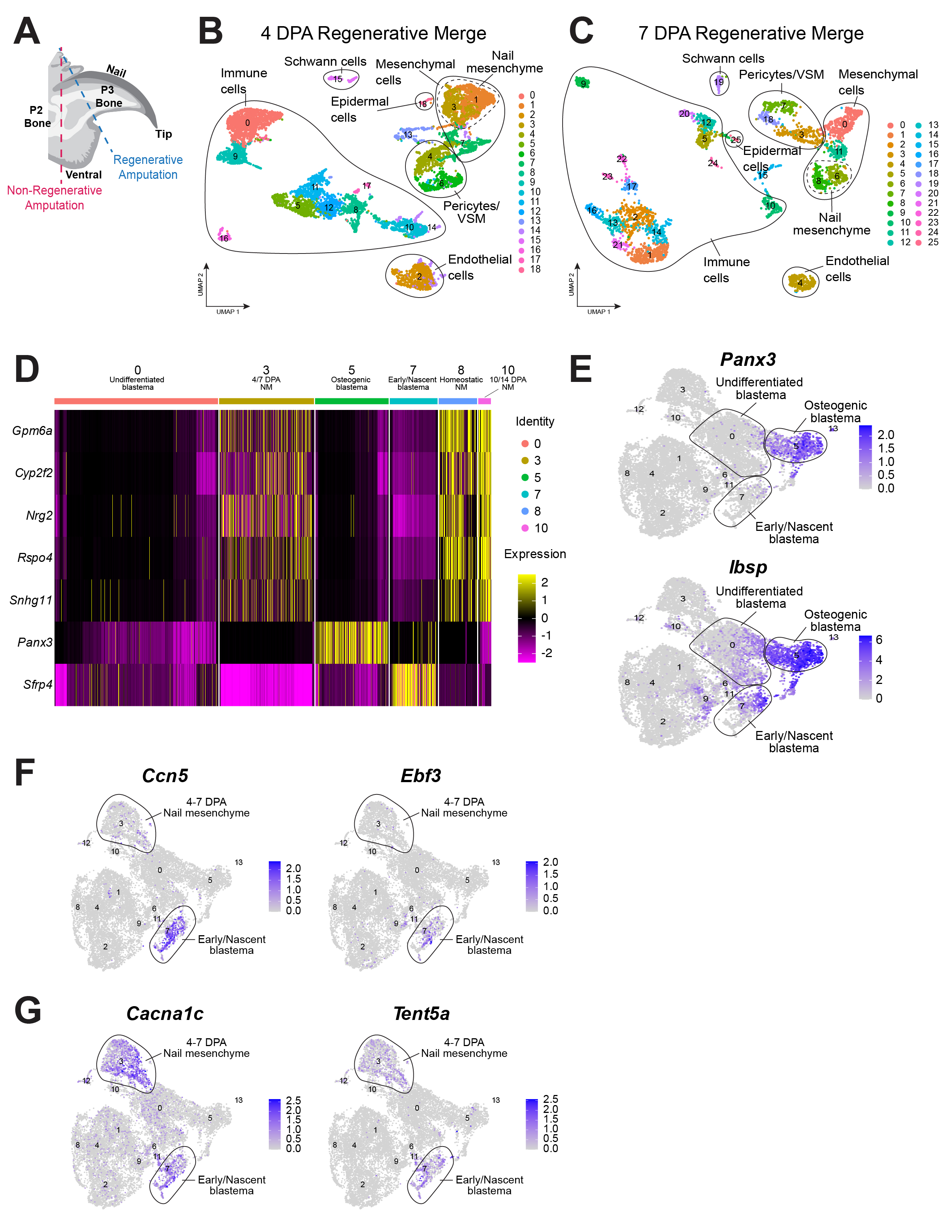

### Supplemental_Figure_2

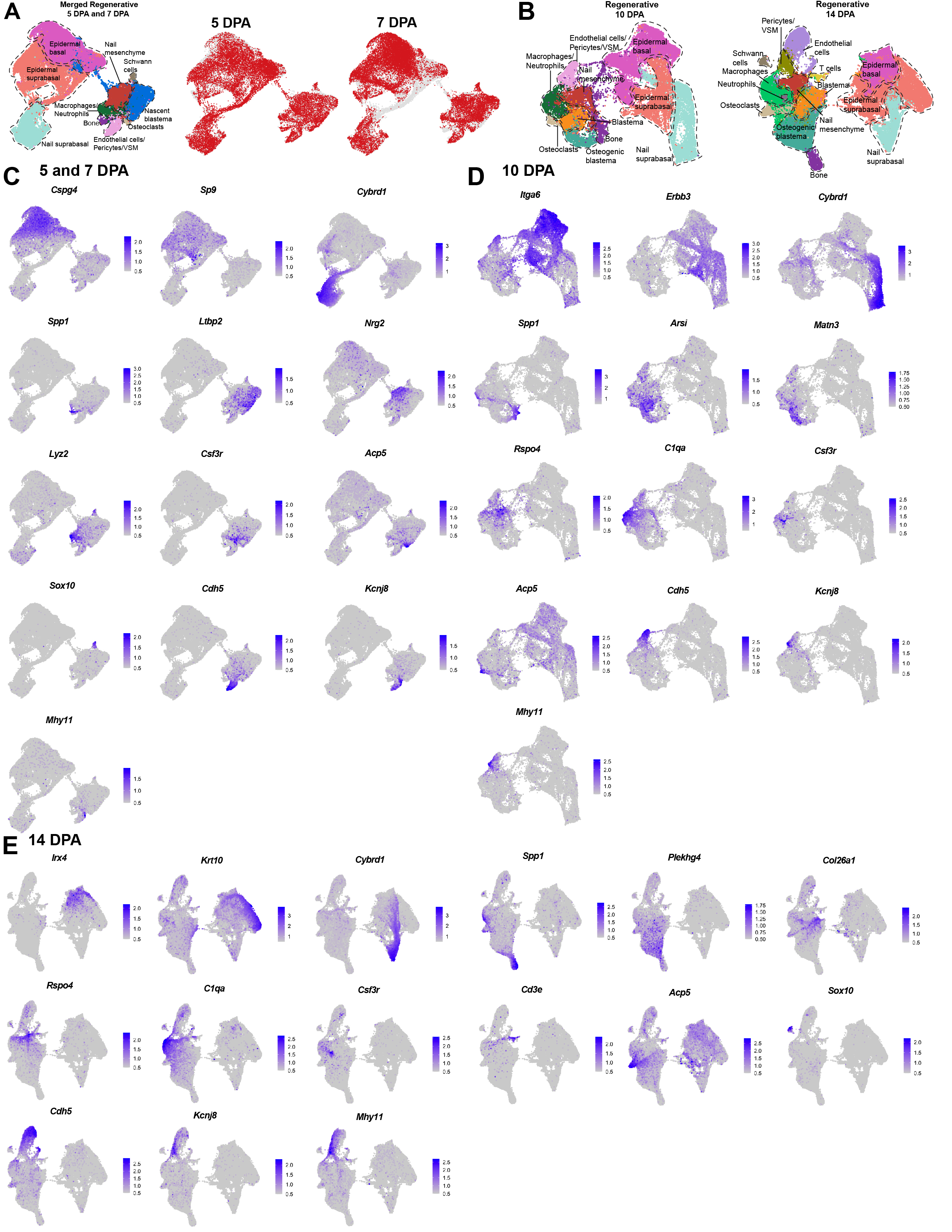

### Supplemental_Figure_3

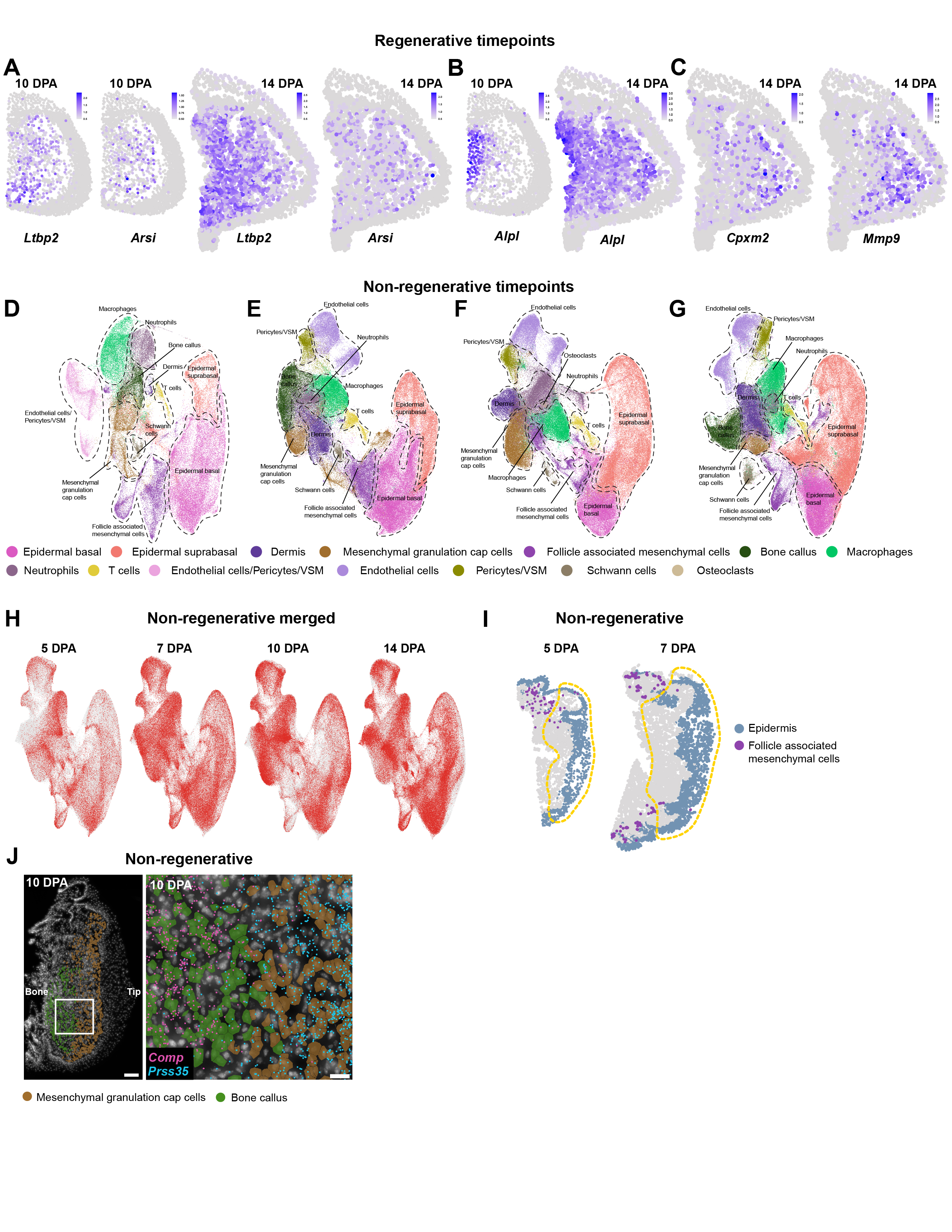

### Supplemental_Figure_4

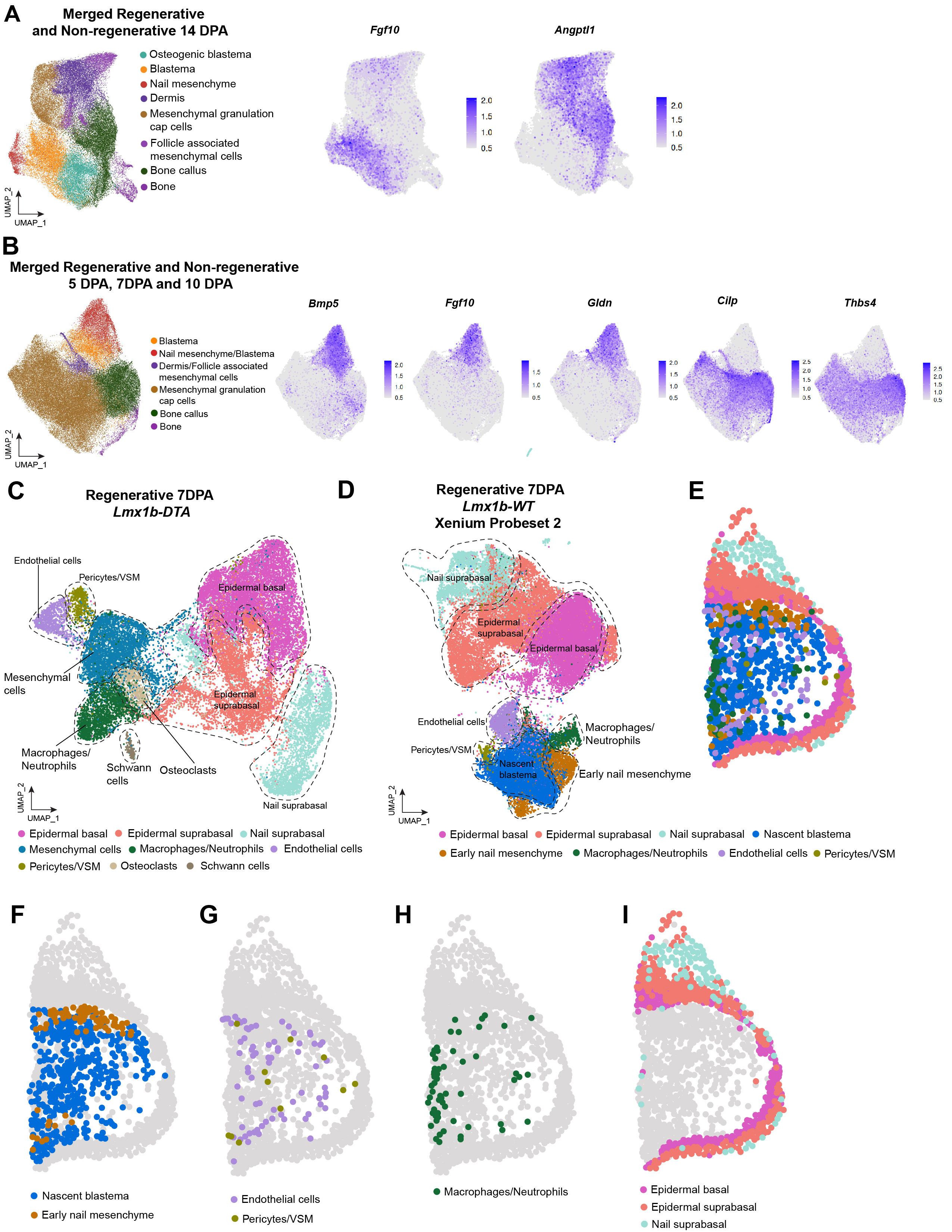

### Supplemental_Figure_5

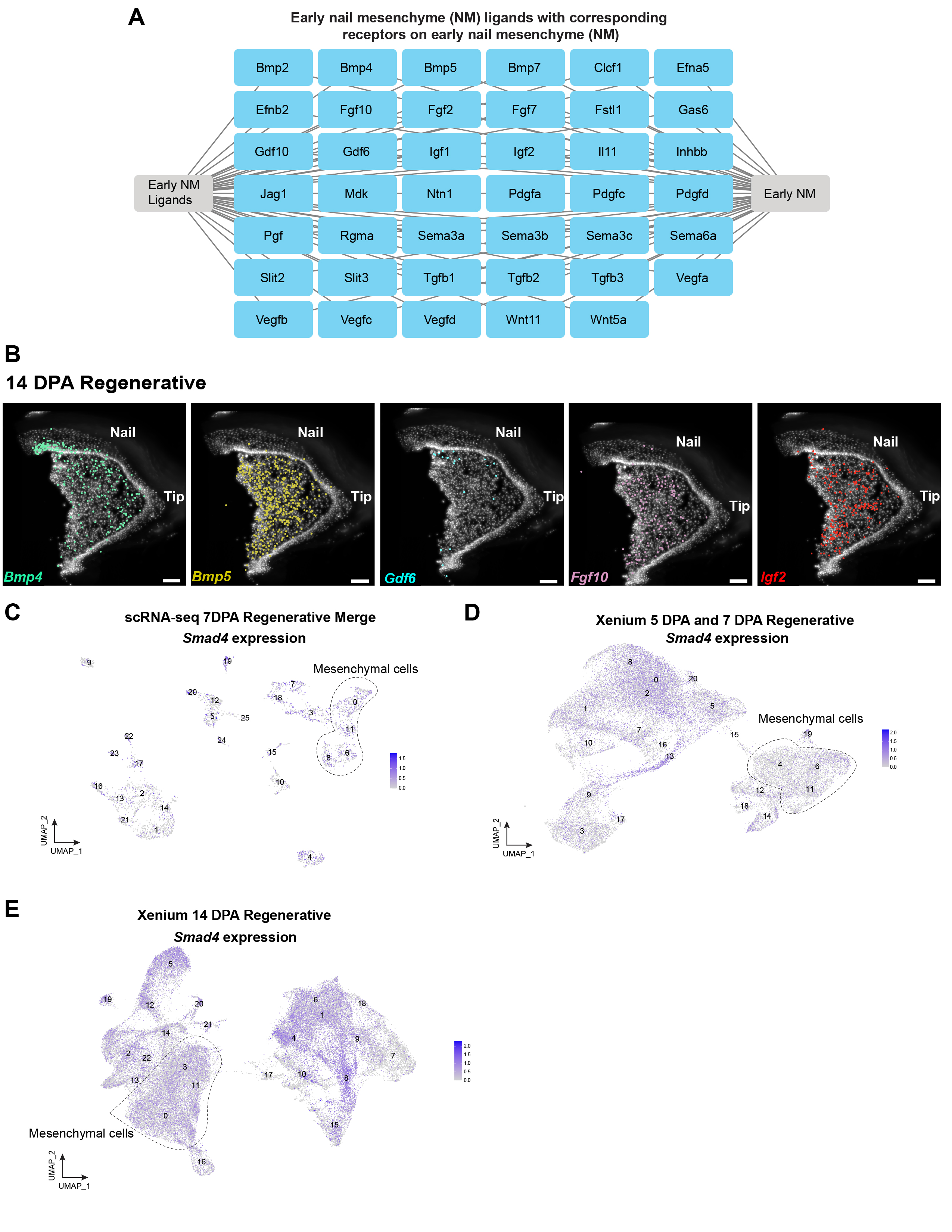

### Supplemental_Figure_6

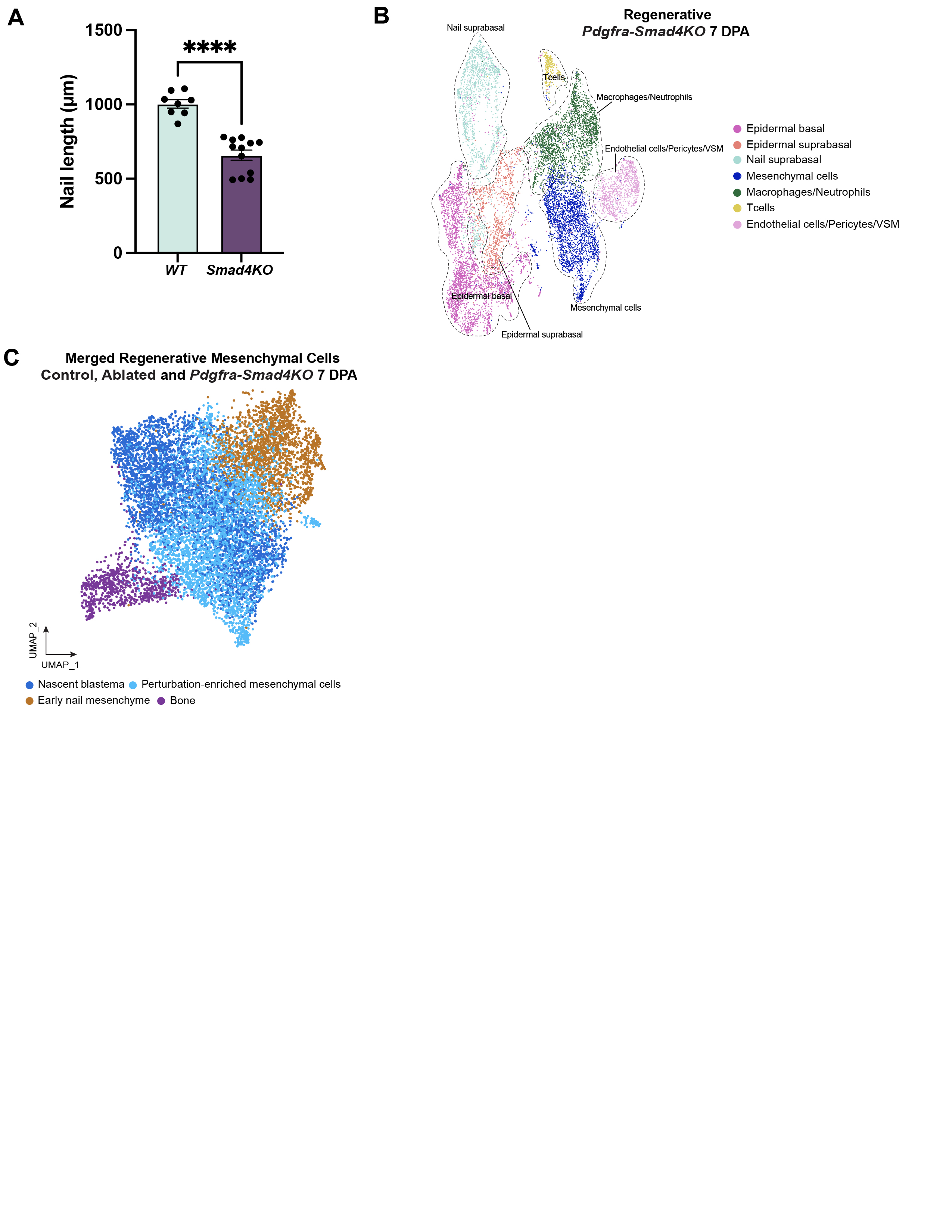
