## Supplementary material for "The nail mesenchyme creates a regeneration-specific ligand environment that orchestrates mammalian regeneration versus fibrotic wound healing": Table_1

**Table 1. Differentially Expressed Genes in Early Nail Mesenchyme or Nascent Blastema.** Shown are mRNAs differentially expressed in a comparison of the early nail mesenchyme (cluster 3) or nascent blastema (cluster 7) compared to all other cells in the merged dataset shown in Figure 1E. Genes were considered significantly differentially enriched if the gene had a Bonferroni corrected p-value (p_val_adj) < 0.05, the average log2FC (avg_log2FC) > 0.6, and the gene was expressed in at least 10% of cells in the queried cluster (pct.1 > 0.1). Also shown are the uncorrected p-value, and percent expression in all other cell types.

**Threshold:**
p_val_adj < 0.05
avg_log2FC > 0.6
pct.1 > 0.1

| Cluster 3 - Early Nail Mesenchyme | | | | | |
|---|---|---|---|---|---|
| Gene | p_val | avg_log2FC | pct.1 | pct.2 | p_val_adj |
| Mir99ahg | 0 | 5.83683939 | 0.913 | 0.049 | 0 |
| Prr16 | 0 | 5.22504494 | 0.883 | 0.074 | 0 |
| Frmpd4 | 0 | 6.04428361 | 0.829 | 0.026 | 0 |
| Cmss1 | 0 | 3.16929888 | 0.901 | 0.109 | 0 |
| Gphn | 0 | 3.58993374 | 0.821 | 0.09 | 0 |
| Masp1 | 0 | 4.15234572 | 0.853 | 0.124 | 0 |
| Camk1d | 0 | 3.06637408 | 0.815 | 0.135 | 0 |
| Ednra | 0 | 3.48135342 | 0.803 | 0.142 | 0 |
| Sfrp2 | 0 | 3.50735074 | 0.86 | 0.201 | 0 |
| Aopep | 0 | 4.17411237 | 0.701 | 0.046 | 0 |
| Steap2 | 0 | 3.23184919 | 0.778 | 0.126 | 0 |
| Setbp1 | 0 | 3.64222782 | 0.829 | 0.177 | 0 |
| Trpm3 | 0 | 5.5941808 | 0.672 | 0.025 | 0 |
| Mllt3 | 0 | 4.36563528 | 0.745 | 0.099 | 0 |
| Robo2 | 0 | 4.42939815 | 0.758 | 0.123 | 0 |
| Rbms3 | 0 | 3.39355446 | 0.859 | 0.228 | 0 |
| Pbx1 | 0 | 3.17021448 | 0.857 | 0.248 | 0 |
| Zfp521 | 0 | 3.71925297 | 0.731 | 0.132 | 0 |
| Cadm2 | 0 | 5.87939511 | 0.618 | 0.022 | 0 |
| Psd3 | 0 | 3.636054 | 0.714 | 0.129 | 0 |
| Ntrk2 | 0 | 3.05327763 | 0.722 | 0.141 | 0 |
| Tshz3 | 0 | 3.68490113 | 0.69 | 0.114 | 0 |
| Magi1 | 0 | 5.33735137 | 0.602 | 0.029 | 0 |
| Nfia | 0 | 2.98656186 | 0.76 | 0.188 | 0 |
| Gpm6b | 0 | 2.72608671 | 0.959 | 0.389 | 0 |
| Lmx1b | 0 | 5.12448747 | 0.595 | 0.027 | 0 |
| Serpina3n | 0 | 3.32256423 | 0.688 | 0.131 | 0 |
| Steap1 | 0 | 3.36819655 | 0.636 | 0.081 | 0 |
| Magi2 | 0 | 3.66332737 | 0.647 | 0.092 | 0 |
| Rorb | 0 | 4.8764947 | 0.635 | 0.08 | 0 |
| Ssbp2 | 0 | 3.19944715 | 0.729 | 0.177 | 0 |
| Trps1 | 0 | 3.74109471 | 0.975 | 0.424 | 0 |
| Mir100hg | 0 | 4.31248341 | 0.583 | 0.034 | 0 |
| Mecom | 0 | 3.53506333 | 0.734 | 0.186 | 0 |
| Kctd8 | 0 | 5.31580642 | 0.578 | 0.031 | 0 |
| Naaladl2 | 0 | 3.58383537 | 0.636 | 0.09 | 0 |
| Glis3 | 0 | 3.55398188 | 0.669 | 0.125 | 0 |
| Tmtc2 | 0 | 4.80771189 | 0.598 | 0.057 | 0 |
| Slc16a2 | 0 | 2.8320856 | 0.693 | 0.153 | 0 |
| Dennd1a | 0 | 3.56643707 | 0.622 | 0.083 | 0 |
| Gm26771 | 0 | 4.05940126 | 0.598 | 0.066 | 0 |
| Cfap69 | 0 | 5.08971289 | 0.559 | 0.031 | 0 |
| Gas5 | 0 | 3.41975783 | 0.581 | 0.056 | 0 |
| Prkg1 | 0 | 3.93377133 | 0.577 | 0.065 | 0 |
| Lama4 | 0 | 2.92981721 | 0.688 | 0.177 | 0 |
| Pde10a | 0 | 4.2479149 | 0.598 | 0.088 | 0 |
| Ifitm1 | 0 | 2.32352136 | 0.723 | 0.216 | 0 |
| Ghr | 0 | 3.1820793 | 0.669 | 0.163 | 0 |
| Plscr4 | 0 | 3.47996574 | 0.6 | 0.096 | 0 |
| Cped1 | 0 | 2.3484743 | 0.82 | 0.317 | 0 |
| Cfb | 0 | 5.00824431 | 0.534 | 0.034 | 0 |
| Sned1 | 0 | 3.03910085 | 0.637 | 0.14 | 0 |
| Lncpint | 0 | 3.33028064 | 0.594 | 0.1 | 0 |
| Pde1a | 0 | 3.38117284 | 0.581 | 0.088 | 0 |
| Tead1 | 0 | 2.373754 | 0.726 | 0.233 | 0 |
| Lrmda | 0 | 4.17407793 | 0.526 | 0.038 | 0 |
| Pdzrn3 | 0 | 3.27860639 | 0.612 | 0.124 | 0 |
| Diaph2 | 0 | 3.08165293 | 0.585 | 0.102 | 0 |
| Cpne8 | 0 | 2.36370705 | 0.705 | 0.222 | 0 |
| Phf21a | 0 | 2.86159478 | 0.604 | 0.126 | 0 |
| Kctd1 | 0 | 1.96711789 | 0.75 | 0.272 | 0 |
| Arl15 | 0 | 3.25321557 | 0.543 | 0.068 | 0 |
| Hdac4 | 0 | 2.47493974 | 0.653 | 0.18 | 0 |
| Epha7 | 0 | 4.86566671 | 0.514 | 0.041 | 0 |
| Sema3a | 0 | 4.95376194 | 0.519 | 0.046 | 0 |
| Sgip1 | 0 | 4.05584144 | 0.523 | 0.051 | 0 |
| Epha5 | 0 | 4.81890897 | 0.54 | 0.068 | 0 |
| Dio2 | 0 | 3.33070507 | 0.554 | 0.082 | 0 |
| Fbxl7 | 0 | 3.6026618 | 0.541 | 0.07 | 0 |
| Cdk8 | 0 | 2.89094476 | 0.565 | 0.097 | 0 |
| Ptprg | 0 | 3.07293665 | 0.581 | 0.113 | 0 |
| Adgrl2 | 0 | 3.24367256 | 0.561 | 0.094 | 0 |
| Bcas3 | 0 | 3.80228692 | 0.537 | 0.071 | 0 |
| Zeb2 | 0 | 2.10139535 | 0.907 | 0.444 | 0 |
| AY036118 | 0 | 2.12994028 | 0.865 | 0.402 | 0 |
| Arhgef12 | 0 | 2.10281332 | 0.677 | 0.216 | 0 |
| Atp6v0c | 0 | 3.21385649 | 0.509 | 0.053 | 0 |
| Plpp3 | 0 | 1.56523548 | 0.828 | 0.38 | 0 |
| Selenow | 0 | 1.6496579 | 0.671 | 0.224 | 0 |
| Selenom | 0 | 1.7535579 | 0.656 | 0.21 | 0 |
| Elob | 0 | 1.61076466 | 0.686 | 0.242 | 0 |
| Trabd2b | 0 | 3.38028746 | 0.521 | 0.079 | 0 |
| Spock1 | 0 | 4.36307437 | 0.493 | 0.052 | 0 |
| Mkx | 0 | 3.14731048 | 0.541 | 0.101 | 0 |
| Pdgfra | 0 | 1.58631695 | 0.824 | 0.384 | 0 |
| Rspo4 | 0 | 3.57154081 | 0.482 | 0.043 | 0 |
| Ltbp1 | 0 | 2.23722469 | 0.765 | 0.328 | 0 |
| Hsd17b4 | 0 | 2.81611476 | 0.553 | 0.117 | 0 |
| Dach1 | 0 | 5.01035789 | 0.457 | 0.022 | 0 |
| Aff2 | 0 | 4.88359941 | 0.459 | 0.025 | 0 |
| Chsy3 | 0 | 4.30336021 | 0.468 | 0.035 | 0 |
| Nbea | 0 | 3.08340731 | 0.518 | 0.085 | 0 |
| Steap4 | 0 | 2.34889686 | 0.578 | 0.147 | 0 |
| Cacna1c | 0 | 2.98023119 | 0.522 | 0.092 | 0 |
| Stag1 | 0 | 2.27923065 | 0.61 | 0.18 | 0 |
| Vcan | 0 | 2.16506007 | 0.828 | 0.398 | 0 |
| Zbtb20 | 0 | 2.92230615 | 0.984 | 0.554 | 0 |
| Lars2 | 0 | 1.54638456 | 0.801 | 0.372 | 0 |
| Rev3l | 0 | 2.74800103 | 0.545 | 0.118 | 0 |
| Sybu | 0 | 3.96809415 | 0.473 | 0.048 | 0 |
| Sulf1 | 0 | 2.5129149 | 0.642 | 0.219 | 0 |
| Rora | 0 | 2.14322687 | 0.814 | 0.392 | 0 |
| Pappa | 0 | 2.12146412 | 0.635 | 0.215 | 0 |
| Robo1 | 0 | 3.04104185 | 0.532 | 0.112 | 0 |
| Cntn1 | 0 | 4.21906735 | 0.46 | 0.043 | 0 |
| Sik3 | 0 | 2.39804674 | 0.564 | 0.15 | 0 |
| Cblb | 0 | 3.04267078 | 0.499 | 0.087 | 0 |
| Runx1t1 | 0 | 2.90113422 | 0.522 | 0.112 | 0 |
| Col23a1 | 0 | 2.66671521 | 0.605 | 0.196 | 0 |
| Mmp16 | 0 | 4.2610169 | 0.45 | 0.042 | 0 |
| Fbxl17 | 0 | 3.08953147 | 0.484 | 0.077 | 0 |
| Kansl1 | 0 | 2.27008604 | 0.569 | 0.163 | 0 |
| Pappa2 | 0 | 3.84449829 | 0.475 | 0.071 | 0 |
| Gulp1 | 0 | 2.3877938 | 0.554 | 0.153 | 0 |
| Pde4d | 0 | 2.63612119 | 0.532 | 0.131 | 0 |
| Bcl2 | 0 | 3.36062808 | 0.473 | 0.073 | 0 |
| Rsrc1 | 0 | 2.47093888 | 0.537 | 0.138 | 0 |
| Gpc6 | 0 | 3.52200552 | 0.546 | 0.147 | 0 |
| Calcrl | 0 | 4.32490026 | 0.449 | 0.051 | 0 |
| Man1a | 0 | 2.14114787 | 0.601 | 0.203 | 0 |
| Slc8a1 | 0 | 4.33927422 | 0.426 | 0.029 | 0 |
| Dip2c | 0 | 2.85541781 | 0.488 | 0.092 | 0 |
| Fam172a | 0 | 2.56929655 | 0.513 | 0.117 | 0 |
| Hpse2 | 0 | 4.26825368 | 0.451 | 0.056 | 0 |
| Focad | 0 | 4.20277823 | 0.43 | 0.037 | 0 |
| Itga1 | 0 | 2.93954265 | 0.473 | 0.081 | 0 |
| Ophn1 | 0 | 3.24689644 | 0.468 | 0.077 | 0 |
| Nrg2 | 0 | 3.78867501 | 0.433 | 0.042 | 0 |
| Igf2 | 0 | 1.79466361 | 0.514 | 0.124 | 0 |
| Cavin3 | 0 | 1.92419633 | 0.542 | 0.152 | 0 |
| Ecrg4 | 0 | 2.73477228 | 0.409 | 0.019 | 0 |
| Cdon | 0 | 2.86990557 | 0.488 | 0.099 | 0 |
| Dlg1 | 0 | 2.62194878 | 0.5 | 0.111 | 0 |
| Tshz2 | 0 | 3.3401285 | 0.467 | 0.079 | 0 |
| Syne1 | 0 | 2.30110962 | 0.568 | 0.18 | 0 |
| Apod | 0 | 1.41003783 | 0.899 | 0.512 | 0 |
| Mcub | 0 | 3.72023158 | 0.432 | 0.045 | 0 |
| Rspo1 | 0 | 3.54880056 | 0.433 | 0.049 | 0 |
| Edil3 | 0 | 2.59956776 | 0.635 | 0.252 | 0 |
| Exoc4 | 0 | 2.94834171 | 0.465 | 0.083 | 0 |
| Exoc6b | 0 | 2.6733922 | 0.486 | 0.104 | 0 |
| 9530026P05 | 0 | 3.92194611 | 0.42 | 0.038 | 0 |
| Mbd5 | 0 | 3.32001142 | 0.434 | 0.053 | 0 |
| Tomm6 | 0 | 2.14198923 | 0.513 | 0.133 | 0 |
| Daam2 | 0 | 2.66912419 | 0.476 | 0.097 | 0 |
| Ncoa7 | 0 | 2.97037066 | 0.468 | 0.091 | 0 |
| Ecpas | 0 | 4.09964618 | 0.4 | 0.025 | 0 |
| Tbc1d5 | 0 | 3.58739356 | 0.414 | 0.043 | 0 |
| Dio3os | 0 | 6.61899795 | 0.375 | 0.004 | 0 |
| Prkce | 0 | 3.6450864 | 0.417 | 0.047 | 0 |
| Tle5 | 0 | 3.96668019 | 0.396 | 0.026 | 0 |
| B3galt1 | 0 | 5.40942061 | 0.38 | 0.014 | 0 |
| 4932438A13 | 0 | 2.34824601 | 0.496 | 0.131 | 0 |
| Atrnl1 | 0 | 2.99352837 | 0.436 | 0.074 | 0 |
| Large1 | 0 | 3.13252451 | 0.414 | 0.052 | 0 |
| Pard3b | 0 | 2.80804533 | 0.452 | 0.091 | 0 |
| Ptprm | 0 | 2.58785888 | 0.462 | 0.103 | 0 |
| Tle4 | 0 | 2.45150482 | 0.475 | 0.116 | 0 |
| Ndufb1-ps | 0 | 3.1817381 | 0.398 | 0.043 | 0 |
| Cntn5 | 0 | 6.05819209 | 0.38 | 0.025 | 0 |
| Slco2a1 | 0 | 2.55923955 | 0.474 | 0.119 | 0 |
| Dipk2a | 0 | 4.914957 | 0.367 | 0.014 | 0 |
| Chl1 | 0 | 3.64211813 | 0.393 | 0.042 | 0 |
| Tcf4 | 0 | 1.84506965 | 0.922 | 0.572 | 0 |
| Nrg3 | 0 | 7.01618516 | 0.358 | 0.008 | 0 |
| Pitx1 | 0 | 2.497751 | 0.434 | 0.089 | 0 |
| Nfix | 0 | 1.33110842 | 0.919 | 0.575 | 0 |
| Pcdh9 | 0 | 5.18772492 | 0.364 | 0.02 | 0 |
| [illegible] | 0 | 3.20843179 | 0.392 | 0.049 | 0 |
| Pde5a | 0 | 3.06045766 | 0.407 | 0.066 | 0 |
| Sgcd | 0 | 2.6708282 | 0.428 | 0.087 | 0 |
| Wwox | 0 | 3.13949287 | 0.398 | 0.058 | 0 |
| Slc9a9 | 0 | 3.66454772 | 0.379 | 0.04 | 0 |
| 4933406I18 | 0 | 5.08661705 | 0.351 | 0.012 | 0 |
| Nhs | 0 | 2.91648893 | 0.406 | 0.073 | 0 |
| Cachd1 | 0 | 2.86988062 | 0.399 | 0.067 | 0 |
| Pde7b | 0 | 3.20682726 | 0.371 | 0.039 | 0 |
| Rnasek | 0 | 3.05298441 | 0.375 | 0.044 | 0 |
| Ncoa1 | 0 | 2.45441972 | 0.429 | 0.098 | 0 |
| Tut4 | 0 | 4.01177549 | 0.353 | 0.023 | 0 |
| Sorcs3 | 0 | 5.92396024 | 0.339 | 0.01 | 0 |
| 2210408F21 | 0 | 5.08208644 | 0.341 | 0.012 | 0 |
| Vti1a | 0 | 3.15630584 | 0.38 | 0.055 | 0 |
| Rnf180 | 0 | 5.30981085 | 0.333 | 0.011 | 0 |
| Tasor | 0 | 4.59296615 | 0.337 | 0.016 | 0 |
| Dnm3 | 0 | 3.29229333 | 0.374 | 0.054 | 0 |
| Ambra1 | 0 | 3.24970662 | 0.366 | 0.046 | 0 |
| Arhgap24 | 0 | 2.60776077 | 0.408 | 0.09 | 0 |
| Nr3c2 | 0 | 4.70432701 | 0.335 | 0.018 | 0 |
| Hdac8 | 0 | 3.68405667 | 0.346 | 0.032 | 0 |
| Gpcpd1 | 0 | 3.36586215 | 0.4 | 0.088 | 0 |
| Trerf1 | 0 | 4.60770716 | 0.326 | 0.015 | 0 |
| Sdk1 | 0 | 3.16709359 | 0.355 | 0.045 | 0 |
| Ror1 | 0 | 3.48298711 | 0.339 | 0.032 | 0 |
| [illegible] | [illegible] | [illegible] | [illegible] | [illegible] | [illegible] |

| Cluster 7 - Nascent Blastema | | | | | |
|---|---|---|---|---|---|
| Gene | p_val | avg_log2FC | pct.1 | pct.2 | p_val_adj |
| Sfrp4 | 0 | 4.25075954 | 0.78 | 0.103 | 0 |
| Csmd1 | 0 | 5.01708095 | 0.684 | 0.057 | 0 |
| Cmss1 | 0 | 2.10840163 | 0.764 | 0.161 | 0 |
| Plcb1 | 0 | 4.22260957 | 0.68 | 0.1 | 0 |
| Nme2 | 0 | 3.0504906 | 0.821 | 0.271 | 0 |
| Tnfsf11 | 0 | 3.69818265 | 0.608 | 0.091 | 0 |
| Ccn5 | 0 | 6.40226722 | 0.522 | 0.01 | 0 |
| Csgalnact1 | 0 | 3.86683114 | 0.592 | 0.095 | 0 |
| Mmp13 | 0 | 3.75335892 | 0.646 | 0.15 | 0 |
| Gas5 | 0 | 2.68736379 | 0.575 | 0.086 | 0 |
| Ptprd | 0 | 3.42159502 | 0.825 | 0.338 | 0 |
| Satb2 | 0 | 3.8312749 | 0.585 | 0.099 | 0 |
| Slit2 | 0 | 3.86538127 | 0.673 | 0.193 | 0 |
| Micos10 | 0 | 3.56489846 | 0.513 | 0.048 | 0 |
| AI506816 | 0 | 4.96590222 | 0.477 | 0.017 | 0 |
| Atp5mpl | 0 | 3.33982168 | 0.489 | 0.05 | 0 |
| Nav3 | 0 | 3.89486696 | 0.471 | 0.038 | 0 |
| Atp5md | 0 | 3.1723531 | 0.485 | 0.052 | 0 |
| Frmd4a | 0 | 3.06285304 | 0.505 | 0.088 | 0 |
| Galnt2 | 0 | 3.71829786 | 0.449 | 0.04 | 0 |
| Morrbid | 0 | 4.65069235 | 0.421 | 0.02 | 0 |
| Ndufb1-ps | 0 | 2.84477032 | 0.443 | 0.06 | 0 |
| Mid1 | 0 | 4.0091177 | 0.418 | 0.036 | 0 |
| Bnc2 | 0 | 3.75147776 | 0.453 | 0.071 | 0 |
| Cct6a | 0 | 3.50980632 | 0.416 | 0.04 | 0 |
| Timm23 | 0 | 3.13513745 | 0.403 | 0.048 | 0 |
| 2610307P16 | 0 | 5.04561646 | 0.36 | 0.012 | 0 |
| Dbndd2 | 0 | 3.93277544 | 0.377 | 0.03 | 0 |
| Tent5a | 0 | 3.83010301 | 0.358 | 0.029 | 0 |
| March3 | 0 | 3.65364131 | 0.365 | 0.042 | 0 |
| Pdzrn4 | 0 | 7.8593723 | 0.322 | 0.006 | 0 |
| Cemip2 | 0 | 3.74688267 | 0.336 | 0.029 | 0 |
| Ccn1 | 0 | 4.30573338 | 0.313 | 0.021 | 0 |
| 1110038B12 | 0 | 4.06485908 | 0.282 | 0.018 | 0 |
| Ly75 | 0 | 5.0492887 | 0.268 | 0.012 | 0 |
| Tenm4 | 0 | 4.746923 | 0.263 | 0.017 | 0 |
| Kng2 | 0 | 4.74959887 | 0.208 | 0.01 | 0 |
| Caln1 | 0 | 5.87342131 | 0.173 | 0.005 | 0 |
| Cd244a | 0 | 6.81834503 | 0.159 | 0.002 | 0 |
| [illegible] | [illegible] | [illegible] | [illegible] | [illegible] | [illegible] |

| | | | | | |
|---|---|---|---|---|---|
| Rbfox2 | 4.47E-229 | 1.82758797 | 0.558 | 0.22 | 8.96E-225 |
| Brinp1 | 1.10E-228 | 5.76982286 | 0.103 | 0.003 | 2.20E-224 |
| Nsd3 | 2.00E-228 | 1.79548627 | 0.43 | 0.122 | 4.02E-224 |
| Xrcc4 | 2.44E-228 | 2.89051051 | 0.229 | 0.035 | 4.90E-224 |
| Prxl2b | 5.98E-228 | 4.4350335 | 0.115 | 0.005 | 1.20E-223 |
| Schip1 | 8.30E-228 | 5.38377527 | 0.177 | 0.02 | 1.67E-223 |
| Prdm5 | 9.80E-228 | 2.45800127 | 0.32 | 0.072 | 1.97E-223 |
| Il16 | 1.49E-227 | 2.95542791 | 0.218 | 0.032 | 2.99E-223 |
| Cpa6 | 1.52E-227 | 2.60699065 | 0.297 | 0.062 | 3.05E-223 |
| Ntng1 | 1.72E-227 | 3.58018795 | 0.203 | 0.027 | 3.46E-223 |
| Ulk2 | 1.62E-226 | 2.19583094 | 0.387 | 0.105 | 3.25E-222 |
| Fndc3a | 2.17E-225 | 1.78111091 | 0.543 | 0.206 | 4.35E-221 |
| Nxn | 2.35E-225 | 2.23267194 | 0.382 | 0.103 | 4.71E-221 |
| Snhg11 | 6.13E-225 | 2.84130111 | 0.21 | 0.029 | 1.23E-220 |
| Kdm4c | 2.01E-224 | 2.78385197 | 0.23 | 0.036 | 4.04E-220 |
| Dkk2 | 8.60E-224 | 3.4397268 | 0.464 | 0.171 | 1.73E-219 |
| Usp25 | 1.83E-222 | 2.16740523 | 0.412 | 0.122 | 3.67E-218 |
| Nexmif | 2.70E-222 | 4.1620364 | 0.127 | 0.008 | 5.41E-218 |
| Scara3 | 3.17E-222 | 1.75276708 | 0.572 | 0.238 | 6.35E-218 |
| Crppa | 3.17E-222 | 3.68944788 | 0.143 | 0.011 | 6.35E-218 |
| Rab5if | 6.97E-222 | 2.8176692 | 0.206 | 0.028 | 1.40E-217 |
| Sdccag8 | 1.31E-221 | 2.93527438 | 0.216 | 0.032 | 2.62E-217 |
| Zfas1 | 1.69E-221 | 3.30920519 | 0.164 | 0.016 | 3.39E-217 |
| Kcnab1 | 3.12E-221 | 3.23382111 | 0.208 | 0.03 | 6.25E-217 |
| Stimate | 1.29E-220 | 3.96810577 | 0.127 | 0.008 | 2.59E-216 |
| Rab11fip2 | 4.40E-220 | 2.25340566 | 0.334 | 0.08 | 8.82E-216 |
| Arhgap12 | 5.72E-220 | 2.74181866 | 0.229 | 0.036 | 1.15E-215 |
| Gldn | 7.72E-220 | 1.93040703 | 0.562 | 0.229 | 1.55E-215 |
| Pakap.1 | 1.52E-219 | 2.72295642 | 0.202 | 0.027 | 3.06E-215 |
| Igf1r | 3.07E-219 | 1.73726654 | 0.561 | 0.221 | 6.16E-215 |
| Gnaq | 3.36E-219 | 1.90380298 | 0.471 | 0.159 | 6.74E-215 |
| Pknox2 | 3.81E-219 | 3.91555919 | 0.133 | 0.01 | 7.65E-215 |
| Gm4117 | 3.81E-218 | 3.71043389 | 0.153 | 0.014 | 7.64E-214 |
| Vwa8 | 5.98E-218 | 3.03297753 | 0.193 | 0.025 | 1.20E-213 |
| Maml3 | 5.05E-217 | 2.68514388 | 0.267 | 0.052 | 1.01E-212 |
| Afdn | 5.78E-217 | 2.48679154 | 0.261 | 0.048 | 1.16E-212 |
| Plaat3 | 6.52E-217 | 4.40621225 | 0.11 | 0.005 | 1.31E-212 |
| Slc25a21 | 2.50E-216 | 3.99126143 | 0.141 | 0.012 | 5.02E-212 |
| [illegible] | [illegible] | [illegible] | [illegible] | [illegible] | [illegible] |

| | | | | | |
|---|---|---|---|---|---|
| Lrp1 | 2.08E-58 | 0.81382334 | 0.842 | 0.724 | 4.18E-54 |
| Fmnl2 | 2.20E-58 | 1.602539 | 0.399 | 0.185 | 4.42E-54 |
| Ecm1 | 2.67E-58 | 0.80885454 | 0.788 | 0.581 | 5.36E-54 |
| Snx24 | 3.04E-58 | 2.04183866 | 0.172 | 0.044 | 6.11E-54 |
| Kmt5a | 3.42E-58 | 1.74128658 | 0.201 | 0.058 | 6.86E-54 |
| Serpine2 | 3.54E-58 | 1.11686328 | 0.78 | 0.541 | 7.11E-54 |
| Basp1 | 3.81E-58 | 1.27543235 | 0.405 | 0.183 | 7.64E-54 |
| Rasal2 | 4.22E-58 | 1.70496628 | 0.254 | 0.086 | 8.48E-54 |
| Stard3nl | 4.29E-58 | 1.42555226 | 0.32 | 0.124 | 8.62E-54 |
| Cacna2d1 | 5.08E-58 | 1.48885879 | 0.422 | 0.2 | 1.02E-53 |
| Prickle1 | 5.23E-58 | 1.961841 | 0.24 | 0.079 | 1.05E-53 |
| Lcorl | 9.08E-58 | 1.7088916 | 0.227 | 0.072 | 1.82E-53 |
| Pfkfb3 | 9.12E-58 | 1.90765642 | 0.197 | 0.057 | 1.83E-53 |
| Myo1d | 9.17E-58 | 1.55406348 | 0.405 | 0.191 | 1.84E-53 |
| Ryk | 1.11E-57 | 1.03351313 | 0.577 | 0.329 | 2.23E-53 |
| Mrc2 | 1.18E-57 | 1.14600052 | 0.476 | 0.237 | 2.36E-53 |
| Memo1 | 1.22E-57 | 1.51820804 | 0.294 | 0.11 | 2.44E-53 |
| Stim2 | 1.38E-57 | 1.77604329 | 0.236 | 0.077 | 2.77E-53 |
| St3gal3 | 2.42E-57 | 2.18413901 | 0.159 | 0.04 | 4.86E-53 |
| Gspt1 | 2.91E-57 | 1.08721611 | 0.488 | 0.246 | 5.84E-53 |
| Adarb1 | 6.45E-57 | 1.54139088 | 0.297 | 0.112 | 1.29E-52 |
| Tacc1 | 7.30E-57 | 1.13646417 | 0.502 | 0.263 | 1.47E-52 |
| Ilrun | 8.15E-57 | 2.1513518 | 0.137 | 0.031 | 1.64E-52 |
| Csf1 | 1.34E-56 | 1.21602357 | 0.386 | 0.171 | 2.69E-52 |
| Ccdc85b | 1.51E-56 | 1.66928522 | 0.226 | 0.072 | 3.04E-52 |
| Dlg1 | 1.63E-56 | 1.38691812 | 0.343 | 0.142 | 3.28E-52 |
| Clint1 | 1.93E-56 | 1.11160533 | 0.466 | 0.227 | 3.88E-52 |
| Cast | 2.05E-56 | 1.11150306 | 0.482 | 0.243 | 4.11E-52 |
| Fam172a | 3.20E-56 | 1.30382587 | 0.355 | 0.148 | 6.43E-52 |
| Nsf | 3.55E-56 | 1.68283825 | 0.243 | 0.082 | 7.12E-52 |
| Sulf1 | 4.04E-56 | 1.24937451 | 0.478 | 0.252 | 8.12E-52 |
| Lars2 | 4.05E-56 | 0.93937302 | 0.655 | 0.405 | 8.13E-52 |
| Psme4 | 4.08E-56 | 1.24958238 | 0.403 | 0.183 | 8.19E-52 |
| Cdk17 | 7.36E-56 | 1.80863869 | 0.217 | 0.068 | 1.48E-51 |
| Fkbp11 | 7.59E-56 | 1.15596061 | 0.451 | 0.217 | 1.52E-51 |
| Itpr1 | 8.03E-56 | 1.62329473 | 0.285 | 0.107 | 1.61E-51 |
| Tent2 | 1.62E-55 | 2.13652099 | 0.133 | 0.029 | 3.24E-51 |
| Pdia4 | 1.84E-55 | 1.25915443 | 0.358 | 0.151 | 3.69E-51 |
| [illegible] | [illegible] | [illegible] | [illegible] | [illegible] | [illegible] |

| | | | | | |
|---|---|---|---|---|---|
| Chd2 | 1.72E-117 | 1.58681343 | 0.364 | 0.145 | 3.46E-113 |
| Rptor | 7.56E-117 | 2.26886558 | 0.174 | 0.039 | 1.52E-112 |
| Gpatch2 | 9.49E-117 | 2.25732827 | 0.177 | 0.04 | 1.90E-112 |
| Bmp7 | 9.68E-117 | 2.61386163 | 0.13 | 0.022 | 1.94E-112 |
| Il1rap | 1.57E-116 | 2.6365194 | 0.147 | 0.028 | 3.14E-112 |
| Nsmaf | 4.10E-116 | 2.32078054 | 0.184 | 0.044 | 8.23E-112 |
| Wipf1 | 6.43E-116 | 1.46417985 | 0.406 | 0.175 | 1.29E-111 |
| Eps15 | 1.01E-115 | 1.6603705 | 0.302 | 0.105 | 2.03E-111 |
| Tcf7l1 | 1.48E-115 | 1.74942192 | 0.282 | 0.094 | 2.98E-111 |
| Fbxl5 | 1.72E-115 | 1.59464302 | 0.314 | 0.112 | 3.45E-111 |
| Rap1gds1 | 1.74E-115 | 1.76522057 | 0.293 | 0.101 | 3.50E-111 |
| Zfp280d | 2.08E-115 | 1.74555384 | 0.289 | 0.098 | 4.18E-111 |
| Rngtt | 2.85E-115 | 2.04108258 | 0.199 | 0.05 | 5.72E-111 |
| Tspan14 | 2.94E-115 | 2.32395392 | 0.184 | 0.044 | 5.91E-111 |
| Ncam1 | 3.50E-115 | 1.39186396 | 0.458 | 0.217 | 7.02E-111 |
| Dsel | 3.58E-115 | 1.72510966 | 0.286 | 0.096 | 7.18E-111 |
| Rnf38 | 9.04E-115 | 1.89244247 | 0.227 | 0.064 | 1.81E-110 |
| Psme4 | 1.03E-114 | 1.40012445 | 0.401 | 0.171 | 2.07E-110 |
| Mindy3 | 3.03E-114 | 2.20114544 | 0.168 | 0.037 | 6.07E-110 |
| A4galt | 3.25E-114 | 1.99877813 | 0.203 | 0.052 | 6.53E-110 |
| Atp2a2 | 3.46E-114 | 1.04344374 | 0.62 | 0.361 | 6.94E-110 |
| Qk | 4.94E-114 | 1.14594623 | 0.6 | 0.351 | 9.92E-110 |
| Zfp445 | 2.05E-113 | 2.1988072 | 0.162 | 0.034 | 4.11E-109 |
| Nup98 | 2.82E-113 | 1.52524137 | 0.288 | 0.096 | 5.67E-109 |
| Usp9x | 4.11E-113 | 1.182733 | 0.497 | 0.246 | 8.25E-109 |
| Mdfic | 4.36E-113 | 1.53013541 | 0.345 | 0.134 | 8.75E-109 |
| Hdac7 | 6.68E-113 | 1.85392943 | 0.235 | 0.069 | 1.34E-108 |
| Flrt3 | 1.12E-112 | 2.14225175 | 0.152 | 0.031 | 2.24E-108 |
| Pdzd2 | 2.53E-112 | 2.04236912 | 0.202 | 0.053 | 5.08E-108 |
| Timm23 | 2.60E-112 | 1.89074605 | 0.2 | 0.052 | 5.21E-108 |
| Parva | 2.75E-112 | 1.27193292 | 0.484 | 0.241 | 5.52E-108 |
| Fgf10 | 3.86E-112 | 2.41565734 | 0.325 | 0.127 | 7.75E-108 |
| Cdh11 | 5.85E-112 | 0.86903975 | 0.77 | 0.52 | 1.17E-107 |
| Pabpn1 | 8.46E-112 | 1.39483949 | 0.361 | 0.143 | 1.70E-107 |
| Rreb1 | 9.42E-112 | 2.08656589 | 0.191 | 0.048 | 1.89E-107 |
| Gng12 | 1.01E-111 | 1.22936816 | 0.47 | 0.227 | 2.04E-107 |
| Usp47 | 1.19E-111 | 1.57311422 | 0.331 | 0.126 | 2.39E-107 |
| Svep1 | 1.46E-111 | 1.61524972 | 0.536 | 0.294 | 2.93E-107 |
| Sp4 | 1.79E-111 | 2.50052169 | 0.144 | 0.028 | 3.59E-107 |
| Pde7a | 3.55E-111 | 1.95438589 | 0.208 | 0.056 | 7.13E-107 |
| Baz2b | 5.61E-111 | 1.33861165 | 0.455 | 0.219 | 1.13E-106 |
| Ern1 | 8.98E-111 | 2.11289477 | 0.186 | 0.046 | 1.80E-106 |
| Map2k4 | 1.02E-110 | 1.7701817 | 0.268 | 0.088 | 2.04E-106 |
| Stac2 | 1.09E-110 | 2.9707157 | 0.101 | 0.014 | 2.18E-106 |
| Dapk1 | 1.82E-110 | 1.94642016 | 0.255 | 0.082 | 3.65E-106 |
| Arhgef11 | 2.25E-110 | 2.01646165 | 0.203 | 0.054 | 4.52E-106 |
| Dennd1b | 4.67E-110 | 2.05098841 | 0.205 | 0.055 | 9.37E-106 |
| Ptpa | 9.25E-110 | 2.07222596 | 0.181 | 0.044 | 1.86E-105 |
| Sh3kbp1 | 1.68E-109 | 1.48900512 | 0.421 | 0.197 | 3.37E-105 |
| Acadsb | 2.50E-109 | 1.58308813 | 0.306 | 0.111 | 5.03E-105 |
| Atf7 | 3.21E-109 | 1.57857922 | 0.33 | 0.127 | 6.44E-105 |
| Clasp2 | 7.35E-109 | 1.55209069 | 0.318 | 0.119 | 1.48E-104 |
| Txnip | 1.75E-108 | 1.16235879 | 0.461 | 0.217 | 3.50E-104 |
| Atrx | 1.93E-108 | 1.13594496 | 0.537 | 0.289 | 3.88E-104 |
| Ascc3 | 1.95E-108 | 1.64271798 | 0.302 | 0.11 | 3.92E-104 |
| Mycbp2 | 2.42E-108 | 1.17644667 | 0.55 | 0.306 | 4.87E-104 |
| Gdf6 | 2.56E-108 | 2.78719636 | 0.111 | 0.017 | 5.14E-104 |
| Tbx2 | 4.25E-108 | 1.68154865 | 0.297 | 0.107 | 8.52E-104 |
| Usp40 | 4.57E-108 | 2.11563776 | 0.184 | 0.046 | 9.18E-104 |
| Akap8l | 5.26E-108 | 1.72274276 | 0.253 | 0.08 | 1.06E-103 |
| Map3k14 | 8.14E-108 | 2.21906735 | 0.165 | 0.038 | 1.63E-103 |
| Peli2 | 9.10E-108 | 1.96715849 | 0.215 | 0.061 | 1.83E-103 |
| Aarsd1 | 9.58E-108 | 2.60635986 | 0.12 | 0.02 | 1.92E-103 |
| Atl3 | 9.64E-108 | 1.61355747 | 0.294 | 0.105 | 1.93E-103 |
| Srr | 1.82E-107 | 2.29140246 | 0.152 | 0.032 | 3.65E-103 |
| Aff4 | 1.82E-107 | 1.20901368 | 0.473 | 0.233 | 3.65E-103 |
| Cxadr | 1.91E-107 | 2.07784266 | 0.175 | 0.042 | 3.83E-103 |
| Zzef1 | 2.23E-107 | 1.77098665 | 0.24 | 0.074 | 4.48E-103 |
| Sufu | 2.28E-107 | 2.29405622 | 0.153 | 0.033 | 4.58E-103 |
| Tlk1 | 2.62E-107 | 1.62320764 | 0.307 | 0.114 | 5.26E-103 |
| Cep170 | 3.75E-107 | 1.70803486 | 0.275 | 0.094 | 7.53E-103 |
| Elovl6 | 3.88E-107 | 2.15128923 | 0.191 | 0.05 | 7.78E-103 |
| Cdk13 | 4.36E-107 | 1.59314817 | 0.298 | 0.108 | 8.75E-103 |
| Clec16a | 4.47E-107 | 2.61872962 | 0.121 | 0.021 | 8.96E-103 |
| Zfhx3 | 4.49E-107 | 1.13926821 | 0.553 | 0.306 | 9.01E-103 |
| Tmtc1 | 4.57E-107 | 2.4440422 | 0.154 | 0.033 | 9.17E-103 |
| Ptbp2 | 4.78E-107 | 1.63499788 | 0.304 | 0.112 | 9.59E-103 |
| Zmym6 | 9.57E-107 | 2.7982485 | 0.104 | 0.015 | 1.92E-102 |
| Gpc4 | 2.05E-106 | 1.51903398 | 0.403 | 0.184 | 4.12E-102 |
| Shprh | 2.21E-106 | 2.21744068 | 0.16 | 0.036 | 4.43E-102 |
| Ddx17 | 3.30E-106 | 1.24972871 | 0.478 | 0.238 | 6.63E-102 |
| Dmxl1 | 4.05E-106 | 1.60607031 | 0.278 | 0.096 | 8.13E-102 |
| Fnbp1 | 4.23E-106 | 1.89438864 | 0.215 | 0.062 | 8.48E-102 |
| Bnc2 | 6.33E-106 | 1.4409452 | 0.241 | 0.074 | 1.27E-101 |
| Ppp1r12b | 9.68E-106 | 2.35703099 | 0.152 | 0.033 | 1.94E-101 |
| Nfatc4 | 1.17E-105 | 1.36488892 | 0.353 | 0.143 | 2.35E-101 |
| Numa1 | 1.30E-105 | 1.57240939 | 0.293 | 0.105 | 2.61E-101 |
| Cdyl2 | 1.34E-105 | 2.34068318 | 0.151 | 0.032 | 2.68E-101 |
| Auts2 | 1.93E-105 | 1.43933812 | 0.487 | 0.262 | 3.88E-101 |
| Ttc7b | 2.41E-105 | 2.33025304 | 0.146 | 0.03 | 4.83E-101 |
| Rex1bd | 2.75E-105 | 1.57196809 | 0.278 | 0.096 | 5.51E-101 |
| Farp2 | 4.78E-105 | 2.07124364 | 0.169 | 0.04 | 9.60E-101 |
| Wnt5a | 9.78E-105 | 0.92581405 | 0.659 | 0.406 | 1.96E-100 |
| Vps13d | 1.10E-104 | 2.0082259 | 0.199 | 0.054 | 2.20E-100 |
| Mfsd14b | 1.40E-104 | 2.14809263 | 0.157 | 0.035 | 2.80E-100 |
| Nbas | 1.54E-104 | 2.09206867 | 0.171 | 0.041 | 3.10E-100 |
| Chd6 | 2.08E-104 | 1.41155931 | 0.348 | 0.141 | 4.17E-100 |
| Abhd18 | 2.62E-104 | 2.46045676 | 0.138 | 0.028 | 5.25E-100 |
| Cnot4 | 2.75E-104 | 1.44951446 | 0.372 | 0.159 | 5.52E-100 |
| Pou2f1 | 3.49E-104 | 2.04718349 | 0.192 | 0.051 | 7.01E-100 |
| Prrx2 | 4.69E-104 | 0.79263027 | 0.743 | 0.484 | 9.41E-100 |
| Hdgfl3 | 7.21E-104 | 1.99557366 | 0.173 | 0.042 | 1.45E-99 |
| Crebbp | 1.24E-103 | 1.16276546 | 0.49 | 0.25 | 2.49E-99 |
| Ccn2 | 1.31E-103 | 2.37040392 | 0.112 | 0.018 | 2.62E-99 |
| Kcnt2 | 1.87E-103 | 2.5246157 | 0.12 | 0.021 | 3.76E-99 |
| Rmdn1 | 1.91E-103 | 2.24482148 | 0.154 | 0.034 | 3.84E-99 |
| Ntn5 | 1.96E-103 | 2.46894909 | 0.12 | 0.021 | 3.93E-99 |
| Pi4ka | 3.33E-103 | 1.95254034 | 0.2 | 0.056 | 6.69E-99 |
| Herc1 | 6.56E-103 | 1.51264133 | 0.33 | 0.132 | 1.32E-98 |
| Hook3 | 9.80E-103 | 1.18955077 | 0.462 | 0.228 | 1.97E-98 |
| Fbxl20 | 2.79E-102 | 1.81851009 | 0.228 | 0.07 | 5.60E-98 |
| Ppm1h | 3.63E-102 | 2.70498833 | 0.116 | 0.02 | 7.29E-98 |
| Fmnl2 | 3.77E-102 | 1.47288829 | 0.387 | 0.174 | 7.57E-98 |
| Abcb7 | 5.66E-102 | 2.47520125 | 0.125 | 0.023 | 1.14E-97 |
| Smarcc2 | 6.89E-102 | 1.63243293 | 0.281 | 0.101 | 1.38E-97 |
| Xpo7 | 7.45E-102 | 1.749053 | 0.235 | 0.074 | 1.49E-97 |
| Itgb8 | 1.18E-101 | 2.24742957 | 0.157 | 0.036 | 2.37E-97 |
| Atf7ip | 1.20E-101 | 1.60869978 | 0.294 | 0.109 | 2.41E-97 |
| [illegible] | 1.50E-101 | 1.76940506 | 0.235 | 0.074 | 3.02E-97 |
| Ano6 | 1.80E-101 | 1.25716553 | 0.459 | 0.228 | 3.61E-97 |
| Micu2 | 2.40E-101 | 1.81913674 | 0.224 | 0.069 | 4.81E-97 |
| Ccn1 | 3.67E-101 | 2.02401775 | 0.132 | 0.025 | 7.36E-97 |
| Cnot2 | 4.76E-101 | 1.77775586 | 0.251 | 0.084 | 9.56E-97 |
| Cxcl5 | 6.22E-101 | 2.82913211 | 0.183 | 0.049 | 1.25E-96 |
| Lin52 | 1.05E-100 | 2.39017926 | 0.136 | 0.028 | 2.10E-96 |
| Mrpl53 | 2.90E-100 | 2.53992241 | 0.113 | 0.019 | 5.82E-96 |
| Mindy1 | 2.91E-100 | 2.09269871 | 0.156 | 0.036 | 5.84E-96 |
| Cavin1 | 3.21E-100 | 1.01327578 | 0.364 | 0.146 | 6.43E-96 |
| Plec | 3.45E-100 | 0.89455072 | 0.628 | 0.379 | 6.92E-96 |
| Syne3 | 3.68E-100 | 2.34727445 | 0.136 | 0.028 | 7.38E-96 |
| Mta3 | 4.16E-100 | 1.94508453 | 0.199 | 0.056 | 8.34E-96 |
| Srgap2 | 4.27E-100 | 1.74529471 | 0.258 | 0.089 | 8.57E-96 |
| Vps28 | 4.31E-100 | 1.80782342 | 0.205 | 0.059 | 8.65E-96 |
| Herc4 | 5.24E-100 | 1.82619896 | 0.208 | 0.061 | 1.05E-95 |
| Camk2d | 6.97E-100 | 1.33915293 | 0.463 | 0.237 | 1.40E-95 |
| Gpr173 | 8.55E-100 | 2.61563219 | 0.112 | 0.019 | 1.72E-95 |
| Herc2 | 8.88E-100 | 1.4107039 | 0.359 | 0.154 | 1.78E-95 |
| Arih1 | 1.29E-99 | 1.27186876 | 0.416 | 0.194 | 2.60E-95 |
| Gigyf2 | 2.39E-99 | 1.55225309 | 0.303 | 0.116 | 4.79E-95 |
| Ythdf3 | 2.55E-99 | 1.55349984 | 0.302 | 0.115 | 5.12E-95 |
| Tnrc6a | 2.76E-99 | 1.13177174 | 0.478 | 0.242 | 5.53E-95 |
| Kirrel | 3.21E-99 | 1.37630709 | 0.39 | 0.178 | 6.43E-95 |
| Stat5a | 7.61E-99 | 1.95201988 | 0.194 | 0.054 | 1.53E-94 |
| Washc2 | 9.96E-99 | 2.00774572 | 0.169 | 0.042 | 2.00E-94 |
| Erg28 | 1.56E-98 | 2.41850799 | 0.128 | 0.025 | 3.14E-94 |
| Eps15l1 | 2.55E-98 | 2.16060278 | 0.155 | 0.036 | 5.12E-94 |
| Spaca6 | 3.02E-98 | 1.74988333 | 0.23 | 0.073 | 6.06E-94 |
| Mapk14 | 3.69E-98 | 1.78412446 | 0.227 | 0.072 | 7.41E-94 |
| Myo6 | 5.09E-98 | 1.57274448 | 0.29 | 0.109 | 1.02E-93 |
| C2cd5 | 5.36E-98 | 2.20619867 | 0.156 | 0.037 | 1.08E-93 |
| Prr14l | 5.50E-98 | 2.09298096 | 0.16 | 0.038 | 1.10E-93 |
| Abca1 | 5.62E-98 | 1.41114004 | 0.347 | 0.146 | 1.13E-93 |
| 9330159F19 | 5.70E-98 | 2.30893838 | 0.141 | 0.03 | 1.14E-93 |
| Anks1 | 6.30E-98 | 2.50129302 | 0.119 | 0.022 | 1.27E-93 |
| Ppp6r3 | 8.77E-98 | 1.62396029 | 0.267 | 0.094 | 1.76E-93 |
| Sfmbt1 | 1.33E-97 | 1.91445452 | 0.199 | 0.057 | 2.67E-93 |
| Zfp950 | 2.99E-97 | 1.7515237 | 0.221 | 0.069 | 6.00E-93 |
| Igsf10 | 3.05E-97 | 0.94268726 | 0.562 | 0.307 | 6.11E-93 |
| Zfp160 | 9.20E-97 | 2.44056729 | 0.119 | 0.022 | 1.85E-92 |
| Ssbp3 | 1.04E-96 | 1.20515383 | 0.427 | 0.205 | 2.08E-92 |
| Gpha2 | 1.06E-96 | 3.01601356 | 0.101 | 0.016 | 2.13E-92 |
| Selenop | 1.10E-96 | 1.0446347 | 0.356 | 0.144 | 2.20E-92 |
| Calm3 | 1.54E-96 | 1.08953826 | 0.479 | 0.245 | 3.09E-92 |
| Phf14 | 1.78E-96 | 1.46929806 | 0.354 | 0.154 | 3.57E-92 |
| Kdm3b | 3.63E-96 | 1.9770472 | 0.179 | 0.048 | 7.28E-92 |
| Smurf2 | 5.58E-96 | 1.42715173 | 0.343 | 0.145 | 1.12E-91 |
| Sod3 | 6.38E-96 | 0.77707595 | 0.65 | 0.381 | 1.28E-91 |
| Zfp398 | 7.62E-96 | 2.23784455 | 0.146 | 0.033 | 1.53E-91 |
| Chm | 7.71E-96 | 1.98339308 | 0.176 | 0.047 | 1.55E-91 |
| Adcy9 | 7.90E-96 | 2.53597053 | 0.116 | 0.021 | 1.59E-91 |
| Coq8b | 2.06E-95 | 2.39499628 | 0.116 | 0.021 | 4.14E-91 |
| Helz | 2.35E-95 | 1.56997451 | 0.27 | 0.098 | 4.72E-91 |
| Map2k5 | 3.17E-95 | 1.90465595 | 0.195 | 0.056 | 6.36E-91 |
| Phf8 | 3.98E-95 | 1.88488609 | 0.191 | 0.054 | 7.98E-91 |
| Rnf150 | 7.29E-95 | 2.15992278 | 0.148 | 0.034 | 1.46E-90 |
| Thap12 | 8.47E-95 | 1.8351735 | 0.186 | 0.051 | 1.70E-90 |
| Dock6 | 1.35E-94 | 1.99597881 | 0.181 | 0.049 | 2.71E-90 |
| Ddr2 | 3.03E-94 | 1.16198153 | 0.469 | 0.243 | 6.08E-90 |
| Cyld | 3.26E-94 | 1.80294595 | 0.213 | 0.066 | 6.53E-90 |
| Lrp6 | 3.29E-94 | 1.31507332 | 0.353 | 0.153 | 6.60E-90 |
| Fbn1 | 3.84E-94 | 0.67503647 | 0.678 | 0.406 | 7.70E-90 |
| Vps41 | 6.30E-94 | 1.62664133 | 0.259 | 0.092 | 1.26E-89 |
| R3hdm2 | 6.36E-94 | 1.40959093 | 0.325 | 0.133 | 1.28E-89 |
| Trir | 7.03E-94 | 1.41039707 | 0.281 | 0.104 | 1.41E-89 |
| Amotl1 | 8.09E-94 | 1.10669337 | 0.487 | 0.257 | 1.62E-89 |
| Mxi1 | 1.60E-93 | 1.59826802 | 0.306 | 0.123 | 3.21E-89 |
| Kat6a | 1.60E-93 | 1.63684575 | 0.249 | 0.086 | 3.22E-89 |
| Kiz | 2.13E-93 | 2.06877115 | 0.166 | 0.043 | 4.27E-89 |
| Ufd1 | 2.87E-93 | 1.97730234 | 0.165 | 0.042 | 5.75E-89 |
| Trip12 | 3.39E-93 | 1.25848225 | 0.42 | 0.206 | 6.81E-89 |
| Ncor1 | 3.88E-93 | 1.02576124 | 0.556 | 0.324 | 7.80E-89 |
| Olfml1 | 7.23E-93 | 2.11415234 | 0.152 | 0.036 | 1.45E-88 |
| R3hcc1l | 7.37E-93 | 2.28021701 | 0.138 | 0.031 | 1.48E-88 |
| B3gnt5 | 9.08E-93 | 2.56707975 | 0.107 | 0.018 | 1.82E-88 |
| Nr2c2 | 1.06E-92 | 1.63268557 | 0.262 | 0.095 | 2.12E-88 |
| Tbc1d4 | 1.25E-92 | 2.40245086 | 0.129 | 0.027 | 2.50E-88 |
| Rfx7 | 2.38E-92 | 1.67488609 | 0.234 | 0.078 | 4.77E-88 |
| Cdkn1b | 2.79E-92 | 1.20901368 | 0.364 | 0.159 | 5.60E-88 |
| Bmp5 | 3.07E-92 | 1.35180854 | 0.54 | 0.336 | 6.17E-88 |
| Ogt | 3.75E-92 | 1.27267614 | 0.394 | 0.185 | 7.52E-88 |
| [illegible] | 4.90E-92 | 2.33293531 | 0.128 | 0.027 | 9.82E-88 |
| Cyb5a | 5.01E-92 | 0.79822699 | 0.706 | 0.478 | 1.01E-87 |
| Cep128 | 7.42E-92 | 2.51516377 | 0.103 | 0.017 | 1.49E-87 |
| Cd248 | 7.46E-92 | 1.21213906 | 0.383 | 0.173 | 1.50E-87 |
| Dcaf5 | 9.51E-92 | 1.95983006 | 0.176 | 0.048 | 1.91E-87 |
| Atp7a | 2.07E-91 | 2.18078635 | 0.144 | 0.034 | 4.15E-87 |
| Npepps | 2.35E-91 | 1.30374813 | 0.369 | 0.167 | 4.71E-87 |
| Fam135a | 2.36E-91 | 1.90706275 | 0.195 | 0.058 | 4.73E-87 |
| Zeb1 | 3.11E-91 | 1.21232051 | 0.407 | 0.197 | 6.25E-87 |
| Ube4a | 5.79E-91 | 1.88273918 | 0.183 | 0.052 | 1.16E-86 |
| Nsmce2 | 5.95E-91 | 1.57733043 | 0.307 | 0.126 | 1.19E-86 |
| Trim2 | 6.09E-91 | 1.4606739 | 0.293 | 0.115 | 1.22E-86 |
| Fam91a1 | 6.17E-91 | 2.08761649 | 0.152 | 0.037 | 1.24E-86 |
| Dse | 8.65E-91 | 1.2119091 | 0.414 | 0.201 | 1.73E-86 |
| Ppp2r3a | 9.06E-91 | 1.48040831 | 0.299 | 0.119 | 1.82E-86 |
| Dcaf1 | 1.05E-90 | 2.29728254 | 0.123 | 0.025 | 2.11E-86 |
| Mdm4 | 1.20E-90 | 1.61512174 | 0.248 | 0.087 | 2.40E-86 |
| Kremen1 | 1.56E-90 | 1.27443466 | 0.354 | 0.155 | 3.13E-86 |
| 6720427I07R | 2.33E-90 | 2.26379912 | 0.127 | 0.026 | 4.67E-86 |
| Snx13 | 3.69E-90 | 1.95629357 | 0.184 | 0.053 | 7.40E-86 |
| Aak1 | 3.74E-90 | 1.52330343 | 0.272 | 0.102 | 7.51E-86 |
| Stx18 | 3.93E-90 | 1.3944642 | 0.337 | 0.145 | 7.89E-86 |
| Map3k2 | 3.94E-90 | 1.35592695 | 0.326 | 0.136 | 7.90E-86 |
| 5930430L01 | 4.07E-90 | 2.10636721 | 0.146 | 0.035 | 8.17E-86 |
| Stxbp5 | 4.11E-90 | 2.07523157 | 0.156 | 0.039 | 8.24E-86 |
| Glg1 | 8.85E-90 | 1.08428215 | 0.472 | 0.249 | 1.78E-85 |
| Plce1 | 9.48E-90 | 2.00862719 | 0.163 | 0.043 | 1.90E-85 |
| Araf | 1.17E-89 | 1.60268559 | 0.271 | 0.103 | 2.36E-85 |
| Ctif | 1.33E-89 | 1.97919347 | 0.169 | 0.046 | 2.67E-85 |
| Fbxo2 | 1.56E-89 | 1.90618439 | 0.183 | 0.052 | 3.14E-85 |
| Capn1 | 2.22E-89 | 1.99901887 | 0.167 | 0.044 | 4.45E-85 |
| Dock7 | 2.33E-89 | 1.48909479 | 0.305 | 0.126 | 4.68E-85 |
| Nhlrc2 | 4.14E-89 | 2.06485243 | 0.155 | 0.039 | 8.30E-85 |
| Tanc2 | 4.55E-89 | 2.1342818 | 0.17 | 0.047 | 9.13E-85 |
| Mosmo | 5.85E-89 | 1.78152067 | 0.183 | 0.052 | 1.17E-84 |
| Mcc | 8.25E-89 | 1.5491775 | 0.266 | 0.099 | 1.66E-84 |
| Zc3h6 | 1.20E-88 | 1.91391497 | 0.185 | 0.054 | 2.40E-84 |
| Lamp2 | 1.24E-88 | 0.66465291 | 0.789 | 0.579 | 2.50E-84 |
| Itsn2 | 1.53E-88 | 1.71589694 | 0.211 | 0.067 | 3.07E-84 |
| Grcc10 | 2.96E-88 | 1.38831252 | 0.317 | 0.132 | 5.95E-84 |
| Jcad | 4.11E-88 | 2.22728785 | 0.118 | 0.024 | 8.26E-84 |
| Snhg1 | 6.65E-88 | 2.25614272 | 0.114 | 0.022 | 1.33E-83 |
| Ccdc136 | 7.10E-88 | 2.4937058 | 0.106 | 0.019 | 1.42E-83 |
| Ccny | 8.23E-88 | 1.40608788 | 0.307 | 0.126 | 1.65E-83 |
| Tasp1 | 8.33E-88 | 1.90235239 | 0.182 | 0.053 | 1.67E-83 |
| Yes1 | 1.70E-87 | 1.51979382 | 0.257 | 0.094 | 3.42E-83 |
| Mcrip1 | 1.98E-87 | 1.52841403 | 0.233 | 0.079 | 3.98E-83 |
| Ttc23 | 2.41E-87 | 2.07075616 | 0.152 | 0.039 | 4.83E-83 |
| Ick | 2.91E-87 | 1.57783663 | 0.245 | 0.087 | 5.84E-83 |
| Prrc2c | 3.48E-87 | 0.8274201 | 0.637 | 0.397 | 6.98E-83 |
| Trim33 | 3.60E-87 | 1.70265432 | 0.227 | 0.078 | 7.23E-83 |
| Sap130 | 3.66E-87 | 2.2622768 | 0.13 | 0.029 | 7.34E-83 |
| Naxd | 4.13E-87 | 1.81019083 | 0.176 | 0.049 | 8.29E-83 |
| Deptor | 4.73E-87 | 2.09546515 | 0.145 | 0.035 | 9.50E-83 |
| Cntln | 6.05E-87 | 1.52295355 | 0.263 | 0.098 | 1.21E-82 |
| Rab3gap1 | 6.50E-87 | 1.66946913 | 0.218 | 0.072 | 1.30E-82 |
| Ccdc85b | 7.57E-87 | 1.68464482 | 0.206 | 0.065 | 1.52E-82 |
| Kif26b | 1.67E-86 | 2.37107044 | 0.117 | 0.024 | 3.35E-82 |
| Tbcd | 1.91E-86 | 1.8725138 | 0.183 | 0.053 | 3.83E-82 |
| Commd10 | 2.97E-86 | 1.82571201 | 0.207 | 0.067 | 5.96E-82 |
| Ranbp9 | 3.65E-86 | 1.5818249 | 0.253 | 0.093 | 7.33E-82 |
| Slc12a6 | 3.99E-86 | 1.97830698 | 0.165 | 0.045 | 8.01E-82 |
| Etl4 | 5.44E-86 | 2.13403124 | 0.219 | 0.075 | 1.09E-81 |
| Tmod2 | 6.83E-86 | 1.45476069 | 0.256 | 0.094 | 1.37E-81 |
| Ankrd28 | 9.85E-86 | 1.43309724 | 0.275 | 0.106 | 1.98E-81 |
| Zfp280c | 1.10E-85 | 2.31076993 | 0.119 | 0.025 | 2.21E-81 |
| Cpq | 1.12E-85 | 1.20680729 | 0.471 | 0.261 | 2.26E-81 |
| Kidins220 | 1.15E-85 | 1.60375676 | 0.24 | 0.085 | 2.30E-81 |
| Fbxl2 | 1.17E-85 | 2.27796104 | 0.127 | 0.028 | 2.35E-81 |
| Tceal9 | 2.37E-85 | 0.86187172 | 0.395 | 0.176 | 4.77E-81 |
| Bcl6 | 3.40E-85 | 1.52792373 | 0.261 | 0.098 | 6.83E-81 |
| Pnpt1 | 3.72E-85 | 2.15074049 | 0.14 | 0.034 | 7.46E-81 |
| Sptbn1 | 5.68E-85 | 0.86049997 | 0.596 | 0.364 | 1.14E-80 |
| Ntrk3 | 5.74E-85 | 2.799047 | 0.107 | 0.02 | 1.15E-80 |
| Mical3 | 9.21E-85 | 1.68959996 | 0.202 | 0.064 | 1.85E-80 |
| Adk | 1.13E-84 | 0.87103838 | 0.63 | 0.399 | 2.26E-80 |
| Fgf13 | 1.85E-84 | 2.90091242 | 0.146 | 0.037 | 3.70E-80 |
| Sos1 | 2.15E-84 | 1.86010852 | 0.181 | 0.053 | 4.30E-80 |
| Togaram1 | 2.67E-84 | 2.31763118 | 0.118 | 0.025 | 5.35E-80 |
| Zdhhc17 | 3.53E-84 | 1.98477353 | 0.152 | 0.04 | 7.08E-80 |

| | | | | | |
|---|---|---|---|---|---|
| Scaf8 | 4.90E-24 | 1.14025806 | 0.203 | 0.094 | 9.83E-20 |
| Gramd3 | 5.86E-24 | 1.51744172 | 0.123 | 0.045 | 1.18E-19 |
| Kdm6a | 6.60E-24 | 1.18705589 | 0.209 | 0.1 | 1.32E-19 |
| Lamc1 | 7.04E-24 | 0.8832828 | 0.342 | 0.199 | 1.41E-19 |
| Vps54 | 7.50E-24 | 1.29558454 | 0.155 | 0.063 | 1.51E-19 |
| Rhoj | 8.33E-24 | 0.94348453 | 0.319 | 0.179 | 1.67E-19 |
| Sdf2l1 | 8.67E-24 | 1.1664855 | 0.169 | 0.072 | 1.74E-19 |
| Eif4ebp2 | 9.24E-24 | 1.4921572 | 0.112 | 0.039 | 1.85E-19 |
| Slc23a2 | 1.11E-23 | 1.15460406 | 0.204 | 0.096 | 2.23E-19 |
| Ern1 | 1.39E-23 | 1.48065942 | 0.142 | 0.057 | 2.79E-19 |
| Pold4 | 1.58E-23 | 0.92312095 | 0.345 | 0.205 | 3.17E-19 |
| L3mbtl3 | 1.67E-23 | 1.29306297 | 0.17 | 0.074 | 3.34E-19 |
| Tjp1 | 1.70E-23 | 1.00130982 | 0.262 | 0.137 | 3.41E-19 |
| Cct2 | 1.74E-23 | 0.68676274 | 0.487 | 0.331 | 3.49E-19 |
| Sec63 | 1.92E-23 | 0.75770341 | 0.403 | 0.251 | 3.86E-19 |
| Lrch2 | 2.05E-23 | 1.51979145 | 0.108 | 0.037 | 4.12E-19 |
| Ptprf | 2.41E-23 | 1.01575843 | 0.246 | 0.125 | 4.84E-19 |
| Wdr75 | 2.43E-23 | 1.23261698 | 0.178 | 0.079 | 4.87E-19 |
| Aff1 | 2.61E-23 | 0.78400458 | 0.318 | 0.176 | 5.24E-19 |
| Xrn2 | 3.74E-23 | 0.85149107 | 0.327 | 0.188 | 7.51E-19 |
| Uri1 | 4.07E-23 | 1.0329654 | 0.229 | 0.114 | 8.17E-19 |
| Prrc1 | 4.76E-23 | 0.80865636 | 0.345 | 0.202 | 9.54E-19 |
| Washc4 | 4.98E-23 | 1.30518388 | 0.134 | 0.052 | 9.98E-19 |
| Akap10 | 5.42E-23 | 1.40816119 | 0.122 | 0.045 | 1.09E-18 |
| Ppp2r5e | 5.47E-23 | 1.0492786 | 0.217 | 0.105 | 1.10E-18 |
| Spats2 | 5.92E-23 | 1.03527239 | 0.202 | 0.095 | 1.19E-18 |
| Ankrd17 | 6.23E-23 | 0.73543568 | 0.414 | 0.261 | 1.25E-18 |
| Nsmce3 | 6.69E-23 | 1.49063201 | 0.101 | 0.034 | 1.34E-18 |
| Zfp532 | 1.07E-22 | 1.41835368 | 0.136 | 0.054 | 2.15E-18 |
| Dlg5 | 1.21E-22 | 1.33781247 | 0.157 | 0.067 | 2.42E-18 |
| Ube2m | 1.27E-22 | 0.76391601 | 0.381 | 0.234 | 2.55E-18 |
| 4930402H24 | 1.30E-22 | 1.13084437 | 0.191 | 0.089 | 2.61E-18 |
| Thsd4 | 1.33E-22 | 1.47481877 | 0.127 | 0.048 | 2.68E-18 |
| Bzw2 | 1.43E-22 | 1.07040888 | 0.207 | 0.099 | 2.88E-18 |
| Rlf | 1.60E-22 | 1.17052898 | 0.185 | 0.085 | 3.21E-18 |
| Map2k4 | 1.73E-22 | 1.0574938 | 0.21 | 0.102 | 3.47E-18 |
| Cdk13 | 1.75E-22 | 1.07426515 | 0.237 | 0.122 | 3.51E-18 |
| Ap1g1 | 1.78E-22 | 1.14643101 | 0.187 | 0.086 | 3.57E-18 |
| Bex3 | 1.89E-22 | 1.12052126 | 0.162 | 0.069 | 3.78E-18 |
| Btaf1 | 1.93E-22 | 1.25064651 | 0.168 | 0.074 | 3.88E-18 |
| Arhgap10 | 2.02E-22 | 1.23111593 | 0.173 | 0.077 | 4.05E-18 |
| Sod3 | 2.63E-22 | 0.65973292 | 0.556 | 0.402 | 5.28E-18 |
| Kitl | 2.67E-22 | 1.65704881 | 0.141 | 0.058 | 5.35E-18 |
| Ankrd44 | 2.85E-22 | 1.31546588 | 0.168 | 0.074 | 5.73E-18 |
| Tmod3 | 2.95E-22 | 0.83419066 | 0.342 | 0.202 | 5.93E-18 |
| Spata13 | 3.32E-22 | 1.31680938 | 0.146 | 0.06 | 6.66E-18 |
| Plau | 3.38E-22 | 0.61874279 | 0.139 | 0.055 | 6.79E-18 |
| Grk5 | 3.72E-22 | 1.14242764 | 0.169 | 0.075 | 7.46E-18 |
| Mgat5 | 4.26E-22 | 1.23585879 | 0.168 | 0.074 | 8.55E-18 |
| Epn2 | 4.50E-22 | 1.11115848 | 0.238 | 0.124 | 9.02E-18 |
| Trip11 | 4.83E-22 | 0.8112685 | 0.331 | 0.193 | 9.70E-18 |
| Supt16 | 5.77E-22 | 0.85574237 | 0.298 | 0.167 | 1.16E-17 |
| Elk3 | 5.87E-22 | 0.83741723 | 0.304 | 0.172 | 1.18E-17 |
| Atrnl1 | 6.36E-22 | 1.05785793 | 0.217 | 0.107 | 1.28E-17 |
| Stxbp5 | 6.57E-22 | 1.44346081 | 0.124 | 0.048 | 1.32E-17 |
| Xpo1 | 7.52E-22 | 1.04273885 | 0.22 | 0.11 | 1.51E-17 |
| Ralgapa1 | 7.54E-22 | 1.04560892 | 0.193 | 0.091 | 1.51E-17 |
| Fmo2 | 8.15E-22 | 1.09265625 | 0.107 | 0.038 | 1.64E-17 |
| Medag | 8.94E-22 | 0.7686608 | 0.339 | 0.203 | 1.79E-17 |
| Zc3h12b | 1.04E-21 | 1.47604042 | 0.113 | 0.041 | 2.09E-17 |
| Map4 | 1.04E-21 | 0.8028793 | 0.405 | 0.261 | 2.09E-17 |
| Kat6b | 1.16E-21 | 1.15360938 | 0.214 | 0.107 | 2.32E-17 |
| Khdrbs1 | 1.17E-21 | 0.71665477 | 0.377 | 0.231 | 2.34E-17 |
| Ptpa | 1.18E-21 | 1.312313 | 0.135 | 0.054 | 2.37E-17 |
| Tpx2 | 1.50E-21 | 1.65925927 | 0.114 | 0.043 | 3.02E-17 |
| Cwc27 | 1.57E-21 | 1.24388852 | 0.157 | 0.068 | 3.14E-17 |
| Grb10 | 1.57E-21 | 0.81265902 | 0.439 | 0.295 | 3.16E-17 |
| Fubp1 | 1.69E-21 | 0.61152408 | 0.423 | 0.266 | 3.38E-17 |
| Cdk6 | 1.90E-21 | 1.10844499 | 0.184 | 0.086 | 3.80E-17 |
| Eif2b3 | 1.91E-21 | 1.11790532 | 0.175 | 0.08 | 3.83E-17 |
| Ubr3 | 2.54E-21 | 0.96564462 | 0.226 | 0.115 | 5.10E-17 |
| Wdpcp | 2.63E-21 | 1.49657941 | 0.113 | 0.042 | 5.27E-17 |
| Sned1 | 2.76E-21 | 1.06332584 | 0.318 | 0.186 | 5.54E-17 |
| Picalm | 2.92E-21 | 0.75861112 | 0.461 | 0.312 | 5.86E-17 |
| Usp47 | 3.48E-21 | 0.87858142 | 0.263 | 0.142 | 6.99E-17 |
| Tgfbr1 | 3.66E-21 | 0.89745854 | 0.274 | 0.151 | 7.34E-17 |
| Lars | 3.74E-21 | 1.04511719 | 0.209 | 0.104 | 7.50E-17 |
| Wdr7 | 4.04E-21 | 1.45346666 | 0.114 | 0.043 | 8.10E-17 |
| Zfc3h1 | 4.08E-21 | 0.96010387 | 0.248 | 0.132 | 8.19E-17 |
| Lnpk | 4.27E-21 | 1.23546108 | 0.139 | 0.057 | 8.57E-17 |
| Tcf7l1 | 4.41E-21 | 1.25127087 | 0.213 | 0.109 | 8.85E-17 |
| Eprs | 4.41E-21 | 0.60294686 | 0.465 | 0.309 | 8.85E-17 |
| Mybbp1a | 5.10E-21 | 0.9621695 | 0.221 | 0.112 | 1.02E-16 |
| Ifnar2 | 5.12E-21 | 0.74095759 | 0.372 | 0.23 | 1.03E-16 |
| Dcaf5 | 5.24E-21 | 1.33015697 | 0.139 | 0.058 | 1.05E-16 |
| Psd3 | 5.25E-21 | 0.68659182 | 0.318 | 0.185 | 1.05E-16 |
| Kmt2e | 6.44E-21 | 0.69375876 | 0.541 | 0.398 | 1.29E-16 |
| Csnk1g3 | 7.10E-21 | 0.90913618 | 0.254 | 0.137 | 1.42E-16 |
| Nin | 7.41E-21 | 1.09826079 | 0.179 | 0.083 | 1.49E-16 |
| Pdha1 | 7.49E-21 | 1.03210202 | 0.416 | 0.28 | 1.50E-16 |
| Ackr3 | 7.89E-21 | 1.10608957 | 0.231 | 0.122 | 1.58E-16 |
| Btg3 | 1.11E-20 | 0.81367922 | 0.302 | 0.173 | 2.22E-16 |
| Sytl2 | 1.29E-20 | 1.40355802 | 0.14 | 0.059 | 2.59E-16 |
| Thsd7a | 1.31E-20 | 1.26758512 | 0.22 | 0.114 | 2.64E-16 |
| Hmbox1 | 1.44E-20 | 1.34805243 | 0.13 | 0.053 | 2.88E-16 |
| Med14 | 1.62E-20 | 1.17587291 | 0.164 | 0.074 | 3.26E-16 |
| Ufd1 | 1.62E-20 | 1.2936326 | 0.128 | 0.051 | 3.26E-16 |
| Eya3 | 1.78E-20 | 1.49054423 | 0.106 | 0.039 | 3.57E-16 |
| Fam98a | 2.58E-20 | 1.25992432 | 0.14 | 0.059 | 5.18E-16 |
| Rasl11a | 2.67E-20 | 0.8204204 | 0.252 | 0.137 | 5.37E-16 |
| Abl1 | 3.08E-20 | 1.0739173 | 0.287 | 0.168 | 6.17E-16 |
| Ranbp1 | 3.13E-20 | 0.63455701 | 0.527 | 0.383 | 6.28E-16 |
| Gm16685 | 3.27E-20 | 1.56025016 | 0.129 | 0.053 | 6.57E-16 |
| Copb1 | 3.52E-20 | 0.6455143 | 0.427 | 0.28 | 7.07E-16 |
| Acaca | 3.95E-20 | 1.1399637 | 0.18 | 0.086 | 7.93E-16 |
| C1qbp | 4.29E-20 | 0.67694537 | 0.359 | 0.22 | 8.61E-16 |
| Alyref | 4.66E-20 | 0.81037696 | 0.308 | 0.181 | 9.36E-16 |
| Tspan5 | 5.42E-20 | 1.25935262 | 0.167 | 0.077 | 1.09E-15 |
| Vps26 | 5.44E-20 | 1.17250389 | 0.156 | 0.07 | 1.09E-15 |
| Ubr5 | 6.33E-20 | 0.74313061 | 0.376 | 0.239 | 1.27E-15 |
| Hacd1 | 6.75E-20 | 0.83354551 | 0.302 | 0.176 | 1.35E-15 |
| Spc24 | 6.84E-20 | 1.13859771 | 0.141 | 0.06 | 1.37E-15 |
| Msi2 | 7.30E-20 | 0.90919911 | 0.257 | 0.141 | 1.47E-15 |
| Phka1 | 7.53E-20 | 1.38880184 | 0.106 | 0.039 | 1.51E-15 |
| Gnptab | 8.35E-20 | 1.4630408 | 0.106 | 0.04 | 1.68E-15 |
| Arhgap31 | 9.18E-20 | 0.92733316 | 0.319 | 0.192 | 1.84E-15 |
| Cep350 | 9.21E-20 | 0.99765254 | 0.196 | 0.097 | 1.85E-15 |
| Csnk1g1 | 9.99E-20 | 1.01052178 | 0.221 | 0.116 | 2.00E-15 |
| Ubn2 | 1.03E-19 | 0.77322959 | 0.376 | 0.241 | 2.08E-15 |
| Hsd17b12 | 1.17E-19 | 0.90329788 | 0.248 | 0.135 | 2.35E-15 |
| Pdlim5 | 1.22E-19 | 1.03921268 | 0.363 | 0.235 | 2.44E-15 |
| Setd5 | 1.23E-19 | 0.83763706 | 0.356 | 0.227 | 2.48E-15 |
| Tmem263 | 1.28E-19 | 0.67606776 | 0.37 | 0.231 | 2.57E-15 |
| Atp8b1 | 1.52E-19 | 1.33758335 | 0.167 | 0.078 | 3.06E-15 |
| Pde3a | 1.55E-19 | 1.06385754 | 0.156 | 0.07 | 3.10E-15 |
| Ddr2 | 1.86E-19 | 0.76058949 | 0.394 | 0.26 | 3.73E-15 |
| Smg7 | 1.98E-19 | 1.12183102 | 0.159 | 0.073 | 3.96E-15 |
| Cd2ap | 1.98E-19 | 0.8522758 | 0.263 | 0.147 | 3.98E-15 |
| Nipbl | 1.99E-19 | 0.7280218 | 0.408 | 0.269 | 3.99E-15 |
| Lyst | 2.04E-19 | 1.4685924 | 0.103 | 0.039 | 4.09E-15 |
| Il6st | 2.06E-19 | 0.78267586 | 0.456 | 0.323 | 4.14E-15 |
| Rhoq | 2.20E-19 | 0.78279768 | 0.325 | 0.197 | 4.41E-15 |
| mt-Nd6 | 2.20E-19 | 1.19574796 | 0.178 | 0.086 | 4.41E-15 |
| Vps13d | 2.20E-19 | 1.19482929 | 0.147 | 0.065 | 4.41E-15 |
| Cdk1 | 2.23E-19 | 1.15809441 | 0.114 | 0.045 | 4.47E-15 |
| Ptpn3 | 2.24E-19 | 1.44988708 | 0.101 | 0.037 | 4.50E-15 |
| Usp3 | 2.86E-19 | 0.88995337 | 0.241 | 0.131 | 5.75E-15 |
| H2afy | 3.65E-19 | 0.72509374 | 0.319 | 0.19 | 7.33E-15 |
| Rbbp6 | 4.36E-19 | 1.29113769 | 0.13 | 0.055 | 8.75E-15 |
| Abhd17c | 4.48E-19 | 0.87835052 | 0.191 | 0.094 | 8.99E-15 |
| Pias2 | 5.21E-19 | 1.0623921 | 0.201 | 0.103 | 1.05E-14 |
| Akap11 | 5.79E-19 | 0.9096838 | 0.212 | 0.109 | 1.16E-14 |
| Tmcc1 | 6.75E-19 | 0.98598043 | 0.246 | 0.137 | 1.36E-14 |
| Usp34 | 7.93E-19 | 0.66908496 | 0.427 | 0.289 | 1.59E-14 |
| Ptprg | 7.98E-19 | 0.83523372 | 0.275 | 0.157 | 1.60E-14 |
| Snx13 | 8.76E-19 | 1.22977565 | 0.142 | 0.063 | 1.76E-14 |
| Cyth1 | 1.08E-18 | 1.263878 | 0.139 | 0.061 | 2.16E-14 |
| Nbea | 1.08E-18 | 0.97716134 | 0.231 | 0.126 | 2.17E-14 |
| Thap12 | 1.15E-18 | 1.20327907 | 0.14 | 0.061 | 2.31E-14 |
| Hdgfl3 | 1.21E-18 | 1.24051162 | 0.125 | 0.052 | 2.44E-14 |
| Fam49b | 1.28E-18 | 1.20698886 | 0.164 | 0.078 | 2.56E-14 |
| Pcdh7 | 1.29E-18 | 1.14057347 | 0.271 | 0.162 | 2.60E-14 |
| Bckdhb | 1.30E-18 | 1.43629181 | 0.101 | 0.038 | 2.61E-14 |
| Hibadh | 1.49E-18 | 1.36831211 | 0.106 | 0.041 | 2.99E-14 |
| Nfia | 1.51E-18 | 0.68687189 | 0.373 | 0.242 | 3.04E-14 |
| Csnk2a2 | 1.52E-18 | 0.95567984 | 0.193 | 0.097 | 3.05E-14 |
| Zfp445 | 1.63E-18 | 1.3276492 | 0.112 | 0.044 | 3.27E-14 |
| Dipk2a | 1.68E-18 | 1.17673742 | 0.118 | 0.048 | 3.37E-14 |
| Pacs1 | 1.72E-18 | 1.26896207 | 0.134 | 0.058 | 3.46E-14 |
| Prrg1 | 1.76E-18 | 1.37468073 | 0.117 | 0.048 | 3.53E-14 |
| Strn3 | 2.56E-18 | 0.70162465 | 0.409 | 0.273 | 5.13E-14 |
| Cfdp1 | 2.62E-18 | 0.6256155 | 0.4 | 0.262 | 5.26E-14 |
| Ubtd2 | 2.71E-18 | 1.09359941 | 0.201 | 0.104 | 5.43E-14 |
| Dst | 2.74E-18 | 0.90742143 | 0.439 | 0.316 | 5.50E-14 |
| Psmd1 | 2.77E-18 | 0.663385 | 0.356 | 0.224 | 5.55E-14 |
| Trerf1 | 2.89E-18 | 1.19709234 | 0.112 | 0.044 | 5.79E-14 |
| Fhit | 3.05E-18 | 1.02653096 | 0.113 | 0.045 | 6.12E-14 |
| Mtmr3 | 4.21E-18 | 0.99229095 | 0.196 | 0.1 | 8.45E-14 |
| Golt1b | 4.48E-18 | 0.81953195 | 0.273 | 0.158 | 8.99E-14 |
| Gmps | 4.72E-18 | 0.89634418 | 0.217 | 0.115 | 9.47E-14 |
| Setdb1 | 5.40E-18 | 1.36609196 | 0.102 | 0.039 | 1.08E-13 |
| Scaf4 | 5.74E-18 | 1.21732061 | 0.127 | 0.054 | 1.15E-13 |
| Tspan11 | 5.93E-18 | 0.90344741 | 0.266 | 0.155 | 1.19E-13 |
| Ubap2 | 6.02E-18 | 0.99956087 | 0.196 | 0.101 | 1.21E-13 |
| Spop | 6.59E-18 | 0.62739161 | 0.437 | 0.3 | 1.32E-13 |
| Scaper | 7.70E-18 | 1.15107218 | 0.142 | 0.064 | 1.55E-13 |
| Gfpt1 | 8.04E-18 | 0.81259684 | 0.254 | 0.144 | 1.61E-13 |
| Lrp4 | 8.19E-18 | 1.2261088 | 0.193 | 0.101 | 1.64E-13 |
| Adamtsl1 | 1.02E-17 | 1.42498357 | 0.13 | 0.057 | 2.04E-13 |
| Pou2f1 | 1.11E-17 | 1.10759505 | 0.139 | 0.062 | 2.22E-13 |
| Foxc1 | 1.13E-17 | 0.91586154 | 0.168 | 0.081 | 2.27E-13 |
| Notch2 | 1.14E-17 | 0.61564956 | 0.349 | 0.218 | 2.28E-13 |
| Ppp4r3b | 1.14E-17 | 1.01853898 | 0.161 | 0.076 | 2.29E-13 |
| Jarid2 | 1.23E-17 | 1.04839702 | 0.182 | 0.092 | 2.47E-13 |
| Bmt2 | 1.40E-17 | 1.3365456 | 0.113 | 0.046 | 2.82E-13 |
| Nudcd2 | 1.42E-17 | 0.74444692 | 0.305 | 0.186 | 2.84E-13 |
| Lsm12 | 1.42E-17 | 0.73958439 | 0.319 | 0.197 | 2.85E-13 |
| Top2b | 1.43E-17 | 0.75845554 | 0.326 | 0.205 | 2.86E-13 |
| Ltbp2 | 1.47E-17 | 0.78921986 | 0.291 | 0.177 | 2.94E-13 |
| Nsmce2 | 1.52E-17 | 0.8004726 | 0.248 | 0.14 | 3.05E-13 |
| Btbd9 | 1.62E-17 | 1.06774864 | 0.163 | 0.079 | 3.25E-13 |
| Nectin3 | 1.83E-17 | 1.08991682 | 0.147 | 0.068 | 3.67E-13 |
| Rcc2 | 1.88E-17 | 0.8675565 | 0.219 | 0.118 | 3.76E-13 |
| Sae1 | 1.95E-17 | 0.92040407 | 0.214 | 0.115 | 3.92E-13 |
| Ermp1 | 1.96E-17 | 1.35328338 | 0.107 | 0.043 | 3.94E-13 |
| Cul3 | 2.11E-17 | 0.7076688 | 0.325 | 0.202 | 4.23E-13 |
| Nrf1 | 2.18E-17 | 1.06672659 | 0.152 | 0.071 | 4.38E-13 |
| Prkd1 | 2.42E-17 | 0.99497779 | 0.15 | 0.07 | 4.86E-13 |
| Ldb2 | 2.72E-17 | 1.41188765 | 0.122 | 0.053 | 5.46E-13 |
| Tmpo | 2.97E-17 | 0.92485044 | 0.2 | 0.106 | 5.95E-13 |
| Mrpl34 | 3.20E-17 | 0.81865528 | 0.249 | 0.142 | 6.43E-13 |
| Dcaf6 | 3.25E-17 | 1.02988057 | 0.165 | 0.081 | 6.51E-13 |
| Lrch3 | 3.95E-17 | 1.02949985 | 0.189 | 0.098 | 7.93E-13 |
| Ak2 | 4.03E-17 | 0.71828594 | 0.271 | 0.158 | 8.09E-13 |
| Tasp1 | 4.11E-17 | 1.21101472 | 0.137 | 0.063 | 8.25E-13 |
| 5031425E22 | 4.14E-17 | 0.9420335 | 0.257 | 0.151 | 8.31E-13 |
| Herc1 | 4.23E-17 | 0.83464473 | 0.255 | 0.147 | 8.48E-13 |
| Dpy19l1 | 4.62E-17 | 0.89814411 | 0.219 | 0.12 | 9.27E-13 |
| Cdc20 | 4.96E-17 | 1.21589282 | 0.107 | 0.043 | 9.95E-13 |
| Rrm1 | 4.98E-17 | 1.10798985 | 0.148 | 0.07 | 9.99E-13 |
| Kpna3 | 5.14E-17 | 0.94061693 | 0.181 | 0.092 | 1.03E-12 |
| Fam13b | 5.49E-17 | 1.1413195 | 0.145 | 0.067 | 1.10E-12 |
| Zmym4 | 5.52E-17 | 1.03586955 | 0.191 | 0.1 | 1.11E-12 |
| Mapkapk3 | 5.88E-17 | 1.37574349 | 0.106 | 0.043 | 1.18E-12 |
| Utrn | 6.20E-17 | 0.74236935 | 0.363 | 0.237 | 1.24E-12 |
| S1pr3 | 6.41E-17 | 0.90900123 | 0.144 | 0.066 | 1.29E-12 |
| Gas7 | 7.58E-17 | 1.15870585 | 0.133 | 0.06 | 1.52E-12 |
| Abce1 | 7.71E-17 | 0.78639117 | 0.243 | 0.138 | 1.55E-12 |
| Htt | 8.08E-17 | 1.0615787 | 0.153 | 0.073 | 1.62E-12 |
| Acvr2a | 8.76E-17 | 1.41524777 | 0.103 | 0.042 | 1.76E-12 |
| Prkca | 9.20E-17 | 0.96537852 | 0.231 | 0.131 | 1.85E-12 |
| Pak1 | 9.30E-17 | 1.09696598 | 0.175 | 0.089 | 1.87E-12 |
| Trappc9 | 9.66E-17 | 1.19780918 | 0.127 | 0.056 | 1.94E-12 |
| Anp32b | 1.16E-16 | 0.61434896 | 0.528 | 0.4 | 2.33E-12 |
| Snapc3 | 1.23E-16 | 1.09237022 | 0.15 | 0.071 | 2.47E-12 |
| Pex7 | 1.33E-16 | 1.20644199 | 0.122 | 0.053 | 2.67E-12 |
| Smad3 | 1.36E-16 | 0.97026176 | 0.206 | 0.112 | 2.73E-12 |
| Tcf7l2 | 1.46E-16 | 0.88307157 | 0.302 | 0.189 | 2.94E-12 |
| Srgn | 1.53E-16 | 1.55357232 | 0.124 | 0.056 | 3.07E-12 |
| Cnot2 | 1.57E-16 | 0.98040179 | 0.186 | 0.097 | 3.15E-12 |
| Mnat1 | 1.57E-16 | 0.92928422 | 0.237 | 0.136 | 3.15E-12 |
| Mindy2 | 1.59E-16 | 1.01068824 | 0.145 | 0.068 | 3.20E-12 |
| Pak2 | 1.68E-16 | 0.67116547 | 0.33 | 0.209 | 3.38E-12 |
| Brwd1 | 2.10E-16 | 0.81513516 | 0.254 | 0.148 | 4.22E-12 |
| Psmd12 | 2.14E-16 | 0.70851438 | 0.304 | 0.188 | 4.29E-12 |
| Nxn | 2.23E-16 | 1.00198603 | 0.225 | 0.128 | 4.48E-12 |
| Psip1 | 2.25E-16 | 0.74660461 | 0.286 | 0.173 | 4.51E-12 |
| Pum1 | 2.27E-16 | 0.69204093 | 0.343 | 0.221 | 4.55E-12 |
| Nav1 | 2.30E-16 | 0.71736739 | 0.336 | 0.216 | 4.62E-12 |
| Cd276 | 2.41E-16 | 0.9975689 | 0.182 | 0.095 | 4.84E-12 |
| Mindy3 | 2.69E-16 | 1.25793577 | 0.112 | 0.047 | 5.40E-12 |
| Sntb2 | 2.77E-16 | 0.6631807 | 0.319 | 0.199 | 5.56E-12 |
| Ctdspl2 | 3.20E-16 | 0.96730673 | 0.185 | 0.097 | 6.42E-12 |
| Hpse2 | 3.31E-16 | 1.05319356 | 0.18 | 0.094 | 6.65E-12 |
| Whrn | 3.44E-16 | 1.7438396 | 0.103 | 0.043 | 6.90E-12 |
| Washc2 | 3.67E-16 | 1.178697 | 0.119 | 0.052 | 7.36E-12 |
| Elp4 | 4.31E-16 | 1.06061551 | 0.124 | 0.055 | 8.65E-12 |
| Tmem87a | 4.91E-16 | 1.17562081 | 0.119 | 0.052 | 9.86E-12 |
| Abtb2 | 4.97E-16 | 1.41977946 | 0.101 | 0.041 | 9.98E-12 |
| Tlk1 | 5.03E-16 | 0.80793235 | 0.229 | 0.129 | 1.01E-11 |
| Plekha5 | 5.47E-16 | 0.93483922 | 0.236 | 0.137 | 1.10E-11 |
| Atf6 | 5.47E-16 | 0.98523171 | 0.185 | 0.097 | 1.10E-11 |
| Cep170 | 6.02E-16 | 0.92164838 | 0.201 | 0.109 | 1.21E-11 |
| Rnf217 | 6.07E-16 | 1.07894629 | 0.147 | 0.071 | 1.22E-11 |
| Slf1 | 6.86E-16 | 1.27463082 | 0.112 | 0.048 | 1.38E-11 |
| Pik3c2a | 7.15E-16 | 0.90390366 | 0.18 | 0.093 | 1.43E-11 |
| Atad2 | 7.17E-16 | 1.13605266 | 0.15 | 0.073 | 1.44E-11 |
| Ezh2 | 7.53E-16 | 0.93079587 | 0.203 | 0.111 | 1.51E-11 |
| Clmp | 7.86E-16 | 0.61104638 | 0.468 | 0.339 | 1.58E-11 |
| Phf20l1 | 8.18E-16 | 0.74112874 | 0.264 | 0.156 | 1.64E-11 |
| Kdm4c | 8.24E-16 | 1.17582529 | 0.12 | 0.053 | 1.65E-11 |
| Kif13b | 8.54E-16 | 1.14899299 | 0.135 | 0.063 | 1.71E-11 |
| Ptar1 | 9.18E-16 | 1.17998783 | 0.105 | 0.044 | 1.84E-11 |
| Vps13c | 9.38E-16 | 1.03920507 | 0.148 | 0.072 | 1.88E-11 |
| Sept9 | 9.50E-16 | 0.83756255 | 0.273 | 0.166 | 1.91E-11 |
| Atxn10 | 9.56E-16 | 0.66862125 | 0.356 | 0.236 | 1.92E-11 |
| Xpo7 | 1.00E-15 | 1.09910412 | 0.169 | 0.087 | 2.02E-11 |
| Ttc7 | 1.03E-15 | 0.98109467 | 0.164 | 0.083 | 2.06E-11 |
| Btrc | 1.06E-15 | 1.15163617 | 0.13 | 0.06 | 2.12E-11 |
| Gigyf2 | 1.10E-15 | 0.85402451 | 0.229 | 0.131 | 2.21E-11 |
| Rasa2 | 1.35E-15 | 0.81981386 | 0.247 | 0.146 | 2.70E-11 |
| Tanc2 | 1.48E-15 | 1.19060851 | 0.124 | 0.056 | 2.98E-11 |
| Tdg | 1.64E-15 | 1.04942558 | 0.147 | 0.072 | 3.30E-11 |
| Abl2 | 2.68E-15 | 0.8249407 | 0.186 | 0.099 | 5.39E-11 |
| Acbd3 | 2.98E-15 | 0.66525011 | 0.319 | 0.204 | 5.98E-11 |
| Agpat5 | 3.15E-15 | 0.82355646 | 0.191 | 0.103 | 6.33E-11 |
| Ndufaf8 | 3.61E-15 | 1.24049292 | 0.103 | 0.044 | 7.25E-11 |
| Mcm4 | 3.81E-15 | 1.30398771 | 0.105 | 0.045 | 7.64E-11 |
| Slc25a24 | 3.87E-15 | 1.00202896 | 0.161 | 0.082 | 7.77E-11 |
| Herc4 | 4.06E-15 | 1.07577984 | 0.147 | 0.073 | 8.15E-11 |

| | | | | | |
|---|---|---|---|---|---|
| Pid1 | 7.35E-69 | 1.50689152 | 0.309 | 0.148 | 1.48E-64 |
| Dhx36 | 8.10E-69 | 1.21698804 | 0.31 | 0.144 | 1.62E-64 |
| Cacna2d1 | 8.18E-69 | 1.21972603 | 0.369 | 0.193 | 1.64E-64 |
| Ppp4r3a | 1.02E-68 | 1.66791707 | 0.162 | 0.05 | 2.05E-64 |
| Tspo | 1.06E-68 | 0.63867874 | 0.739 | 0.584 | 2.14E-64 |
| Wac | 1.24E-68 | 1.04148877 | 0.398 | 0.209 | 2.49E-64 |
| Senp5 | 1.42E-68 | 1.60426463 | 0.181 | 0.061 | 2.85E-64 |
| Ptpra | 1.44E-68 | 1.07048357 | 0.361 | 0.18 | 2.88E-64 |
| Sall1 | 1.62E-68 | 1.81902256 | 0.147 | 0.043 | 3.25E-64 |
| Ubn2 | 1.99E-68 | 1.13111584 | 0.413 | 0.228 | 3.99E-64 |
| Tnks2 | 2.08E-68 | 1.18019267 | 0.324 | 0.154 | 4.18E-64 |
| Arid3a | 2.08E-68 | 1.90994386 | 0.132 | 0.036 | 4.18E-64 |
| Inhbb | 2.37E-68 | 1.77486402 | 0.227 | 0.091 | 4.75E-64 |
| Ppp1r12a | 2.37E-68 | 1.13238843 | 0.369 | 0.19 | 4.76E-64 |
| Lrrk1 | 3.46E-68 | 1.47505175 | 0.232 | 0.092 | 6.95E-64 |
| Abl1 | 3.50E-68 | 1.33321045 | 0.254 | 0.105 | 7.01E-64 |
| Spop | 3.59E-68 | 0.95029123 | 0.484 | 0.286 | 7.21E-64 |
| Tiam2 | 6.14E-68 | 1.43792995 | 0.226 | 0.088 | 1.23E-63 |
| Cenpx | 6.88E-68 | 1.71395815 | 0.152 | 0.046 | 1.38E-63 |
| Acer3 | 7.57E-68 | 1.58463617 | 0.204 | 0.075 | 1.52E-63 |
| Galnt7 | 1.08E-67 | 1.77905318 | 0.153 | 0.047 | 2.18E-63 |
| Nod1 | 1.46E-67 | 1.97222498 | 0.123 | 0.032 | 2.93E-63 |
| Camkk2 | 1.96E-67 | 1.0401286 | 0.368 | 0.187 | 3.92E-63 |
| Arid5b | 2.66E-67 | 0.75233002 | 0.558 | 0.346 | 5.35E-63 |
| March6 | 3.07E-67 | 1.43348305 | 0.224 | 0.088 | 6.15E-63 |
| Map4k5 | 3.17E-67 | 1.30746975 | 0.255 | 0.107 | 6.37E-63 |
| Dlg4 | 3.33E-67 | 1.5853587 | 0.184 | 0.064 | 6.69E-63 |
| Dram1 | 4.80E-67 | 1.4090096 | 0.226 | 0.089 | 9.63E-63 |
| Rarg | 9.04E-67 | 0.99889416 | 0.405 | 0.217 | 1.81E-62 |
| Lcorl | 1.02E-66 | 1.5758252 | 0.189 | 0.067 | 2.05E-62 |
| Ift172 | 2.10E-66 | 2.00183663 | 0.116 | 0.029 | 4.20E-62 |
| Rab43 | 2.66E-66 | 1.93317633 | 0.128 | 0.035 | 5.33E-62 |
| Rgl1 | 3.02E-66 | 1.3561201 | 0.262 | 0.113 | 6.06E-62 |
| Ttc21b | 3.57E-66 | 2.14867802 | 0.105 | 0.024 | 7.17E-62 |
| Dhx57 | 4.76E-66 | 2.14173195 | 0.106 | 0.025 | 9.55E-62 |
| Il11ra1 | 4.89E-66 | 0.71275367 | 0.576 | 0.358 | 9.80E-62 |
| Ift80 | 5.90E-66 | 1.90371191 | 0.13 | 0.036 | 1.18E-61 |
| Dpp8 | 9.40E-66 | 1.61865119 | 0.18 | 0.062 | 1.89E-61 |
| Plekhm3 | 1.01E-65 | 2.08761649 | 0.115 | 0.029 | 2.03E-61 |
| Kdm5c | 1.21E-65 | 1.52302242 | 0.211 | 0.081 | 2.44E-61 |
| mt-Nd6 | 1.24E-65 | 1.71906518 | 0.203 | 0.077 | 2.49E-61 |
| Hip1 | 1.29E-65 | 1.65359139 | 0.189 | 0.068 | 2.58E-61 |
| Hmbox1 | 1.68E-65 | 1.74165799 | 0.15 | 0.046 | 3.37E-61 |
| Phf20l1 | 2.03E-65 | 1.23158685 | 0.306 | 0.145 | 4.07E-61 |
| Mark1 | 2.45E-65 | 1.88044152 | 0.14 | 0.041 | 4.91E-61 |
| Cpt1c | 4.11E-65 | 1.88457058 | 0.135 | 0.039 | 8.25E-61 |
| Slc6a17 | 5.17E-65 | 1.17783997 | 0.266 | 0.114 | 1.04E-60 |
| Pcdh7 | 5.20E-65 | 1.62208726 | 0.305 | 0.152 | 1.04E-60 |
| Flrt2 | 5.54E-65 | 1.30956554 | 0.309 | 0.148 | 1.11E-60 |
| Mthfs | 5.56E-65 | 1.9454637 | 0.138 | 0.04 | 1.12E-60 |
| Snapc3 | 7.35E-65 | 1.66010006 | 0.18 | 0.063 | 1.48E-60 |
| Phf3 | 7.80E-65 | 1.08966092 | 0.35 | 0.178 | 1.57E-60 |
| Mocs2 | 9.00E-65 | 1.03769233 | 0.389 | 0.208 | 1.81E-60 |
| Slmap | 1.28E-64 | 1.29813202 | 0.281 | 0.129 | 2.57E-60 |
| Pde5a | 1.34E-64 | 1.22828727 | 0.31 | 0.149 | 2.69E-60 |
| Pgf | 1.35E-64 | 1.87787092 | 0.119 | 0.031 | 2.70E-60 |
| Ddhd2 | 1.44E-64 | 1.46803217 | 0.209 | 0.08 | 2.88E-60 |
| Agps | 3.21E-64 | 1.49720072 | 0.205 | 0.078 | 6.44E-60 |
| Taf1 | 3.49E-64 | 1.33235463 | 0.241 | 0.1 | 7.00E-60 |
| Trim23 | 3.82E-64 | 2.02294483 | 0.111 | 0.028 | 7.67E-60 |
| Baz2a | 1.07E-63 | 1.57150843 | 0.192 | 0.071 | 2.15E-59 |
| Lrba | 1.29E-63 | 2.04914235 | 0.114 | 0.029 | 2.58E-59 |
| Kdm2a | 1.49E-63 | 1.1269445 | 0.347 | 0.177 | 3.00E-59 |
| Trip4 | 1.55E-63 | 1.59096834 | 0.177 | 0.062 | 3.11E-59 |
| Bbx | 1.84E-63 | 1.25777946 | 0.307 | 0.149 | 3.70E-59 |
| Vps13a | 2.32E-63 | 1.8003218 | 0.141 | 0.042 | 4.65E-59 |
| Usp24 | 2.46E-63 | 1.6514753 | 0.16 | 0.052 | 4.94E-59 |
| Fbxw11 | 2.69E-63 | 1.46417985 | 0.217 | 0.086 | 5.41E-59 |
| Nav3 | 3.29E-63 | 2.17771821 | 0.154 | 0.05 | 6.61E-59 |
| Mpdz | 4.15E-63 | 1.48783111 | 0.201 | 0.076 | 8.33E-59 |
| Kitl | 5.05E-63 | 1.60692002 | 0.157 | 0.051 | 1.01E-58 |
| Stk38 | 6.06E-63 | 1.28558254 | 0.259 | 0.113 | 1.22E-58 |
| Phf12 | 6.10E-63 | 1.48877183 | 0.198 | 0.074 | 1.22E-58 |
| Auh | 6.81E-63 | 1.50272031 | 0.21 | 0.082 | 1.37E-58 |
| Nfatc3 | 7.25E-63 | 1.44368153 | 0.216 | 0.085 | 1.45E-58 |
| Ahi1 | 9.38E-63 | 1.49695488 | 0.203 | 0.078 | 1.88E-58 |
| Gmds | 9.44E-63 | 1.47050945 | 0.247 | 0.107 | 1.89E-58 |
| Tmem161b | 9.61E-63 | 2.02308005 | 0.111 | 0.028 | 1.93E-58 |
| Rprd1a | 9.86E-63 | 1.79675518 | 0.138 | 0.041 | 1.98E-58 |
| Hira | 1.02E-62 | 1.81863728 | 0.133 | 0.038 | 2.04E-58 |
| Epc2 | 1.26E-62 | 1.25925314 | 0.258 | 0.112 | 2.53E-58 |
| Arhgap44 | 1.32E-62 | 2.11054289 | 0.103 | 0.025 | 2.64E-58 |
| Lmbrd2 | 1.49E-62 | 1.95694865 | 0.114 | 0.03 | 2.99E-58 |
| Csnk2a1 | 1.61E-62 | 1.04703468 | 0.363 | 0.19 | 3.24E-58 |
| Hey2 | 1.66E-62 | 2.02308005 | 0.108 | 0.027 | 3.34E-58 |
| Frs2 | 1.66E-62 | 1.46139472 | 0.195 | 0.073 | 3.34E-58 |
| Fyb | 1.82E-62 | 2.10628625 | 0.108 | 0.027 | 3.65E-58 |
| Alg13 | 1.96E-62 | 2.03478705 | 0.112 | 0.029 | 3.93E-58 |
| Crispld2 | 2.27E-62 | 0.75870656 | 0.749 | 0.56 | 4.56E-58 |
| S100a1 | 2.85E-62 | 0.88813391 | 0.423 | 0.239 | 5.72E-58 |
| Dph6 | 4.04E-62 | 1.91379049 | 0.131 | 0.038 | 8.10E-58 |
| Flot1 | 4.06E-62 | 1.08839779 | 0.333 | 0.166 | 8.14E-58 |
| Phip | 4.80E-62 | 1.13407009 | 0.366 | 0.196 | 9.63E-58 |
| Tubgcp5 | 8.04E-62 | 1.85217252 | 0.132 | 0.039 | 1.61E-57 |
| Smap1 | 8.40E-62 | 0.85588304 | 0.471 | 0.279 | 1.69E-57 |
| Atxn2 | 8.84E-62 | 1.18844138 | 0.305 | 0.148 | 1.77E-57 |
| Pgpep1 | 9.01E-62 | 1.95739978 | 0.116 | 0.031 | 1.81E-57 |
| Stard13 | 1.30E-61 | 1.25637496 | 0.233 | 0.097 | 2.62E-57 |
| Gja1 | 1.43E-61 | 1.07779453 | 0.504 | 0.33 | 2.86E-57 |
| Rnd3 | 1.75E-61 | 0.90490554 | 0.479 | 0.292 | 3.50E-57 |
| Rars2 | 1.89E-61 | 2.03395594 | 0.108 | 0.027 | 3.80E-57 |
| Ifi27 | 2.07E-61 | 0.96205144 | 0.47 | 0.293 | 4.16E-57 |
| Rspry1 | 2.48E-61 | 1.29363796 | 0.238 | 0.101 | 4.98E-57 |
| Zfyve1 | 3.40E-61 | 1.99441916 | 0.112 | 0.029 | 6.83E-57 |
| Nsg1 | 3.70E-61 | 0.75457351 | 0.541 | 0.342 | 7.43E-57 |
| Elmsan1 | 6.18E-61 | 1.61527758 | 0.164 | 0.056 | 1.24E-56 |
| Ap1g1 | 6.76E-61 | 1.43785256 | 0.203 | 0.079 | 1.36E-56 |
| Ptpn4 | 7.86E-61 | 1.68382166 | 0.143 | 0.044 | 1.58E-56 |
| Tfcp2 | 9.25E-61 | 1.69121999 | 0.113 | 0.03 | 1.86E-56 |
| Bmpr1a | 9.49E-61 | 1.11716077 | 0.321 | 0.159 | 1.90E-56 |
| Igfbp2 | 1.07E-60 | 0.84513596 | 0.254 | 0.11 | 2.14E-56 |
| Arhgap35 | 1.15E-60 | 1.5139097 | 0.197 | 0.075 | 2.31E-56 |
| Fam49b | 1.26E-60 | 1.48706956 | 0.188 | 0.07 | 2.53E-56 |
| Zfp236 | 1.29E-60 | 1.84333281 | 0.129 | 0.038 | 2.59E-56 |
| Bmpr2 | 1.43E-60 | 0.92787698 | 0.449 | 0.265 | 2.87E-56 |
| Fryl | 1.51E-60 | 1.59890681 | 0.178 | 0.064 | 3.03E-56 |
| Ncoa3 | 1.55E-60 | 1.37643695 | 0.22 | 0.09 | 3.11E-56 |
| Ift88 | 1.74E-60 | 2.12274544 | 0.104 | 0.026 | 3.49E-56 |
| Cdkl3 | 1.97E-60 | 2.04548532 | 0.109 | 0.028 | 3.96E-56 |
| Itpr2 | 2.02E-60 | 1.77496738 | 0.14 | 0.043 | 4.06E-56 |
| Ggnbp2 | 2.28E-60 | 0.99933514 | 0.396 | 0.219 | 4.58E-56 |
| Gripap1 | 2.40E-60 | 1.5589776 | 0.173 | 0.061 | 4.81E-56 |
| Arhgef10l | 2.45E-60 | 1.71752085 | 0.149 | 0.048 | 4.91E-56 |
| Cradd | 2.84E-60 | 2.10662784 | 0.104 | 0.026 | 5.69E-56 |
| Ttc37 | 3.01E-60 | 1.84167298 | 0.13 | 0.038 | 6.05E-56 |
| Rnf217 | 3.48E-60 | 1.61414733 | 0.175 | 0.063 | 6.98E-56 |
| Fktn | 4.19E-60 | 1.98398967 | 0.108 | 0.028 | 8.41E-56 |
| Mindy2 | 4.39E-60 | 1.41532368 | 0.171 | 0.06 | 8.81E-56 |
| Epc1 | 4.78E-60 | 1.25138507 | 0.254 | 0.112 | 9.59E-56 |
| Pnisr | 5.70E-60 | 0.96587173 | 0.422 | 0.242 | 1.14E-55 |
| Ezh1 | 6.22E-60 | 1.77051109 | 0.133 | 0.04 | 1.25E-55 |
| Ints8 | 6.65E-60 | 1.66976398 | 0.149 | 0.048 | 1.33E-55 |
| Ep300 | 6.82E-60 | 1.19543179 | 0.268 | 0.122 | 1.37E-55 |
| Alg10b | 7.42E-60 | 1.81557013 | 0.12 | 0.034 | 1.49E-55 |
| Zmynd8 | 7.59E-60 | 1.30908857 | 0.232 | 0.097 | 1.52E-55 |
| Dcaf8 | 7.72E-60 | 1.13680504 | 0.307 | 0.15 | 1.55E-55 |
| Gopc | 9.74E-60 | 1.41993293 | 0.213 | 0.086 | 1.95E-55 |
| Cspp1 | 1.01E-59 | 1.26710524 | 0.272 | 0.126 | 2.03E-55 |
| Fkbp15 | 1.25E-59 | 1.75274934 | 0.135 | 0.041 | 2.51E-55 |
| Ctdspl2 | 1.87E-59 | 1.36210458 | 0.216 | 0.088 | 3.75E-55 |
| Gtf2ird1 | 1.93E-59 | 1.41531167 | 0.211 | 0.085 | 3.87E-55 |
| Nprl3 | 2.06E-59 | 1.98221134 | 0.106 | 0.027 | 4.13E-55 |
| Ckap5 | 2.11E-59 | 1.44134621 | 0.206 | 0.082 | 4.23E-55 |
| Sp3 | 2.72E-59 | 1.08631735 | 0.308 | 0.15 | 5.46E-55 |
| Bcl9 | 2.93E-59 | 1.64427187 | 0.153 | 0.051 | 5.88E-55 |
| Pbxip1 | 3.92E-59 | 0.96453307 | 0.393 | 0.218 | 7.87E-55 |
| Spopl | 4.33E-59 | 1.57426233 | 0.171 | 0.061 | 8.69E-55 |
| Klhl13 | 4.60E-59 | 2.00551541 | 0.118 | 0.033 | 9.24E-55 |
| Pum1 | 4.95E-59 | 0.98336444 | 0.382 | 0.209 | 9.94E-55 |
| Rab11fip3 | 4.98E-59 | 1.8484006 | 0.122 | 0.035 | 9.99E-55 |
| Slf1 | 5.24E-59 | 1.79198451 | 0.135 | 0.041 | 1.05E-54 |
| Dnm2 | 5.75E-59 | 1.20385144 | 0.275 | 0.128 | 1.15E-54 |
| Vezt | 7.68E-59 | 1.76687847 | 0.135 | 0.042 | 1.54E-54 |
| Tbl1xr1 | 7.80E-59 | 1.25238542 | 0.273 | 0.128 | 1.57E-54 |
| Map6 | 8.54E-59 | 1.69353112 | 0.144 | 0.046 | 1.71E-54 |
| Armc8 | 8.63E-59 | 1.64926948 | 0.151 | 0.05 | 1.73E-54 |
| Fam222b | 1.15E-58 | 1.72721425 | 0.138 | 0.043 | 2.31E-54 |
| Dlc1 | 1.19E-58 | 0.95429416 | 0.355 | 0.19 | 2.40E-54 |
| Nr3c1 | 1.58E-58 | 1.04059965 | 0.401 | 0.23 | 3.17E-54 |
| Kank2 | 1.83E-58 | 1.15081376 | 0.28 | 0.131 | 3.67E-54 |
| Arhgap5 | 2.04E-58 | 1.04578594 | 0.334 | 0.172 | 4.10E-54 |
| Nkd1 | 3.33E-58 | 1.08836653 | 0.269 | 0.123 | 6.68E-54 |
| Impact | 3.61E-58 | 1.1681173 | 0.267 | 0.122 | 7.24E-54 |
| Gnl2 | 5.56E-58 | 1.36175878 | 0.232 | 0.1 | 1.12E-53 |
| Sbspon | 5.68E-58 | 1.82093963 | 0.196 | 0.079 | 1.14E-53 |
| Mrpl58 | 7.41E-58 | 1.8225067 | 0.116 | 0.032 | 1.49E-53 |
| Kif13a | 7.41E-58 | 1.66776701 | 0.151 | 0.05 | 1.49E-53 |
| Bmp4 | 8.61E-58 | 0.76078736 | 0.379 | 0.2 | 1.73E-53 |
| Sik1 | 9.08E-58 | 1.56979003 | 0.172 | 0.062 | 1.82E-53 |
| Zfp871 | 1.04E-57 | 1.46137849 | 0.191 | 0.073 | 2.09E-53 |
| Brms1l | 1.24E-57 | 1.49174811 | 0.18 | 0.067 | 2.49E-53 |
| Celf1 | 1.28E-57 | 1.06934998 | 0.341 | 0.18 | 2.57E-53 |
| Cgnl1 | 1.65E-57 | 1.57846327 | 0.163 | 0.058 | 3.31E-53 |
| Ppp3cb | 2.20E-57 | 1.18191299 | 0.292 | 0.143 | 4.42E-53 |
| Yeats2 | 2.51E-57 | 1.68258219 | 0.153 | 0.052 | 5.04E-53 |
| Ddx42 | 2.97E-57 | 1.20562354 | 0.267 | 0.123 | 5.96E-53 |
| Hltf | 3.61E-57 | 1.62226323 | 0.153 | 0.052 | 7.24E-53 |
| Dip2b | 3.66E-57 | 1.60727458 | 0.162 | 0.057 | 7.34E-53 |
| Xpo4 | 4.18E-57 | 1.6600132 | 0.16 | 0.056 | 8.38E-53 |
| Ggps1 | 5.04E-57 | 1.43707949 | 0.188 | 0.072 | 1.01E-52 |
| Arfgef2 | 7.10E-57 | 1.64885861 | 0.147 | 0.049 | 1.42E-52 |
| Dpy19l4 | 7.91E-57 | 1.64251709 | 0.148 | 0.049 | 1.59E-52 |
| Zhx3 | 8.70E-57 | 1.5248062 | 0.178 | 0.066 | 1.75E-52 |
| Lrch1 | 1.12E-56 | 1.79266222 | 0.124 | 0.037 | 2.25E-52 |
| Tbc1d22a | 2.46E-56 | 1.99469456 | 0.101 | 0.026 | 4.93E-52 |
| Pikfyve | 2.46E-56 | 1.46708851 | 0.184 | 0.07 | 4.94E-52 |
| [illegible] | [illegible] | [illegible] | [illegible] | [illegible] | [illegible] |

[illegible]

| | | | | | |
|---|---|---|---|---|---|
| Setd2 | 5.86E-42 | 1.04914235 | 0.277 | 0.149 | 1.18E-37 |
| Ccdc47 | 5.89E-42 | 0.9936923 | 0.258 | 0.133 | 1.18E-37 |
| Zzz3 | 6.43E-42 | 1.03347775 | 0.253 | 0.13 | 1.29E-37 |
| Emp2 | 6.87E-42 | 0.92788227 | 0.391 | 0.247 | 1.38E-37 |
| Frk | 7.18E-42 | 1.63737256 | 0.118 | 0.041 | 1.44E-37 |
| Crabp1 | 1.29E-41 | 0.82508646 | 0.359 | 0.211 | 2.60E-37 |
| Ptprf | 1.44E-41 | 1.10572587 | 0.238 | 0.119 | 2.89E-37 |
| Synj1 | 1.47E-41 | 1.419366 | 0.14 | 0.054 | 2.94E-37 |
| Ids | 1.55E-41 | 1.02747141 | 0.243 | 0.122 | 3.11E-37 |
| Plppr3 | 1.58E-41 | 1.62829923 | 0.107 | 0.035 | 3.16E-37 |
| Snx27 | 1.63E-41 | 1.25964109 | 0.173 | 0.074 | 3.27E-37 |
| Hectd1 | 1.85E-41 | 0.74625385 | 0.401 | 0.248 | 3.72E-37 |
| Manea | 2.30E-41 | 1.50444372 | 0.126 | 0.046 | 4.61E-37 |
| Stat2 | 3.33E-41 | 1.53676785 | 0.114 | 0.039 | 6.69E-37 |
| Tmem30a | 3.52E-41 | 0.73387474 | 0.421 | 0.268 | 7.06E-37 |
| Supt20 | 3.65E-41 | 1.1944419 | 0.189 | 0.086 | 7.32E-37 |
| Tbc1d19 | 3.76E-41 | 1.61618294 | 0.111 | 0.037 | 7.55E-37 |
| Ttc3 | 3.79E-41 | 0.82975918 | 0.403 | 0.256 | 7.60E-37 |
| Eloc | 4.06E-41 | 0.84402792 | 0.257 | 0.13 | 8.15E-37 |
| Shoc2 | 5.63E-41 | 1.05423122 | 0.237 | 0.119 | 1.13E-36 |
| Gabpb2 | 5.64E-41 | 1.30762805 | 0.162 | 0.068 | 1.13E-36 |
| Ube2v1 | 8.20E-41 | 1.62365806 | 0.103 | 0.034 | 1.65E-36 |
| Scaf4 | 9.80E-41 | 1.46842307 | 0.131 | 0.049 | 1.97E-36 |
| Tnfrsf11b | 1.32E-40 | 0.84236212 | 0.175 | 0.075 | 2.65E-36 |
| Rnf111 | 1.49E-40 | 1.31166885 | 0.162 | 0.068 | 2.99E-36 |
| Acvrl1 | 1.81E-40 | 1.28540797 | 0.163 | 0.069 | 3.62E-36 |
| Rtn3 | 1.93E-40 | 0.84442143 | 0.377 | 0.233 | 3.88E-36 |
| Arhgef17 | 2.33E-40 | 1.25514581 | 0.174 | 0.076 | 4.67E-36 |
| Kmt2a | 2.35E-40 | 0.86610962 | 0.303 | 0.168 | 4.72E-36 |
| Slc30a9 | 2.41E-40 | 1.1574079 | 0.207 | 0.098 | 4.83E-36 |
| Sp1 | 2.72E-40 | 1.22145789 | 0.173 | 0.076 | 5.45E-36 |
| Agk | 2.92E-40 | 1.52163012 | 0.125 | 0.046 | 5.85E-36 |
| Uri1 | 3.64E-40 | 1.13604455 | 0.219 | 0.108 | 7.31E-36 |
| Wdr35 | 4.54E-40 | 1.54884224 | 0.112 | 0.039 | 9.11E-36 |
| Mfn1 | 5.22E-40 | 1.59256472 | 0.108 | 0.037 | 1.05E-35 |
| Slc38a6 | 5.95E-40 | 1.41241842 | 0.126 | 0.047 | 1.19E-35 |
| Tbc1d9b | 6.24E-40 | 1.33661535 | 0.149 | 0.061 | 1.25E-35 |
| Myof | 6.67E-40 | 0.8742637 | 0.357 | 0.218 | 1.34E-35 |
| Dcaf17 | 7.75E-40 | 1.42052462 | 0.127 | 0.047 | 1.56E-35 |
| Ggt7 | 7.83E-40 | 1.58609223 | 0.107 | 0.036 | 1.57E-35 |
| Mid1 | 8.02E-40 | 1.5834158 | 0.128 | 0.049 | 1.61E-35 |
| Arhgef7 | 8.54E-40 | 1.31112407 | 0.156 | 0.065 | 1.71E-35 |
| Cdk19 | 1.02E-39 | 1.38123377 | 0.149 | 0.061 | 2.04E-35 |
| St7l | 1.02E-39 | 1.31688888 | 0.157 | 0.066 | 2.05E-35 |
| Tmod3 | 1.45E-39 | 0.87821894 | 0.331 | 0.196 | 2.92E-35 |
| Ints6 | 1.47E-39 | 0.90042184 | 0.212 | 0.101 | 2.94E-35 |
| Slc6a6 | 1.59E-39 | 0.63099492 | 0.589 | 0.43 | 3.19E-35 |
| Itfg1 | 1.79E-39 | 1.01100722 | 0.249 | 0.13 | 3.60E-35 |
| Adamtsl5 | 2.33E-39 | 1.2894404 | 0.157 | 0.066 | 4.68E-35 |
| Atg16l1 | 2.91E-39 | 1.49569082 | 0.122 | 0.045 | 5.84E-35 |
| Filip1l | 3.51E-39 | 0.92093503 | 0.242 | 0.123 | 7.05E-35 |
| Tia1 | 3.65E-39 | 0.97147585 | 0.251 | 0.131 | 7.32E-35 |
| Parp3 | 4.01E-39 | 0.96709169 | 0.266 | 0.142 | 8.05E-35 |
| Cog4 | 4.16E-39 | 1.45751611 | 0.127 | 0.048 | 8.35E-35 |
| Taok3 | 4.87E-39 | 1.29096027 | 0.174 | 0.078 | 9.77E-35 |
| Klhl12 | 6.04E-39 | 1.58823784 | 0.104 | 0.035 | 1.21E-34 |
| Creb1 | 7.88E-39 | 1.01180298 | 0.23 | 0.116 | 1.58E-34 |
| Gng4 | 1.14E-38 | 1.60692002 | 0.101 | 0.033 | 2.28E-34 |
| Foxj3 | 1.30E-38 | 1.30633265 | 0.154 | 0.065 | 2.60E-34 |
| Rufy2 | 1.46E-38 | 1.58480955 | 0.108 | 0.037 | 2.92E-34 |
| Tbc1d12 | 1.55E-38 | 1.31520354 | 0.176 | 0.08 | 3.11E-34 |
| Kdm5b | 1.93E-38 | 0.93346039 | 0.284 | 0.158 | 3.87E-34 |
| Rmnd5a | 2.01E-38 | 1.08656589 | 0.21 | 0.102 | 4.03E-34 |
| Med15 | 2.06E-38 | 1.19684002 | 0.179 | 0.081 | 4.13E-34 |
| Odf2 | 2.11E-38 | 1.31120882 | 0.163 | 0.071 | 4.24E-34 |
| Ip6k1 | 2.56E-38 | 1.00484736 | 0.245 | 0.129 | 5.13E-34 |
| Gtf2a1 | 2.86E-38 | 1.29981955 | 0.152 | 0.064 | 5.74E-34 |
| Mfsd14a | 2.87E-38 | 1.49321952 | 0.108 | 0.038 | 5.75E-34 |
| Chd8 | 3.96E-38 | 1.10293883 | 0.203 | 0.098 | 7.96E-34 |
| Fam76b | 4.24E-38 | 1.25361286 | 0.161 | 0.07 | 8.50E-34 |
| 2610037D02 | 4.96E-38 | 1.3855394 | 0.134 | 0.053 | 9.95E-34 |
| Acp2 | 4.98E-38 | 1.24260228 | 0.15 | 0.062 | 9.99E-34 |
| Map3k7 | 4.98E-38 | 1.16091258 | 0.188 | 0.088 | 1.00E-33 |
| Zfp318 | 5.05E-38 | 1.27061026 | 0.151 | 0.063 | 1.01E-33 |
| Usp48 | 5.81E-38 | 1.29859378 | 0.163 | 0.071 | 1.16E-33 |
| Itpr1 | 5.98E-38 | 1.07687873 | 0.213 | 0.105 | 1.20E-33 |
| Gcc2 | 7.13E-38 | 1.10757564 | 0.201 | 0.097 | 1.43E-33 |
| Atp8b1 | 8.61E-38 | 1.2264288 | 0.166 | 0.073 | 1.73E-33 |
| Cul5 | 9.14E-38 | 0.89947491 | 0.283 | 0.157 | 1.83E-33 |
| Hp1bp3 | 1.36E-37 | 0.66585874 | 0.419 | 0.268 | 2.72E-33 |
| Sobp | 1.47E-37 | 1.24061688 | 0.182 | 0.085 | 2.94E-33 |
| Lamc3 | 1.81E-37 | 1.09670803 | 0.182 | 0.084 | 3.63E-33 |
| Tfap2a | 1.89E-37 | 1.54191115 | 0.111 | 0.04 | 3.79E-33 |
| Rab14 | 2.02E-37 | 0.64529558 | 0.485 | 0.337 | 4.06E-33 |
| Usp15 | 2.14E-37 | 1.13750994 | 0.183 | 0.084 | 4.29E-33 |
| Prpf6 | 2.35E-37 | 1.12720787 | 0.186 | 0.087 | 4.71E-33 |
| Smarca1 | 2.51E-37 | 1.46159668 | 0.111 | 0.04 | 5.03E-33 |
| Kifap3 | 3.38E-37 | 0.98493975 | 0.241 | 0.127 | 6.78E-33 |
| Thrap3 | 3.76E-37 | 0.8249826 | 0.337 | 0.202 | 7.54E-33 |
| Pdxdc1 | 3.86E-37 | 0.99283877 | 0.246 | 0.131 | 7.74E-33 |
| Rab3gap2 | 4.86E-37 | 1.32738162 | 0.143 | 0.059 | 9.76E-33 |
| Tfdp1 | 4.90E-37 | 1.51281462 | 0.106 | 0.037 | 9.83E-33 |
| Ciz1 | 4.99E-37 | 1.43294626 | 0.117 | 0.044 | 1.00E-32 |
| Med27 | 5.43E-37 | 1.42256076 | 0.121 | 0.046 | 1.09E-32 |
| Fblim1 | 6.81E-37 | 1.49408133 | 0.113 | 0.041 | 1.37E-32 |
| Ago1 | 7.83E-37 | 1.50592439 | 0.109 | 0.039 | 1.57E-32 |
| Pon2 | 1.08E-36 | 0.85946205 | 0.3 | 0.172 | 2.16E-32 |
| Vamp4 | 1.14E-36 | 1.10999755 | 0.196 | 0.095 | 2.28E-32 |
| Zkscan1 | 1.38E-36 | 1.32518246 | 0.136 | 0.055 | 2.77E-32 |
| Cept1 | 1.58E-36 | 1.24178742 | 0.17 | 0.077 | 3.16E-32 |
| Qrich1 | 1.64E-36 | 1.09507014 | 0.208 | 0.104 | 3.29E-32 |
| Rftl | 1.84E-36 | 1.55021968 | 0.106 | 0.038 | 3.69E-32 |
| Peak1 | 1.91E-36 | 0.85178933 | 0.299 | 0.172 | 3.83E-32 |
| Hnrnpr | 2.24E-36 | 0.79922384 | 0.352 | 0.216 | 4.49E-32 |
| Nol8 | 2.36E-36 | 1.29403677 | 0.143 | 0.059 | 4.73E-32 |
| Dnmt3a | 2.79E-36 | 1.03371231 | 0.211 | 0.105 | 5.60E-32 |
| Khsrp | 3.22E-36 | 1.17252476 | 0.175 | 0.08 | 6.46E-32 |
| Tet3 | 3.42E-36 | 1.22566776 | 0.153 | 0.066 | 6.87E-32 |
| Daam1 | 3.72E-36 | 1.23582587 | 0.156 | 0.068 | 7.46E-32 |
| Adam33 | 3.72E-36 | 1.03168314 | 0.171 | 0.077 | 7.47E-32 |
| Golph3l | 3.73E-36 | 1.59568463 | 0.101 | 0.035 | 7.49E-32 |
| Iqsec1 | 3.76E-36 | 1.07915492 | 0.191 | 0.092 | 7.54E-32 |
| Dnmt1 | 3.88E-36 | 1.28833001 | 0.127 | 0.049 | 7.78E-32 |
| Rufy1 | 3.89E-36 | 1.52795576 | 0.108 | 0.039 | 7.80E-32 |
| Strn | 4.35E-36 | 1.45464133 | 0.116 | 0.044 | 8.73E-32 |
| Esco1 | 4.51E-36 | 1.30359907 | 0.13 | 0.052 | 9.04E-32 |
| Gna12 | 4.57E-36 | 1.14196638 | 0.189 | 0.091 | 9.16E-32 |
| Usf3 | 5.42E-36 | 1.47867941 | 0.114 | 0.043 | 1.09E-31 |
| Mib1 | 8.90E-36 | 0.87420747 | 0.291 | 0.166 | 1.79E-31 |
| Zscan26 | 1.37E-35 | 1.19966195 | 0.158 | 0.07 | 2.75E-31 |
| Dpm1 | 1.42E-35 | 0.88134678 | 0.26 | 0.142 | 2.85E-31 |
| Pik3ca | 1.44E-35 | 0.93366513 | 0.252 | 0.137 | 2.90E-31 |
| Tnfrsf1a | 2.00E-35 | 0.67230591 | 0.444 | 0.302 | 4.01E-31 |
| Dgkh | 2.06E-35 | 1.37815987 | 0.125 | 0.05 | 4.13E-31 |
| Ap2a1 | 2.89E-35 | 1.0458372 | 0.194 | 0.094 | 5.79E-31 |
| Pip4k2a | 3.04E-35 | 1.43170624 | 0.116 | 0.044 | 6.09E-31 |
| Ndrg3 | 3.36E-35 | 1.05221082 | 0.203 | 0.101 | 6.75E-31 |
| Cd99l2 | 3.40E-35 | 1.37471822 | 0.119 | 0.046 | 6.83E-31 |
| Prr14 | 3.49E-35 | 1.17231627 | 0.162 | 0.073 | 7.01E-31 |
| Fyn | 4.05E-35 | 0.85761089 | 0.298 | 0.173 | 8.13E-31 |
| Bcorl1 | 4.40E-35 | 1.49594241 | 0.112 | 0.042 | 8.82E-31 |
| Prkaa1 | 4.98E-35 | 1.32529225 | 0.13 | 0.053 | 9.99E-31 |
| Optn | 5.44E-35 | 1.39967949 | 0.127 | 0.051 | 1.09E-30 |
| Slc17a5 | 6.17E-35 | 1.33864896 | 0.133 | 0.054 | 1.24E-30 |
| Trp53bp1 | 6.98E-35 | 1.38768171 | 0.119 | 0.046 | 1.40E-30 |
| Mthfr | 8.21E-35 | 1.40703594 | 0.114 | 0.043 | 1.65E-30 |
| Fam126a | 9.35E-35 | 1.05616277 | 0.272 | 0.157 | 1.88E-30 |
| Rc3h2 | 9.58E-35 | 0.96996126 | 0.256 | 0.142 | 1.92E-30 |
| Zhx1 | 9.60E-35 | 1.25204359 | 0.141 | 0.06 | 1.93E-30 |
| Dact3 | 9.76E-35 | 1.11965032 | 0.175 | 0.082 | 1.96E-30 |
| Rif1 | 1.27E-34 | 1.27278331 | 0.155 | 0.069 | 2.55E-30 |
| Rcor3 | 1.29E-34 | 1.10563543 | 0.182 | 0.087 | 2.59E-30 |
| Shwap | 1.31E-34 | 1.06702505 | 0.195 | 0.096 | 2.64E-30 |
| Abi2 | 1.40E-34 | 1.1516105 | 0.17 | 0.079 | 2.80E-30 |
| Gnptab | 1.41E-34 | 1.52522605 | 0.101 | 0.036 | 2.83E-30 |
| Mark2 | 1.92E-34 | 1.08179445 | 0.209 | 0.107 | 3.86E-30 |
| Ankfy1 | 2.83E-34 | 1.18903362 | 0.15 | 0.066 | 5.69E-30 |
| Cdc42bpb | 3.38E-34 | 1.20204967 | 0.154 | 0.069 | 6.78E-30 |
| Gosr1 | 3.44E-34 | 1.46676764 | 0.109 | 0.041 | 6.90E-30 |
| Klhl26 | 4.41E-34 | 1.42823896 | 0.11 | 0.041 | 8.85E-30 |
| Mxra7 | 5.26E-34 | 0.79148512 | 0.382 | 0.251 | 1.06E-29 |
| Ccdc91 | 5.49E-34 | 1.44045077 | 0.113 | 0.043 | 1.10E-29 |
| Zfp740 | 9.26E-34 | 1.24091807 | 0.14 | 0.06 | 1.86E-29 |
| Synrg | 9.55E-34 | 1.42928505 | 0.105 | 0.039 | 1.92E-29 |
| Ell | 1.12E-33 | 1.38159343 | 0.122 | 0.049 | 2.24E-29 |
| Ankle2 | 1.39E-33 | 1.25772897 | 0.132 | 0.055 | 2.79E-29 |
| Tbc1d8b | 1.44E-33 | 1.32239659 | 0.14 | 0.06 | 2.88E-29 |
| Ptbp3 | 1.51E-33 | 0.67870693 | 0.398 | 0.261 | 3.04E-29 |
| Txndc11 | 1.59E-33 | 1.05185673 | 0.216 | 0.114 | 3.20E-29 |
| Zc3h7a | 1.64E-33 | 0.95589294 | 0.253 | 0.141 | 3.29E-29 |
| Wwp1 | 1.66E-33 | 1.08070766 | 0.189 | 0.093 | 3.32E-29 |
| Rhobtb3 | 1.73E-33 | 1.11782828 | 0.183 | 0.089 | 3.47E-29 |
| Epn2 | 1.76E-33 | 0.98680957 | 0.224 | 0.119 | 3.53E-29 |
| Foxk1 | 1.78E-33 | 1.42765397 | 0.108 | 0.041 | 3.57E-29 |
| Sesn1 | 1.88E-33 | 1.55772061 | 0.115 | 0.045 | 3.76E-29 |
| Heatr5a | 1.90E-33 | 1.17334905 | 0.163 | 0.075 | 3.80E-29 |
| Qser1 | 2.17E-33 | 1.16667522 | 0.159 | 0.073 | 4.36E-29 |
| Hivep1 | 2.34E-33 | 1.18292416 | 0.152 | 0.068 | 4.69E-29 |
| Usp46 | 2.38E-33 | 1.44849822 | 0.106 | 0.04 | 4.77E-29 |
| Cstf3 | 2.44E-33 | 1.29460875 | 0.157 | 0.072 | 4.90E-29 |
| Cdk12 | 2.55E-33 | 0.74874741 | 0.315 | 0.189 | 5.11E-29 |
| Lix1l | 2.67E-33 | 0.69261198 | 0.391 | 0.255 | 5.35E-29 |
| Stx17 | 2.92E-33 | 1.33585575 | 0.125 | 0.051 | 5.86E-29 |
| Ccdc74a | 3.08E-33 | 1.20858586 | 0.149 | 0.066 | 6.19E-29 |
| Sorbs3 | 3.14E-33 | 1.44172499 | 0.102 | 0.037 | 6.30E-29 |
| Srpk2 | 4.15E-33 | 0.86092108 | 0.304 | 0.183 | 8.34E-29 |
| Ankrd13c | 4.32E-33 | 1.16378491 | 0.163 | 0.076 | 8.67E-29 |
| Anapc4 | 4.51E-33 | 1.10970118 | 0.173 | 0.083 | 9.05E-29 |
| Nim1k | 4.81E-33 | 1.29050496 | 0.108 | 0.04 | 9.66E-29 |
| Brd4 | 5.12E-33 | 0.69594595 | 0.372 | 0.238 | 1.03E-28 |
| Agl | 5.98E-33 | 1.17199909 | 0.112 | 0.043 | 1.20E-28 |
| Dennd5a | 6.34E-33 | 0.86652227 | 0.282 | 0.164 | 1.27E-28 |
| Akt2 | 6.54E-33 | 0.93906795 | 0.223 | 0.118 | 1.31E-28 |
| Wasf2 | 8.31E-33 | 0.74063814 | 0.393 | 0.261 | 1.67E-28 |
| Ldlr | 8.41E-33 | 1.15020927 | 0.159 | 0.073 | 1.69E-28 |
| Palld | 8.93E-33 | 0.80956431 | 0.336 | 0.21 | 1.79E-28 |
| Tbc1d23 | 9.14E-33 | 1.15168353 | 0.153 | 0.069 | 1.83E-28 |
| Epb41 | 9.48E-33 | 1.06775802 | 0.177 | 0.085 | 1.90E-28 |
| Npat | 1.01E-32 | 1.38823099 | 0.11 | 0.042 | 2.02E-28 |
| Dicer1 | 1.09E-32 | 1.12964278 | 0.161 | 0.075 | 2.20E-28 |
| E2f3 | 1.15E-32 | 1.50857396 | 0.103 | 0.038 | 2.31E-28 |
| Sptlc1 | 1.23E-32 | 1.35333408 | 0.114 | 0.045 | 2.48E-28 |
| Fads1 | 1.44E-32 | 1.0711016 | 0.185 | 0.091 | 2.89E-28 |
| Ext2 | 1.53E-32 | 0.85171279 | 0.28 | 0.163 | 3.06E-28 |
| Rp2 | 1.72E-32 | 1.30735283 | 0.122 | 0.05 | 3.46E-28 |
| Sltm | 1.75E-32 | 0.82140858 | 0.334 | 0.21 | 3.51E-28 |
| Ltn1 | 1.82E-32 | 1.26364686 | 0.133 | 0.057 | 3.66E-28 |
| Zcchc14 | 1.88E-32 | 0.79430566 | 0.307 | 0.185 | 3.78E-28 |
| Senp1 | 2.18E-32 | 1.43492604 | 0.105 | 0.04 | 4.37E-28 |
| Arid1a | 2.31E-32 | 0.81366242 | 0.285 | 0.167 | 4.63E-28 |
| Memo1 | 2.32E-32 | 1.10945377 | 0.208 | 0.109 | 4.65E-28 |
| Supt6 | 2.36E-32 | 1.00176758 | 0.195 | 0.099 | 4.74E-28 |
| Mef2a | 2.42E-32 | 0.60260924 | 0.488 | 0.348 | 4.86E-28 |
| Gm26782 | 2.45E-32 | 1.3891169 | 0.105 | 0.04 | 4.92E-28 |
| Jak2 | 2.65E-32 | 0.88879885 | 0.244 | 0.135 | 5.33E-28 |
| Pogz | 2.68E-32 | 1.3608506 | 0.12 | 0.049 | 5.38E-28 |
| Map3k20 | 2.82E-32 | 1.31150589 | 0.117 | 0.046 | 5.65E-28 |
| Zadh2 | 3.09E-32 | 1.25559322 | 0.138 | 0.06 | 6.20E-28 |
| Evi5 | 3.16E-32 | 0.97808456 | 0.224 | 0.12 | 6.35E-28 |
| Cul3 | 4.37E-32 | 0.77852634 | 0.318 | 0.195 | 8.77E-28 |
| Ttc5 | 4.56E-32 | 1.41961429 | 0.103 | 0.038 | 9.15E-28 |
| Gigyf1 | 5.03E-32 | 1.12576363 | 0.153 | 0.07 | 1.01E-27 |
| Hnrnpd | 5.40E-32 | 0.72817959 | 0.318 | 0.194 | 1.08E-27 |
| Arhgap21 | 5.91E-32 | 1.06986951 | 0.183 | 0.09 | 1.19E-27 |
| Stk39 | 8.28E-32 | 1.29425484 | 0.132 | 0.057 | 1.66E-27 |
| C330007P06 | 9.00E-32 | 1.20114544 | 0.151 | 0.069 | 1.81E-27 |
| Tor1aip1 | 9.46E-32 | 0.61274626 | 0.407 | 0.269 | 1.90E-27 |
| Klhl22 | 1.02E-31 | 1.18879434 | 0.146 | 0.066 | 2.04E-27 |
| Nek1 | 1.04E-31 | 1.27052905 | 0.134 | 0.058 | 2.08E-27 |
| Zfp652 | 1.08E-31 | 0.88741916 | 0.222 | 0.119 | 2.17E-27 |
| Ephb3 | 1.20E-31 | 1.21801926 | 0.128 | 0.054 | 2.41E-27 |
| Slc33a1 | 1.36E-31 | 1.34541005 | 0.117 | 0.048 | 2.73E-27 |
| Esyt2 | 1.61E-31 | 0.7937726 | 0.337 | 0.214 | 3.23E-27 |
| Clint1 | 1.73E-31 | 0.69383367 | 0.355 | 0.226 | 3.46E-27 |
| Flii | 1.97E-31 | 0.98299284 | 0.203 | 0.106 | 3.96E-27 |
| Mtus1 | 2.11E-31 | 1.45112369 | 0.116 | 0.047 | 4.24E-27 |
| Phactr2 | 2.13E-31 | 1.01248388 | 0.192 | 0.097 | 4.27E-27 |
| Nsf | 2.18E-31 | 1.13880968 | 0.168 | 0.081 | 4.38E-27 |
| Cnot1 | 2.21E-31 | 1.00442443 | 0.218 | 0.118 | 4.43E-27 |
| Usp37 | 2.50E-31 | 1.20754321 | 0.13 | 0.056 | 5.01E-27 |
| March3 | 3.05E-31 | 1.28526175 | 0.124 | 0.052 | 6.12E-27 |
| Cul2 | 3.17E-31 | 1.33322845 | 0.122 | 0.051 | 6.37E-27 |
| Psen2 | 3.25E-31 | 1.15715457 | 0.148 | 0.067 | 6.51E-27 |
| Dcun1d3 | 3.79E-31 | 1.30345953 | 0.116 | 0.047 | 7.61E-27 |
| Arhgef40 | 3.80E-31 | 0.80745217 | 0.275 | 0.16 | 7.63E-27 |
| Kif21a | 4.05E-31 | 1.1815699 | 0.158 | 0.074 | 8.12E-27 |
| Eea1 | 4.32E-31 | 0.70664002 | 0.365 | 0.238 | 8.67E-27 |
| Kdm3a | 4.37E-31 | 1.15368665 | 0.163 | 0.078 | 8.77E-27 |
| Rbl2 | 4.60E-31 | 1.1627049 | 0.146 | 0.066 | 9.24E-27 |
| Kif1bp | 5.37E-31 | 1.2754182 | 0.127 | 0.054 | 1.08E-26 |
| Plcb3 | 5.59E-31 | 1.38497258 | 0.11 | 0.044 | 1.12E-26 |
| Naa16 | 6.18E-31 | 1.36555181 | 0.119 | 0.049 | 1.24E-26 |
| Frmd6 | 6.27E-31 | 0.63099894 | 0.451 | 0.318 | 1.26E-26 |
| Nktr | 7.33E-31 | 0.62318657 | 0.47 | 0.337 | 1.47E-26 |
| Nek7 | 7.34E-31 | 0.9148061 | 0.248 | 0.141 | 1.47E-26 |
| Paxbp1 | 7.67E-31 | 1.19712808 | 0.145 | 0.066 | 1.54E-26 |
| Cerk | 1.02E-30 | 1.27763833 | 0.129 | 0.056 | 2.05E-26 |
| Mtss1 | 1.07E-30 | 1.25799286 | 0.132 | 0.057 | 2.14E-26 |
| Cpne3 | 1.10E-30 | 0.9195431 | 0.221 | 0.119 | 2.20E-26 |
| Rpe | 1.16E-30 | 1.30318797 | 0.115 | 0.047 | 2.32E-26 |
| Tbcel | 1.28E-30 | 1.25288901 | 0.123 | 0.052 | 2.57E-26 |
| Pcgf5 | 1.35E-30 | 1.11177333 | 0.166 | 0.08 | 2.72E-26 |
| Aldh3a2 | 1.40E-30 | 1.22030306 | 0.128 | 0.055 | 2.80E-26 |
| Extl3 | 1.44E-30 | 0.96382053 | 0.203 | 0.106 | 2.89E-26 |
| Kansl1l | 1.49E-30 | 1.07374069 | 0.224 | 0.125 | 2.99E-26 |
| Fndc4 | 1.55E-30 | 1.03380066 | 0.175 | 0.086 | 3.12E-26 |
| Utp11 | 2.07E-30 | 1.18304363 | 0.129 | 0.056 | 4.15E-26 |
| Mia3 | 2.08E-30 | 0.74885273 | 0.329 | 0.208 | 4.17E-26 |
| Fam110b | 2.53E-30 | 1.28903467 | 0.128 | 0.055 | 5.08E-26 |
| Sin3a | 2.53E-30 | 1.22186288 | 0.132 | 0.058 | 5.08E-26 |
| Gatad2b | 2.60E-30 | 0.73330158 | 0.338 | 0.215 | 5.23E-26 |
| Traf3 | 3.47E-30 | 1.1766924 | 0.144 | 0.066 | 6.97E-26 |
| Adam10 | 3.63E-30 | 0.94187102 | 0.229 | 0.128 | 7.28E-26 |
| Abl2 | 3.91E-30 | 1.01887856 | 0.184 | 0.094 | 7.84E-26 |
| Amn1 | 4.03E-30 | 1.25863554 | 0.15 | 0.07 | 8.09E-26 |
| Tmem63a | 4.06E-30 | 1.05790495 | 0.167 | 0.081 | 8.15E-26 |
| St3gal2 | 5.24E-30 | 1.0612891 | 0.183 | 0.093 | 1.05E-25 |
| Ppp2r5b | 7.54E-30 | 1.27850904 | 0.124 | 0.053 | 1.51E-25 |
| Eif2ak2 | 7.68E-30 | 1.34342769 | 0.118 | 0.049 | 1.54E-25 |
| Lyn | 7.98E-30 | 1.22950909 | 0.124 | 0.053 | 1.60E-25 |
| Nudcd3 | 9.11E-30 | 0.81103071 | 0.272 | 0.161 | 1.83E-25 |
| Ctnna1 | 1.14E-29 | 0.6618735 | 0.397 | 0.269 | 2.30E-25 |
| Cxxc5 | 1.41E-29 | 0.81144091 | 0.256 | 0.148 | 2.83E-25 |
| Nek9 | 1.63E-29 | 1.06293615 | 0.167 | 0.082 | 3.27E-25 |
| Rhou | 1.85E-29 | 1.04370846 | 0.152 | 0.072 | 3.71E-25 |
| Prkci | 1.88E-29 | 1.18984121 | 0.133 | 0.059 | 3.77E-25 |
| Hgsnat | 1.89E-29 | 0.77206849 | 0.275 | 0.163 | 3.79E-25 |
| Lats2 | 2.05E-29 | 0.8209964 | 0.242 | 0.137 | 4.11E-25 |
| Csnk2a2 | 2.18E-29 | 1.00312995 | 0.183 | 0.093 | 4.38E-25 |
| Mbtps2 | 2.62E-29 | 1.4236093 | 0.105 | 0.042 | 5.25E-25 |
| Map1a | 2.75E-29 | 1.09721374 | 0.154 | 0.073 | 5.51E-25 |
| Mrc2 | 2.85E-29 | 0.60051337 | 0.365 | 0.236 | 5.72E-25 |
| Arap1 | 2.99E-29 | 1.05291739 | 0.163 | 0.08 | 6.00E-25 |
| Tmem47 | 3.05E-29 | 0.87801859 | 0.224 | 0.124 | 6.11E-25 |
| Atp2c1 | 3.30E-29 | 0.94489468 | 0.207 | 0.111 | 6.62E-25 |
| Cnot10 | 3.44E-29 | 1.21906735 | 0.123 | 0.053 | 6.91E-25 |
| Tet2 | 3.96E-29 | 1.03359352 | 0.177 | 0.09 | 7.96E-25 |
| Taf2 | 4.42E-29 | 1.21636897 | 0.128 | 0.056 | 8.86E-25 |
| Ctcf | 4.56E-29 | 0.85561161 | 0.25 | 0.144 | 9.15E-25 |
| Zcchc24 | 5.73E-29 | 0.71725414 | 0.338 | 0.218 | 1.15E-24 |
| Aifm1 | 5.73E-29 | 1.27535181 | 0.12 | 0.051 | 1.15E-24 |
| Dixdc1 | 5.94E-29 | 1.19963397 | 0.123 | 0.053 | 1.19E-24 |
| Sptlc2 | 6.24E-29 | 1.02924279 | 0.202 | 0.109 | 1.25E-24 |
| Gnl3l | 6.48E-29 | 1.15880959 | 0.132 | 0.059 | 1.30E-24 |
| Ppfia1 | 6.62E-29 | 1.00261721 | 0.186 | 0.097 | 1.33E-24 |
| Paip1 | 7.23E-29 | 0.93336359 | 0.203 | 0.109 | 1.45E-24 |
| Scarb2 | 7.24E-29 | 0.73894354 | 0.277 | 0.165 | 1.45E-24 |
| Lgals3bp | 7.27E-29 | 0.90164192 | 0.201 | 0.107 | 1.46E-24 |
| Smad4 | 8.45E-29 | 0.84514603 | 0.251 | 0.146 | 1.70E-24 |
| Rapgef6 | 1.04E-28 | 0.97397178 | 0.216 | 0.12 | 2.09E-24 |
| Xpo1 | 1.08E-28 | 0.99621996 | 0.199 | 0.106 | 2.16E-24 |
| Mier1 | 1.35E-28 | 0.63500681 | 0.365 | 0.24 | 2.72E-24 |
| Stk24 | 1.58E-28 | 0.87998453 | 0.227 | 0.128 | 3.17E-24 |
| Uhrf2 | 1.59E-28 | 0.68906143 | 0.333 | 0.213 | 3.19E-24 |
| Nfx1 | 1.85E-28 | 0.99156507 | 0.178 | 0.091 | 3.71E-24 |
| Ncstn | 1.95E-28 | 0.89433781 | 0.215 | 0.118 | 3.92E-24 |
| Smarcad1 | 1.99E-28 | 1.10364288 | 0.152 | 0.073 | 4.00E-24 |
| Phactr4 | 2.06E-28 | 1.17795581 | 0.137 | 0.063 | 4.13E-24 |
| Sept10 | 2.32E-28 | 1.08431698 | 0.167 | 0.084 | 4.66E-24 |
| Ahcyl1 | 2.34E-28 | 0.79799093 | 0.254 | 0.149 | 4.70E-24 |
| Rfc1 | 2.72E-28 | 0.91441235 | 0.203 | 0.11 | 5.45E-24 |
| Zbtb7c | 2.77E-28 | 1.35710262 | 0.112 | 0.047 | 5.56E-24 |
| Sestd1 | 2.79E-28 | 0.99603101 | 0.187 | 0.098 | 5.61E-24 |
| Ccar1 | 3.12E-28 | 0.68597336 | 0.348 | 0.228 | 6.26E-24 |
| Acin1 | 3.18E-28 | 0.69884293 | 0.364 | 0.243 | 6.39E-24 |
| Mark3 | 3.57E-28 | 0.86662622 | 0.252 | 0.148 | 7.17E-24 |
| B3glct | 3.91E-28 | 0.81641724 | 0.26 | 0.154 | 7.84E-24 |
| Zfp592 | 4.27E-28 | 1.22714065 | 0.116 | 0.049 | 8.56E-24 |
| Sh3bp5 | 4.56E-28 | 0.92188016 | 0.194 | 0.103 | 9.15E-24 |
| Syap1 | 4.64E-28 | 0.94378935 | 0.205 | 0.111 | 9.31E-24 |
| Mau2 | 4.85E-28 | 1.16750454 | 0.13 | 0.058 | 9.73E-24 |
| Zfp277 | 7.07E-28 | 1.23585447 | 0.121 | 0.053 | 1.42E-23 |
| Plekhm2 | 8.06E-28 | 1.02197214 | 0.156 | 0.076 | 1.62E-23 |
| Nfyc | 1.09E-27 | 1.09044932 | 0.169 | 0.086 | 2.18E-23 |
| Atl3 | 1.21E-27 | 1.30176014 | 0.116 | 0.05 | 2.43E-23 |
| Snx15 | 1.26E-27 | 1.0549247 | 0.149 | 0.072 | 2.52E-23 |
| Slc39a14 | 1.27E-27 | 0.69402677 | 0.367 | 0.246 | 2.55E-23 |
| Adam22 | 1.32E-27 | 1.38838449 | 0.12 | 0.053 | 2.65E-23 |
| Slc9a1 | 1.38E-27 | 1.09512855 | 0.148 | 0.071 | 2.76E-23 |
| Vps53 | 1.54E-27 | 1.24391373 | 0.112 | 0.047 | 3.09E-23 |
| Tgoln1 | 1.60E-27 | 0.61395827 | 0.366 | 0.244 | 3.21E-23 |
| Kpna4 | 1.67E-27 | 0.79330344 | 0.277 | 0.169 | 3.35E-23 |
| Cpxm2 | 1.72E-27 | 1.37989429 | 0.103 | 0.042 | 3.46E-23 |
| Pank2 | 1.75E-27 | 1.01628326 | 0.17 | 0.086 | 3.50E-23 |
| Cuedc1 | 1.92E-27 | 0.87961915 | 0.223 | 0.126 | 3.86E-23 |
| Ccnh | 2.20E-27 | 1.05417794 | 0.163 | 0.082 | 4.42E-23 |
| Gnai3 | 2.34E-27 | 0.66186051 | 0.35 | 0.231 | 4.69E-23 |
| Slc44a1 | 2.79E-27 | 0.66920032 | 0.403 | 0.282 | 5.60E-23 |
| Sash1 | 2.79E-27 | 0.6941224 | 0.298 | 0.186 | 5.61E-23 |
| Arl5a | 3.25E-27 | 0.91953649 | 0.21 | 0.116 | 6.52E-23 |
| Tpp2 | 3.30E-27 | 0.80628582 | 0.253 | 0.15 | 6.61E-23 |
| Mtor | 3.71E-27 | 1.306115 | 0.109 | 0.046 | 7.44E-23 |
| Kdm5a | 3.80E-27 | 0.80135892 | 0.278 | 0.171 | 7.63E-23 |
| Polr2a | 4.05E-27 | 0.62450285 | 0.317 | 0.199 | 8.13E-23 |
| Pnpla8 | 4.34E-27 | 0.7985295 | 0.251 | 0.149 | 8.72E-23 |
| Foxk2 | 4.71E-27 | 1.1180897 | 0.134 | 0.062 | 9.46E-23 |
| Cenpc1 | 5.09E-27 | 1.06508389 | 0.154 | 0.076 | 1.02E-22 |
| Birc3 | 5.21E-27 | 0.73115425 | 0.227 | 0.127 | 1.05E-22 |
| Chst11 | 5.44E-27 | 0.84421999 | 0.187 | 0.099 | 1.09E-22 |
| Gdi1 | 6.50E-27 | 1.02838379 | 0.161 | 0.081 | 1.30E-22 |
| Impad1 | 7.52E-27 | 0.71025542 | 0.313 | 0.202 | 1.51E-22 |
| Hdac1 | 8.68E-27 | 0.74145617 | 0.284 | 0.176 | 1.74E-22 |
| Zfyve16 | 8.71E-27 | 1.16541773 | 0.13 | 0.06 | 1.75E-22 |
| Arhgap29 | 9.55E-27 | 0.97159323 | 0.18 | 0.095 | 1.92E-22 |
| Pla2g4a | 9.97E-27 | 1.06233627 | 0.168 | 0.087 | 2.00E-22 |
| Aqr | 1.00E-26 | 1.10287751 | 0.145 | 0.07 | 2.01E-22 |
| Klhl7 | 1.48E-26 | 1.2545417 | 0.119 | 0.053 | 2.96E-22 |
| Crebzf | 1.57E-26 | 1.14998304 | 0.135 | 0.064 | 3.14E-22 |
| Gm26532 | 1.58E-26 | 0.96468796 | 0.226 | 0.13 | 3.17E-22 |
| Bod1l | 1.88E-26 | 0.70789798 | 0.297 | 0.187 | 3.78E-22 |
| Pcnt | 2.13E-26 | 1.27153477 | 0.101 | 0.042 | 4.28E-22 |
| Msx2 | 2.31E-26 | 0.85267327 | 0.19 | 0.102 | 4.64E-22 |
| Lrp5 | 2.52E-26 | 1.13828285 | 0.138 | 0.065 | 5.06E-22 |
| Wdr26 | 2.82E-26 | 0.64111256 | 0.382 | 0.263 | 5.65E-22 |
| Ufl1 | 3.04E-26 | 0.9306956 | 0.192 | 0.104 | 6.10E-22 |
| Ptgr1 | 3.47E-26 | 1.2488307 | 0.104 | 0.044 | 6.97E-22 |
| Dhx40 | 3.73E-26 | 0.84336464 | 0.215 | 0.121 | 7.48E-22 |
| Gpr153 | 3.85E-26 | 0.76710698 | 0.262 | 0.159 | 7.73E-22 |
| Tm9sf2 | 3.88E-26 | 0.62985424 | 0.368 | 0.25 | 7.78E-22 |
| Rgs17 | 4.03E-26 | 1.05877411 | 0.14 | 0.067 | 8.09E-22 |
| Synj2bp | 4.22E-26 | 0.96220024 | 0.184 | 0.098 | 8.47E-22 |
| Rhot1 | 4.41E-26 | 0.99514746 | 0.17 | 0.088 | 8.85E-22 |
| Camsap2 | 4.82E-26 | 0.90258738 | 0.187 | 0.101 | 9.67E-22 |
| Ddx10 | 4.88E-26 | 1.11339689 | 0.146 | 0.072 | 9.79E-22 |
| Ercc5 | 5.09E-26 | 1.18229647 | 0.12 | 0.054 | 1.02E-21 |
| Rara | 5.98E-26 | 0.91593896 | 0.192 | 0.104 | 1.20E-21 |
| Ppm1b | 6.38E-26 | 1.01513574 | 0.162 | 0.082 | 1.28E-21 |
| Stxbp3 | 6.76E-26 | 0.9769343 | 0.172 | 0.09 | 1.36E-21 |
| Yap1 | 6.94E-26 | 0.68643042 | 0.297 | 0.188 | 1.39E-21 |
| Upf2 | 7.14E-26 | 0.90000773 | 0.202 | 0.113 | 1.43E-21 |
| Yme1l1 | 9.58E-26 | 0.70633242 | 0.292 | 0.185 | 1.92E-21 |
| Chd3 | 1.11E-25 | 0.61658985 | 0.384 | 0.266 | 2.24E-21 |
| Zbtb44 | 1.21E-25 | 0.94103335 | 0.174 | 0.091 | 2.42E-21 |
| Uimc1 | 1.29E-25 | 0.87921734 | 0.205 | 0.115 | 2.59E-21 |
| Steap3 | 1.34E-25 | 0.9323165 | 0.179 | 0.095 | 2.68E-21 |
| Aatf | 1.38E-25 | 1.18327902 | 0.122 | 0.056 | 2.76E-21 |
| Spast | 1.60E-25 | 1.02019097 | 0.152 | 0.076 | 3.20E-21 |
| Rint1 | 1.63E-25 | 1.27153477 | 0.108 | 0.047 | 3.28E-21 |
| Stam2 | 1.70E-25 | 1.09396454 | 0.138 | 0.066 | 3.41E-21 |
| Tns2 | 1.96E-25 | 1.09194667 | 0.141 | 0.068 | 3.93E-21 |
| Mapk8ip3 | 2.13E-25 | 1.18791513 | 0.116 | 0.052 | 4.28E-21 |
| Phldb2 | 2.33E-25 | 1.14102445 | 0.131 | 0.062 | 4.68E-21 |
| Tcf7l2 | 2.38E-25 | 0.76493392 | 0.288 | 0.184 | 4.78E-21 |
| Smchd1 | 2.57E-25 | 0.75141096 | 0.261 | 0.16 | 5.16E-21 |
| Faf2 | 2.91E-25 | 0.87299691 | 0.205 | 0.115 | 5.85E-21 |
| Zfx | 2.98E-25 | 1.02004123 | 0.173 | 0.092 | 5.99E-21 |
| Slc25a36 | 3.21E-25 | 1.00117431 | 0.159 | 0.081 | 6.44E-21 |
| N4bp2 | 3.33E-25 | 1.0613514 | 0.149 | 0.075 | 6.69E-21 |
| Smurf1 | 3.60E-25 | 1.15988941 | 0.119 | 0.054 | 7.23E-21 |
| Ndnf | 3.87E-25 | 1.09301384 | 0.283 | 0.181 | 7.77E-21 |
| Sgms1 | 4.48E-25 | 1.09023244 | 0.15 | 0.075 | 8.98E-21 |
| Farsb | 4.61E-25 | 1.19063778 | 0.112 | 0.05 | 9.26E-21 |
| Ctdspl | 5.93E-25 | 0.85881113 | 0.221 | 0.128 | 1.19E-20 |
| Wipi1 | 6.49E-25 | 0.63960032 | 0.348 | 0.236 | 1.30E-20 |
| Rev1 | 6.56E-25 | 1.05768746 | 0.143 | 0.071 | 1.32E-20 |
| Brwd1 | 6.57E-25 | 0.86258667 | 0.239 | 0.144 | 1.32E-20 |
| Casc3 | 7.28E-25 | 1.09276928 | 0.135 | 0.065 | 1.46E-20 |
| Pde4a | 8.15E-25 | 1.11372605 | 0.124 | 0.058 | 1.64E-20 |
| Raf1 | 8.59E-25 | 0.73226306 | 0.254 | 0.155 | 1.72E-20 |
| Zfp106 | 8.62E-25 | 0.71268548 | 0.277 | 0.173 | 1.73E-20 |
| Rapgef1 | 9.00E-25 | 1.03314187 | 0.154 | 0.079 | 1.81E-20 |
| Tnks1bp1 | 1.07E-24 | 1.03646852 | 0.147 | 0.074 | 2.15E-20 |
| Supt16 | 1.08E-24 | 0.71890924 | 0.265 | 0.164 | 2.16E-20 |
| Leng8 | 1.15E-24 | 1.05487872 | 0.143 | 0.071 | 2.32E-20 |
| Flywch1 | 1.22E-24 | 1.08753458 | 0.138 | 0.067 | 2.46E-20 |
| Kpna6 | 1.38E-24 | 1.16977205 | 0.112 | 0.05 | 2.77E-20 |
| Lemd3 | 1.39E-24 | 1.10750651 | 0.13 | 0.062 | 2.79E-20 |
| Pum2 | 1.40E-24 | 0.70940876 | 0.317 | 0.209 | 2.81E-20 |
| Fbxo3 | 1.48E-24 | 0.98058968 | 0.155 | 0.079 | 2.97E-20 |
| Gtf3c1 | 1.56E-24 | 1.19658454 | 0.112 | 0.05 | 3.12E-20 |
| Hyou1 | 1.82E-24 | 1.10753037 | 0.122 | 0.057 | 3.66E-20 |
| Xpo6 | 2.43E-24 | 0.97824981 | 0.159 | 0.083 | 4.88E-20 |
| Tmem245 | 2.48E-24 | 0.85945409 | 0.165 | 0.086 | 4.98E-20 |
| Vps50 | 2.64E-24 | 1.29975304 | 0.104 | 0.046 | 5.29E-20 |
| Stox2 | 2.86E-24 | 1.08516157 | 0.174 | 0.095 | 5.74E-20 |
| Snx4 | 3.25E-24 | 0.77700573 | 0.229 | 0.136 | 6.52E-20 |
| Crtc3 | 3.30E-24 | 0.78987829 | 0.24 | 0.145 | 6.62E-20 |
| Ube4b | 3.99E-24 | 0.97607888 | 0.172 | 0.093 | 8.02E-20 |
| Lpin2 | 4.07E-24 | 0.94200443 | 0.185 | 0.102 | 8.18E-20 |
| Dnajc18 | 5.10E-24 | 1.16024616 | 0.116 | 0.053 | 1.02E-19 |
| Fnbp4 | 5.29E-24 | 0.96572634 | 0.173 | 0.094 | 1.06E-19 |
| Dusp16 | 5.73E-24 | 1.13393143 | 0.122 | 0.057 | 1.15E-19 |
| Edem3 | 5.73E-24 | 1.09166696 | 0.136 | 0.067 | 1.15E-19 |
| Nup214 | 6.06E-24 | 1.20469233 | 0.107 | 0.047 | 1.22E-19 |
| Mnat1 | 6.36E-24 | 0.85790009 | 0.222 | 0.132 | 1.28E-19 |
| Spata6 | 6.99E-24 | 1.15447304 | 0.135 | 0.066 | 1.40E-19 |
| Larp4b | 7.28E-24 | 0.73170339 | 0.275 | 0.175 | 1.46E-19 |
| Dnajc24 | 9.29E-24 | 1.13418447 | 0.124 | 0.059 | 1.86E-19 |
| Usp6nl | 9.38E-24 | 1.22471391 | 0.109 | 0.049 | 1.88E-19 |
| Ice1 | 9.78E-24 | 1.03347775 | 0.136 | 0.067 | 1.96E-19 |
| Slc4a4 | 1.38E-23 | 1.08673423 | 0.128 | 0.062 | 2.77E-19 |
| Creb3l2 | 1.56E-23 | 0.69655489 | 0.291 | 0.189 | 3.14E-19 |
| Zdhhc1 | 1.69E-23 | 1.17467323 | 0.106 | 0.047 | 3.39E-19 |
| Zc3h14 | 1.93E-23 | 0.81370951 | 0.207 | 0.119 | 3.87E-19 |
| Sspn | 1.96E-23 | 0.79660031 | 0.2 | 0.114 | 3.93E-19 |
| Pgk1 | 1.99E-23 | 0.69299854 | 0.207 | 0.119 | 3.99E-19 |
| Dchs1 | 2.35E-23 | 0.93785901 | 0.157 | 0.082 | 4.71E-19 |
| Secisbp2l | 2.44E-23 | 0.90879927 | 0.181 | 0.1 | 4.89E-19 |
| Suz12 | 2.82E-23 | 0.91143829 | 0.181 | 0.101 | 5.65E-19 |
| Zic5 | 3.08E-23 | 1.14660816 | 0.113 | 0.052 | 6.18E-19 |
| Emc2 | 3.15E-23 | 0.67741933 | 0.266 | 0.167 | 6.33E-19 |
| Slk | 3.43E-23 | 0.73505009 | 0.259 | 0.162 | 6.88E-19 |
| Kat7 | 3.49E-23 | 1.12389121 | 0.118 | 0.055 | 7.00E-19 |
| Spen | 3.50E-23 | 0.84958825 | 0.187 | 0.104 | 7.03E-19 |
| Mlxip | 4.11E-23 | 0.99233599 | 0.144 | 0.073 | 8.25E-19 |

| | | | | | |
|---|---|---|---|---|---|
| Exoc5 | 4.45E-23 | 0.71766785 | 0.264 | 0.166 | 8.92E-19 |
| Wnt5b | 4.71E-23 | 1.14390527 | 0.222 | 0.135 | 9.46E-19 |
| Tmem45a | 5.45E-23 | 0.78358718 | 0.272 | 0.175 | 1.09E-18 |
| 5730455P16 | 5.57E-23 | 1.10007661 | 0.122 | 0.058 | 1.12E-18 |
| Gnai1 | 5.99E-23 | 1.15007125 | 0.122 | 0.059 | 1.20E-18 |
| Pdgfd | 6.07E-23 | 1.36156478 | 0.131 | 0.065 | 1.22E-18 |
| Zcchc8 | 6.14E-23 | 1.13915251 | 0.11 | 0.05 | 1.23E-18 |
| Sel1l | 6.61E-23 | 0.71618131 | 0.272 | 0.173 | 1.33E-18 |
| Oxr1 | 7.17E-23 | 0.91738376 | 0.192 | 0.11 | 1.44E-18 |
| Zfp644 | 7.82E-23 | 0.70756316 | 0.28 | 0.18 | 1.57E-18 |
| Rc3h1 | 7.93E-23 | 0.72080356 | 0.259 | 0.163 | 1.59E-18 |
| Sdha | 8.27E-23 | 0.70842653 | 0.266 | 0.168 | 1.66E-18 |
| Tmtc3 | 8.30E-23 | 1.18879434 | 0.101 | 0.044 | 1.66E-18 |
| Mast2 | 8.35E-23 | 0.88315376 | 0.193 | 0.11 | 1.68E-18 |
| March7 | 9.22E-23 | 0.6985821 | 0.25 | 0.154 | 1.85E-18 |
| Adam17 | 9.77E-23 | 0.90984057 | 0.183 | 0.103 | 1.96E-18 |
| Ccnt2 | 9.96E-23 | 0.8521491 | 0.195 | 0.111 | 2.00E-18 |
| 2610507B11 | 1.03E-22 | 0.79532337 | 0.213 | 0.125 | 2.06E-18 |
| Scfd1 | 1.10E-22 | 0.80653984 | 0.223 | 0.134 | 2.20E-18 |
| Wdr20 | 1.10E-22 | 1.08092474 | 0.126 | 0.061 | 2.21E-18 |
| Fip1l1 | 1.17E-22 | 0.71480008 | 0.261 | 0.165 | 2.35E-18 |
| Orc4 | 1.18E-22 | 1.00632926 | 0.144 | 0.074 | 2.37E-18 |
| Phtf1 | 1.27E-22 | 0.98279985 | 0.148 | 0.076 | 2.54E-18 |
| Mvp | 1.30E-22 | 0.77281112 | 0.2 | 0.116 | 2.60E-18 |
| Ric1 | 1.33E-22 | 0.82207344 | 0.186 | 0.105 | 2.66E-18 |
| Katnbl1 | 1.52E-22 | 1.11922078 | 0.128 | 0.063 | 3.04E-18 |
| Diaph1 | 1.89E-22 | 0.87875546 | 0.189 | 0.108 | 3.80E-18 |
| 1810013L24 | 2.02E-22 | 0.86283137 | 0.181 | 0.101 | 4.06E-18 |
| Srsf4 | 2.04E-22 | 0.91075886 | 0.169 | 0.092 | 4.09E-18 |
| Rab8b | 2.27E-22 | 0.65761118 | 0.241 | 0.147 | 4.56E-18 |
| Ube2w | 2.38E-22 | 0.80847346 | 0.225 | 0.136 | 4.77E-18 |
| Il1r1 | 2.41E-22 | 0.71472331 | 0.294 | 0.195 | 4.83E-18 |
| Dlg5 | 2.54E-22 | 1.02371975 | 0.13 | 0.065 | 5.09E-18 |
| Rsu1 | 2.78E-22 | 0.67404808 | 0.323 | 0.22 | 5.58E-18 |
| Chuk | 3.74E-22 | 1.00192034 | 0.155 | 0.083 | 7.51E-18 |
| Ogdh | 4.12E-22 | 0.72172291 | 0.261 | 0.166 | 8.27E-18 |
| Arhgap17 | 4.40E-22 | 0.91163882 | 0.167 | 0.092 | 8.83E-18 |
| Azin2 | 4.75E-22 | 0.97568018 | 0.152 | 0.08 | 9.54E-18 |
| Cdk5rap2 | 4.79E-22 | 1.00649801 | 0.13 | 0.065 | 9.61E-18 |
| Vps37a | 4.87E-22 | 0.88032328 | 0.17 | 0.094 | 9.77E-18 |
| Fzd2 | 5.01E-22 | 0.68205207 | 0.255 | 0.161 | 1.00E-17 |
| Blvrb | 5.67E-22 | 0.62552879 | 0.32 | 0.216 | 1.14E-17 |
| Hbs1l | 5.93E-22 | 0.97677537 | 0.144 | 0.075 | 1.19E-17 |
| Wdr60 | 6.55E-22 | 1.16242963 | 0.11 | 0.051 | 1.31E-17 |
| Efna1 | 6.68E-22 | 1.06815957 | 0.122 | 0.059 | 1.34E-17 |
| Pwwp2a | 6.94E-22 | 0.92884811 | 0.149 | 0.078 | 1.39E-17 |
| Axin1 | 7.22E-22 | 1.02551874 | 0.126 | 0.062 | 1.45E-17 |
| Med14 | 7.89E-22 | 0.94298189 | 0.14 | 0.072 | 1.58E-17 |
| Lgr4 | 7.96E-22 | 1.05399175 | 0.152 | 0.081 | 1.60E-17 |
| Msantd2 | 8.36E-22 | 1.16565635 | 0.102 | 0.046 | 1.68E-17 |
| Sulf2 | 8.81E-22 | 0.99678694 | 0.127 | 0.063 | 1.77E-17 |
| Lars | 9.15E-22 | 0.87720241 | 0.179 | 0.101 | 1.84E-17 |
| Cpsf7 | 9.51E-22 | 0.864267 | 0.181 | 0.102 | 1.91E-17 |
| Sec61a2 | 1.08E-21 | 0.90042184 | 0.168 | 0.093 | 2.18E-17 |
| Trim3 | 1.19E-21 | 1.00493587 | 0.136 | 0.069 | 2.38E-17 |
| Taf3 | 1.31E-21 | 0.93856393 | 0.164 | 0.09 | 2.63E-17 |
| Mapk8 | 1.33E-21 | 0.91120025 | 0.167 | 0.092 | 2.68E-17 |
| Drg1 | 1.41E-21 | 0.87122455 | 0.183 | 0.105 | 2.82E-17 |
| Dstyk | 1.45E-21 | 0.90132153 | 0.163 | 0.089 | 2.91E-17 |
| Hipk3 | 1.48E-21 | 0.70716302 | 0.245 | 0.153 | 2.98E-17 |
| Mamdc2 | 1.70E-21 | 0.86429143 | 0.101 | 0.046 | 3.41E-17 |
| Dact1 | 1.81E-21 | 0.91681674 | 0.194 | 0.114 | 3.62E-17 |
| Atp6v1h | 1.84E-21 | 0.82948593 | 0.203 | 0.121 | 3.69E-17 |
| Agfg1 | 3.04E-21 | 0.84053425 | 0.182 | 0.104 | 6.11E-17 |
| Cfap97 | 3.07E-21 | 1.15411191 | 0.101 | 0.046 | 6.16E-17 |
| Stat6 | 3.45E-21 | 0.99469456 | 0.125 | 0.062 | 6.92E-17 |
| Ep400 | 3.55E-21 | 0.68181959 | 0.251 | 0.159 | 7.12E-17 |
| Zc3h13 | 3.83E-21 | 0.70110704 | 0.253 | 0.16 | 7.68E-17 |
| Ppme1 | 4.15E-21 | 0.8311489 | 0.176 | 0.099 | 8.32E-17 |
| Trpc4ap | 4.58E-21 | 0.76423876 | 0.214 | 0.129 | 9.19E-17 |
| Rai1 | 4.77E-21 | 0.90764861 | 0.159 | 0.087 | 9.58E-17 |
| Golga4 | 5.02E-21 | 0.77105162 | 0.226 | 0.14 | 1.01E-16 |
| Golga5 | 5.69E-21 | 0.93117343 | 0.155 | 0.084 | 1.14E-16 |
| Cdadc1 | 6.41E-21 | 1.06563047 | 0.106 | 0.05 | 1.29E-16 |
| Fxr1 | 6.57E-21 | 0.64856189 | 0.283 | 0.187 | 1.32E-16 |
| Ppp4r1 | 6.62E-21 | 1.0972643 | 0.124 | 0.062 | 1.33E-16 |
| Lpcat3 | 7.35E-21 | 1.03261645 | 0.126 | 0.063 | 1.47E-16 |
| Rcor1 | 7.79E-21 | 0.89175606 | 0.175 | 0.099 | 1.56E-16 |
| Fam45a | 7.89E-21 | 1.12075962 | 0.111 | 0.053 | 1.58E-16 |
| Smim1 | 8.49E-21 | 1.05963472 | 0.12 | 0.059 | 1.70E-16 |
| Hoxa10 | 1.00E-20 | 0.67343454 | 0.23 | 0.142 | 2.02E-16 |
| Usp4 | 1.07E-20 | 0.91481146 | 0.156 | 0.085 | 2.15E-16 |
| Hnrnph2 | 1.16E-20 | 0.68972246 | 0.245 | 0.155 | 2.33E-16 |
| Stt3b | 1.19E-20 | 0.65585079 | 0.301 | 0.204 | 2.38E-16 |
| Cluap1 | 1.32E-20 | 1.11744836 | 0.103 | 0.048 | 2.64E-16 |
| Bcor | 1.35E-20 | 0.79467294 | 0.186 | 0.108 | 2.71E-16 |
| Bcl3 | 1.48E-20 | 0.64220623 | 0.187 | 0.108 | 2.98E-16 |
| Rbm10 | 1.51E-20 | 1.15322287 | 0.102 | 0.047 | 3.03E-16 |
| Cacul1 | 1.60E-20 | 0.95825854 | 0.131 | 0.067 | 3.22E-16 |
| Npas2 | 1.64E-20 | 1.09918303 | 0.108 | 0.051 | 3.29E-16 |
| Hoxa9 | 1.81E-20 | 0.87443228 | 0.162 | 0.09 | 3.64E-16 |
| Pank3 | 1.92E-20 | 0.89330448 | 0.152 | 0.083 | 3.85E-16 |
| Pex13 | 1.99E-20 | 0.97228088 | 0.136 | 0.071 | 3.99E-16 |
| Exoc3 | 2.09E-20 | 0.82841297 | 0.178 | 0.102 | 4.20E-16 |
| Cul4a | 2.39E-20 | 0.83681634 | 0.191 | 0.112 | 4.79E-16 |
| Nfxl1 | 2.49E-20 | 1.12734625 | 0.112 | 0.054 | 5.00E-16 |
| Eif4enif1 | 2.67E-20 | 0.908851 | 0.14 | 0.074 | 5.35E-16 |
| Tspan11 | 2.85E-20 | 0.69144061 | 0.24 | 0.152 | 5.71E-16 |
| Dmtf1 | 2.85E-20 | 1.00686828 | 0.129 | 0.066 | 5.73E-16 |
| Fmc1 | 3.27E-20 | 0.92280661 | 0.128 | 0.065 | 6.56E-16 |
| Col24a1 | 3.29E-20 | 1.1309119 | 0.112 | 0.055 | 6.60E-16 |
| Cramp1l | 3.42E-20 | 1.08831294 | 0.101 | 0.047 | 6.85E-16 |
| Nus1 | 3.85E-20 | 0.69712219 | 0.235 | 0.148 | 7.73E-16 |
| Cep85 | 3.91E-20 | 1.04738403 | 0.118 | 0.059 | 7.86E-16 |
| Epm2aip1 | 4.02E-20 | 1.04914235 | 0.113 | 0.055 | 8.07E-16 |
| Marf1 | 4.36E-20 | 0.81648159 | 0.175 | 0.1 | 8.74E-16 |
| Kpna3 | 4.49E-20 | 0.91927888 | 0.16 | 0.089 | 9.01E-16 |
| Fubp3 | 4.73E-20 | 0.92408487 | 0.148 | 0.08 | 9.48E-16 |
| 2310033P09 | 4.81E-20 | 1.03354549 | 0.119 | 0.06 | 9.66E-16 |
| Tmx4 | 5.00E-20 | 0.82644143 | 0.181 | 0.105 | 1.00E-15 |
| Thoc1 | 5.16E-20 | 0.99693155 | 0.14 | 0.075 | 1.04E-15 |
| 1500011B03 | 6.99E-20 | 1.07272917 | 0.109 | 0.053 | 1.40E-15 |
| Rin2 | 7.15E-20 | 0.63455352 | 0.289 | 0.195 | 1.44E-15 |
| Gatad1 | 8.61E-20 | 0.66728704 | 0.288 | 0.195 | 1.73E-15 |
| Foxo3 | 8.97E-20 | 0.82522633 | 0.183 | 0.107 | 1.80E-15 |
| Pdia4 | 9.37E-20 | 0.66261501 | 0.24 | 0.152 | 1.88E-15 |
| Nrbp2 | 9.91E-20 | 1.03238612 | 0.122 | 0.062 | 1.99E-15 |
| Elmo2 | 1.00E-19 | 0.9761629 | 0.125 | 0.064 | 2.01E-15 |
| Cpsf6 | 1.03E-19 | 0.68861141 | 0.256 | 0.167 | 2.07E-15 |
| Elf4 | 1.06E-19 | 1.10819677 | 0.106 | 0.051 | 2.12E-15 |
| Ablim1 | 1.15E-19 | 0.95934568 | 0.123 | 0.062 | 2.30E-15 |
| Hcfc1 | 1.18E-19 | 0.90967845 | 0.14 | 0.075 | 2.37E-15 |
| Atad1 | 1.18E-19 | 0.84750848 | 0.181 | 0.106 | 2.37E-15 |
| Ptpn12 | 1.22E-19 | 0.7248892 | 0.251 | 0.164 | 2.45E-15 |
| Ivd | 1.33E-19 | 1.06877115 | 0.109 | 0.053 | 2.66E-15 |
| 1110008P14 | 1.36E-19 | 0.7283396 | 0.264 | 0.176 | 2.74E-15 |
| Bcl2l1 | 1.40E-19 | 1.03958171 | 0.111 | 0.054 | 2.81E-15 |
| Cnot6l | 1.65E-19 | 0.88161186 | 0.178 | 0.104 | 3.30E-15 |
| Ism1 | 1.68E-19 | 0.93392689 | 0.186 | 0.111 | 3.37E-15 |
| Hcfc2 | 1.69E-19 | 1.11327268 | 0.106 | 0.051 | 3.40E-15 |
| Dnajb2 | 1.72E-19 | 1.01589054 | 0.114 | 0.057 | 3.46E-15 |
| Acbd6 | 1.86E-19 | 0.66475016 | 0.252 | 0.163 | 3.74E-15 |
| Rras | 1.90E-19 | 0.65225219 | 0.281 | 0.19 | 3.81E-15 |
| Cry2 | 2.11E-19 | 0.94233273 | 0.137 | 0.073 | 4.24E-15 |
| Fam120a | 2.41E-19 | 0.64810771 | 0.291 | 0.199 | 4.83E-15 |
| Tmco3 | 2.52E-19 | 0.92203043 | 0.134 | 0.071 | 5.07E-15 |
| Eng | 2.53E-19 | 0.80096259 | 0.17 | 0.097 | 5.08E-15 |
| Palm | 2.71E-19 | 0.96676582 | 0.135 | 0.072 | 5.44E-15 |
| Podn | 2.93E-19 | 0.91657323 | 0.125 | 0.064 | 5.88E-15 |
| Echs1 | 3.07E-19 | 0.66410145 | 0.234 | 0.148 | 6.16E-15 |
| Gtf2f2 | 3.09E-19 | 0.95465548 | 0.132 | 0.07 | 6.20E-15 |
| Pde4dip | 3.18E-19 | 0.84655116 | 0.151 | 0.083 | 6.38E-15 |
| Cep164 | 3.32E-19 | 0.86810389 | 0.159 | 0.09 | 6.67E-15 |
| Cabin1 | 3.63E-19 | 0.977315 | 0.115 | 0.057 | 7.29E-15 |
| Trim24 | 3.75E-19 | 1.03663593 | 0.115 | 0.058 | 7.52E-15 |
| Ndufs1 | 3.92E-19 | 0.87805622 | 0.158 | 0.089 | 7.86E-15 |
| Clpx | 4.69E-19 | 0.93463046 | 0.135 | 0.072 | 9.42E-15 |
| Smo | 5.13E-19 | 0.65989514 | 0.249 | 0.162 | 1.03E-14 |
| Dars | 5.14E-19 | 0.87754271 | 0.163 | 0.093 | 1.03E-14 |
| Cryzl1 | 5.16E-19 | 0.89797827 | 0.149 | 0.082 | 1.04E-14 |
| Rnf13 | 5.38E-19 | 0.66728915 | 0.262 | 0.173 | 1.08E-14 |
| Magt1 | 6.31E-19 | 0.67204213 | 0.256 | 0.169 | 1.27E-14 |
| Ergic1 | 6.88E-19 | 0.71530748 | 0.209 | 0.129 | 1.38E-14 |
| Adamts9 | 7.04E-19 | 1.00240809 | 0.178 | 0.106 | 1.41E-14 |
| Txndc15 | 7.10E-19 | 0.66302446 | 0.24 | 0.154 | 1.42E-14 |
| Zc3h7b | 7.27E-19 | 0.94729853 | 0.133 | 0.071 | 1.46E-14 |
| Tank | 8.24E-19 | 0.9457492 | 0.138 | 0.074 | 1.65E-14 |
| Rusc2 | 8.31E-19 | 1.06386385 | 0.112 | 0.056 | 1.67E-14 |
| Mapre3 | 9.48E-19 | 0.96240502 | 0.133 | 0.071 | 1.90E-14 |
| Zkscan3 | 1.02E-18 | 0.78248156 | 0.173 | 0.101 | 2.04E-14 |
| Zfp322a | 1.02E-18 | 1.00904069 | 0.117 | 0.06 | 2.05E-14 |
| Lgals9 | 1.05E-18 | 0.76340489 | 0.184 | 0.11 | 2.11E-14 |
| Klc1 | 1.09E-18 | 0.70768981 | 0.228 | 0.145 | 2.18E-14 |
| Rufy3 | 1.23E-18 | 0.98912855 | 0.118 | 0.06 | 2.47E-14 |
| Myo1e | 1.24E-18 | 0.73226551 | 0.238 | 0.155 | 2.49E-14 |
| Fut11 | 1.26E-18 | 0.86506686 | 0.143 | 0.079 | 2.52E-14 |
| Pex14 | 1.42E-18 | 1.04247862 | 0.122 | 0.063 | 2.86E-14 |
| Tnpo3 | 1.57E-18 | 0.69702954 | 0.203 | 0.125 | 3.16E-14 |
| Synpo | 1.86E-18 | 0.73572575 | 0.205 | 0.127 | 3.74E-14 |
| Mtf2 | 1.89E-18 | 0.87531289 | 0.157 | 0.09 | 3.78E-14 |
| Evc2 | 1.91E-18 | 1.04330541 | 0.108 | 0.054 | 3.84E-14 |
| Bmyc | 2.01E-18 | 0.85093396 | 0.159 | 0.091 | 4.04E-14 |
| N4bp1 | 2.13E-18 | 0.93310191 | 0.119 | 0.061 | 4.27E-14 |
| Acadl | 2.27E-18 | 0.75935789 | 0.2 | 0.123 | 4.56E-14 |
| Tacc2 | 2.39E-18 | 0.84315573 | 0.149 | 0.083 | 4.80E-14 |
| Uba6 | 2.48E-18 | 0.93160192 | 0.126 | 0.066 | 4.97E-14 |
| D430042O09 | 2.64E-18 | 0.94446337 | 0.115 | 0.059 | 5.29E-14 |
| Osbpl1a | 2.81E-18 | 0.6923236 | 0.216 | 0.136 | 5.64E-14 |
| Abca5 | 2.89E-18 | 1.05313321 | 0.104 | 0.051 | 5.79E-14 |
| Cdyl | 3.64E-18 | 0.90269878 | 0.135 | 0.073 | 7.31E-14 |
| Trappc10 | 3.75E-18 | 1.06756782 | 0.111 | 0.056 | 7.53E-14 |
| 4833420G17 | 3.78E-18 | 0.90033864 | 0.138 | 0.076 | 7.59E-14 |
| Csnk1g3 | 3.85E-18 | 0.70785892 | 0.214 | 0.135 | 7.72E-14 |
| Fbxo8 | 4.12E-18 | 0.96383503 | 0.117 | 0.061 | 8.28E-14 |
| Snap23 | 4.23E-18 | 0.74368445 | 0.206 | 0.129 | 8.48E-14 |
| Ublcp1 | 4.77E-18 | 0.75587801 | 0.179 | 0.106 | 9.57E-14 |
| Tead3 | 5.13E-18 | 0.99771411 | 0.11 | 0.056 | 1.03E-13 |
| Trim56 | 5.54E-18 | 0.93130586 | 0.112 | 0.057 | 1.11E-13 |
| Prpf8 | 6.00E-18 | 0.73147628 | 0.195 | 0.119 | 1.20E-13 |
| Gtf3c2 | 6.60E-18 | 0.79546599 | 0.168 | 0.098 | 1.32E-13 |
| Grb10 | 7.04E-18 | 0.60693247 | 0.384 | 0.293 | 1.41E-13 |
| Rom1 | 7.07E-18 | 0.73650914 | 0.127 | 0.067 | 1.42E-13 |
| Sec16a | 8.67E-18 | 0.85242133 | 0.143 | 0.08 | 1.74E-13 |
| Nr1d2 | 1.16E-17 | 0.7561104 | 0.191 | 0.117 | 2.33E-13 |
| Rnf170 | 1.18E-17 | 1.01954756 | 0.111 | 0.057 | 2.37E-13 |
| Polr2b | 1.52E-17 | 0.97673385 | 0.118 | 0.062 | 3.04E-13 |
| Ppp1r9b | 1.89E-17 | 0.8673662 | 0.136 | 0.075 | 3.80E-13 |
| Srebf2 | 2.56E-17 | 0.78166204 | 0.17 | 0.102 | 5.13E-13 |
| Plaa | 2.60E-17 | 0.80735567 | 0.163 | 0.096 | 5.21E-13 |
| Ktn1 | 3.19E-17 | 0.6532671 | 0.277 | 0.193 | 6.41E-13 |
| Bdp1 | 3.31E-17 | 0.76643887 | 0.186 | 0.114 | 6.64E-13 |
| Uggt1 | 3.56E-17 | 0.76045276 | 0.167 | 0.099 | 7.14E-13 |
| Plin3 | 3.57E-17 | 0.69338066 | 0.204 | 0.128 | 7.16E-13 |
| Dusp7 | 3.77E-17 | 0.79668423 | 0.148 | 0.084 | 7.57E-13 |
| Klhl5 | 3.96E-17 | 0.88635941 | 0.15 | 0.086 | 7.95E-13 |
| Ly6c1 | 4.10E-17 | 0.61653222 | 0.45 | 0.36 | 8.22E-13 |
| Ttc39b | 4.47E-17 | 0.75918664 | 0.163 | 0.096 | 8.96E-13 |
| Trf | 4.75E-17 | 1.33938976 | 0.122 | 0.066 | 9.54E-13 |
| Ppil4 | 4.78E-17 | 0.7078263 | 0.193 | 0.12 | 9.59E-13 |
| 5031425E22 | 4.90E-17 | 0.6844742 | 0.227 | 0.148 | 9.83E-13 |
| Brcc3 | 5.01E-17 | 0.97178346 | 0.106 | 0.054 | 1.01E-12 |
| Arih2 | 5.67E-17 | 0.77245584 | 0.169 | 0.101 | 1.14E-12 |
| Alg14 | 5.88E-17 | 0.85376748 | 0.136 | 0.076 | 1.18E-12 |
| Atp6v1a | 5.91E-17 | 0.62895567 | 0.238 | 0.158 | 1.19E-12 |
| Ube2g1 | 6.74E-17 | 0.74512689 | 0.208 | 0.133 | 1.35E-12 |
| Swap70 | 6.77E-17 | 0.8311332 | 0.144 | 0.082 | 1.36E-12 |
| Mgrn1 | 7.13E-17 | 0.812992 | 0.147 | 0.084 | 1.43E-12 |
| Hnrnph3 | 8.38E-17 | 0.68836149 | 0.211 | 0.135 | 1.68E-12 |
| Papss1 | 8.73E-17 | 0.793812 | 0.206 | 0.132 | 1.75E-12 |
| Chst2 | 8.95E-17 | 0.90682181 | 0.148 | 0.086 | 1.80E-12 |
| Tjp1 | 9.52E-17 | 0.806461 | 0.139 | 0.078 | 1.91E-12 |
| Ldb1 | 9.87E-17 | 0.70940297 | 0.185 | 0.114 | 1.98E-12 |
| Smarcc1 | 1.00E-16 | 0.69260769 | 0.196 | 0.123 | 2.01E-12 |
| Ssbp4 | 1.01E-16 | 0.6540299 | 0.24 | 0.16 | 2.02E-12 |
| Cat | 1.19E-16 | 0.88380461 | 0.14 | 0.079 | 2.40E-12 |
| Zdhhc6 | 1.39E-16 | 0.96688466 | 0.114 | 0.061 | 2.78E-12 |
| Lats1 | 1.44E-16 | 0.78566801 | 0.157 | 0.092 | 2.89E-12 |
| Maoa | 1.68E-16 | 0.96533859 | 0.116 | 0.062 | 3.36E-12 |
| Smad2 | 1.78E-16 | 0.84798376 | 0.149 | 0.087 | 3.56E-12 |
| Cpeb4 | 2.07E-16 | 0.64599674 | 0.189 | 0.117 | 4.16E-12 |
| Unc93b1 | 2.11E-16 | 0.89979371 | 0.121 | 0.066 | 4.23E-12 |
| Camsap1 | 2.12E-16 | 0.98316831 | 0.109 | 0.057 | 4.26E-12 |
| Ubac2 | 2.16E-16 | 0.81659006 | 0.16 | 0.096 | 4.34E-12 |
| Cpeb1 | 2.18E-16 | 1.12663379 | 0.102 | 0.052 | 4.37E-12 |
| Banp | 2.42E-16 | 0.91509438 | 0.112 | 0.059 | 4.85E-12 |
| Sec23b | 2.83E-16 | 0.83757906 | 0.156 | 0.092 | 5.68E-12 |
| Siah1a | 3.35E-16 | 0.87717242 | 0.136 | 0.077 | 6.72E-12 |
| Ganab | 3.43E-16 | 0.73577626 | 0.191 | 0.121 | 6.87E-12 |
| Ccpg1 | 3.50E-16 | 0.8632758 | 0.138 | 0.078 | 7.02E-12 |
| Tbce | 3.63E-16 | 0.98377916 | 0.104 | 0.054 | 7.29E-12 |
| Adgrl1 | 3.84E-16 | 0.62397115 | 0.226 | 0.148 | 7.70E-12 |
| Btbd1 | 3.95E-16 | 0.65549156 | 0.229 | 0.152 | 7.92E-12 |
| Suco | 4.30E-16 | 0.72418019 | 0.163 | 0.097 | 8.64E-12 |
| Rilpl1 | 4.44E-16 | 0.91163882 | 0.111 | 0.059 | 8.91E-12 |
| Vapb | 4.67E-16 | 0.62872439 | 0.216 | 0.14 | 9.37E-12 |
| Rad17 | 4.88E-16 | 1.03723558 | 0.103 | 0.053 | 9.80E-12 |
| Cdc23 | 5.00E-16 | 0.9571241 | 0.109 | 0.057 | 1.00E-11 |
| Lztr1 | 5.12E-16 | 0.88594321 | 0.126 | 0.07 | 1.03E-11 |
| Ccdc102a | 5.91E-16 | 0.8759826 | 0.119 | 0.064 | 1.19E-11 |
| Nvl | 6.27E-16 | 0.84893478 | 0.129 | 0.072 | 1.26E-11 |
| Ubp1 | 6.44E-16 | 0.77881545 | 0.167 | 0.101 | 1.29E-11 |
| Pcyt2 | 6.86E-16 | 0.9569197 | 0.104 | 0.054 | 1.38E-11 |
| Msi2 | 7.19E-16 | 0.81229575 | 0.132 | 0.074 | 1.44E-11 |
| Kmt2d | 8.58E-16 | 0.83372959 | 0.13 | 0.073 | 1.72E-11 |
| Papss2 | 1.01E-15 | 0.78696389 | 0.144 | 0.084 | 2.02E-11 |
| Capza1 | 1.14E-15 | 0.61472158 | 0.229 | 0.153 | 2.29E-11 |
| Zfp651 | 1.18E-15 | 0.74797281 | 0.16 | 0.097 | 2.37E-11 |
| Tef | 1.63E-15 | 0.85970841 | 0.14 | 0.081 | 3.26E-11 |
| Morc3 | 1.67E-15 | 0.86186378 | 0.12 | 0.066 | 3.36E-11 |
| Arid4a | 1.79E-15 | 0.67309214 | 0.199 | 0.128 | 3.59E-11 |
| Rnf115 | 2.04E-15 | 0.84750848 | 0.145 | 0.086 | 4.10E-11 |
| Rxra | 2.27E-15 | 0.95415565 | 0.106 | 0.056 | 4.55E-11 |
| Golga3 | 2.34E-15 | 0.75512025 | 0.152 | 0.091 | 4.70E-11 |
| Pcbd2 | 2.53E-15 | 0.72049733 | 0.172 | 0.106 | 5.07E-11 |
| Snrnp200 | 2.60E-15 | 0.85397139 | 0.126 | 0.071 | 5.21E-11 |
| Dguok | 2.62E-15 | 0.78186833 | 0.154 | 0.093 | 5.26E-11 |
| Akap8 | 2.74E-15 | 0.79620605 | 0.134 | 0.077 | 5.50E-11 |
| Setd1b | 3.03E-15 | 0.71790317 | 0.162 | 0.098 | 6.08E-11 |
| Epb41l1 | 3.13E-15 | 0.92031237 | 0.107 | 0.057 | 6.29E-11 |
| Rnpc3 | 3.14E-15 | 0.92052791 | 0.108 | 0.058 | 6.31E-11 |
| Fam117b | 3.16E-15 | 0.85304867 | 0.143 | 0.084 | 6.34E-11 |
| Dnajc7 | 3.18E-15 | 0.63044411 | 0.212 | 0.139 | 6.37E-11 |
| Btbd10 | 3.56E-15 | 0.94169606 | 0.11 | 0.059 | 7.15E-11 |
| Setd3 | 3.83E-15 | 0.64475209 | 0.205 | 0.134 | 7.68E-11 |
| Rhoj | 3.99E-15 | 0.60137904 | 0.257 | 0.179 | 8.02E-11 |
| Dnm1l | 4.42E-15 | 0.75846519 | 0.161 | 0.098 | 8.87E-11 |
| Ipo9 | 4.49E-15 | 0.88994375 | 0.115 | 0.063 | 9.00E-11 |
| Ifnar1 | 4.55E-15 | 0.80309866 | 0.141 | 0.082 | 9.13E-11 |
| Sdk2 | 4.70E-15 | 0.83419611 | 0.163 | 0.101 | 9.43E-11 |
| Mtmr11 | 5.43E-15 | 0.79498288 | 0.136 | 0.079 | 1.09E-10 |
| Brip1os | 5.87E-15 | 0.81659403 | 0.151 | 0.091 | 1.18E-10 |
| Usp7 | 6.09E-15 | 0.63851003 | 0.224 | 0.151 | 1.22E-10 |
| Ppp6c | 6.71E-15 | 0.72814834 | 0.176 | 0.111 | 1.35E-10 |
| Prkd3 | 6.72E-15 | 0.69610865 | 0.178 | 0.112 | 1.35E-10 |
| Hoxa13 | 6.87E-15 | 0.68748335 | 0.173 | 0.108 | 1.38E-10 |
| Angpt1 | 7.49E-15 | 0.84888557 | 0.143 | 0.085 | 1.50E-10 |
| Ttc7 | 8.66E-15 | 0.83763824 | 0.138 | 0.081 | 1.74E-10 |
| Lgals8 | 9.83E-15 | 0.73271024 | 0.154 | 0.093 | 1.97E-10 |
| Ccdc186 | 1.04E-14 | 0.65066744 | 0.19 | 0.122 | 2.09E-10 |
| Tle2 | 1.07E-14 | 0.92670627 | 0.103 | 0.055 | 2.15E-10 |
| Sec22a | 1.20E-14 | 0.92575993 | 0.124 | 0.071 | 2.41E-10 |
| Arfgap3 | 1.22E-14 | 0.61202531 | 0.219 | 0.146 | 2.46E-10 |
| Iws1 | 1.43E-14 | 0.76903443 | 0.14 | 0.083 | 2.86E-10 |
| Slc22a17 | 1.43E-14 | 0.68130258 | 0.187 | 0.12 | 2.86E-10 |
| Cul7 | 1.74E-14 | 0.90437082 | 0.106 | 0.058 | 3.49E-10 |
| Uvssa | 1.96E-14 | 0.7611912 | 0.136 | 0.08 | 3.94E-10 |
| Fbxo4 | 2.01E-14 | 0.83655088 | 0.125 | 0.071 | 4.04E-10 |
| Hadha | 2.12E-14 | 0.60794063 | 0.208 | 0.138 | 4.25E-10 |
| Dhx30 | 2.36E-14 | 0.74622515 | 0.156 | 0.095 | 4.74E-10 |
| Aldh1a3 | 2.62E-14 | 0.64742623 | 0.17 | 0.107 | 5.27E-10 |
| Gabpa | 2.70E-14 | 0.74948614 | 0.14 | 0.083 | 5.41E-10 |
| Jmy | 2.71E-14 | 0.71062576 | 0.163 | 0.102 | 5.43E-10 |
| Znrf1 | 2.73E-14 | 0.66113505 | 0.215 | 0.145 | 5.48E-10 |
| Arhgef19 | 2.85E-14 | 0.74494345 | 0.152 | 0.092 | 5.73E-10 |
| Cyb5b | 3.29E-14 | 0.72332033 | 0.168 | 0.106 | 6.60E-10 |
| 1700025G04 | 3.86E-14 | 0.63571231 | 0.222 | 0.152 | 7.74E-10 |
| Naa35 | 3.91E-14 | 0.7357981 | 0.141 | 0.084 | 7.85E-10 |
| Npr2 | 4.50E-14 | 0.68349287 | 0.161 | 0.1 | 9.04E-10 |
| Psmd5 | 5.15E-14 | 0.87233917 | 0.11 | 0.061 | 1.03E-09 |
| Ift74 | 5.29E-14 | 0.69874577 | 0.163 | 0.102 | 1.06E-09 |
| Mapk9 | 5.47E-14 | 0.8023956 | 0.117 | 0.066 | 1.10E-09 |
| Mcu | 6.03E-14 | 0.8076411 | 0.126 | 0.073 | 1.21E-09 |
| Atp11b | 6.16E-14 | 0.68628399 | 0.157 | 0.097 | 1.24E-09 |
| Nin | 6.22E-14 | 0.77571277 | 0.138 | 0.082 | 1.25E-09 |
| Arl2 | 6.28E-14 | 0.6089964 | 0.202 | 0.133 | 1.26E-09 |
| Srebf1 | 6.53E-14 | 0.73521971 | 0.147 | 0.089 | 1.31E-09 |
| Phrf1 | 6.56E-14 | 0.84429535 | 0.114 | 0.064 | 1.32E-09 |
| Gpbp1l1 | 6.68E-14 | 0.73364052 | 0.155 | 0.096 | 1.34E-09 |
| Afap1 | 6.81E-14 | 0.83833469 | 0.142 | 0.086 | 1.37E-09 |
| Spats2l | 8.98E-14 | 0.62629173 | 0.271 | 0.195 | 1.80E-09 |
| Smarcd2 | 9.07E-14 | 0.74198371 | 0.13 | 0.076 | 1.82E-09 |
| Ltbr | 9.40E-14 | 0.68904276 | 0.163 | 0.102 | 1.89E-09 |
| Src | 1.01E-13 | 0.90724928 | 0.104 | 0.057 | 2.02E-09 |
| Tmem127 | 1.13E-13 | 0.80159645 | 0.119 | 0.068 | 2.26E-09 |
| Ldlrad3 | 1.22E-13 | 0.83341365 | 0.119 | 0.068 | 2.45E-09 |
| Osbpl8 | 1.59E-13 | 0.64285732 | 0.196 | 0.131 | 3.18E-09 |
| Dcaf10 | 1.60E-13 | 0.75591456 | 0.148 | 0.091 | 3.21E-09 |
| Eif4e3 | 1.77E-13 | 0.72500492 | 0.15 | 0.093 | 3.55E-09 |
| Antxr2 | 2.27E-13 | 0.63238837 | 0.171 | 0.11 | 4.56E-09 |
| Cpeb2 | 2.38E-13 | 0.77927591 | 0.157 | 0.099 | 4.77E-09 |
| Zswim6 | 2.57E-13 | 0.6480856 | 0.156 | 0.097 | 5.15E-09 |
| Ptpn13 | 2.67E-13 | 0.69422584 | 0.159 | 0.1 | 5.36E-09 |
| Git2 | 2.92E-13 | 0.76986995 | 0.142 | 0.087 | 5.86E-09 |
| Dgcr2 | 3.01E-13 | 0.69527136 | 0.157 | 0.099 | 6.04E-09 |
| Ccnc | 3.96E-13 | 0.89452221 | 0.104 | 0.058 | 7.94E-09 |
| Pphln1 | 4.32E-13 | 0.84243155 | 0.109 | 0.062 | 8.66E-09 |
| Rnf145 | 4.44E-13 | 0.75433559 | 0.143 | 0.088 | 8.91E-09 |
| Dbp | 4.82E-13 | 0.7308105 | 0.125 | 0.073 | 9.67E-09 |
| Ehmt2 | 5.72E-13 | 0.66606161 | 0.167 | 0.107 | 1.15E-08 |
| Exosc10 | 7.30E-13 | 0.86032303 | 0.103 | 0.058 | 1.46E-08 |
| Trip10 | 7.50E-13 | 0.75906312 | 0.133 | 0.08 | 1.50E-08 |
| Emsy | 8.21E-13 | 0.65808989 | 0.178 | 0.117 | 1.65E-08 |
| Lrrfip2 | 8.49E-13 | 0.61257169 | 0.216 | 0.149 | 1.70E-08 |
| Cyth1 | 8.50E-13 | 0.88346432 | 0.106 | 0.06 | 1.71E-08 |
| Irak3 | 8.56E-13 | 0.86920865 | 0.112 | 0.064 | 1.72E-08 |
| Stam | 9.82E-13 | 0.86248771 | 0.104 | 0.059 | 1.97E-08 |
| 0610010K14 | 9.96E-13 | 0.64953489 | 0.163 | 0.105 | 2.00E-08 |
| Megf8 | 1.09E-12 | 0.85160511 | 0.108 | 0.061 | 2.20E-08 |
| Tmem184c | 1.27E-12 | 0.75169034 | 0.125 | 0.075 | 2.55E-08 |
| Cnrip1 | 1.31E-12 | 0.73228524 | 0.132 | 0.08 | 2.64E-08 |
| Taz | 1.63E-12 | 0.79722397 | 0.112 | 0.065 | 3.27E-08 |
| Pex2 | 1.70E-12 | 0.73095019 | 0.136 | 0.084 | 3.41E-08 |
| Glyr1 | 1.87E-12 | 0.61710449 | 0.197 | 0.133 | 3.74E-08 |
| D5Ertd579e | 1.95E-12 | 0.69517529 | 0.161 | 0.104 | 3.90E-08 |
| Arsk | 1.98E-12 | 0.82506157 | 0.107 | 0.061 | 3.96E-08 |
| 3830406C13 | 2.11E-12 | 0.7485879 | 0.12 | 0.071 | 4.24E-08 |
| Ctnnbl1 | 2.26E-12 | 0.77984118 | 0.119 | 0.071 | 4.53E-08 |
| Ebag9 | 2.58E-12 | 0.83988692 | 0.112 | 0.065 | 5.17E-08 |
| Spire1 | 3.37E-12 | 0.71164145 | 0.143 | 0.089 | 6.77E-08 |
| Glod4 | 3.50E-12 | 0.63006846 | 0.17 | 0.112 | 7.03E-08 |
| Eif2s3x | 3.54E-12 | 0.67549149 | 0.153 | 0.098 | 7.11E-08 |
| Crnkl1 | 3.66E-12 | 0.80639221 | 0.112 | 0.065 | 7.35E-08 |
| Nav2 | 4.06E-12 | 0.71312188 | 0.138 | 0.085 | 8.14E-08 |
| Sun1 | 4.88E-12 | 0.73541473 | 0.14 | 0.088 | 9.80E-08 |
| Tm9sf4 | 4.90E-12 | 0.61942495 | 0.186 | 0.125 | 9.84E-08 |
| Egfr | 5.22E-12 | 0.63179469 | 0.16 | 0.103 | 1.05E-07 |
| Aldh7a1 | 5.72E-12 | 0.63688193 | 0.154 | 0.098 | 1.15E-07 |
| Hbb-bs | 5.79E-12 | 1.07218682 | 0.211 | 0.289 | 1.16E-07 |
| Zswim8 | 7.08E-12 | 0.65196463 | 0.15 | 0.096 | 1.42E-07 |
| Atxn7l3 | 7.10E-12 | 0.70099406 | 0.136 | 0.084 | 1.42E-07 |
| Abhd14b | 7.14E-12 | 0.68202493 | 0.133 | 0.082 | 1.43E-07 |
| Slc25a24 | 7.71E-12 | 0.69607574 | 0.131 | 0.08 | 1.55E-07 |
| Spats2 | 8.44E-12 | 0.67145606 | 0.149 | 0.095 | 1.69E-07 |
| Dram2 | 1.04E-11 | 0.86004227 | 0.108 | 0.063 | 2.09E-07 |
| Nek6 | 1.06E-11 | 0.7245816 | 0.127 | 0.078 | 2.13E-07 |
| Lypla1 | 1.06E-11 | 0.72209377 | 0.131 | 0.081 | 2.13E-07 |
| Hk1 | 1.27E-11 | 0.70573452 | 0.118 | 0.071 | 2.54E-07 |
| Sil1 | 1.32E-11 | 0.73769709 | 0.127 | 0.078 | 2.64E-07 |
| Rnpep | 1.45E-11 | 0.80247391 | 0.104 | 0.061 | 2.90E-07 |
| Ide | 1.64E-11 | 0.75596456 | 0.127 | 0.078 | 3.29E-07 |
| Gmps | 1.78E-11 | 0.63232044 | 0.171 | 0.114 | 3.58E-07 |
| Glb1 | 1.94E-11 | 0.76194163 | 0.111 | 0.066 | 3.88E-07 |
| Dnajc14 | 1.95E-11 | 0.75323877 | 0.108 | 0.064 | 3.91E-07 |
| Ate1 | 2.03E-11 | 0.70464965 | 0.12 | 0.073 | 4.08E-07 |
| Tfdp2 | 2.14E-11 | 0.73058635 | 0.142 | 0.091 | 4.29E-07 |
| Cox6b2 | 2.20E-11 | 0.80092394 | 0.132 | 0.082 | 4.42E-07 |
| Trmt1l | 2.34E-11 | 0.82240227 | 0.115 | 0.069 | 4.70E-07 |
| Soat1 | 3.23E-11 | 0.62829923 | 0.145 | 0.093 | 6.48E-07 |
| S1pr2 | 4.14E-11 | 0.70783034 | 0.12 | 0.074 | 8.31E-07 |
| Riok1 | 5.25E-11 | 0.71472331 | 0.121 | 0.074 | 1.05E-06 |
| Pcdh19 | 5.54E-11 | 0.88578728 | 0.123 | 0.076 | 1.11E-06 |
| Rcbtb2 | 5.61E-11 | 0.68657227 | 0.128 | 0.08 | 1.13E-06 |
| Inpp5a | 5.69E-11 | 0.60646209 | 0.149 | 0.097 | 1.14E-06 |
| Atp6v0a2 | 5.74E-11 | 0.66992851 | 0.14 | 0.089 | 1.15E-06 |
| Setd7 | 5.96E-11 | 0.65096289 | 0.152 | 0.1 | 1.20E-06 |
| Mettl16 | 6.05E-11 | 0.77865564 | 0.104 | 0.061 | 1.21E-06 |
| Trrap | 7.38E-11 | 0.6534502 | 0.14 | 0.09 | 1.48E-06 |
| Tmem63b | 8.54E-11 | 0.69927386 | 0.13 | 0.082 | 1.71E-06 |
| Sec24a | 8.95E-11 | 0.68411489 | 0.15 | 0.099 | 1.80E-06 |
| Adamtsl4 | 1.10E-10 | 0.76280154 | 0.11 | 0.067 | 2.21E-06 |
| Mfap3 | 1.23E-10 | 0.75259957 | 0.103 | 0.061 | 2.47E-06 |
| Cpne2 | 1.27E-10 | 0.61310355 | 0.15 | 0.098 | 2.55E-06 |
| Ncbp3 | 1.55E-10 | 0.70221987 | 0.119 | 0.074 | 3.11E-06 |
| Mob1b | 1.58E-10 | 0.64141354 | 0.136 | 0.088 | 3.17E-06 |
| Carm1 | 1.69E-10 | 0.75925672 | 0.103 | 0.062 | 3.39E-06 |
| Ubap2 | 2.20E-10 | 0.60131491 | 0.152 | 0.101 | 4.42E-06 |
| Prkaca | 2.35E-10 | 0.64481328 | 0.15 | 0.099 | 4.72E-06 |
| Tram2 | 2.40E-10 | 0.69628968 | 0.136 | 0.088 | 4.81E-06 |
| Ilf3 | 2.42E-10 | 0.61921063 | 0.152 | 0.101 | 4.86E-06 |
| Fam98b | 2.54E-10 | 0.74284867 | 0.101 | 0.06 | 5.10E-06 |
| Snrk | 3.03E-10 | 0.62341025 | 0.144 | 0.094 | 6.09E-06 |
| Plekhg2 | 3.11E-10 | 0.65493534 | 0.127 | 0.081 | 6.23E-06 |
| Nploc4 | 3.20E-10 | 0.72721425 | 0.102 | 0.061 | 6.42E-06 |
| Dcaf7 | 4.09E-10 | 0.72571223 | 0.112 | 0.069 | 8.21E-06 |
| Napg | 5.04E-10 | 0.7081954 | 0.122 | 0.077 | 1.01E-05 |
| Pgap2 | 5.63E-10 | 0.6470439 | 0.128 | 0.082 | 1.13E-05 |
| Pcgf3 | 6.11E-10 | 0.65682492 | 0.127 | 0.082 | 1.23E-05 |
| Fbxo9 | 6.92E-10 | 0.70903172 | 0.119 | 0.075 | 1.39E-05 |
| Taok2 | 7.67E-10 | 0.74931844 | 0.101 | 0.061 | 1.54E-05 |
| Mtmr6 | 1.03E-09 | 0.60709897 | 0.148 | 0.099 | 2.07E-05 |
| Ccdc28b | 1.28E-09 | 0.71377811 | 0.102 | 0.062 | 2.57E-05 |
| Dlat | 1.62E-09 | 0.60217821 | 0.142 | 0.095 | 3.26E-05 |
| Ctps | 1.70E-09 | 0.64382504 | 0.135 | 0.089 | 3.42E-05 |
| Gcnt2 | 1.83E-09 | 0.67327544 | 0.13 | 0.086 | 3.68E-05 |
| Myo9b | 1.92E-09 | 0.62096944 | 0.141 | 0.094 | 3.86E-05 |
| Bms1 | 1.93E-09 | 0.68956948 | 0.104 | 0.065 | 3.87E-05 |
| Cdc42se2 | 1.93E-09 | 0.66658247 | 0.123 | 0.079 | 3.88E-05 |
| Afg3l1 | 2.03E-09 | 0.62319119 | 0.127 | 0.082 | 4.07E-05 |
| Casp8ap2 | 2.05E-09 | 0.60178837 | 0.133 | 0.087 | 4.11E-05 |
| Tmem38b | 2.12E-09 | 0.70518794 | 0.105 | 0.065 | 4.25E-05 |
| Uben2a | 2.42E-09 | 0.60213712 | 0.126 | 0.082 | 4.85E-05 |
| Ap4s1 | 2.73E-09 | 0.65773314 | 0.13 | 0.086 | 5.47E-05 |
| Clcn5 | 3.67E-09 | 0.64937682 | 0.104 | 0.065 | 7.37E-05 |
| Tubgcp4 | 3.82E-09 | 0.6673173 | 0.107 | 0.067 | 7.66E-05 |
| Otud4 | 4.02E-09 | 0.65682492 | 0.112 | 0.071 | 8.07E-05 |
| Slc35a1 | 4.58E-09 | 0.69198576 | 0.106 | 0.067 | 9.20E-05 |
| Fam160b1 | 4.74E-09 | 0.62177194 | 0.11 | 0.07 | 9.51E-05 |
| Slc25a46 | 5.30E-09 | 0.77849476 | 0.105 | 0.066 | 0.00010629 |
| Acyp1 | 5.94E-09 | 0.67963955 | 0.102 | 0.064 | 0.00011913 |
| Psen1 | 6.16E-09 | 0.63495574 | 0.123 | 0.08 | 0.00012355 |
| Sympk | 6.19E-09 | 0.71757483 | 0.104 | 0.066 | 0.0001243 |
| Camk2g | 6.72E-09 | 0.60592703 | 0.14 | 0.095 | 0.00013493 |
| Gpr107 | 6.82E-09 | 0.62094039 | 0.129 | 0.086 | 0.00013676 |
| Ibtk | 7.00E-09 | 0.65490005 | 0.122 | 0.08 | 0.00014045 |
| Vta1 | 7.21E-09 | 0.68167691 | 0.132 | 0.089 | 0.00014465 |
| Ephb2 | 8.89E-09 | 0.75067322 | 0.104 | 0.066 | 0.0001784 |
| Galnt11 | 1.40E-08 | 0.66378887 | 0.105 | 0.067 | 0.00028023 |
| Zbtb8os | 1.54E-08 | 0.66687859 | 0.112 | 0.073 | 0.00030808 |
| Gpatch11 | 2.30E-08 | 0.62967713 | 0.11 | 0.071 | 0.00046129 |
| Mtmr2 | 2.45E-08 | 0.62337844 | 0.117 | 0.077 | 0.00049164 |
| Capn7 | 3.49E-08 | 0.66699673 | 0.106 | 0.069 | 0.0006998 |
| Mpp1 | 3.59E-08 | 0.62960345 | 0.107 | 0.069 | 0.0007213 |
| Myo18a | 5.86E-08 | 0.60751064 | 0.123 | 0.083 | 0.00117629 |
| Ankrd44 | 8.27E-08 | 0.62128081 | 0.113 | 0.075 | 0.00166019 |
| Zfp9 | 8.94E-08 | 0.61161835 | 0.109 | 0.072 | 0.00179441 |

| | | | | | |
|---|---|---|---|---|---|
| H2afy2 | 9.23E-08 | 0.62135335 | 0.104 | 0.068 | 0.00185143 |
| Pik3r3 | 1.04E-07 | 0.70313336 | 0.106 | 0.07 | 0.00207721 |
| Lrif1 | 1.37E-07 | 0.62873167 | 0.114 | 0.077 | 0.00274969 |
| Foxn2 | 1.85E-07 | 0.63643065 | 0.109 | 0.073 | 0.00370765 |
| Fbxo28 | 2.62E-07 | 0.60168337 | 0.104 | 0.069 | 0.00525642 |
| Cdk17 | 2.89E-07 | 0.6452454 | 0.108 | 0.072 | 0.00580025 |
| Col8a1 | 1.83E-06 | 0.81423313 | 0.296 | 0.247 | 0.03682212 |
