## Supplementary material for "The nail mesenchyme creates a regeneration-specific ligand environment that orchestrates mammalian regeneration versus fibrotic wound healing": Table_2

**Table 2. *Probes used for Xenium single-cell spatial transcriptomics* (Related to Figures 2-7 and Supplemental Figures 2-6).** Shown are the genes included in the custom Xenium panel 1, comprised of probes for 480 genes. Included are Ensembl IDs for all genes used, as well as the number of probes used for each gene.The number of probes assigned to each gene was optimized individually to achieve strong detection and adequate isoform coverage, while minimizing optical crowding. Probe selection was guided by the 10x Xenium panel designer. The genes highlighted in yellow represent bioactive ligands that are targeted by the Xenium panel and predictive models of the early regenerating digit tip.

| | Custom Xenium Panel 1 | |
|---|---|---|
| **Genes** | **Ensembl ID** | **# Probes** |
| Bdnf | ENSMUSG00000048482 | 8 |
| Bmp7 | ENSMUSG00000008999 | 8 |
| Cck | ENSMUSG00000032532 | 5 |
| Edn3 | ENSMUSG00000027524 | 8 |
| Ednra | ENSMUSG00000031616 | 8 |
| Erbb3 | ENSMUSG00000018166 | 8 |
| Fgf18 | ENSMUSG00000057967 | 8 |
| Fzd4 | ENSMUSG00000049791 | 8 |
| Gdf6 | ENSMUSG00000051279 | 8 |
| Igf2 | ENSMUSG00000048583 | 8 |
| Inhbb | ENSMUSG00000037035 | 8 |
| Lgr5 | ENSMUSG00000020140 | 8 |
| Lgr6 | ENSMUSG00000042793 | 8 |
| Ngfr | ENSMUSG00000000120 | 8 |
| Nrg2 | ENSMUSG00000060275 | 8 |
| Ntrk2 | ENSMUSG00000055254 | 8 |
| Pth1r | ENSMUSG00000032492 | 8 |
| Pthlh | ENSMUSG00000048776 | 7 |
| Rspo1 | ENSMUSG00000028871 | 6 |
| Rspo4 | ENSMUSG00000032852 | 8 |
| Sema3a | ENSMUSG00000028883 | 8 |
| Slit2 | ENSMUSG00000031558 | 8 |
| Wnt11 | ENSMUSG00000015957 | 8 |
| Lmx1b | ENSMUSG00000038765 | 8 |
| Pappa2 | ENSMUSG00000073530 | 8 |
| Cxcl12 | ENSMUSG00000061353 | 8 |
| Sp9 | ENSMUSG00000068859 | 8 |
| Krt5 | ENSMUSG00000061527 | 8 |
| Krt14 | ENSMUSG00000045545 | 8 |
| Krt10 | ENSMUSG00000019761 | 6 |
| Krt17 | ENSMUSG00000035557 | 7 |
| Sox2 | ENSMUSG00000074637 | 8 |
| Sox10 | ENSMUSG00000033006 | 8 |
| Plp1 | ENSMUSG00000031425 | 8 |
| Lyve1 | ENSMUSG00000030787 | 8 |
| Flt4 | ENSMUSG00000020357 | 8 |
| Prox1 | ENSMUSG00000010175 | 8 |
| Pdpn | ENSMUSG00000028583 | 8 |
| Pecam1 | ENSMUSG00000020717 | 8 |
| Cdh5 | ENSMUSG00000031871 | 8 |
| Tie1 | ENSMUSG00000033191 | 8 |
| Tek | ENSMUSG00000006386 | 8 |
| Ly6a | ENSMUSG00000075602 | 7 |
| Prrx2 | ENSMUSG00000039476 | 5 |
| Pdgfra | ENSMUSG00000029231 | 8 |
| Hic1 | ENSMUSG00000043099 | 8 |
| Msx1 | ENSMUSG00000048450 | 4 |
| Mcam | ENSMUSG00000032135 | 8 |
| Rgs5 | ENSMUSG00000026678 | 6 |
| Myh11 | ENSMUSG00000018830 | 8 |
| Kcnj8 | ENSMUSG00000030247 | 8 |
| Pdgfrb | ENSMUSG00000024620 | 8 |
| Mki67 | ENSMUSG00000031004 | 8 |
| Pcna | ENSMUSG00000027342 | 8 |
| Alpl | ENSMUSG00000028766 | 8 |
| Col1a1 | ENSMUSG00000001506 | 3 |
| Acan | ENSMUSG00000030607 | 8 |
| Spp1 | ENSMUSG00000029304 | 8 |
| Sp7 | ENSMUSG00000060284 | 8 |
| Chad | ENSMUSG00000039084 | 8 |
| Dmp1 | ENSMUSG00000029307 | 8 |
| Tnfsf11 | ENSMUSG00000022015 | 8 |
| Tnfrsf11a | ENSMUSG00000026321 | 8 |
| Ctsk | ENSMUSG00000028111 | 8 |
| Mmp9 | ENSMUSG00000017737 | 8 |
| Acp5 | ENSMUSG00000001348 | 8 |
| Nfatc1 | ENSMUSG00000033016 | 8 |
| Ptprc | ENSMUSG00000026395 | 8 |
| Cd3e | ENSMUSG00000032093 | 8 |
| Cd3g | ENSMUSG00000002033 | 8 |
| Cxcr6 | ENSMUSG00000048521 | 8 |
| Ly6g | ENSMUSG00000022582 | 8 |
| Cd68 | ENSMUSG00000018774 | 8 |
| Cd14 | ENSMUSG00000051439 | 8 |
| Aif1 | ENSMUSG00000024397 | 7 |
| Itgam | ENSMUSG00000030786 | 8 |
| Lyz2 | ENSMUSG00000069516 | 4 |
| Tnf | ENSMUSG00000024401 | 8 |
| Ptgs2 | ENSMUSG00000032487 | 8 |
| Nos2 | ENSMUSG00000020826 | 8 |
| Cxcl2 | ENSMUSG00000058427 | 8 |
| Tlr2 | ENSMUSG00000027995 | 8 |
| Fcgr3 | ENSMUSG00000059498 | 8 |
| Mrc1 | ENSMUSG00000026712 | 8 |
| Arg1 | ENSMUSG00000019987 | 8 |
| Pparg | ENSMUSG00000000440 | 8 |
| Chil3 | ENSMUSG00000040809 | 8 |
| Retnla | ENSMUSG00000061100 | 7 |
| Il10 | ENSMUSG00000016529 | 8 |
| C1qb | ENSMUSG00000036905 | 6 |
| Cndp2 | ENSMUSG00000024644 | 8 |
| Csf1r | ENSMUSG00000024621 | 8 |
| F13a1 | ENSMUSG00000039109 | 8 |
| Fcgr2b | ENSMUSG00000026656 | 8 |
| Ms4a6d | ENSMUSG00000024679 | 8 |
| Ms4a7 | ENSMUSG00000024672 | 8 |
| Msr1 | ENSMUSG00000025044 | 8 |
| Pf4 | ENSMUSG00000029373 | 5 |
| Trem2 | ENSMUSG00000023992 | 3 |
| Arg2 | ENSMUSG00000021125 | 8 |
| Cd274 | ENSMUSG00000016496 | 8 |
| Csf3r | ENSMUSG00000028859 | 8 |
| Entpd1 | ENSMUSG00000048120 | 8 |
| Hdc | ENSMUSG00000027360 | 8 |
| Lcn2 | ENSMUSG00000026822 | 7 |
| Plk3 | ENSMUSG00000028680 | 8 |
| Trem1 | ENSMUSG00000042265 | 8 |
| Cxcr2 | ENSMUSG00000026180 | 8 |
| Fpr2 | ENSMUSG00000052270 | 8 |
| Upp1 | ENSMUSG00000020407 | 8 |
| Wfdc21 | ENSMUSG00000051748 | 3 |
| Cd83 | ENSMUSG00000015396 | 8 |
| Etv3 | ENSMUSG00000003382 | 8 |
| Ramp3 | ENSMUSG00000041046 | 8 |
| Tnip3 | ENSMUSG00000044162 | 8 |
| Ccl3 | ENSMUSG00000000982 | 7 |
| Ccl4 | ENSMUSG00000018930 | 4 |
| Ccl9 | ENSMUSG00000019122 | 8 |
| Il1a | ENSMUSG00000027399 | 8 |
| Il1b | ENSMUSG00000027398 | 8 |
| Osm | ENSMUSG00000058755 | 8 |
| Rtn4 | ENSMUSG00000020458 | 8 |
| Tnfsf12 | ENSMUSG00000097328 | 8 |
| Vegfa | ENSMUSG00000023951 | 8 |
| Tnfrsf1a | ENSMUSG00000030341 | 8 |
| Tmem119 | ENSMUSG00000054675 | 8 |
| Clec11a | ENSMUSG00000004473 | 8 |
| Comp | ENSMUSG00000031849 | 8 |
| Cpz | ENSMUSG00000036596 | 8 |
| H19 | ENSMUSG00000000031 | 8 |
| Matn4 | ENSMUSG00000016995 | 7 |
| Sfrp2 | ENSMUSG00000027996 | 5 |
| Sfrp4 | ENSMUSG00000021319 | 8 |
| Col9a2 | ENSMUSG00000028626 | 7 |
| Lama1 | ENSMUSG00000032796 | 8 |
| Slc22a6 | ENSMUSG00000024650 | 8 |
| Aldh1a2 | ENSMUSG00000013584 | 8 |
| Bmp6 | ENSMUSG00000039004 | 8 |
| Clcn2 | ENSMUSG00000022843 | 8 |
| Coch | ENSMUSG00000020953 | 8 |
| Crabp2 | ENSMUSG00000004885 | 8 |
| Gja1 | ENSMUSG00000050953 | 8 |
| Gjb2 | ENSMUSG00000046352 | 8 |
| Gjb6 | ENSMUSG00000040055 | 8 |
| Kcnj13 | ENSMUSG00000079436 | 8 |
| Osr1 | ENSMUSG00000048387 | 8 |
| Slc6a13 | ENSMUSG00000030108 | 8 |
| Wnt6 | ENSMUSG00000033227 | 8 |
| Adamts5 | ENSMUSG00000022894 | 8 |
| Dpt | ENSMUSG00000026574 | 8 |
| Pcolce2 | ENSMUSG00000015354 | 8 |
| Dpp4 | ENSMUSG00000035000 | 8 |
| Cxcl13 | ENSMUSG00000023078 | 8 |
| Ly6c1 | ENSMUSG00000079018 | 7 |
| Bmp4 | ENSMUSG00000021835 | 8 |
| Gpnmb | ENSMUSG00000029816 | 8 |
| Fbln7 | ENSMUSG00000027386 | 8 |
| Fmod | ENSMUSG00000041559 | 8 |
| Msln | ENSMUSG00000063011 | 8 |
| Itga6 | ENSMUSG00000027111 | 8 |
| Itgb4 | ENSMUSG00000020758 | 8 |
| Krt19 | ENSMUSG00000020911 | 8 |
| Sfrp5 | ENSMUSG00000018822 | 6 |
| Itga7 | ENSMUSG00000025348 | 8 |
| Sox8 | ENSMUSG00000024176 | 8 |
| Etv1 | ENSMUSG00000004151 | 8 |
| Sox9 | ENSMUSG00000000567 | 8 |
| Col11a1 | ENSMUSG00000027966 | 8 |
| Col26a1 | ENSMUSG00000004415 | 8 |
| A2m | ENSMUSG00000030111 | 8 |
| Kng2 | ENSMUSG00000060459 | 8 |
| Aqp4 | ENSMUSG00000024411 | 8 |
| Abhd3 | ENSMUSG00000002475 | 8 |
| Arxes2 | ENSMUSG00000048040 | 8 |
| Agt | ENSMUSG00000031980 | 8 |
| Cadm2 | ENSMUSG00000064115 | 8 |
| Car2 | ENSMUSG00000027562 | 8 |
| Cd38 | ENSMUSG00000029084 | 8 |
| Cryab | ENSMUSG00000032060 | 7 |
| Cxcl14 | ENSMUSG00000021508 | 8 |
| Gfap | ENSMUSG00000020932 | 8 |
| Slc1a3 | ENSMUSG00000005360 | 8 |
| Cdh1 | ENSMUSG00000000303 | 8 |
| Myoc | ENSMUSG00000026697 | 8 |
| Pdgfc | ENSMUSG00000028019 | 8 |
| Ltbp2 | ENSMUSG00000002020 | 8 |
| Fbxo17 | ENSMUSG00000030598 | 8 |
| Pycr1 | ENSMUSG00000025140 | 8 |
| Arsi | ENSMUSG00000036412 | 8 |
| Matn3 | ENSMUSG00000020583 | 8 |
| Cdkn2b | ENSMUSG00000073802 | 7 |
| Zfp354c | ENSMUSG00000044807 | 8 |
| Hebp2 | ENSMUSG00000019853 | 8 |
| Prss35 | ENSMUSG00000033491 | 8 |
| Wnt16 | ENSMUSG00000029671 | 8 |
| Camk4 | ENSMUSG00000038128 | 8 |
| Serinc2 | ENSMUSG00000023232 | 8 |
| Prkg2 | ENSMUSG00000029334 | 8 |
| Ttll5 | ENSMUSG00000012609 | 8 |
| Igdcc4 | ENSMUSG00000032816 | 8 |
| Cpa6 | ENSMUSG00000042501 | 8 |
| Mpped2 | ENSMUSG00000016386 | 8 |
| Bcl11b | ENSMUSG00000048251 | 8 |
| Nell2 | ENSMUSG00000022454 | 8 |
| Sec16b | ENSMUSG00000026589 | 8 |
| Tmeff1 | ENSMUSG00000028347 | 8 |
| Bnc2 | ENSMUSG00000028487 | 8 |
| Fbxo36 | ENSMUSG00000073633 | 8 |
| Dync1i1 | ENSMUSG00000029757 | 8 |
| Kcnj15 | ENSMUSG00000062609 | 8 |
| Capn6 | ENSMUSG00000067276 | 8 |
| Cad | ENSMUSG00000013629 | 8 |
| Fhod3 | ENSMUSG00000034295 | 8 |
| Ccn5 | ENSMUSG00000027656 | 8 |
| Efna4 | ENSMUSG00000028040 | 6 |
| Npr3 | ENSMUSG00000022206 | 8 |
| Pamr1 | ENSMUSG00000027188 | 8 |
| Col24a1 | ENSMUSG00000028197 | 8 |
| Serp2 | ENSMUSG00000052584 | 4 |
| Stk26 | ENSMUSG00000031112 | 8 |
| Pak3 | ENSMUSG00000031284 | 8 |
| Adamtsl3 | ENSMUSG00000070469 | 8 |
| Aff3 | ENSMUSG00000037138 | 8 |
| Brinp3 | ENSMUSG00000035131 | 8 |
| Rcor2 | ENSMUSG00000024968 | 8 |
| Arxes1 | ENSMUSG00000048355 | 8 |
| Asphd2 | ENSMUSG00000029348 | 8 |
| Tnfaip8l3 | ENSMUSG00000074345 | 8 |
| Hs3st3b1 | ENSMUSG00000070407 | 8 |
| Unc5c | ENSMUSG00000059921 | 8 |
| Cxadr | ENSMUSG00000022865 | 8 |
| Gjb5 | ENSMUSG00000042357 | 8 |
| Gpr27 | ENSMUSG00000072875 | 8 |
| Zfp185 | ENSMUSG00000031351 | 8 |
| Bmp3 | ENSMUSG00000029335 | 8 |
| Plekhg4 | ENSMUSG00000014782 | 8 |
| Farp2 | ENSMUSG00000034066 | 8 |
| Col9a1 | ENSMUSG00000026147 | 8 |
| Colq | ENSMUSG00000057606 | 8 |
| Gpr173 | ENSMUSG00000056679 | 8 |
| Gzmc | ENSMUSG00000079186 | 8 |
| Aqp5 | ENSMUSG00000044217 | 5 |
| Galnt3 | ENSMUSG00000026994 | 8 |
| Fat3 | ENSMUSG00000074505 | 8 |
| Ucn2 | ENSMUSG00000049699 | 7 |
| Bdkrb1 | ENSMUSG00000041347 | 8 |
| Rab27b | ENSMUSG00000024511 | 8 |
| Stc2 | ENSMUSG00000020303 | 8 |
| Pank1 | ENSMUSG00000033610 | 8 |
| Hp | ENSMUSG00000031722 | 6 |
| Angptl1 | ENSMUSG00000033544 | 8 |
| Aqp2 | ENSMUSG00000023013 | 4 |
| Ccl8 | ENSMUSG00000009185 | 5 |
| Cd34 | ENSMUSG00000016494 | 8 |
| Cilp | ENSMUSG00000042254 | 8 |
| Clec3b | ENSMUSG00000025784 | 2 |
| Galnt16 | ENSMUSG00000021130 | 8 |
| Mmp11 | ENSMUSG00000000901 | 8 |
| Sgsm2 | ENSMUSG00000038351 | 8 |
| Shox2 | ENSMUSG00000027833 | 8 |
| Sulf2 | ENSMUSG00000006800 | 8 |
| Thbs4 | ENSMUSG00000021702 | 8 |
| Tgfb2 | ENSMUSG00000039239 | 8 |
| Tgfb3 | ENSMUSG00000021253 | 8 |
| Bmp1 | ENSMUSG00000022098 | 8 |
| Bmp2 | ENSMUSG00000027358 | 8 |
| Bmp5 | ENSMUSG00000032179 | 8 |
| Gdf11 | ENSMUSG00000025352 | 8 |
| Smad1 | ENSMUSG00000031681 | 8 |
| Smad3 | ENSMUSG00000032402 | 8 |
| Smad4 | ENSMUSG00000024515 | 8 |
| Smad6 | ENSMUSG00000036867 | 6 |
| Smad9 | ENSMUSG00000027796 | 8 |
| Mapk10 | ENSMUSG00000046709 | 8 |
| Mapk12 | ENSMUSG00000022610 | 8 |
| Mapk13 | ENSMUSG00000004864 | 8 |
| Map3k7 | ENSMUSG00000028284 | 8 |
| Fbn2 | ENSMUSG00000024598 | 8 |
| Lrrc17 | ENSMUSG00000039883 | 8 |
| Adgrg6 | ENSMUSG00000039116 | 8 |
| Lhx9 | ENSMUSG00000019230 | 8 |
| Galnt17 | ENSMUSG00000034040 | 8 |
| Cyp2f2 | ENSMUSG00000052974 | 8 |
| Dio3 | ENSMUSG00000075707 | 8 |
| Emx2 | ENSMUSG00000043969 | 8 |
| Dlx5 | ENSMUSG00000029755 | 7 |
| Gpha2 | ENSMUSG00000024784 | 6 |
| Steap1 | ENSMUSG00000015652 | 8 |
| Pcsk2 | ENSMUSG00000027419 | 8 |
| Gpm6a | ENSMUSG00000031517 | 8 |
| Penk | ENSMUSG00000045573 | 8 |
| Pitx1 | ENSMUSG00000021506 | 2 |
| Corin | ENSMUSG00000005220 | 8 |
| Spock3 | ENSMUSG00000054162 | 8 |
| Snhg11 | ENSMUSG00000044349 | 8 |
| Alx4 | ENSMUSG00000040310 | 8 |
| Tox3 | ENSMUSG00000043668 | 8 |
| Zcchc18 | ENSMUSG00000031428 | 8 |
| Hs3st3a1 | ENSMUSG00000047759 | 8 |
| Ntn5 | ENSMUSG00000070564 | 7 |
| Plxdc1 | ENSMUSG00000017417 | 8 |
| Pcdh10 | ENSMUSG00000049100 | 8 |
| Adam33 | ENSMUSG00000027318 | 8 |
| Macb | ENSMUSG00000040147 | 8 |
| Tmem132c | ENSMUSG00000034324 | 8 |
| Gna14 | ENSMUSG00000024697 | 8 |
| Tbx4 | ENSMUSG00000000094 | 8 |
| Ccdc74a | ENSMUSG00000041617 | 8 |
| Aff2 | ENSMUSG00000031189 | 8 |
| Zic2 | ENSMUSG00000061524 | 7 |
| Chd5 | ENSMUSG00000005045 | 8 |
| Emx2os | ENSMUSG00000087095 | 8 |
| Ackr4 | ENSMUSG00000079355 | 8 |
| Lamc3 | ENSMUSG00000026840 | 8 |
| Scube1 | ENSMUSG00000016763 | 8 |
| Kcp | ENSMUSG00000059022 | 8 |
| Chodl | ENSMUSG00000022860 | 8 |
| Kera | ENSMUSG00000019932 | 8 |
| Foxf2 | ENSMUSG00000038402 | 8 |
| Vipr2 | ENSMUSG00000011171 | 8 |
| Adh1 | ENSMUSG00000074207 | 8 |
| Nlgn1 | ENSMUSG00000063887 | 8 |
| Klhl32 | ENSMUSG00000040387 | 8 |
| Ccdc136 | ENSMUSG00000029769 | 8 |
| Cntfr | ENSMUSG00000028444 | 8 |
| Rab40b | ENSMUSG00000025170 | 8 |
| Negr1 | ENSMUSG00000040037 | 8 |
| Hs3st6 | ENSMUSG00000039628 | 3 |
| Tbxa2r | ENSMUSG00000034881 | 6 |
| Dbx2 | ENSMUSG00000045608 | 8 |
| Gfra2 | ENSMUSG00000022103 | 8 |
| Rab30 | ENSMUSG00000030643 | 8 |
| Dipk1b | ENSMUSG00000036186 | 7 |
| Hmgcs2 | ENSMUSG00000027875 | 8 |
| Fez1 | ENSMUSG00000032118 | 8 |
| Rgs17 | ENSMUSG00000019775 | 8 |
| Csmd3 | ENSMUSG00000022311 | 8 |
| Dpep1 | ENSMUSG00000019278 | 8 |
| Tle2 | ENSMUSG00000034771 | 8 |
| Sema3b | ENSMUSG00000057969 | 8 |
| Zic5 | ENSMUSG00000041703 | 8 |
| Focad | ENSMUSG00000038368 | 8 |
| Sall1 | ENSMUSG00000031665 | 8 |
| Plppr3 | ENSMUSG00000035835 | 7 |
| Hoxd13 | ENSMUSG00000001819 | 8 |
| Gaa | ENSMUSG00000025579 | 8 |
| Galnt9 | ENSMUSG00000033316 | 8 |
| Ubtd2 | ENSMUSG00000044949 | 8 |
| Tmem26 | ENSMUSG00000060044 | 8 |
| Il17d | ENSMUSG00000050222 | 5 |
| Mrc2 | ENSMUSG00000020695 | 8 |
| Arsj | ENSMUSG00000046561 | 8 |
| Hoxa11 | ENSMUSG00000038210 | 8 |
| Sh3rf3 | ENSMUSG00000037990 | 8 |
| Omd | ENSMUSG00000048368 | 8 |
| Cpxm2 | ENSMUSG00000030862 | 8 |
| Tnn | ENSMUSG00000026725 | 8 |
| Ptpru | ENSMUSG00000028909 | 8 |
| Hmgcll1 | ENSMUSG00000007908 | 8 |
| Ror2 | ENSMUSG00000021464 | 8 |
| Ptgfr | ENSMUSG00000028036 | 8 |
| Tspan11 | ENSMUSG00000030351 | 8 |
| Has1 | ENSMUSG00000003665 | 8 |
| Wnt5b | ENSMUSG00000030170 | 8 |
| Mmp23 | ENSMUSG00000029061 | 6 |
| Hoxa13 | ENSMUSG00000038203 | 8 |
| Ssc5d | ENSMUSG00000035279 | 8 |
| Pcdh18 | ENSMUSG00000037892 | 8 |
| C1qtnf2 | ENSMUSG00000046491 | 6 |
| Tcea3 | ENSMUSG00000001604 | 8 |
| Pcdh17 | ENSMUSG00000035566 | 8 |
| Cys1 | ENSMUSG00000062563 | 8 |
| Epha5 | ENSMUSG00000029245 | 8 |
| Ntrk3 | ENSMUSG00000059146 | 8 |
| Mc5r | ENSMUSG00000007480 | 8 |
| Ereg | ENSMUSG00000029377 | 8 |
| Lif | ENSMUSG00000034394 | 8 |
| Rspo2 | ENSMUSG00000051920 | 8 |
| Rspo3 | ENSMUSG00000019880 | 8 |
| Sema3d | ENSMUSG00000040254 | 8 |
| Col2a1 | ENSMUSG00000022483 | 8 |
| Cxcr4 | ENSMUSG00000045382 | 8 |
| Ppp1r1b | ENSMUSG00000061718 | 8 |
| Col11a2 | ENSMUSG00000024330 | 8 |
| Msmp | ENSMUSG00000078719 | 5 |
| Col9a3 | ENSMUSG00000027570 | 8 |
| Smpd3 | ENSMUSG00000031906 | 8 |
| Cenpc1 | ENSMUSG00000029253 | 8 |
| Cnmd | ENSMUSG00000022025 | 8 |
| Fzd9 | ENSMUSG00000049551 | 8 |
| Chrdl1 | ENSMUSG00000031283 | 8 |
| Mmp3 | ENSMUSG00000043613 | 8 |
| Gldn | ENSMUSG00000046167 | 8 |
| Itgbl1 | ENSMUSG00000032925 | 8 |
| Crabp1 | ENSMUSG00000032291 | 6 |
| Mesp1 | ENSMUSG00000022887 | 8 |
| Ostn | ENSMUSG00000052276 | 8 |
| Ccnb1ip1 | ENSMUSG00000071470 | 8 |
| Cybrd1 | ENSMUSG00000027015 | 8 |
| Gdnf | ENSMUSG00000022144 | 8 |
| Il1rl1 | ENSMUSG00000026069 | 8 |
| Tnmd | ENSMUSG00000031250 | 8 |
| Mkx | ENSMUSG00000061013 | 8 |
| Ndnf | ENSMUSG00000049001 | 8 |
| Scx | ENSMUSG00000034161 | 3 |
| Nova1 | ENSMUSG00000021047 | 8 |
| Adgrd1 | ENSMUSG00000044017 | 8 |
| Il33 | ENSMUSG00000024810 | 8 |
| Lrrn4cl | ENSMUSG00000071656 | 8 |
| Osr2 | ENSMUSG00000022330 | 8 |
| Gzmd | ENSMUSG00000059256 | 8 |
| Mcpt8 | ENSMUSG00000022157 | 8 |
| Gzme | ENSMUSG00000022156 | 8 |
| Grem1 | ENSMUSG00000074934 | 8 |
| Fam180a | ENSMUSG00000047420 | 8 |
| Car3 | ENSMUSG00000027559 | 8 |
| Slc36a2 | ENSMUSG00000020264 | 8 |
| Cdh15 | ENSMUSG00000031962 | 8 |
| Phex | ENSMUSG00000057457 | 8 |
| Panx3 | ENSMUSG00000011118 | 8 |

| | | |
|---|---|---|
| *Wnt10b* | ENSMUSG00000022996 | 8 |
| *Bmp8a* | ENSMUSG00000032726 | 8 |
| *Ube2ql1* | ENSMUSG00000052981 | 8 |
| *Bglap3* | ENSMUSG00000074489 | 3 |
| *Six4* | ENSMUSG00000034460 | 8 |
| *Bglap2* | ENSMUSG00000074486 | 2 |
| *Ucma* | ENSMUSG00000026668 | 8 |
| *Ihh* | ENSMUSG00000006538 | 8 |
| *Slc38a3* | ENSMUSG00000010064 | 8 |
| *Dkk1* | ENSMUSG00000024868 | 8 |
| *Mia* | ENSMUSG00000089661 | 3 |
| *Gm28153* | ENSMUSG00000099895 | 6 |
| *Foxa3* | ENSMUSG00000040891 | 8 |
| *Sim2* | ENSMUSG00000062713 | 8 |
| *Slc13a5* | ENSMUSG00000020805 | 8 |
| *Foxd1* | ENSMUSG00000078302 | 8 |
| *Shisa3* | ENSMUSG00000050010 | 8 |
| *Gdf10* | ENSMUSG00000021943 | 8 |
| *Igf1r* | ENSMUSG00000005533 | 8 |
| *Fgf2* | ENSMUSG00000037225 | 8 |
| *Ror1* | ENSMUSG00000035305 | 8 |
| *Fgfr3* | ENSMUSG00000054252 | 8 |
| *Wnt5a* | ENSMUSG00000021994 | 8 |
| *Tjp1* | ENSMUSG00000030516 | 8 |
| *Tjp2* | ENSMUSG00000024812 | 8 |
| *S100a8* | ENSMUSG00000056054 | 4 |
| *Retnlg* | ENSMUSG00000022651 | 5 |
| *Lrg1* | ENSMUSG00000037095 | 5 |
| *Il23a* | ENSMUSG00000025383 | 8 |
| *Cxcl3* | ENSMUSG00000029379 | 8 |
| *Slpi* | ENSMUSG00000017002 | 6 |
| *Bglap* | ENSMUSG00000074483 | 3 |
| *Nrg3* | ENSMUSG00000041014 | 8 |
| *Cd300lf* | ENSMUSG00000047798 | 8 |
| *C1qa* | ENSMUSG00000036887 | 7 |
| *Gm26771* | ENSMUSG00000097250 | 8 |
| *Capg4* | ENSMUSG00000032911 | 8 |
| *Mepe* | ENSMUSG00000053863 | 8 |
| *Col10a1* | ENSMUSG00000039462 | 8 |
| *Shh* | ENSMUSG00000002633 | 8 |
| *Mc4r* | ENSMUSG00000047259 | 8 |
| *Irx4* | ENSMUSG00000021604 | 8 |
| *Irx1* | ENSMUSG00000060969 | 8 |
| *Sox21* | ENSMUSG00000061517 | 8 |
| *Epgn* | ENSMUSG00000035020 | 8 |
| *Fam110c* | ENSMUSG00000036136 | 8 |
| *Edn2* | ENSMUSG00000028635 | 8 |
| *Lypd5* | ENSMUSG00000030484 | 5 |
| *Sdr9c7* | ENSMUSG00000040127 | 8 |
| *Ocstamp* | ENSMUSG00000027670 | 8 |
| *Oscar* | ENSMUSG00000054594 | 8 |
| *Cyp1a1* | ENSMUSG00000032315 | 8 |
| *Chp2* | ENSMUSG00000030865 | 8 |
| *Gkn3* | ENSMUSG00000030048 | 8 |
| *Pcp4* | ENSMUSG00000090223 | 3 |
| *P2rx1* | ENSMUSG00000020787 | 8 |
| *Dlk1* | ENSMUSG00000040856 | 8 |
| *Kcna1* | ENSMUSG00000047976 | 8 |
| *Fgf10* | ENSMUSG00000021732 | 8 |
| *Cfh* | ENSMUSG00000026365 | 8 |

**Table 2. *Probes used for Xenium single-cell spatial transcriptomics* (Related to Supplemental Figure 5).** Shown are the genes included in the custom Xenium panel 2, comprised of probes for 473 genes. Included are Ensembl IDs for all genes used, as well as the number of probes used for each gene. The number of probes assigned to each gene was optimized individually to achieve strong detection and adequate isoform coverage, while minimizing optical crowding. Probe selection was guided by the 10x Xenium panel designer. The genes highlighted in yellow represent bioactive ligands that are targeted by the Xenium panel and predictive models of the early regenerating digit tip.

| Custom Xenium Panel 2 | | |
|---|---|---|
| **Genes** | **Ensembl ID** | **# Probes** |
| *Gm26771* | ENSMUSG00000097250 | 5 |
| *Msx1* | ENSMUSG00000048450 | 6 |
| *S100a4* | ENSMUSG00000001020 | 4 |
| *Tnmd* | ENSMUSG00000031250 | 6 |
| *Vim* | ENSMUSG00000026728 | 4 |
| *Sfrp2* | ENSMUSG00000027996 | 5 |
| *Ccl4* | ENSMUSG00000018930 | 3 |
| *Cd14* | ENSMUSG00000051439 | 5 |
| *Krt10* | ENSMUSG00000019761 | 6 |
| *Rgs5* | ENSMUSG00000026678 | 6 |
| *Slpi* | ENSMUSG00000017002 | 6 |
| *Bdnf* | ENSMUSG00000048482 | 8 |
| *Bmp7* | ENSMUSG00000008999 | 8 |
| *Cck* | ENSMUSG00000032532 | 8 |
| *Edn3* | ENSMUSG00000027524 | 8 |
| *Ednra* | ENSMUSG00000031616 | 8 |
| *Erbb3* | ENSMUSG00000018166 | 8 |
| *Fgf18* | ENSMUSG00000057967 | 8 |
| *Fzd4* | ENSMUSG00000049791 | 8 |
| *Gdf6* | ENSMUSG00000051279 | 8 |
| *Igf2* | ENSMUSG00000048583 | 8 |
| *Inhbb* | ENSMUSG00000037035 | 8 |
| *Lgr5* | ENSMUSG00000020140 | 8 |
| *Lgr6* | ENSMUSG00000042793 | 8 |
| *Ngfr* | ENSMUSG00000000120 | 8 |
| *Nrg2* | ENSMUSG00000060275 | 8 |
| *Ntrk2* | ENSMUSG00000055254 | 8 |
| *Pth1r* | ENSMUSG00000032492 | 8 |
| *Pthlh* | ENSMUSG00000048776 | 8 |
| *Rspo1* | ENSMUSG00000028871 | 8 |
| *Rspo4* | ENSMUSG00000032852 | 8 |
| *Sema3a* | ENSMUSG00000028883 | 8 |
| *Slit2* | ENSMUSG00000031558 | 8 |
| *Wnt11* | ENSMUSG00000015957 | 8 |
| *Lmx1b* | ENSMUSG00000038765 | 8 |
| *Pappa2* | ENSMUSG00000073530 | 8 |
| *Cxcl12* | ENSMUSG00000061353 | 8 |
| *Sp9* | ENSMUSG00000068859 | 8 |
| *Krt5* | ENSMUSG00000061527 | 8 |
| *Krt14* | ENSMUSG00000045545 | 8 |
| *Krt17* | ENSMUSG00000035557 | 8 |
| *Sox2* | ENSMUSG00000074637 | 8 |
| *Sox10* | ENSMUSG00000033006 | 8 |
| *Plp1* | ENSMUSG00000031425 | 8 |
| *Lyve1* | ENSMUSG00000030787 | 8 |
| *Flt4* | ENSMUSG00000020357 | 8 |
| *Prox1* | ENSMUSG00000010175 | 8 |
| *Pdpn* | ENSMUSG00000028583 | 8 |
| *Pecam1* | ENSMUSG00000020717 | 8 |
| *Cdh5* | ENSMUSG00000031871 | 8 |
| *Tie1* | ENSMUSG00000033191 | 8 |
| *Tek* | ENSMUSG00000006386 | 8 |
| *Ly6a* | ENSMUSG00000075602 | 7 |
| *Prrx2* | ENSMUSG00000039476 | 8 |
| *Pdgfra* | ENSMUSG00000029231 | 8 |
| *Hic1* | ENSMUSG00000043099 | 8 |
| *Mcam* | ENSMUSG00000032135 | 8 |
| *Myh11* | ENSMUSG00000018830 | 8 |
| *Kcnj8* | ENSMUSG00000030247 | 8 |
| *Pdgfrb* | ENSMUSG00000024620 | 8 |
| *Mki67* | ENSMUSG00000031004 | 8 |
| *Pcna* | ENSMUSG00000027342 | 8 |
| *Alpl* | ENSMUSG00000028766 | 8 |
| *Acan* | ENSMUSG00000030607 | 8 |
| *Spp1* | ENSMUSG00000029304 | 8 |
| *Sp7* | ENSMUSG00000060284 | 8 |
| *Chad* | ENSMUSG00000039084 | 8 |
| *Dmp1* | ENSMUSG00000029307 | 8 |
| *Tnfsf11* | ENSMUSG00000022015 | 8 |
| *Tnfrsf11a* | ENSMUSG00000026321 | 8 |
| *Ctsk* | ENSMUSG00000028111 | 8 |
| *Mmp9* | ENSMUSG00000017737 | 8 |
| *Acp5* | ENSMUSG00000001348 | 8 |
| *Nfatc1* | ENSMUSG00000033016 | 8 |
| *Ptprc* | ENSMUSG00000026395 | 8 |
| *Cd3e* | ENSMUSG00000032093 | 8 |
| *Cd3g* | ENSMUSG00000002033 | 8 |
| *Cxcr6* | ENSMUSG00000048521 | 8 |
| *Cd68* | ENSMUSG00000018774 | 8 |
| *Aif1* | ENSMUSG00000024397 | 8 |
| *Itgam* | ENSMUSG00000030786 | 8 |
| *Lyz2* | ENSMUSG00000069516 | 8 |
| *Tnf* | ENSMUSG00000024401 | 8 |
| *Ptgs2* | ENSMUSG00000032487 | 8 |
| *Nos2* | ENSMUSG00000020826 | 8 |
| *Tlr2* | ENSMUSG00000027995 | 8 |
| *Fcgr3* | ENSMUSG00000059498 | 8 |
| *Mrc1* | ENSMUSG00000026712 | 8 |
| *Arg1* | ENSMUSG00000019987 | 8 |
| *Pparg* | ENSMUSG00000000440 | 8 |
| *Chil3* | ENSMUSG00000040809 | 8 |
| *C1qb* | ENSMUSG00000036905 | 8 |
| *Cndp2* | ENSMUSG00000024644 | 8 |
| *Csf1r* | ENSMUSG00000024621 | 8 |
| *F13a1* | ENSMUSG00000039109 | 8 |
| *Fcgr2b* | ENSMUSG00000026656 | 8 |
| *Ms4a6d* | ENSMUSG00000024679 | 8 |
| *Ms4a7* | ENSMUSG00000024672 | 8 |
| *Msr1* | ENSMUSG00000025044 | 8 |
| *Pf4* | ENSMUSG00000029373 | 5 |
| *Trem2* | ENSMUSG00000023992 | 8 |
| *Arg2* | ENSMUSG00000021125 | 8 |
| *Cd274* | ENSMUSG00000016496 | 8 |
| *Csf3r* | ENSMUSG00000028859 | 8 |
| *Entpd1* | ENSMUSG00000048120 | 8 |
| *Hdc* | ENSMUSG00000027360 | 8 |
| *Lcn2* | ENSMUSG00000026822 | 8 |
| *Plk3* | ENSMUSG00000028680 | 8 |
| *Trem1* | ENSMUSG00000042265 | 8 |
| *Cxcr2* | ENSMUSG00000026180 | 8 |
| *Fpr2* | ENSMUSG00000052270 | 8 |
| *Upp1* | ENSMUSG00000020407 | 8 |
| *Wfdc21* | ENSMUSG00000051748 | 3 |
| *Cd83* | ENSMUSG00000015396 | 8 |
| *Etv3* | ENSMUSG00000003382 | 8 |
| *Ramp3* | ENSMUSG00000041046 | 8 |
| *Tnip3* | ENSMUSG00000044162 | 8 |
| *Ccl9* | ENSMUSG00000019122 | 8 |
| *Il1a* | ENSMUSG00000027399 | 8 |
| *Osm* | ENSMUSG00000058755 | 8 |
| *Rtn4* | ENSMUSG00000020458 | 8 |
| *Tnfsf12* | ENSMUSG00000097328 | 8 |
| *Vegfa* | ENSMUSG00000023951 | 8 |
| *Tnfrsf1a* | ENSMUSG00000030341 | 8 |
| *Tmem119* | ENSMUSG00000054675 | 8 |
| *Clec11a* | ENSMUSG00000004473 | 8 |
| *Comp* | ENSMUSG00000031849 | 8 |
| *Cpz* | ENSMUSG00000036596 | 8 |
| *H19* | ENSMUSG00000000031 | 8 |
| *Matn4* | ENSMUSG00000016995 | 8 |
| *Sfrp4* | ENSMUSG00000021319 | 8 |
| *Col9a2* | ENSMUSG00000028626 | 8 |
| *Aldh1a2* | ENSMUSG00000013584 | 8 |
| *Bmp6* | ENSMUSG00000039004 | 8 |
| *Clcn2* | ENSMUSG00000022843 | 8 |
| *Coch* | ENSMUSG00000020953 | 8 |
| *Crabp2* | ENSMUSG00000004885 | 8 |
| *Gja1* | ENSMUSG00000050953 | 8 |
| *Gjb2* | ENSMUSG00000046352 | 8 |
| *Gjb6* | ENSMUSG00000040055 | 8 |
| *Osr1* | ENSMUSG00000048387 | 8 |
| *Slc6a13* | ENSMUSG00000030108 | 8 |
| *Wnt6* | ENSMUSG00000033227 | 8 |
| *Adamts5* | ENSMUSG00000022894 | 8 |
| *Dpt* | ENSMUSG00000026574 | 8 |
| *Pcolce2* | ENSMUSG00000015354 | 8 |
| *Dpp4* | ENSMUSG00000035000 | 8 |
| *Cxcl13* | ENSMUSG00000023078 | 8 |
| *Ly6c1* | ENSMUSG00000079018 | 8 |
| *Bmp4* | ENSMUSG00000021835 | 8 |
| *Gpnmb* | ENSMUSG00000029816 | 8 |
| *Fbln7* | ENSMUSG00000027386 | 8 |
| *Fmod* | ENSMUSG00000041559 | 8 |
| *Itga6* | ENSMUSG00000027111 | 8 |
| *Itgb4* | ENSMUSG00000020758 | 8 |
| *Krt19* | ENSMUSG00000020911 | 8 |
| *Sfrp5* | ENSMUSG00000018822 | 8 |
| *Itga7* | ENSMUSG00000025348 | 8 |
| *Sox8* | ENSMUSG00000024176 | 8 |
| *Etv1* | ENSMUSG00000004151 | 8 |
| *Sox9* | ENSMUSG00000000567 | 8 |
| *Col11a1* | ENSMUSG00000027966 | 8 |
| *Col26a1* | ENSMUSG00000004415 | 8 |
| *Kng2* | ENSMUSG00000060459 | 8 |
| *Abhd3* | ENSMUSG00000002475 | 8 |
| *Arxes2* | ENSMUSG00000048040 | 8 |
| *Agt* | ENSMUSG00000031980 | 8 |
| *Cadm2* | ENSMUSG00000064115 | 8 |
| *Car2* | ENSMUSG00000027562 | 8 |
| *Cd38* | ENSMUSG00000029084 | 8 |
| *Cryab* | ENSMUSG00000032060 | 8 |
| *Cxcl14* | ENSMUSG00000021508 | 8 |
| *Gfap* | ENSMUSG00000020932 | 8 |
| *Slc1a3* | ENSMUSG00000005360 | 8 |
| *Cdh1* | ENSMUSG00000000303 | 8 |
| *Myoc* | ENSMUSG00000026697 | 8 |
| *Pdgfc* | ENSMUSG00000028019 | 8 |
| *Ltbp2* | ENSMUSG00000002020 | 8 |
| *Fbxo17* | ENSMUSG00000030598 | 8 |
| *Pycr1* | ENSMUSG00000025140 | 8 |
| *Arsi* | ENSMUSG00000036412 | 8 |
| *Matn3* | ENSMUSG00000020583 | 8 |
| *Cdkn2b* | ENSMUSG00000073802 | 8 |
| *Zfp354c* | ENSMUSG00000044807 | 8 |
| *Hebp2* | ENSMUSG00000019853 | 8 |
| *Prss35* | ENSMUSG00000033491 | 8 |
| *Wnt16* | ENSMUSG00000029671 | 8 |
| *Camk4* | ENSMUSG00000038128 | 8 |
| *Serinc2* | ENSMUSG00000023232 | 8 |
| *Prkg2* | ENSMUSG00000029334 | 8 |
| *Ttll5* | ENSMUSG00000012609 | 8 |
| *Igdcc4* | ENSMUSG00000032816 | 8 |
| *Cpa6* | ENSMUSG00000042501 | 8 |
| *Mpped2* | ENSMUSG00000016386 | 8 |
| *Bcl11b* | ENSMUSG00000048251 | 8 |
| *Nell2* | ENSMUSG00000022454 | 8 |
| *Sec16b* | ENSMUSG00000026589 | 8 |
| *Tmeff1* | ENSMUSG00000028347 | 8 |
| *Bnc2* | ENSMUSG00000028487 | 8 |
| *Fbxo36* | ENSMUSG00000073633 | 8 |
| *Dync1i1* | ENSMUSG00000029757 | 8 |
| *Kcnj15* | ENSMUSG00000062609 | 8 |
| *Capn6* | ENSMUSG00000067276 | 8 |
| *Cad* | ENSMUSG00000013629 | 8 |
| *Fhod3* | ENSMUSG00000034295 | 8 |
| *Ccn5* | ENSMUSG00000027656 | 8 |
| *Efna4* | ENSMUSG00000028040 | 8 |
| *Npr3* | ENSMUSG00000022206 | 8 |
| *Pamr1* | ENSMUSG00000027188 | 8 |
| *Col24a1* | ENSMUSG00000028197 | 8 |
| *Serp2* | ENSMUSG00000052584 | 5 |
| *Stk26* | ENSMUSG00000031112 | 8 |
| *Pak3* | ENSMUSG00000031284 | 8 |
| *Adamtsl3* | ENSMUSG00000070469 | 8 |
| *Aff3* | ENSMUSG00000037138 | 8 |
| *Brinp3* | ENSMUSG00000035131 | 8 |
| *Rcor2* | ENSMUSG00000024968 | 8 |
| *Arxes1* | ENSMUSG00000048355 | 8 |
| *Asphd2* | ENSMUSG00000029348 | 8 |
| *Tnfaip8l3* | ENSMUSG00000074345 | 8 |
| *Hs3st3b1* | ENSMUSG00000070407 | 8 |
| *Unc5c* | ENSMUSG00000059921 | 8 |
| *Cxadr* | ENSMUSG00000022865 | 8 |
| *Gjb5* | ENSMUSG00000042357 | 8 |
| *Gpr27* | ENSMUSG00000072875 | 8 |
| *Zfp185* | ENSMUSG00000031351 | 8 |
| *Bmp3* | ENSMUSG00000029335 | 8 |
| *Plekhg4* | ENSMUSG00000014782 | 8 |
| *Farp2* | ENSMUSG00000034066 | 8 |
| *Col9a1* | ENSMUSG00000026147 | 8 |
| *Gpr173* | ENSMUSG00000056679 | 8 |
| *Gzmc* | ENSMUSG00000079186 | 8 |
| *Aqp5* | ENSMUSG00000044217 | 8 |
| *Galnt3* | ENSMUSG00000026994 | 8 |
| *Fat3* | ENSMUSG00000074505 | 8 |
| *Ucn2* | ENSMUSG00000049699 | 8 |
| *Bdkrb1* | ENSMUSG00000041347 | 8 |
| *Rab27b* | ENSMUSG00000024511 | 8 |
| *Stc2* | ENSMUSG00000020303 | 8 |
| *Pank1* | ENSMUSG00000033610 | 8 |
| *Hp* | ENSMUSG00000031722 | 8 |
| *Angptl1* | ENSMUSG00000033544 | 8 |
| *Aqp2* | ENSMUSG00000023013 | 8 |
| *Ccl8* | ENSMUSG00000009185 | 4 |
| *Cd34* | ENSMUSG00000016494 | 8 |
| *Cilp* | ENSMUSG00000042254 | 8 |
| *Clec3b* | ENSMUSG00000025784 | 8 |
| *Galnt16* | ENSMUSG00000021130 | 8 |
| *Mmp11* | ENSMUSG00000000901 | 8 |
| *Sgsm2* | ENSMUSG00000038351 | 8 |
| *Shox2* | ENSMUSG00000027833 | 8 |
| *Sulf2* | ENSMUSG00000006800 | 8 |
| *Thbs4* | ENSMUSG00000021702 | 8 |
| *Tgfb2* | ENSMUSG00000039239 | 8 |
| *Tgfb3* | ENSMUSG00000021253 | 8 |
| *Bmp1* | ENSMUSG00000022098 | 8 |
| *Bmp2* | ENSMUSG00000027358 | 8 |
| *Bmp5* | ENSMUSG00000032179 | 8 |
| *Gdf11* | ENSMUSG00000025352 | 8 |
| *Smad1* | ENSMUSG00000031681 | 8 |
| *Smad3* | ENSMUSG00000032402 | 8 |
| *Smad4* | ENSMUSG00000024515 | 8 |
| *Smad6* | ENSMUSG00000036867 | 8 |
| *Smad9* | ENSMUSG00000027796 | 8 |
| *Mapk10* | ENSMUSG00000046709 | 8 |
| *Mapk12* | ENSMUSG00000022610 | 8 |
| *Mapk13* | ENSMUSG00000004864 | 8 |
| *Map3k7* | ENSMUSG00000028284 | 8 |
| *Fbn2* | ENSMUSG00000024598 | 8 |
| *Lrrc17* | ENSMUSG00000039883 | 8 |
| *Adgrg6* | ENSMUSG00000039116 | 8 |
| *Lhx9* | ENSMUSG00000019230 | 8 |
| *Galnt17* | ENSMUSG00000034040 | 8 |
| *Cyp2f2* | ENSMUSG00000052974 | 8 |
| *Dio3* | ENSMUSG00000075707 | 8 |
| *Emx2* | ENSMUSG00000043969 | 8 |
| *Dlx5* | ENSMUSG00000029755 | 8 |
| *Steap1* | ENSMUSG00000015652 | 8 |
| *Pcsk2* | ENSMUSG00000027419 | 8 |
| *Gpm6a* | ENSMUSG00000031517 | 8 |
| *Penk* | ENSMUSG00000045573 | 8 |
| *Pitx1* | ENSMUSG00000021506 | 8 |
| *Corin* | ENSMUSG00000005220 | 8 |
| *Spock3* | ENSMUSG00000054162 | 8 |
| *Snhg11* | ENSMUSG00000044349 | 8 |
| *Alx4* | ENSMUSG00000040310 | 8 |
| *Tox3* | ENSMUSG00000043668 | 8 |
| *Zcchc18* | ENSMUSG00000031428 | 8 |
| *Ntn5* | ENSMUSG00000070564 | 8 |
| *Plxdc1* | ENSMUSG00000017417 | 8 |
| *Pcdh10* | ENSMUSG00000049100 | 8 |
| *Adam33* | ENSMUSG00000027318 | 8 |
| *Maob* | ENSMUSG00000040147 | 8 |
| *Tmem132c* | ENSMUSG00000034324 | 8 |
| *Gna14* | ENSMUSG00000024697 | 8 |
| *Tbx4* | ENSMUSG00000000094 | 8 |
| *Ccdc74a* | ENSMUSG00000041617 | 8 |
| *Aff2* | ENSMUSG00000031189 | 8 |
| *Zic2* | ENSMUSG00000061524 | 8 |
| *Chd5* | ENSMUSG00000005045 | 8 |
| *Emx2os* | ENSMUSG00000087095 | 8 |
| *Ackr4* | ENSMUSG00000079355 | 8 |
| *Lamc3* | ENSMUSG00000026840 | 8 |
| *Scube1* | ENSMUSG00000016763 | 8 |
| *Kcp* | ENSMUSG00000059022 | 8 |
| *Chodl* | ENSMUSG00000022860 | 8 |
| *Kera* | ENSMUSG00000019932 | 8 |
| *Foxf2* | ENSMUSG00000038402 | 8 |
| *Adh1* | ENSMUSG00000074207 | 8 |
| *Nlgn1* | ENSMUSG00000063887 | 8 |
| *Klhl32* | ENSMUSG00000040387 | 8 |
| *Ccdc136* | ENSMUSG00000029769 | 8 |
| *Cntfr* | ENSMUSG00000028444 | 8 |
| *Rab40b* | ENSMUSG00000025170 | 8 |
| *Negr1* | ENSMUSG00000040037 | 8 |
| *Hs3st6* | ENSMUSG00000039628 | 8 |
| *Tbxa2r* | ENSMUSG00000034881 | 8 |
| *Dbx2* | ENSMUSG00000045608 | 8 |
| *Gfra2* | ENSMUSG00000022103 | 8 |
| *Rab30* | ENSMUSG00000030643 | 8 |
| *Dipk1b* | ENSMUSG00000036186 | 8 |
| *Hmgcs2* | ENSMUSG00000027875 | 8 |
| *Fez1* | ENSMUSG00000032118 | 8 |
| *Csmd3* | ENSMUSG00000022311 | 8 |
| *Dpep1* | ENSMUSG00000019278 | 8 |
| *Tle2* | ENSMUSG00000034771 | 8 |
| *Sema3b* | ENSMUSG00000057969 | 8 |
| *Zic5* | ENSMUSG00000041703 | 8 |
| *Focad* | ENSMUSG00000038368 | 8 |
| *Sall1* | ENSMUSG00000031665 | 8 |
| *Plppr3* | ENSMUSG00000035835 | 8 |
| *Hoxd13* | ENSMUSG00000001819 | 8 |
| *Gaa* | ENSMUSG00000025579 | 8 |
| *Galnt9* | ENSMUSG00000033316 | 8 |
| *Ubtd2* | ENSMUSG00000044949 | 7 |
| *Tmem26* | ENSMUSG00000060044 | 8 |
| *Il17d* | ENSMUSG00000050222 | 6 |
| *Mrc2* | ENSMUSG00000020695 | 8 |
| *Arsj* | ENSMUSG00000046561 | 8 |
| *Hoxa11* | ENSMUSG00000038210 | 8 |
| *Sh3rf3* | ENSMUSG00000037990 | 8 |
| *Omd* | ENSMUSG00000048368 | 8 |
| *Cpxm2* | ENSMUSG00000030862 | 8 |
| *Tnn* | ENSMUSG00000026725 | 8 |
| *Ptpru* | ENSMUSG00000028909 | 8 |
| *Hmgcll1* | ENSMUSG00000007908 | 8 |
| *Ror2* | ENSMUSG00000021464 | 8 |
| *Ptgfr* | ENSMUSG00000028036 | 8 |
| *Tspan11* | ENSMUSG00000030351 | 8 |
| *Has1* | ENSMUSG00000003665 | 8 |
| *Wnt5b* | ENSMUSG00000030170 | 8 |
| *Mmp23* | ENSMUSG00000029061 | 8 |
| *Hoxa13* | ENSMUSG00000038203 | 8 |
| *Ssc5d* | ENSMUSG00000035279 | 8 |
| *Pcdh18* | ENSMUSG00000037892 | 8 |
| *C1qtnf2* | ENSMUSG00000046491 | 8 |
| *Tcea3* | ENSMUSG00000001604 | 8 |
| *Pcdh17* | ENSMUSG00000035566 | 8 |
| *Cys1* | ENSMUSG00000062563 | 8 |
| *Epha5* | ENSMUSG00000029245 | 8 |
| *Ntrk3* | ENSMUSG00000059146 | 8 |
| *Mc5r* | ENSMUSG00000007480 | 8 |
| *Ereg* | ENSMUSG00000029377 | 8 |
| *Lif* | ENSMUSG00000034394 | 8 |
| *Rspo3* | ENSMUSG00000019880 | 8 |
| *Sema3d* | ENSMUSG00000040254 | 8 |
| *Col2a1* | ENSMUSG00000022483 | 8 |
| *Cxcr4* | ENSMUSG00000045382 | 8 |
| *Ppp1r1b* | ENSMUSG00000061718 | 8 |
| *Col11a2* | ENSMUSG00000024330 | 8 |
| *Col9a3* | ENSMUSG00000027570 | 8 |
| *Smpd3* | ENSMUSG00000031906 | 8 |
| *Cenpc1* | ENSMUSG00000029253 | 8 |
| *Fzd9* | ENSMUSG00000049551 | 8 |
| *Chrdl1* | ENSMUSG00000031283 | 8 |
| *Mmp3* | ENSMUSG00000043613 | 8 |
| *Gldn* | ENSMUSG00000046167 | 8 |
| *Itgbl1* | ENSMUSG00000032925 | 8 |
| *Crabp1* | ENSMUSG00000032291 | 6 |
| *Masp1* | ENSMUSG00000022887 | 8 |
| *Ostn* | ENSMUSG00000052276 | 8 |
| *Cybrd1* | ENSMUSG00000027015 | 8 |
| *Gdnf* | ENSMUSG00000022144 | 8 |
| *Il1rl1* | ENSMUSG00000026069 | 8 |
| *Mkx* | ENSMUSG00000061013 | 8 |
| *Ndnf* | ENSMUSG00000049001 | 8 |
| *Scx* | ENSMUSG00000034161 | 7 |
| *Nova1* | ENSMUSG00000021047 | 8 |
| *Adgrd1* | ENSMUSG00000044017 | 8 |
| *Il33* | ENSMUSG00000024810 | 8 |
| *Lrrn4cl* | ENSMUSG00000071656 | 8 |
| *Osr2* | ENSMUSG00000022330 | 8 |
| *Grem1* | ENSMUSG00000074934 | 8 |
| *Fam180a* | ENSMUSG00000047420 | 8 |
| *Car3* | ENSMUSG00000027559 | 8 |
| *Slc36a2* | ENSMUSG00000020264 | 8 |
| *Phex* | ENSMUSG00000057457 | 8 |
| *Panx3* | ENSMUSG00000011118 | 8 |
| *Wnt10b* | ENSMUSG00000022996 | 8 |
| *Bmp8a* | ENSMUSG00000032726 | 8 |
| *Ube2ql1* | ENSMUSG00000052981 | 8 |
| *Bglap3* | ENSMUSG00000074489 | 8 |
| *Six4* | ENSMUSG00000034460 | 8 |
| *Bglap2* | ENSMUSG00000074486 | 6 |
| *Ihh* | ENSMUSG00000006538 | 8 |
| *Slc38a3* | ENSMUSG00000010064 | 8 |
| *Sim2* | ENSMUSG00000062713 | 8 |
| *Slc13a5* | ENSMUSG00000020805 | 8 |
| *Foxd1* | ENSMUSG00000078302 | 8 |
| *Shisa3* | ENSMUSG00000050010 | 8 |
| *Gdf10* | ENSMUSG00000021943 | 8 |
| *Igf1r* | ENSMUSG00000005533 | 8 |
| *Fgf2* | ENSMUSG00000037225 | 8 |
| *Ror1* | ENSMUSG00000035305 | 8 |
| *Fgfr3* | ENSMUSG00000054252 | 8 |
| *Wnt5a* | ENSMUSG00000021994 | 8 |
| *Tjp1* | ENSMUSG00000030516 | 8 |
| *Tjp2* | ENSMUSG00000024812 | 8 |

| | | |
|---|---|---|
| *Retnlg* | ENSMUSG00000022651 | 7 |
| *Lrg1* | ENSMUSG00000037095 | 8 |
| *Cxcl3* | ENSMUSG00000029379 | 8 |
| *Bglap* | ENSMUSG00000074483 | 5 |
| *Cd300lf* | ENSMUSG00000047798 | 8 |
| *C1qa* | ENSMUSG00000036887 | 8 |
| *Cspg4* | ENSMUSG00000032911 | 8 |
| *Shh* | ENSMUSG00000002633 | 8 |
| *Irx4* | ENSMUSG00000021604 | 8 |
| *Irx1* | ENSMUSG00000060969 | 8 |
| *Sox21* | ENSMUSG00000061517 | 8 |
| *Epgn* | ENSMUSG00000035020 | 8 |
| *Fam110c* | ENSMUSG00000036136 | 8 |
| *Edn2* | ENSMUSG00000028635 | 8 |
| *Lypd5* | ENSMUSG00000030484 | 8 |
| *Sdr9c7* | ENSMUSG00000040127 | 8 |
| *Ocstamp* | ENSMUSG00000027670 | 8 |
| *Oscar* | ENSMUSG00000054594 | 8 |
| *Cyp1a1* | ENSMUSG00000032315 | 8 |
| *Chp2* | ENSMUSG00000030865 | 8 |
| *Gkn3* | ENSMUSG00000030048 | 8 |
| *Pcp4* | ENSMUSG00000090223 | 8 |
| *P2rx1* | ENSMUSG00000020787 | 8 |
| *Dlk1* | ENSMUSG00000040856 | 8 |
| *Kcna1* | ENSMUSG00000047976 | 8 |
| *Fgf10* | ENSMUSG00000021732 | 8 |
| *Cfh* | ENSMUSG00000026365 | 8 |
| *Efna5* | ENSMUSG00000048915 | 8 |
| *Efnb1* | ENSMUSG00000031217 | 8 |
| *Efnb2* | ENSMUSG00000001300 | 8 |
| *Fgf7* | ENSMUSG00000027208 | 8 |
| *Fstl1* | ENSMUSG00000022816 | 8 |
| *Gas6* | ENSMUSG00000031451 | 8 |
| *Inhba* | ENSMUSG00000041324 | 8 |
| *Mdk* | ENSMUSG00000027239 | 8 |
| *Nrg1* | ENSMUSG00000062991 | 8 |
| *Ntn1* | ENSMUSG00000020902 | 8 |
| *Sema6d* | ENSMUSG00000027200 | 8 |
| *Vegfc* | ENSMUSG00000031520 | 8 |
| *Vegfd* | ENSMUSG00000031380 | 8 |
| *Trp63* | ENSMUSG00000022510 | 8 |
| *Itgb1* | ENSMUSG00000025809 | 8 |
| *Cdh3* | ENSMUSG00000061048 | 8 |
| *Acta2* | ENSMUSG00000035783 | 8 |
| *Fn1* | ENSMUSG00000026193 | 8 |
| *Tnc* | ENSMUSG00000028364 | 8 |
| *Lef1* | ENSMUSG00000027985 | 8 |
| *Nog* | ENSMUSG00000048616 | 8 |
| *Tbx18* | ENSMUSG00000032419 | 8 |
| *Krt16* | ENSMUSG00000053797 | 8 |
| *Krt1* | ENSMUSG00000046834 | 8 |
| *Lor* | ENSMUSG00000043165 | 8 |
| *Flg* | ENSMUSG00000102439 | 8 |
| *Mmp2* | ENSMUSG00000031740 | 8 |
| *Thy1* | ENSMUSG00000032011 | 8 |
