## Supplementary material for "The nail mesenchyme creates a regeneration-specific ligand environment that orchestrates mammalian regeneration versus fibrotic wound healing": Table_3

**Table 3. Ligands expressed in the Early Nail Mesenchyme or Nascent Blastema.** Shown are the ligands expressed by the early nail mesenchyme (cluster 3) or nascent blastema (cluster 7) from the merged scRNASeq dataset shown in Figure 1E. Ligands were identified using a previously published database (see Methods) and were considered to be expressed if the mRNA was detected in at least 5% of the cells within the cluster. The gene abundance is shown for each ligand.

**Threshold:**
>5% expression

| Cluster 3 - Early Nail Mesenchyme | |
|---|---|
| **Ligand** | **Gene Abundance (%)** |
| Agrn | 5.72 |
| Angpt1 | 14.26 |
| Artn | 5.04 |
| Bmp2 | 34.17 |
| Bmp4 | 37.86 |
| Bmp5 | 53.96 |
| Bmp7 | 13.03 |
| Cck | 16.53 |
| Ccl2 | 9.47 |
| Ccl25 | 8.97 |
| Ccl7 | 6.02 |
| Clcf1 | 7.38 |
| Csf1 | 20.59 |
| Cx3cl1 | 14.57 |
| Cxcl1 | 14.57 |
| Cxcl12 | 57.10 |
| Cxcl2 | 5.29 |
| Cxcl5 | 18.25 |
| Eda | 16.96 |
| Efna1 | 12.17 |
| Efna2 | 5.65 |
| Efna5 | 52.98 |
| Efnb1 | 26.06 |
| Efnb2 | 40.50 |
| Fgf10 | 32.45 |
| Fgf18 | 8.24 |
| Fgf2 | 30.61 |
| Fgf7 | 31.78 |
| Fstl1 | 93.61 |
| Gas6 | 35.46 |
| Gdf10 | 6.27 |
| Gdf6 | 11.12 |
| Hgf | 11.62 |
| Igf1 | 19.85 |
| Igf2 | 51.38 |
| Il11 | 5.35 |
| Il15 | 6.15 |
| Inhba | 23.42 |
| Inhbb | 22.68 |
| Jag1 | 14.69 |
| Mdk | 57.96 |
| Mif | 30.55 |
| Ngf | 5.78 |
| Nrg1 | 28.21 |
| Nrg2 | 43.33 |
| Nrg3 | 35.83 |
| Ntn1 | 21.27 |
| Pdgfa | 6.82 |
| Pdgfc | 8.36 |
| Pdgfd | 13.09 |
| Pgf | 11.92 |
| Ptn | 14.51 |
| Rgma | 8.17 |
| Rspo1 | 43.27 |
| Rspo2 | 11.99 |
| Rspo3 | 47.51 |
| Rspo4 | 48.25 |
| Rtn4 | 59.68 |
| Sema3a | 51.87 |
| Sema3b | 9.47 |
| Sema3c | 8.91 |
| Sema3f | 5.10 |
| Sema4b | 6.76 |
| Sema4c | 9.47 |
| Sema5a | 8.54 |
| Sema6a | 12.85 |
| Sema6d | 22.19 |
| Slit2 | 19.98 |
| Slit3 | 10.14 |
| Tgfb1 | 11.62 |
| Tgfb2 | 41.92 |
| Tgfb3 | 27.17 |
| Tnfsf11 | 9.53 |
| Tnfsf12 | 19.11 |
| Vegfa | 38.17 |
| Vegfb | 8.91 |
| Vegfc | 26.98 |
| Vegfd | 7.25 |
| Wnt11 | 10.02 |
| Wnt5a | 65.89 |

| Cluster 7 - Nascent Blastema | |
|---|---|
| **Ligand** | **Gene Abundance (%)** |
| Angpt1 | 33.70 |
| Angpt2 | 7.18 |
| Bmp2 | 13.63 |
| Bmp4 | 5.23 |
| Bmp5 | 55.11 |
| Ccl11 | 5.47 |
| Ccl19 | 5.72 |
| Ccl2 | 21.05 |
| Ccl25 | 5.96 |
| Ccl7 | 14.72 |
| Clcf1 | 11.31 |
| Csf1 | 38.56 |
| Cxcl1 | 31.14 |
| Cxcl12 | 46.23 |
| Cxcl16 | 13.26 |
| Cxcl2 | 10.46 |
| Cxcl5 | 18.98 |
| Eda | 23.60 |
| Efna5 | 41.61 |
| Efnb1 | 19.46 |
| Ereg | 5.60 |
| Fgf10 | 16.67 |
| Fgf2 | 9.12 |
| Fgf7 | 54.87 |
| Fstl1 | 85.40 |
| Gas6 | 20.44 |
| Gdf10 | 13.26 |
| Hbegf | 13.38 |
| Hgf | 16.67 |
| Igf1 | 47.81 |
| Il11 | 25.30 |
| Il1b | 8.15 |
| Il6 | 14.84 |
| Inhba | 48.42 |
| Jag1 | 19.59 |
| Kng1 | 5.72 |
| Lif | 8.39 |
| Mdk | 26.64 |
| Mif | 64.11 |
| Ngf | 16.06 |
| Npy | 5.84 |
| Nrg1 | 6.20 |
| Ntn1 | 7.06 |
| Pdgfa | 33.45 |
| Pdgfc | 32.48 |
| Pdgfd | 13.02 |
| Ptn | 17.15 |
| Rspo3 | 24.82 |
| Rtn4 | 63.75 |
| Sema3c | 6.08 |
| Sema4c | 7.66 |
| Sema5a | 17.88 |
| Sema6d | 12.29 |
| Sema7a | 18.13 |
| Slit2 | 67.27 |
| Slit3 | 42.58 |
| Tgfb1 | 49.64 |
| Tgfb2 | 24.82 |
| Tgfb3 | 8.64 |
| Tnfsf11 | 60.83 |
| Tnfsf12 | 8.88 |
| Vegfa | 27.74 |
| Vegfc | 7.54 |
| Wnt4 | 16.30 |
| Wnt5a | 46.47 |

| Common Ligands |
|---|
| Angpt1 |
| Bmp2 |
| Bmp4 |
| Bmp5 |
| Ccl2 |
| Ccl25 |
| Ccl7 |
| Clcf1 |
| Csf1 |
| Cxcl1 |
| Cxcl12 |
| Cxcl2 |
| Cxcl5 |
| Eda |
| Efna5 |
| Efnb1 |
| Fgf10 |
| Fgf2 |
| Fgf7 |
| Fstl1 |
| Gas6 |
| Gdf10 |
| Hgf |
| Igf1 |
| Il11 |
| Inhba |
| Jag1 |
| Mdk |
| Mif |
| Ngf |
| Nrg1 |
| Ntn1 |
| Pdgfa |
| Pdgfc |
| Pdgfd |
| Ptn |
| Rspo3 |
| Rtn4 |
| Sema3c |
| Sema4c |
| Sema5a |
| Sema6d |
| Slit2 |
| Slit3 |
| Tgfb1 |
| Tgfb2 |
| Tgfb3 |
| Tnfsf11 |
| Tnfsf12 |
| Vegfa |
| Vegfc |
| Wnt5a |
