## Supplementary material for "The nail mesenchyme creates a regeneration-specific ligand environment that orchestrates mammalian regeneration versus fibrotic wound healing": Table_4

**Table 4. Receptors expressed in the Early Nail Mesenchyme or Nascent Blastema.** Shown are the receptors expressed by the early nail mesenchyme (cluster 3) or nascent blastema (cluster 7) from the merged scRNASeq dataset shown in Figure 1E. Receptors were identified using a previously published database (see Methods) and were considered to be expressed if the mRNA was detected in at least 20% of the cells within the cluster. The gene abundance is shown for each ligand.

| **Threshold:** |
|---|
| >20% expression |

| Cluster 3 - Early Nail Mesenchyme | |
|---|---|
| **Receptor** | **Gene Abundance (%)** |
| Calcrl | 44.87 |
| Ntrk2 | 72.16 |
| Bmpr1a | 32.08 |
| Bmpr2 | 44.93 |
| Acvr1 | 45.79 |
| Il6st | 48.31 |
| Ghr | 66.87 |
| Smo | 24.89 |
| Lrp6 | 35.28 |
| Notch2 | 21.76 |
| Ednra | 80.27 |
| Epha5 | 53.96 |
| Epha7 | 51.38 |
| Epha4 | 22.07 |
| F2r | 30.92 |
| Fgfr1 | 60.54 |
| Fgfr2 | 61.03 |
| Nrp1 | 47.02 |
| Axl | 33.56 |
| Tgfbr2 | 28.95 |
| Tgfbr1 | 27.84 |
| Gfra1 | 23.42 |
| Ifnar2 | 32.33 |
| Ifngr1 | 20.28 |
| Igf1r | 56.05 |
| Il13ra1 | 29.01 |
| Il1r1 | 29.38 |
| Tgfbr3 | 46.71 |
| Tnfrsf1a | 44.38 |
| Lrp1 | 94.71 |
| Ar | 28.15 |
| Gabbr1 | 21.76 |
| Thra | 48.00 |
| Neo1 | 42.84 |
| Osmr | 48.43 |
| Pdgfra | 82.36 |
| Pdgfrb | 39.58 |
| Nrp2 | 44.74 |
| Pth1r | 24.77 |
| Plxna2 | 27.47 |
| Robo1 | 53.17 |
| Robo2 | 75.78 |
| Ryk | 36.82 |
| Sfrp1 | 34.23 |
| Sfrp2 | 85.99 |
| Fzd1 | 46.28 |
| Fzd2 | 25.51 |
| Ror1 | 33.93 |
| Ror2 | 32.82 |

| Cluster 7 - Nascent Blastema | |
|---|---|
| **Receptor** | **Gene Abundance (%)** |
| Egfr | 46.96 |
| Bmpr1a | 25.30 |
| Bmpr2 | 38.08 |
| Acvr1 | 36.01 |
| Il6st | 45.62 |
| Lifr | 33.09 |
| Ghr | 32.12 |
| Ptch1 | 21.41 |
| Lrp6 | 23.11 |
| Notch2 | 34.91 |
| Fgfr1 | 69.95 |
| Fgfr2 | 21.05 |
| Nrp1 | 36.98 |
| Axl | 22.02 |
| Tgfbr1 | 27.37 |
| Ifnar2 | 37.23 |
| Ifngr1 | 37.23 |
| Ifngr2 | 27.37 |
| Igf1r | 29.20 |
| Igf2r | 22.51 |
| Il13ra1 | 31.39 |
| Il1r1 | 40.39 |
| Tgfbr3 | 45.01 |
| Tnfrsf1a | 25.67 |
| Lrp1 | 84.18 |
| Neo1 | 29.68 |
| Unc5b | 33.45 |
| Osmr | 23.84 |
| Pdgfra | 44.28 |
| Pdgfrb | 56.93 |
| Nrp2 | 38.56 |
| Pth1r | 32.97 |
| Plxna2 | 33.82 |
| Robo1 | 21.53 |
| Tnfrsf12a | 46.11 |
| Tnfrsf21 | 24.94 |
| Ryk | 57.66 |
| Sfrp1 | 32.73 |
| Fzd1 | 30.41 |

| **Common Receptors** |
|---|
| Bmpr1a |
| Bmpr2 |
| Acvr1 |
| Il6st |
| Ghr |
| Lrp6 |
| Notch2 |
| Fgfr1 |
| Fgfr2 |
| Nrp1 |
| Axl |
| Tgfbr1 |
| Ifnar2 |
| Ifngr1 |
| Igf1r |
| Il13ra1 |
| Il1r1 |
| Tgfbr3 |
| Tnfrsf1a |
| Lrp1 |
| Neo1 |
| Osmr |
| Pdgfra |
| Pdgfrb |
| Nrp2 |
| Pth1r |
| Plxna2 |
| Robo1 |
| Ryk |
| Sfrp1 |
| Fzd1 |
