## Supplementary material for "The nail mesenchyme creates a regeneration-specific ligand environment that orchestrates mammalian regeneration versus fibrotic wound healing": Table_5

**Table 5. Ligand-Receptor Modelling between the Early Nail Mesenchyme and Nascent Blastema.** Predicted ligand and receptor communication models were generated from the merged scRNASeq dataset in Figure 1E using the ligands and ligand receptors defined in Tables S3 and S4, as described in the Methods. Shown are ligand-receptor communication models for autocrine and paracrine interactions between the early nail mesenchyme and the nascent blastema. Gene abundance for the listed ligands and receptors are indicated.

**Thresholds:**
Ligands considered expressed if in >5% of cells in source cluster
Receptors considered expressed if in > 20% of cells in target cluster

| Autocrine Early Nail Mesenchyme Interactions | | | |
|---|---|---|---|
| **Early nail mesenchyme ligands with corresponding receptors in early nail mesenchyme** | | | |
| Ligand | Ligand abundance (%) | Receptor | Receptor abundance (%) |
| Inhbb | 22.68 | Acvr1 | 45.79 |
| Bmp7 | 13.03 | Acvr1 | 45.79 |
| Gas6 | 35.46 | Axl | 33.56 |
| Bmp7 | 13.03 | Bmpr1a | 32.08 |
| Bmp5 | 53.96 | Bmpr1a | 32.08 |
| Bmp4 | 37.86 | Bmpr1a | 32.08 |
| Bmp2 | 34.17 | Bmpr1a | 32.08 |
| Gdf6 | 11.12 | Bmpr1a | 32.08 |
| Fstl1 | 93.61 | Bmpr2 | 44.93 |
| Gdf6 | 11.12 | Bmpr2 | 44.93 |
| Bmp2 | 34.17 | Bmpr2 | 44.93 |
| Bmp7 | 13.03 | Bmpr2 | 44.93 |
| Bmp4 | 37.86 | Bmpr2 | 44.93 |
| Efnb2 | 40.50 | Epha4 | 22.07 |
| Efna5 | 52.98 | Epha5 | 53.96 |
| Efna5 | 52.98 | Epha7 | 51.38 |
| Fgf2 | 30.61 | Fgfr1 | 60.54 |
| Fgf10 | 32.45 | Fgfr2 | 61.03 |
| Fgf2 | 30.61 | Fgfr2 | 61.03 |
| Fgf7 | 31.78 | Fgfr2 | 61.03 |
| Wnt5a | 65.89 | Fzd2 | 25.51 |
| Igf2 | 51.38 | Igf1r | 56.05 |
| Igf1 | 19.85 | Igf1r | 56.05 |
| Clcf1 | 7.38 | Il6st | 48.31 |
| Il11 | 5.35 | Il6st | 48.31 |
| Mdk | 57.96 | Lrp1 | 94.71 |
| Wnt5a | 65.89 | Lrp6 | 35.28 |
| Wnt11 | 10.02 | Lrp6 | 35.28 |
| Rgma | 8.17 | Neo1 | 42.84 |
| Ntn1 | 21.27 | Neo1 | 42.84 |
| Jag1 | 14.69 | Notch2 | 21.76 |
| Vegfd | 7.25 | Nrp1 | 47.02 |
| Sema3a | 51.87 | Nrp1 | 47.02 |
| Pgf | 11.92 | Nrp1 | 47.02 |
| Vegfc | 26.98 | Nrp1 | 47.02 |
| Vegfa | 38.17 | Nrp1 | 47.02 |
| Vegfb | 8.91 | Nrp1 | 47.02 |
| Vegfa | 38.17 | Nrp2 | 44.74 |
| Vegfd | 7.25 | Nrp2 | 44.74 |
| Vegfc | 26.98 | Nrp2 | 44.74 |
| Pgf | 11.92 | Nrp2 | 44.74 |
| Pdgfc | 8.36 | Pdgfra | 82.36 |
| Pdgfa | 6.82 | Pdgfra | 82.36 |
| Pdgfd | 13.09 | Pdgfrb | 39.58 |
| Sema3a | 51.87 | Plxna2 | 27.47 |
| Sema3c | 8.91 | Plxna2 | 27.47 |
| Sema6a | 12.85 | Plxna2 | 27.47 |
| Sema3b | 9.47 | Plxna2 | 27.47 |
| Slit2 | 19.98 | Robo1 | 53.17 |
| Slit3 | 10.14 | Robo1 | 53.17 |
| Slit2 | 19.98 | Robo2 | 75.78 |
| Slit3 | 10.14 | Robo2 | 75.78 |
| Wnt5a | 65.89 | Ror1 | 33.93 |
| Wnt5a | 65.89 | Ror2 | 32.82 |
| Wnt5a | 65.89 | Ryk | 36.82 |
| Gdf10 | 6.27 | Tgfbr1 | 27.84 |
| Tgfb3 | 27.17 | Tgfbr1 | 27.84 |
| Tgfb1 | 11.62 | Tgfbr1 | 27.84 |
| Tgfb2 | 41.92 | Tgfbr1 | 27.84 |
| Tgfb3 | 27.17 | Tgfbr2 | 28.95 |
| Tgfb2 | 41.92 | Tgfbr2 | 28.95 |
| Gdf10 | 6.27 | Tgfbr2 | 28.95 |
| Tgfb1 | 11.62 | Tgfbr2 | 28.95 |
| Tgfb1 | 11.62 | Tgfbr3 | 46.71 |
| Tgfb2 | 41.92 | Tgfbr3 | 46.71 |

| Autocrine Nascent Blastema Interactions | | | |
|---|---|---|---|
| **Nascent blastema ligands with corresponding receptors in nascent blastema** | | | |
| Ligand | Ligand abundance (%) | Receptor | Receptor abundance (%) |
| Gas6 | 20.44 | Axl | 22.02 |
| Bmp4 | 5.23 | Bmpr1a | 25.30 |
| Bmp5 | 55.11 | Bmpr1a | 25.30 |
| Bmp2 | 13.63 | Bmpr1a | 25.30 |
| Fstl1 | 85.40 | Bmpr2 | 38.08 |
| Bmp4 | 5.23 | Bmpr2 | 38.08 |
| Bmp2 | 13.63 | Bmpr2 | 38.08 |
| Ereg | 5.60 | Egfr | 46.96 |
| Hbegf | 13.38 | Egfr | 46.96 |
| Fgf2 | 9.12 | Fgfr1 | 69.95 |
| Fgf2 | 9.12 | Fgfr2 | 21.05 |
| Fgf10 | 16.67 | Fgfr2 | 21.05 |
| Fgf7 | 54.87 | Fgfr2 | 21.05 |
| Igf1 | 47.81 | Igf1r | 29.20 |
| Il1b | 8.15 | Il1r1 | 40.39 |
| Il11 | 25.30 | Il6st | 45.62 |
| Lif | 8.39 | Il6st | 45.62 |
| Il6 | 14.84 | Il6st | 45.62 |
| Clcf1 | 11.31 | Il6st | 45.62 |
| Clcf1 | 11.31 | Lifr | 33.09 |
| Lif | 8.39 | Lifr | 33.09 |
| Mdk | 26.64 | Lrp1 | 84.18 |
| Wnt4 | 16.30 | Lrp6 | 23.11 |
| Wnt5a | 46.47 | Lrp6 | 23.11 |
| Ntn1 | 7.06 | Neo1 | 29.68 |
| Jag1 | 19.59 | Notch2 | 34.91 |
| Vegfa | 27.74 | Nrp1 | 36.98 |
| Vegfc | 7.54 | Nrp1 | 36.98 |
| Vegfc | 7.54 | Nrp2 | 38.56 |
| Vegfa | 27.74 | Nrp2 | 38.56 |
| Pdgfc | 32.48 | Pdgfra | 44.28 |
| Pdgfa | 33.45 | Pdgfra | 44.28 |
| Pdgfd | 13.02 | Pdgfrb | 56.93 |
| Sema3c | 6.08 | Plxna2 | 33.82 |
| Slit3 | 42.58 | Robo1 | 21.53 |
| Slit2 | 67.27 | Robo1 | 21.53 |
| Wnt5a | 46.47 | Ryk | 57.66 |
| Gdf10 | 13.26 | Tgfbr1 | 27.37 |
| Tgfb3 | 8.64 | Tgfbr1 | 27.37 |
| Tgfb1 | 49.64 | Tgfbr1 | 27.37 |
| Tgfb2 | 24.82 | Tgfbr1 | 27.37 |
| Tgfb2 | 24.82 | Tgfbr3 | 45.01 |
| Tgfb1 | 49.64 | Tgfbr3 | 45.01 |
| Tnfsf12 | 8.88 | Tnfrsf12a | 46.11 |
| Ntn1 | 7.06 | Unc5b | 33.45 |

| Paracrine Early Nail Mesenchyme-Nascent Blastema Interactions | | | |
|---|---|---|---|
| **Early nail mesenchyme ligand interactions with receptors in nascent blastema** | | | |
| Ligand | Ligand abundance (%) | Receptor | Receptor abundance (%) |
| Inhbb | 22.68 | Acvr1 | 36.01 |
| Bmp7 | 13.03 | Acvr1 | 36.01 |
| Gas6 | 35.46 | Axl | 22.02 |
| Bmp7 | 13.03 | Bmpr1a | 25.30 |
| Bmp4 | 37.86 | Bmpr1a | 25.30 |
| Bmp5 | 53.96 | Bmpr1a | 25.30 |
| Bmp2 | 34.17 | Bmpr1a | 25.30 |
| Gdf6 | 11.12 | Bmpr1a | 25.30 |
| Bmp4 | 37.86 | Bmpr2 | 38.08 |
| Gdf6 | 11.12 | Bmpr2 | 38.08 |
| Bmp2 | 34.17 | Bmpr2 | 38.08 |
| Bmp7 | 13.03 | Bmpr2 | 38.08 |
| Fstl1 | 93.61 | Bmpr2 | 38.08 |
| Fgf2 | 30.61 | Fgfr1 | 69.95 |
| Fgf7 | 31.78 | Fgfr2 | 21.05 |
| Fgf10 | 32.45 | Fgfr2 | 21.05 |
| Fgf2 | 30.61 | Fgfr2 | 21.05 |
| Igf2 | 51.38 | Igf1r | 29.20 |
| Igf1 | 19.85 | Igf1r | 29.20 |
| Igf2 | 51.38 | Igf2r | 22.51 |
| Clcf1 | 7.38 | Il6st | 45.62 |
| Il11 | 5.35 | Il6st | 45.62 |
| Clcf1 | 7.38 | Lifr | 33.09 |
| Mdk | 57.96 | Lrp1 | 84.18 |
| Wnt5a | 65.89 | Lrp6 | 23.11 |
| Wnt11 | 10.02 | Lrp6 | 23.11 |
| Rgma | 8.17 | Neo1 | 29.68 |
| Ntn1 | 21.27 | Neo1 | 29.68 |
| Jag1 | 14.69 | Notch2 | 34.91 |
| Vegfb | 8.91 | Nrp1 | 36.98 |
| Pgf | 11.92 | Nrp1 | 36.98 |
| Vegfc | 26.98 | Nrp1 | 36.98 |
| Vegfd | 7.25 | Nrp1 | 36.98 |
| Sema3a | 51.87 | Nrp1 | 36.98 |
| Vegfa | 38.17 | Nrp1 | 36.98 |
| Pgf | 11.92 | Nrp2 | 38.56 |
| Vegfc | 26.98 | Nrp2 | 38.56 |
| Vegfa | 38.17 | Nrp2 | 38.56 |
| Vegfd | 7.25 | Nrp2 | 38.56 |
| Pdgfc | 8.36 | Pdgfra | 44.28 |
| Pdgfa | 6.82 | Pdgfra | 44.28 |
| Pdgfd | 13.09 | Pdgfrb | 56.93 |
| Sema3c | 8.91 | Plxna2 | 33.82 |
| Sema3b | 9.47 | Plxna2 | 33.82 |
| Sema6a | 12.85 | Plxna2 | 33.82 |
| Sema3a | 51.87 | Plxna2 | 33.82 |
| Slit3 | 10.14 | Robo1 | 21.53 |
| Slit2 | 19.98 | Robo1 | 21.53 |
| Wnt5a | 65.89 | Ryk | 57.66 |
| Tgfb2 | 41.92 | Tgfbr1 | 27.37 |
| Gdf10 | 6.27 | Tgfbr1 | 27.37 |
| Tgfb3 | 27.17 | Tgfbr1 | 27.37 |
| Tgfb1 | 11.62 | Tgfbr1 | 27.37 |
| Tgfb1 | 11.62 | Tgfbr3 | 45.01 |
| Tgfb2 | 41.92 | Tgfbr3 | 45.01 |
| Tnfsf12 | 19.11 | Tnfrsf12a | 46.11 |
| Ntn1 | 21.27 | Unc5b | 33.45 |

| Paracrine Nascent Blastema-Early Nail Mesenchyme Interactions | | | |
|---|---|---|---|
| **Early nail mesenchyme ligand interactions with receptors in nascent blastema** | | | |
| Ligand | Ligand abundance (%) | Receptor | Receptor abundance (%) |
| Gas6 | 20.44 | Axl | 33.56 |
| Bmp5 | 55.11 | Bmpr1a | 32.08 |
| Bmp4 | 5.23 | Bmpr1a | 32.08 |
| Bmp2 | 13.63 | Bmpr1a | 32.08 |
| Bmp2 | 13.63 | Bmpr2 | 44.93 |
| Fstl1 | 85.40 | Bmpr2 | 44.93 |
| Bmp4 | 5.23 | Bmpr2 | 44.93 |
| Efna5 | 41.61 | Epha5 | 53.96 |
| Efna5 | 41.61 | Epha7 | 51.38 |
| Fgf2 | 9.12 | Fgfr1 | 60.54 |
| Fgf2 | 9.12 | Fgfr2 | 61.03 |
| Fgf10 | 16.67 | Fgfr2 | 61.03 |
| Fgf7 | 54.87 | Fgfr2 | 61.03 |
| Wnt5a | 46.47 | Fzd2 | 25.51 |
| Igf1 | 47.81 | Igf1r | 56.05 |
| Il1b | 8.15 | Il1r1 | 29.38 |
| Clcf1 | 11.31 | Il6st | 48.31 |
| Il6 | 14.84 | Il6st | 48.31 |
| Lif | 8.39 | Il6st | 48.31 |
| Il11 | 25.30 | Il6st | 48.31 |
| Mdk | 26.64 | Lrp1 | 94.71 |
| Wnt5a | 46.47 | Lrp6 | 35.28 |
| Wnt4 | 16.30 | Lrp6 | 35.28 |
| Ntn1 | 7.06 | Neo1 | 42.84 |
| Jag1 | 19.59 | Notch2 | 21.76 |
| Vegfa | 27.74 | Nrp1 | 47.02 |
| Vegfc | 7.54 | Nrp1 | 47.02 |
| Vegfa | 27.74 | Nrp2 | 44.74 |
| Vegfc | 7.54 | Nrp2 | 44.74 |
| Pdgfa | 33.45 | Pdgfra | 82.36 |
| Pdgfc | 32.48 | Pdgfra | 82.36 |
| Pdgfd | 13.02 | Pdgfrb | 39.58 |
| Sema3c | 6.08 | Plxna2 | 27.47 |
| Slit2 | 67.27 | Robo1 | 53.17 |
| Slit3 | 42.58 | Robo1 | 53.17 |
| Slit3 | 42.58 | Robo2 | 75.78 |
| Slit2 | 67.27 | Robo2 | 75.78 |
| Wnt5a | 46.47 | Ror1 | 33.93 |
| Wnt5a | 46.47 | Ror2 | 32.82 |
| Wnt5a | 46.47 | Ryk | 36.82 |
| Tgfb2 | 24.82 | Tgfbr1 | 27.84 |
| Tgfb3 | 8.64 | Tgfbr1 | 27.84 |
| Tgfb1 | 49.64 | Tgfbr1 | 27.84 |
| Gdf10 | 13.26 | Tgfbr1 | 27.84 |
| Tgfb3 | 8.64 | Tgfbr2 | 28.95 |
| Gdf10 | 13.26 | Tgfbr2 | 28.95 |
| Tgfb1 | 49.64 | Tgfbr2 | 28.95 |
| Tgfb2 | 24.82 | Tgfbr2 | 28.95 |
| Tgfb2 | 24.82 | Tgfbr3 | 46.71 |
| Tgfb1 | 49.64 | Tgfbr3 | 46.71 |
