## Supplementary material for "The nail mesenchyme creates a regeneration-specific ligand environment that orchestrates mammalian regeneration versus fibrotic wound healing": Table_6

**Table 6**. ***Shown are Xenium probes differentially detected in a comparison between nascent blastema and perturbation enriched mesenchymal cells*****. (Related to Figure 7)** Shown are Xenium probes differentially detected in a comparison between the nascent blastema and perturbation enriched mesenchymal cell clusters shown in the UMAP of the merged 7 DPA *Lmx1b-DTA*, *Pdgfra-Smad4KO* and control Xenium single cell datasets as shown in Suppl. Figure 6C. Probes were considered to be significantly enriched if the gene had a Bonferroni-adjusted p-value of less than 0.05 (p adj < 0.05) and the gene was expressed in at least 5% of nascent blastema cells. Also shown are the average fold-change (avg. fold-change) and the

| Genes enriched in nascent blastema | | | | | |
|---|---|---|---|---|---|
| **Genes** | **avg. log2 fold-change** | **avg. fold-change** | **% Nascent blastema** | **% Perturbation enriched mesenchymal cells** | **p value adj.** |
| *Crabp1* | 3.304853094 | 9.882342751 | 37.6 | 4.9 | 1.04E-249 |
| *Ednra* | 1.750051327 | 3.363705331 | 19.4 | 5.9 | 3.44E-64 |
| *Gdf6* | 1.599787756 | 3.030987192 | 6.1 | 2 | 2.98E-16 |
| *Fgf10* | 1.595397089 | 3.021776774 | 11.7 | 3.8 | 6.90E-33 |
| *Bmp2* | 1.537974502 | 2.903865238 | 19.2 | 7.2 | 1.20E-48 |
| *Ndnf* | 1.527014989 | 2.88188944 | 8.2 | 2.9 | 4.76E-20 |
| *Pdgfra* | 1.400937528 | 2.64073133 | 41.2 | 16.1 | 1.07E-120 |
| *Sfrp2* | 1.290411278 | 2.445977746 | 8.8 | 3.9 | 8.45E-15 |
| *Lamc3* | 1.225283608 | 2.338014064 | 8.5 | 3.5 | 1.17E-15 |
| *Lhx9* | 1.196575395 | 2.291949719 | 5.5 | 2.4 | 7.21E-09 |
| *Zic2* | 1.148797764 | 2.217290447 | 6.2 | 2.7 | 9.64E-10 |
| *Wnt5a* | 1.115769069 | 2.16710502 | 53.6 | 26.8 | 5.39E-124 |
| *Pdpn* | 1.077137991 | 2.109846427 | 11.4 | 5.4 | 2.78E-17 |
| *Msx1* | 1.064725207 | 2.091771418 | 5.3 | 2.5 | 5.19E-07 |
| *Alx4* | 1.035059425 | 2.049198052 | 9.6 | 4.6 | 7.35E-14 |
| *Dpt* | 0.961560647 | 1.947415389 | 16.5 | 8.4 | 2.16E-22 |
| *Tmem119* | 0.87771228 | 1.837459282 | 35.3 | 19.3 | 8.03E-50 |
| *Gldn* | 0.863532524 | 1.819487984 | 12.5 | 6.9 | 2.74E-13 |
| *Masp1* | 0.810825005 | 1.754214303 | 62 | 35.7 | 2.24E-107 |
| *Mmp3* | 0.752219683 | 1.684382373 | 19.7 | 11.5 | 5.99E-19 |
| *Penk* | 0.703033749 | 1.627924451 | 7 | 3.5 | 1.80E-08 |
| *Bmp5* | 0.674622691 | 1.596179269 | 26.2 | 16.7 | 1.66E-20 |
| *Cntfr* | 0.662337018 | 1.58264427 | 15.4 | 9.2 | 1.29E-12 |
| *Cfh* | 0.660414115 | 1.58053624 | 10.3 | 6.6 | 9.13E-06 |
| *Bmp4* | 0.654830541 | 1.57443101 | 14.3 | 8.1 | 1.80E-13 |
| *Igf2* | 0.653719462 | 1.573218942 | 5.7 | 3.6 | 0.011562089 |
| *Col1a1* | 0.61365991 | 1.530136025 | 98.6 | 83.5 | 1.18E-176 |
| *Pdgfrb* | 0.605407428 | 1.52140835 | 39.8 | 27 | 6.48E-29 |
| *Tnn* | 0.491914467 | 1.406309821 | 80.4 | 63.6 | 6.91E-74 |
| *Col11a1* | 0.302363522 | 1.233163012 | 52 | 43.7 | 6.34E-12 |
